# Within-Individual Organization of the Human Cerebral Cortex: Networks, Global Topography, and Function

**DOI:** 10.1101/2023.08.08.552437

**Authors:** Jingnan Du, Lauren M. DiNicola, Peter A. Angeli, Noam Saadon-Grosman, Wendy Sun, Stephanie Kaiser, Joanna Ladopoulou, Aihuiping Xue, B.T. Thomas Yeo, Mark C. Eldaief, Randy L. Buckner

## Abstract

The human cerebral cortex is populated by specialized regions that are organized into networks. Here we estimated networks using a Multi-Session Hierarchical Bayesian sModel (MS-HBM) applied to intensively sampled within-individual functional MRI (fMRI) data. The network estimation procedure was initially developed and tested in two participants (each scanned 31 times) and then prospectively applied to 15 new participants (each scanned 8 to 11 times). Detailed analysis of the networks revealed a global organization. Locally organized first-order sensory and motor networks were surrounded by spatially adjacent second-order networks that also linked to distant regions. Third-order networks each possessed regions distributed widely throughout association cortex. Moreover, regions of distinct third-order networks displayed side-by-side juxtapositions with a pattern that repeated similarly across multiple cortical zones. We refer to these as Supra-Areal Association Megaclusters (SAAMs). Within each SAAM, two candidate control regions were typically adjacent to three separate domain-specialized regions. Independent task data were analyzed to explore functional response properties. The somatomotor and visual first-order networks responded to body movements and visual stimulation, respectively. A subset of the second-order networks responded to transients in an oddball detection task, consistent with a role in orienting to salient or novel events. The third-order networks, including distinct regions within each SAAM, showed two levels of functional specialization. Regions linked to candidate control networks responded to working memory load across multiple stimulus domains. The remaining regions within each SAAM did not track working memory load but rather dissociated across language, social, and spatial / episodic processing domains. These results support a model of the cerebral cortex in which progressively higher-order networks nest outwards from primary sensory and motor cortices. Within the apex zones of association cortex there is specialization of large-scale networks that divides domain-flexible from domain-specialized regions repeatedly across parietal, temporal, and prefrontal cortices. We discuss implications of these findings including how repeating organizational motifs may emerge during development.

---

The primate cerebral cortex is populated by specialized networks that support sensory, motor and higher-order cognitive and affective functions. Characterizing how the networks and their interconnected regions are organized on the cortical surface began more than a century ago with landmark studies of myelogenetic and architectonic patterns (e.g., Flechsig 1901; Campbell 1905; Brodmann 1909; von Economo and Koskinas 1925; von Bonin and Bailey 1947) and continued with modern systems neuroscience integration of anatomical projection data and insights from study of brain lesions (e.g., Geschwind 1965; Ungerleider and Desimone 1986; Goldman-Rakic 1988; Mesulam 1990; 1998; Van Essen et al. 1992; Pandya et al. 2015; Giarrocco and Averbeck 2023). Over the past decades our laboratory, and the field more broadly, has undertaken data collection efforts and analyses of neuroimaging data with the goal to improve understanding of human network organization and provide non-invasive approaches to measure brain organization for clinical use.

It is beyond the present scope to cover the extensive literature that has evolved, but it is important to interpret the current effort with awareness that new details and revisions emerge incrementally as the methods and data quality progress. Our efforts presented here reflect another step in that progression. The specific challenge in examining the details of network organization in humans is that the methods are indirect and limited, and often noisy within individuals. Despite limitations, advances in structural, diffusion, and functional MRI (fMRI) provide valuable information about human cortical organization, albeit with ambiguities consistent with the complexity of cortical architecture and the low resolution of the techniques. Resting-state functional connectivity MRI (fcMRI), based on measuring spontaneous correlated fluctuations between brain regions, has been especially useful for estimating networks (Biswal et al. 1995; see also Fox and Raichle 2007; Van Dijk et al. 2010; Buckner et al. 2013; Murphy et al. 2013; Smith et al. 2013; Power et al. 2014). Explorations in group-averaged fcMRI data, with sample sizes that range from ten to thousands of participants, reveal network estimates that are consistent across analytical approaches and datasets (e.g., Beckmann et al. 2005; Damoiseaux et al. 2006; Yeo et al. 2011; Power et al. 2011; Doucet et al. 2011; Miller et al. 2016; Glasser et al. 2016). Moreover, estimated networks show similarities to directly observed anatomical projection patterns from tracer injections in the monkey, providing support that they reflect, to a first approximation, anatomically connected networks (Vincent et al. 2007; Buckner et al. 2008; Binder et al. 2009; Margulies et al. 2009; Hutchinson et al. 2012; Ghahremani et al. 2017; Liu et al. 2019; Buckner and Margulies 2019; Du and Buckner 2021). Correspondence is far from perfect and there are unresolved aspects to how indirect human network estimates link to anatomy, a theme that we will return to in the discussion.

A recent advance in the field is to use within-individual estimates of networks without recourse to averaging across participants. Architectonic fields tile the cortical mantle with variability in their exact locations, sizes, and borders between individuals (Rademacher et al 1993; Rajkowska and Goldman-Rakic 1995; Amunts et al. 1999; 2000; 2020; Caspers et al. 2006; Fischl et al. 2008; Henssen et al. 2016; Palomero-Gallagher et al. 2019). Spatial blurring – inherent in group-averaging – impedes the ability to estimate details of network organization. Precision neuroimaging, involving intensive sampling and analysis of data within the individual, preserves idiosyncratic anatomical features.

Within-individual approaches have been the mainstay in human neuroimaging studies of sensory and motor systems (e.g., Sereno et al. 1995; Rao et al. 1995; DeYoe et al. 1996; Engel, Glover and Wandell 1997; Kanwisher, McDermott, and Chun 1997; Epstein and Kanwisher 1998) and emerged later as viable to estimate task-based responses in higher-order association cortex (e.g., Fedorenko et al. 2010; 2012; Blank et al. 2013; Peer et al. 2015; Michalka et al. 2015; Huth et al. 2016). Within-individual precision mapping using fcMRI only became emphasized recently, even though the first report was within individuals (Biswal et al. 1995). Following a landmark demonstration that intensive repeat scanning is possible (Poldrack et al. 2015; Laumann et al. 2015), multiple groups have pursued within-individual characterization of network organization (e.g., Braga and Buckner 2017; Gordon et al. 2017; Braga et al. 2019; 2020; Smith et al. 2021; Somers et al. 2021; Noyce et al. 2022; Gordon et al. 2023; for further discussion see Gratton and Braga 2021; Laumann, Zorumski, and Dosenbach 2023).

Here we continue the investigation of the detailed organization of the cerebral cortex using within-individual approaches. There are multiple goals and methodological innovations that steer this work. First, we employ deep, intensive imaging to boost the signal-to-noise (SNR) within individual participants. Each new participant was scanned on at least 8 separate occasions and often more. Second, we applied a novel Multi-Session Hierarchical Bayesian Model (MS-HBM; Kong et al. 2019) to automatically estimate networks in the intensively sampled participants. Specifically, the number of networks estimated was set at 15 to capture established networks sometimes missed in simpler network parcellations, as will be detailed within the methods. Third, to enable clinical translational research, we developed an empirical method and projected all network estimates from the surface back into the native-space volume of individual participants, as is needed for presurgical planning and neuromodulation. Fourth, inspired by the possibility to chart global spatial relations between networks (e.g., Margulies et al. 2016), we also plotted the resulting network estimates on the fully flattened cortical surface (Van Essen and Maunsell 1980; Fischl et al. 1999). As the results will reveal, there are repeating patterns of spatial juxtapositions among networks that provide insight into their evolutionary and developmental origins. Finally, we collected and examined task data within the same intensively sampled participants to test whether within-individual network estimates predict functional response patterns and also to explore between network functional dissociations.

The raw data and our provisional network parcellations generated through this research effort are provided to the community as an open resource

## Methods

### Overview

We sought to estimate networks within individuals with high precision. The analyses proceeded in three stages: (1) a refinement stage established the methods for estimating networks, (2) an implementation stage applied the methods prospectively to 15 new participants, and (3) a functional testing stage explored functional response properties and dissociations between networks.

In the refinement stage, previously reported datasets (N = 2; Braga et al. 2019; Xue et al. 2021) were analyzed to establish a novel MS-HBM network estimate that incorporated priors for 15 distinct networks (as contrast to 10 networks used in earlier work). Each of the participants performed 31 independent MRI sessions allowing considerable data to test for within-individual reliability.

In the implementation stage, the 15-network MS-HBM model was prospectively applied to 15 new participants that were each scanned 8-11 times. The model was estimated for each participant in a fully automated fashion, and the networks were confirmed using model-free seed-region based functional connectivity. Following network estimation, the overlap and variability of each network across individuals were examined.

In the final functional testing stage, an extensive battery of tasks was administered and analyzed within each individual to explore whether the estimated networks predicted functional responses.

### Participants

Seventeen native English-speaking volunteers participated for payment. History of a neurologic or psychiatric illness was an exclusion. Participants provided informed consent using protocols approved by the Institutional Review Board of Harvard University. For the refinement stage data, 2 right-handed adult women ages 22 - 23 yr participated (data previously reported in Braga et al. 2019 and Xue et al. 2021). The refinement stage data participants are labeled S1 and S2 to match Xue et al. (2021). For the implementation stage data, 15 right-handed adults ages 18 – 34 yr participated (mean = 22.1 yr, SD = 3.9 yr, 9 women). Participants came from diverse racial and ethnic backgrounds (9 of the 17 individuals self-reported as non-white and / or Hispanic). A subset of the participants contributing implementation stage data also enrolled in a study of motor movement mapping (Saadon-Grosman et al. 2022). The implementation stage participants are labeled P1 to P15.

### MRI Data Acquisition

Data were acquired at the Harvard Center for Brain Science using a 3T Siemens Prisma-fit MRI scanner. A 64-channel phased-array head-neck coil (Siemens Healthcare, Erlangen, Germany) was used in the refinement stage and for a subset of motor task sessions in the implementation stage. A 32**-**channel phased-array head coil (Siemens Healthcare, Erlangen, Germany) was used to acquire all other data in the implementation stage. For functional neuroimaging, the differences between these two coils are minimal and the data were treated as comparable. Foam and inflated padding mitigated head motion. Participants were instructed to remain still and alert and to look at a rear-projected display through a mirror attached to the head coil. The display had a resolution of 1280 x 1024 pixels and screen width of 43 cm, resulting in an effective viewing distance of 104 cm (54 pixels per visual degree). Eyes were video recorded using an Eyelink 1000 Plus with Long-Range Mount (SR Research, Ottawa, Ontario, Canada), and alertness was scored during each functional run. MRI data quality was monitored during the scan using Framewise Integrated Real-time MRI Monitoring (FIRMM; Dosenbach et al. 2017).

#### Refinement Stage Data

Each participant (S1 and S2) was scanned across 31 MRI sessions over 28-40 wks with no sessions on consecutive days. Each session involved multiple resting-state fixation runs to be used for functional connectivity analysis, for a total of 63 functional MRI (fMRI) runs obtained for each individual. fMRI data were acquired using blood oxygenation level-dependent (BOLD) contrast (Kwong et al. 1992; Ogawa et al. 1992). A custom multiband gradient-echo echo-planar pulse sequence provided by the Center for Magnetic Resonance Research (CMRR) at the University of Minnesota was used (Xu et al. 2012; Van Essen et al. 2013; see also Setsompop et al. 2012): voxel size = 2.4 mm, repetition time (TR) = 1,000 ms, echo time (TE) = 32.6 ms, flip-angle = 64°, matrix 88 x 88 x 65, anterior-to-posterior (AP) phase encoding, multislice 5x acceleration, fully covering the cerebrum and cerebellum. Signal dropout was minimized by automatically (van der Kouwe et al. 2005) selecting a slice 25° from the anterior-posterior commissural plane toward the coronal plane (Weiskopf et al. 2006; Mennes et al. 2014). Each run lasted 7 min 2 sec (422 frames with the first 12 frames removed for T1 equilibration). A dual-gradient-echo B0 fieldmap was acquired to correct for spatial distortions: TE = 4.45 and 6.91 ms with slice prescription / spatial resolution matched to the BOLD sequence. During BOLD scanning, participants fixated a centrally presented plus sign (black on a gray background). The scanner room was illuminated.

A rapid T1w structural scan was obtained using a multi-echo magnetization prepared rapid acquisition gradient echo (ME-MPRAGE) three-dimensional sequence (van der Kouwe et al. 2008): voxel size = 1.2 mm, TR = 2,200 ms, TE = 1.57, 3.39, 5.21, 7.03 ms, TI = 1,100 ms, flip-angle = 7°, matrix 192 x 192 x 176, in-plane generalized auto-calibrating partial parallel acquisition (GRAPPA) acceleration = 4.

#### Implementation Stage Data

Each participant (P1 to P15) was scanned across 8-11 sessions most often over 6 to 10 wks. A few participants had longer gaps between the first and last MRI sessions up to one year. Each session involved multiple fMRI runs to be used for functional connectivity analysis, for a total of 17 to 24 resting-state fixation runs obtained for each individual. BOLD acquisition parameters were similar to the refinement stage data: voxel size = 2.4 mm, TR = 1,000 ms, TE = 33.0 ms, flip-angle = 64°, matrix 92 × 92 × 65 (FOV = 221 × 221), 65 slices covering the full cerebrum and cerebellum. Each resting-state fixation run again lasted 7 min 2 sec (422 frames with the first 12 frames removed for T1 equilibration). Dual-gradient-echo B0 fieldmaps were also acquired with parameters matched to the refinement stage. The first two sessions of P12 were acquired in a different FOV (211 × 211); therefore, the matrix for both BOLD runs and field maps was: 88 × 88 × 65 and BOLD TE = 32.6 ms, matching S1 and S2. The change in FOV did not affect the quality of registration or impact the analyses in any way we could detect.

High-resolution T1w and T2w scans were acquired for the implementation stage data based on the Human Connectome Project (HCP; Harms et al. 2018). T1w MPRAGE parameters: voxel size = 0.8 mm, TR = 2,500 ms, TE = 1.81, 3.60, 5.39, and 7.18 ms, TI = 1,000 ms, flip-angle = 8°, matrix 320 × 320 × 208, 144, in-plane GRAPPA acceleration = 2. T2w sampling perfection with application-optimized contrasts using different flip angle evolution sequence (SPACE) parameters: voxel size = 0.8 mm, TR=3,200 ms, TE=564 ms, 208 slices, matrix=320 x 300 x 208, in-plane GRAPPA acceleration = 2. Rapid T1w structural scans were also obtained as backup using the refinement stage sequence but with matrix 192 x 192 x 144.

#### Functional Testing Stage Data

To explore functional response properties, extensive task-based BOLD fMRI data were collected on participants P1 to P15. Task runs used the same sequence as the resting-state fixation runs, ensuring the estimated networks would be spatially aligned to the task-based data. Details of the task designs, stimuli and run structure are described below under *Task Paradigms*.

### Exclusion Criteria and Quality Control

Each BOLD fMRI run was examined for quality. Exclusion criteria generally consisted of the parameters reported in Xue et al. (2021) including: 1) maximum absolute motion > 1.8 mm and 2) slice-based SNR < 130. Runs with SNR > 100 but also SNR < 130 were retained if motion and visual inspection indicated adequate quality. For the functional testing stage data, the maximum absolute motion for the Episodic Projection task was > 2.5 mm given their long duration. One borderline motor run (P2) was included with motion of 1.9 mm as the motion was largely due to a linear drift. For the refinement stage, usable resting-state runs were 62 (S1) and 61 (S2) runs. For the implementation stage, usable resting-state runs ranged from 15 (P11) to 24 (P12) runs. For the functional testing stage, usable task runs ranged from 18 (P5) to 70 (P12) runs (see Table 1). All data exclusions were finalized prior to functional connectivity and task response analyses.

**Table 1.** Functional data analyzed for each participant.

| Participant | Fixation | Motor | Visual | Oddball | N-Back | Theory-of-Mind | Sentence Processing | Episodic Projection |
| --- | --- | --- | --- | --- | --- | --- | --- | --- |
| S1 | **62(63)** | - | - | - | - | - | - | - |
| S2 | **61(63)** | - | - | - | - | - | - | - |
| P1 | **17(17)** | *0(0)* | **5(5)** | **5(5)** | **8(8)** | **8(8)** | **6(6)** | **10(10)** |
| P2 | **16(17)** | **11(12)** | **5(5)** | **5(5)** | **8(8)** | **8(8)** | **6(6)** | **10(10)** |
| P3 | **19(22)** | **10(12)** | **5(5)** | **4(5)** | **8(8)** | **7(8)** | **6(6)** | **8(10)** |
| P4 | **20(22)** | **10(12)** | **5(5)** | **5(5)** | **8(8)** | **8(8)** | **5(6)** | **9(10)** |
| P5 | **22(22)** | **8(12)** | *0(5)* | **4(5)** | **6(7)** | *8(8)* | *3(6)* | *7(10)* |
| P6 | **21(22)** | **12(12)** | **5(5)** | **5(5)** | **8(8)** | **8(8)** | **6(6)** | **10(10)** |
| P7 | **22(22)** | **12(12)** | **5(5)** | **5(5)** | **8(8)** | **8(8)** | **6(6)** | **10(10)** |
| P8 | **21(22)** | **12(12)** | **5(5)** | **5(5)** | **8(8)** | **8(8)** | **6(6)** | **10(10)** |
| P9 | **20(22)** | **12(12)** | **5(5)** | **5(5)** | **8(8)** | **8(8)** | **6(6)** | **8(10)** |
| P10 | **23(23)** | **24(24)** | *0(5)* | *2(5)* | **7(8)** | **8(8)** | **12(12)** | **10(10)** |
| P11 | **15(20)** | *3(12)* | *3(5)* | **3(5)** | **8(8)** | *8(8)* | *6(6)* | *7(10)* |
| P12 | **24(24)** | **24(24)** | **5(5)** | **4(5)** | **8(8)** | **8(8)** | **11(12)** | **10(10)** |
| P13 | **22(22)** | **12(12)** | **5(5)** | **5(5)** | **8(8)** | **8(8)** | **6(6)** | **10(10)** |
| P14 | **19(19)** | **9(11)** | **5(5)** | **5(5)** | **8(8)** | **8(8)** | **6(6)** | **10(10)** |
| P15 | **20(22)** | **12(12)** | **5(5)** | **3(5)** | **8(8)** | **8(8)** | **6(6)** | **10(10)** |

Notes: Numbers show fMRI runs available for analysis after exclusions; numbers in brackets are the total scanned runs. Bold indicates data were included in final analyses; italics indicates that the task was excluded for that participant. The Theory-of-Mind numbers combine the Pain and False Belief task runs. P10 and P12 had 24 Motor runs and up to 12 Sentence Processing runs due to their participation in Saadon-Grosman et al. (2022).

### Data Processing and Registration that Minimizes Spatial Blurring

Data were processed using an in-house preprocessing pipeline (“iProc”) that preserved spatial details by minimizing blurring and multiple interpolations (described in detail in Braga et al. 2019). For the refinement stage data (S1 and S2), the processed data were taken directly from Xue et al. (2021). For the implementation stage data (P1 to P15), the changes in processing included the use of the high resolution T1w and T2w structural images. For one participant (P12), the registration failed with the 0.8 mm T1w image and their 1.2 mm image was used as a back-up. For another participant (P1), only the 0.8 mm T1w image was used without a paired T2w image.

Data were interpolated to a 1-mm isotropic T1w native-space atlas (with all processing steps composed into a single interpolation) that was then projected using FreeSurfer v6.0.0 to the fsaverage6 cortical surface (40,962 vertices per hemisphere; Fischl et al. 1999). Four transformation matrices were calculated: 1) a motion correction matrix for each volume to the run’s middle volume [linear registration, 6 degrees of freedom (DOF); MCFLIRT, FSL], 2) a matrix for field-map-unwarping the run’s middle volume, correcting for field inhomogeneities caused by susceptibility gradients (FUGUE, FSL), 3) a matrix for registering the field-map-unwarped middle BOLD volume to the within-individual mean BOLD template (12 DOF; FLIRT, FSL), and 4) a matrix for registering the mean BOLD template to the participant’s T1w native-space image which was resampled to 1.0 mm isotropic resolution (6 DOF; using boundary-based registration, Freesurfer). The individual-specific mean BOLD template was created by averaging all field-map-unwarped middle volumes after being registered to an upsampled 1.2 mm and unwarped mid-volume template (an interim target, selected from a low motion run, typically acquired close to a field map).

For resting-state fixation runs, confounding variables including 6 head motion parameters, whole-brain, ventricular signal, deep cerebral white matter signal, and their temporal derivatives were calculated from the BOLD data in T1w native space. The signals were regressed out from the BOLD data using 3dTproject (AFNI; Cox et al. 1996; 2012). The residual BOLD data were then bandpass filtered at 0.01–0.1-Hz using 3dBandpass (AFNI; Cox et al. 1996; 2012). For task data runs, only whole-brain signal was regressed out (see DiNicola et al. 2020). Data registered to the T1w native-space atlas were resampled to the fsaverage6 standardized cortical surface mesh using trilinear interpolation (featuring 40,962 vertices per hemisphere; Fischl et al. 1999) and then surface-smoothed using a 2-mm full-width-at-half-maximum (FWHM) Gaussian kernel. The iProc pipeline thus allowed for high-resolution and robustly aligned BOLD data, with minimal interpolation and signal loss, output to two relevant spaces: the native space and the fsaverage6 cortical surface. Analyses were performed on the fsaverage6 cortical surface, but the network estimates (parcellations) were projected back into the individual participant’s native space allowing both surface-based and volume visualization.

### Individualized Network Estimates of the Cerebral Cortex

The MS-HBM was implemented to estimate cortical networks (Kong et al. 2019). The MS-HBM was independently implemented for the refinement stage data (S1 and S2) and then subsequently for the implementation stage data (in three separate groups P1-P5, P6-P10, and P11-P15). Estimating the model separately for multiple small groups allowed for prospective replication. As the results will reveal, the procedure was robust.

First, the connectivity profile of each vertex on the fsaverage6 cortical surface was estimated as its functional connectivity to 1,175 regions of interest (ROIs) that uniformly distributed across the fsaverage5 surface meshes (Yeo et al. 2011). For each run of data, the Pearson’s correlation coefficients between the fMRI time series at each vertex (40,962 vertices / hemisphere) and the 1,175 ROIs were computed. The resulting 40,962 x 1,175 correlation matrix was then binarized by keeping the top 10% of the correlations to obtain the functional connectivity profiles (Yeo et al. 2011).

Next, the expectation-maximization (EM) algorithm for estimating parameters in the MS-HBM was initialized with a group-level parcellation from the HCP S900 data release (that itself used the clustering algorithm from our previous study; Yeo et al. 2011). It is important to note that the goal of applying the model in this study was to obtain the best estimate of networks within each individual participant’s dataset, not to train parameters and apply them to unseen data from new participants (see Kong et al. 2019). In this analysis, as with our previous study using this approach (Xue et al. 2021), we did not include the validation step described in Kong et al. (2019), so no spatial smoothness prior was applied. Only the training step described in Kong et al. (2019) was conducted here. A network label assignment for each vertex was obtained for each participant within the training step.

#### Refinement Stage Data

Data from the two participants (S1 and S2) were analyzed together using the same MS-HBM model. The data were used to estimate and compare 15-network and 10-network MS-HBM models, as well as to explore reliability of model estimates when subsets of data were analyzed. The results from these initial two participants guided the subsequent processing of the implementation stage data.

The specific impetus for exploring a 15-network model was that networks at or near to the insula did not distinguish multiple networks that had been reported in the literature, variably labeled the Cingular-Opercular Network and Salience Network (Seeley et al. 2007; See also Seeley 2019), as well as established distinctions at or around primary visual and somatomotor^1^ cortex. The 15 candidate networks explored here are labeled^2^: Somatomotor-A (SMOT-A), Somatomotor-B (SMOT-B), Premotor-Posterior Parietal Rostral (PM-PPr), Cingular-Opercular (CG-OP), Salience / Parietal Memory Network (SAL / PMN), Dorsal Attention-A (dATN-A), Dorsal Attention-B (dATN-B), Frontoparietal Network-A (FPN-A)^3^, Frontoparietal Network-B (FPN-B), Default Network-A (DN-A), Default Network-B (DN-B), Language (LANG), Visual-Central (VIS-C), Visual-Peripheral (VIS-P), and Auditory (AUD).

#### Implementation Stage Data: Discovery, Replication and Triplication Datasets

A key aspect of our methods is generalization and replication. The 15 participants in the implementation stage data were divided into discovery, replication and triplication datasets of 5 participants each^4^. The MS-HBM model, initialized with a 15-network group-level parcellation obtained from the HCP S900 data, was applied independently to the three separate datasets.

### Model-Free Seed-Region Based Confirmation of the Networks

When employing the MS-HBM, there are assumptions about the organization of the brain from the group prior, how many networks should be estimated, and assignment of vertices to only a single network. The idiosyncratic patterns of estimated networks thus could be distorted or fail to capture features of the underlying correlation matrix. To confirm that the individual network estimates were not obligated by the assumptions, a model-free seed-region based analysis was conducted using the same data as the MS-HBM model, mirroring the procedures outlined by Braga and Buckner (2017). The results were expected to converge if the model did not overly bias network assignments and diverge if the assignments mismatched the underlying data patterns. Model-free seed-region based confirmation thus served as a check to ensure network estimates properly captured individual correlation patterns.

For this control check, the pair-wise *Pearson* correlation coefficients between the fMRI time courses at each surface vertex were calculated for each resting-state fixation run, yielding an 81,924 x 81,924 matrix (40,962 vertices / hemisphere). The matrix was then Fisher r-to-z transformed and averaged across all runs to yield a single best estimate of the within-individual correlation matrix. This averaged matrix was used to explore network organization. The mean correlation maps were assigned to a cortical template combining left and right hemispheres of the fsaverage6 surface into the CIFTI format to interactively explore correlation maps using the Connectome Workbench’s wb_view software (Glasser et al. 2013; Marcus et al. 2011). Seed regions with robust functional connectivity correlation maps were manually selected within MS-HBM network boundaries. Anterior and posterior seed regions were recorded and visualized for each network in all the participants. Thresholds were set at z(r) > 0.2 for all seed regions. The color scales of correlation maps were thresholded between 0.2 and 0.6, using the Jet look-up table (colorbar) for visualization.

### Visualization Within the Individual Native-Space Volume

Networks were first estimated and analyzed for each individual on the normalized fsaverage6 surface of FreeSurfer. Surface-based analyses allowed comparisons across individuals and utilization of the group-based priors for initialization of the MS-HBM. However, many applications require network assignments to be utilized within the native-space anatomy of the individual’s own volume (e.g., for presurgical planning and neuromodulation targeting). Given these needs, we devised a robust empirical procedure to project the network estimates back into each individual’s native-space T1w anatomical volume.

We constructed three separate images within the native-space volume that each varied from 0-255 in one of the three Cartesian x, y, and z coordinate axes (e.g., the X-coordinate image possessed a volume that linearly varied in the X-dimension going from 0 to 255 with no other variation across the image volume). Each separate axis-volume was then projected to the fsaverage6 surface using mri_vol2surf and mri_surf2surf (FreeSurfer v6.0.0) with the same spatial transformation used for the projection of the participant’s BOLD fMRI data onto the fsaverage6 surface. Nearest neighbor interpolation was used. The matrices for this projection were taken from each participant’s processing pipeline (iProc).

In this manner, x, y, and z volume coordinates were obtained on the surface using the exact same spatial transformation matrix as originally applied to the BOLD data. We assigned each surface network label to its corresponding x, y, and z coordinates in the native-space volume. This resulted in a sparse 256 x 256 x 256 matrix in the volume, which was filled in using nearest neighbor interpolation (Matlab knnsearch). We then masked this with the individual’s FSL-reoriented and binarized cortical ribbon generated by FreeSurfer during preprocessing. As a control check, the final native-space network estimates were projected back to the surface and compared to the original MS-HBM surface estimates for each participant to ensure no spatial distortions.

The resulting estimates of networks in volume space are provided as a reference in the Supplemental Materials. Specifically, the parcellation results from MS-HBM were overlaid onto each individual’s T1w structural image. Sagittal, axial, and coronal slices were chosen to show common landmarks in each individual (midline, left and right insula, anterior commissure, primary sensory and motor cortices).

### Signal-to-Noise Ratio (SNR) Maps

Data using BOLD-contrast (T2* images) and echo-planar imaging result in variable distortion and signal dropout due to magnetic susceptibility artifacts, especially near the sinus and ear canals (e.g., Ojemann et al. 1997). Vertex-based SNR maps were computed by taking the preprocessed time series from each resting-state fixation run (prior to regressing out confounding variables) and dividing the mean signal at each vertex by its standard deviation over time. The SNR maps were then averaged across the runs, resulting in an aggregate within-individual SNR map on the fsaverage6 surface. To visualize these effects in the native anatomy, surface maps were projected to the native-space volume using the procedure described above. The only difference is that linear interpolation (Matlab scatteredInterpolant) was used to fill in the sparsely filled 256 x 256 x 256 matrix.

### Variability in Network Estimates Between Individuals

To measure spatial variability across individuals, overlap maps of network assignments were computed. For each individual, the spatial extent of their estimated network was plotted simultaneously with all other participants and the percentage of overlap computed. In addition, the individual networks were plotted next to one another to appreciate the commonalities across individuals as well as the idiosyncratic features of each individual’s estimate (available in the Supplemental Materials).

Overlap maps were also computed for the model-free seed-region correlation maps. These maps make no assumption of a winner-take-all network assignment so provide a different view of network consistency or inconsistency across participants. For this final analysis, each individual’s seed-region correlation map for each network was thresholded at z(r) > 0.2 and the overlap across participants plotted. The analysis was performed separately for both the anterior and posterior seed regions for each network.

### Visualization on the Flattened Cortical Surface

The human cerebral cortex is a complex structure with numerous sulci and gyri that can make it difficult to appreciate topographic patterns, including patterns that evolve over medial to lateral views and through complex structures like the insula. To appreciate global topographic relations, a flattened surface was created by editing the inflated surface file using the “TKSurfer” tool of FreeSurfer v6.0.0. Five linear cuts were made on the midline of the inflated cortical surface (see Fig. 22), including one along the calcarine sulcus and four roughly equally spaced cuts radiating out from the medial wall. Next, a circular cut was made on the midline to allow the surface to unravel. Finally, the “mris_flatten” tool of FreeSurfer v6.0.0 was employed to create the flattened surface. This procedure was performed separately for the left and right hemispheres.

### Task Paradigms

Following estimation of within-individual networks, functional response properties were explored in independent task-based data collected on the same individuals. The task paradigms were chosen based on literature review and our prior studies because of their ability to differentially activate distinct networks, and to do so robustly. A second feature of the selected task paradigms is that they were amenable to repeat testing either because extensive novel stimuli could be constructed (e.g., sentences, question probes) or, by their nature, were resilient to habituation even after many repetitions (e.g., flickering visual stimuli). Task details are described below.

#### Somatomotor Topography

The motor task extended from Buckner et al. (2011) to examine the organization of the foot, glute, hand and tongue representations. Novel targeting of the glute representation allowed an intermediate body position to be mapped between the hand and foot (as reported earlier in Saadon-Grosman et al. 2022). The goal of this task paradigm was to activate somatotopic portions of SMOT-A and SMOT-B.

Following extensive pre-scan training, participants performed six types of active movements in the scanner: 1,2) left and right finger taps (thumb to index and thumb to middle), 3,4) left and right toes plantar flexion and dorsiflexion, 5) tongue movements from right to left (touching the premolar upper teeth), and 6) contraction and relaxation of their gluteal muscles. Each movement type was performed repeatedly across 10-sec movement blocks. Prior to each movement block, a 2-sec visual cue of a drawn body part with a text label informed the participant to initiate one of the six movement types. The fixation crosshair then changed to a slow flickering black circle to pace the movements. The onset of the black circle cued movement of thumb to index finger, toes plantarflexion, tongue to the right and glutes contraction. The offset of the black circle cued movement of thumb to middle finger, toes dorsiflexion, tongue to the left and glutes relaxation. After five cycles, the word ‘END’ instructed movement cessation. Twenty-four movement blocks (4 per movement type) occurred within each run, with 16-sec blocks of passive fixation following each set of six movement blocks. Runs began and ended with fixation yielding 5 fixation blocks per run.

Each run lasted 7 min 8 sec (428 frames with the first 12 frames removed for T1 equilibration). Six motor runs were collected with full counterbalanced orders of movement conditions on each day. Runs were excluded from analysis if participants missed or failed to respond to cues, as confirmed by operators observing their alertness and movements from the control room.

#### Visual Topography

A visual retinotopic stimulation task was used to map visual cortex (similar to Fox et al. 1987; Engel et al. 1997). Our design had three levels of eccentricity stimuli (to map eccentricity gradients that span the V1, V2, V3 cluster) and separate vertical versus horizontal meridian stimuli (to map polar angle reversals that separate the borders of V1, V2, and V3; Tootell et al. 1995; see also Wandell and Winawer 2011). The goal of this task was to activate retinotopic portions of VIS-C and VIS-P.

The basic stimulus consisted of a circular checkerboard that expanded outwards from the central fixation point to approximate cortical expansion in visual cortex. Moving from center, the radius ring of the checkerboard became larger by a log step of 0.29. The resulting checkerboard was rendered out to 36 even rings cropped to a resolution of 1024 x 1024 pixels. To localize the meridians, two wedges masked the checkerboard each covering 0.5° to 16.2° of eccentricity and 11.2° of polar angle. Horizontal wedges were centered at polar angles 360° and 180°; vertical wedges at 0° and 90°. To localize polar angle, the checkerboard was masked with a circular ring, which increased in size with increasing eccentricity. The center ring covered 0.5° to 1.6°, the middle ring 1.6° to 5.1°, and the peripheral ring 5.1° to 16.2°.

Each run consisted of 10 10-sec blocks of visual stimulation (2 blocks of each of the 5 conditions). The beginning, middle, and end of each run included a 20-sec block of extended fixation. During stimulation the checkerboard changed 6 times per sec in the order: white/black, color, black/white, color, white/black, black/white. The black center fixation dot unpredictably changed to gray (every 1 to 5 sec). To ensure continuous fixation, participants pressed a button every time the fixation dot changed to gray. The primary contrasts of interest were horizontal versus vertical meridian blocks, and separately the three eccentricity blocks versus each other.

Each run lasted 4 min 30 sec (270 frames with the first 6 frames removed for T1 equilibration). Five runs were collected for each participant. Runs were excluded from analysis if participants missed trials and the eye video recordings indicated drowsiness. Lights within the scanner room were off during visual topography mapping, and a black occluding board was inserted into the scanner to prevent any light reflections.

#### Oddball Task

The oddball task explored detection of transient responses to salient, visual oddball targets that were uncommon relative to irrelevant non-targets and distracting non-targets (similar to Wynn et al. 2015). The goal of the task was to activate the SAL / PMN and CG-OP networks. Both networks have regions at or near the anterior insula and have been variably associated with response to task-relevant transients (see Dosenbach et al. 2006; Seeley et al. 2007; Seeley 2019 for discussion).

Participants viewed a sequence of uppercase letters O and K in either black or red. Participants pressed a button using their right index finger when a red K appeared and withheld their responses to all the other letter-color combinations. The random trial ordering was set using Optseq (Dale 1999). In each run, 10% of the trials were target red Ks, 10% were lure red Os, 40% were distractor black Ks, and 40% were distractor black Os. The contrast of interest was the target red Ks versus all other trials coded as the implicit baseline.

Each run lasted 5 min 50 sec (350 frames with the first 6 frames removed for T1 equilibration). Following 6 sec of fixation overlapping the initial stabilization frames, a block of 20 sec of fixation was followed by a continuous extended block of 300 1-sec trials (0.15 sec presentation of the letter followed by 1.85 sec of fixation), and then a final 20-sec block of extended fixation. Before the first trial, a 2-sec start cue (1 sec “Begin”, 1 sec fixation) was presented, as well as a similar “End” cue after the final trial. Thus, the design was a rapid, event-related paradigm sandwiched between blocks of extended fixation. Five runs were collected for each participant. Runs were excluded from the analysis if participants missed more than six targets within a task run, which accounted for 20% of the total targets.

#### Working Memory (N-Back) Task

The working memory (N-Back) task was extended from Cohen et al. (1994) and Braver et al. (1997) to explore demands on cognitive control under varied levels of memory load. Specifically, the N-Back task utilized a 2-back versus 0-back comparison to target FPN-A and FPN-B. In addition, following the design of the HCP N-Back task, multiple stimulus types / matching rules were included to explore whether the load effect was domain-flexible or domain-specialized (Barch et al. 2013).

Stimuli were presented sequentially in the center of the computer screen. Participants decided whether the current stimulus matched a consistent template target (the 0-Back or low load condition) or whether the current stimulus matched the stimulus presented two stimuli back in the sequence (the 2-Back or high load condition). Participants maintained fixation on a central crosshair throughout the run.

The stimuli varied across four conditions (Face, Word, Scene, and Letter) that were each presented in separate blocks. Faces and scenes were color images, with scenes showing both indoor and outdoor spaces and chosen not to feature people (faces from HCP; Barch et al. 2013; scenes generously provided by the Konkle laboratory; Konkle et al. 2012; Josephs and Konkle 2020). Letters included subsets of consonants, and words featured 1-syllable words from 10-word sets matched for length and frequency (as reported by the Corpus of Contemporary English; Davies 2008, vDecember 2015). In all but the Word condition, participants matched the stimuli to an exact stimulus referent, or the exact stimulus presented two trials before. For the Word condition, the participants decided if the current word rhymed with the target (e.g., “dream” would be a positive match with “steam”).

Each N-Back run featured 8 blocks (a 0-Back and a 2-Back for each of the four stimulus categories). Each block included a cue and 9 trials. During the first cue stimulus, participants also saw the block type, either 2-Back or 0-Back. During 2-Back blocks, participants looked for matches (identical images or rhyming words) with the stimulus 2 trials back, and during 0-Back blocks, participants looked for matches to the cue. The background was black (matching the HCP format). All blocks included 2 target and 2 lure (repeated non-target) trials. Targets and lures were equally likely to appear in each viable trial position within and across runs. Participants pressed a button for every trial, indicating match (right index finger) or no-match (left index finger).

Each run lasted 4 min 44 sec (284 frames with the first 12 frames removed for T1 equilibration). Following 12 sec of fixation overlapping the initial stabilization frames, an additional block of 12 sec of fixation was followed by blocks of the N-Back task interspersed with 15-sec fixation blocks (the fixation blocks came after two 25-sec N-Back task blocks). Across runs, 0-Back and 2-Back blocks, categories, and their interactions were counterbalanced. Each trial was 2.5 sec in duration (2 sec of stimulus presentation followed by 0.5 sec of fixation). The fixation crosshair was white for the extended fixation blocks and green during the N-Back task blocks. Within a run, all categories were seen before a category repeated. Eight runs were collected for each participant. Runs where participants missed responses in more than two trials were excluded from analysis.

#### Sentence Processing Task

The Sentence Processing task was adapted from Fedorenko et al. (2010; 2012) to examine domain-specialized processing related to accessing word meaning and phrase-level meaning. The target task involved sentences presented one word at a time. The reference control task was presentation of nonword strings that were matched in length and visually similar. The goal of this task was to activate the LANG network (see Braga et al. 2020).

Participants passively read real sentences (“IN THE MORNING THE TAILOR WAS SHOWING DIFFERENT FABRICS TO THE CUSTOMER”) or pronounceable nonword strings (“SMOLE MUFRISONA VEDER SMOP FO BON FE PAME OMOSTREME GURY U FO”). The centered stimuli were presented one word (or nonword) at a time (0.45 sec per word). After each word or nonword string, a cue appeared for 0.50 sec, prompting the participants to make a right index finger button press. Stimuli, generously provided by the Fedorenko laboratory, never repeated. Word or nonword strings (6 sec each) were presented in 18-sec blocks of 3 strings. Extended fixation blocks (18 sec each) appeared at the start of each run and after every fourth string block. The primary comparison of interest was the contrast between sentence and nonword blocks.

Each run lasted 5 min 0 sec (300 frames with the first 12 frames removed for T1 equilibration). Six runs were collected for each participant. Runs were excluded if the participant did not read the stimuli (observed through eye monitoring) or missed responses.

#### Theory-of-Mind Task

The Theory-of-Mind tasks were adopted from Saxe and colleagues to explore domain-specialized processing associated with representation of other’s mental states (Saxe and Kanwisher 2003; Dodell-Feder et al. 2011; Bruneau et al. 2012; Jacoby et al. 2016). In the False Belief paradigm, participants viewed a brief story and then, on a separate screen, a question about that story. In the False Belief condition, the target stories described events surrounding a person’s perspective, followed by a question about the thoughts and beliefs of that person. In the control False Photo condition, stories described similar situations involving objects (e.g., in photos, on maps, and in descriptions). In the Emotional / Physical Pain Stories paradigm (subsequently abbreviated ‘Pain’), the target stories described a situation that evoked personal emotional pain (Emo Pain condition) and were contrasted with control stories of similar length and complexity involving physical pain (Phys Pain condition). Participants rated the level of pain from “None” to “A Lot” during the question period. These two paradigms yield similar task activation maps (Jacoby et al. 2016). Here the task contrasts of False Belief versus False Photo and Emo Pain versus Phys Pain were combined with the goal to activate DN-B (extending from DiNicola et al. 2020).

Each run consisted of a series of stories and questions (15 sec per individual story / question pairing). For both paradigms, each run included 5 target trials (False Belief or Emo Pain) and 5 control trials (False Photo or Phys Pain). 15-sec fixation periods occurred between trials. Stimuli, generously provided by the Saxe laboratory, never repeated.

Each run lasted 5 min 18 sec (318 frames with the first 12 frames removed for T1 equilibration). Eight runs were collected for each participant – 4 of the False Belief paradigm and 4 of the Pain paradigm. We implemented an exclusion criterion to exclude any run with more than one missed trial. No runs met this criterion.

#### Episodic Projection Task

The Episodic Projection task was adapted from Andrews-Hanna et al. (2010) and DiNicola et al (2020) to encourage processes related to remembering the past and imagining the future (prospection). In the target task conditions, participants viewed a brief scenario that oriented to a situation in the past (Past Self) or future (Future Self) simultaneously with a question about the event that encouraged participants to imagine the specific scenario described. The similarly structured control condition asked the participants about a present situation (Present Self). The task contrasts of Past Self versus Present Self and Future Self versus Present Self were combined with the goal to activate DN-A (extending from DiNicola et al. 2020). Of relevance, detailed behavioral analysis of these contrasts has suggested the main component process tracking increased response in DN-A is the process of mentally constructing scenes (DiNicola et al. 2023; see also Hassabis and Maguire 2007). Thus, the task contrast used here taps into domain-specialized processing related to spatial / scene processing (see Hassabis and Maguire 2009 for discussion).

Each run contained a series of scenarios with questions (10 sec of scenario / question presentation, followed by 10 sec of fixation). 30 questions appeared per run, with 3 per each condition of relevance (Past Self, Future Self, Present Self). Additional conditions were included towards goals distinct from those targeted here. For our analyses, we focus on the condition contrasts that have previously dissociated DN-A from DN-B in DiNicola et al. (2020). All scenarios were unique.

Each run lasted 10 min 17 sec (617 frames with the first 12 frames removed for T1 equilibration). Ten runs were collected for each participant that included 90 relevant trials across runs (30 of each of the 3 conditions). Runs with more than two missed trials were excluded.

### Within-individual Task Activation Analysis

Functional task data were analyzed using the general linear model (GLM) as implemented by FSL’s first-level FEAT (FSL version 5.0.4; Woolrich et al. 2001). All conditions were included in each model design, even those not relevant to the contrasts of interest, except for the Oddball Effect task contrast which coded the targets against the implicit baseline. The data were high-pass filtered using a cutoff of 100 sec (0.01-Hz) to remove low-frequency noise within each run. GLM outputs included, for each contrast, β-values for each vertex that were converted, within FEAT, to *z*-values. Within each participant, *z*-value maps from all runs were averaged together using *fslmaths* (Smith et al. 2004) to create a single cross-session map for each contrast of interest. For the N-Back task, GLM outputs included *z*-value maps for each trial block, which were averaged by condition across runs. A single cross-session contrast map was then created by taking the difference between condition mean maps.

Task contrasts were designed to functionally target specific networks and dissociate response properties between networks. Two convergent methods were used for visualization and quantification. First, *z*-value maps were compared visually by overlaying the borders of networks onto the task contrast maps on the same cortical surface (fsaverage6 cortical surface). This form of visualization allowed comprehensive assessment of task response patterns. Contrast *z*-value maps were manually thresholded to best demonstrate the task activation patterns for each participant. The PSYCH-FIXED look-up table within Connectome Workbench was used for the color scale.

Second, *a priori* networks within-individuals were used to formally quantify differences in response levels between networks, including direct tests for significant differences between networks and between task contrasts. For each task contrast, the average *z*-value was calculated for all vertices within each selected network, combining across hemispheres. Mean *z*-values were computed for each task run, and the cross-run mean *z*-values for each network was then plotted in a bar graph, along with the standard error of the mean across participants. This analysis has the advantage of quantifying the magnitude and variance of the response in each *a priori* defined network for each participant, without any subjective decisions.

For both approaches to task response analysis, the networks were defined within the individuals prior to examination of the task maps, to avoid the possibility of bias.

### Software and Statistical Analysis

Functional connectivity between brain regions was calculated in MATLAB (version 2019a; http://www.mathworks.com; MathWorks, Natick, MA) using Pearson’s product moment correlations. FreeSurfer v6.0.0, FSL, and AFNI were used during data processing. The estimates of networks in volume space were visualized in FreeView v6.0.0. The estimates of networks on the cortical space were visualized in Connectome Workbench v1.3.2. Statistical analyses were performed using R v3.6.2. Model-free seed-region confirmations were performed in Connectome Workbench v1.3.2. Network parcellation was performed using code from Kong et al. (2019) on Github: (https://github.com/ThomasYeoLab/CBIG/tree/master/stable_projects/brain_parcellation/Kong2019_MS HBM).

## Results

### Networks Can Be Estimated Robustly Within Individuals

Networks were estimated for the refinement stage data using a 15-network MS-HBM model. Figs. 1 through 6 display the main results for S1 and S2 on the surface, and the Supplementary Materials display the comprehensive results and quality control visualizations on the surface and in the native volume.

**Figure 1.**
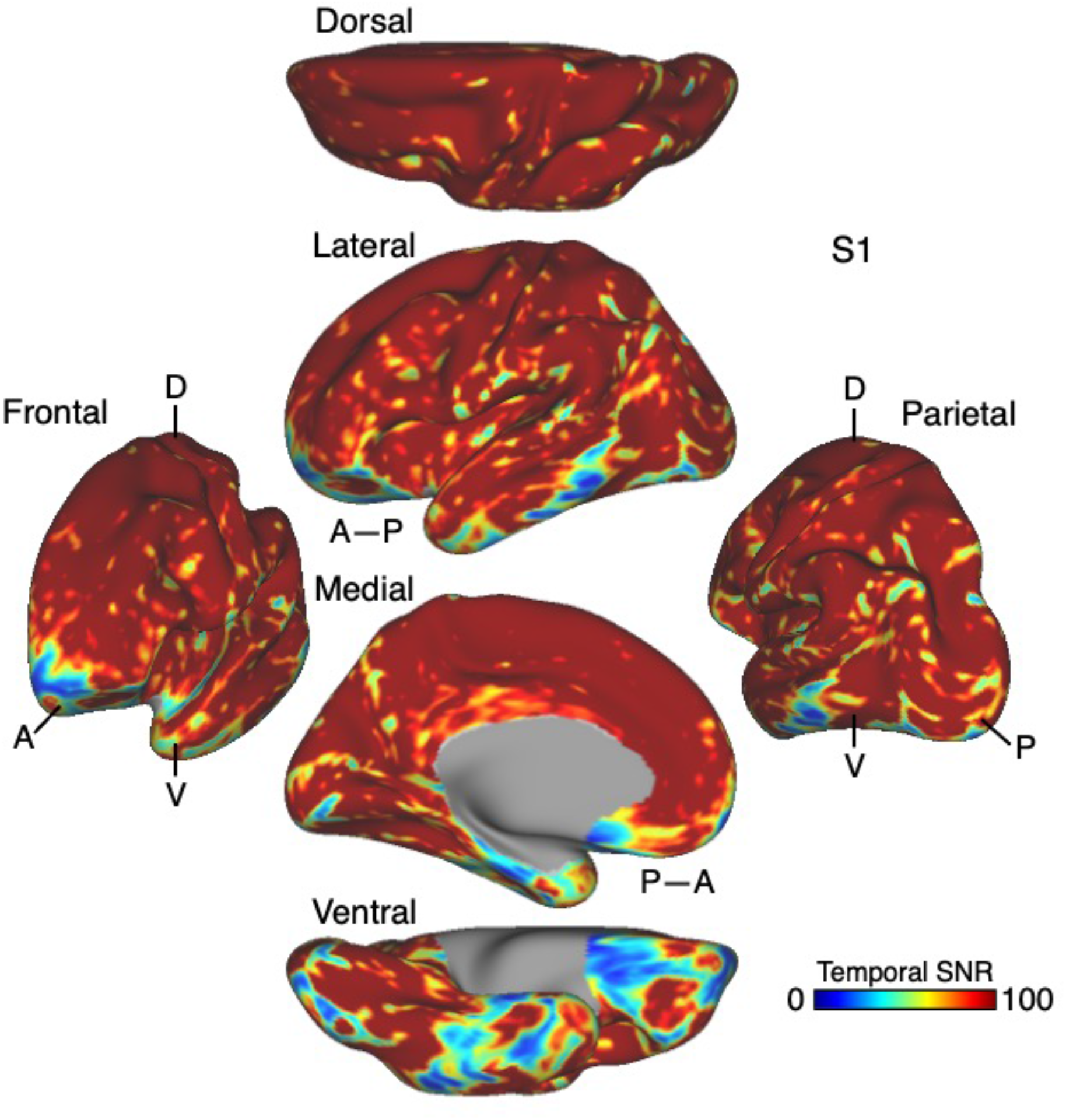
Temporal signal-to-noise ratio (SNR) map for S1. The mean estimate of temporal SNR for the fMRI data is illustrated for multiple views of the left hemisphere on the inflated cortical surface (from 62 runs collected over 31 days). Note the low SNR within the orbitofrontal cortex and the temporal pole. This pattern is typical of the data across all participants in the present work and should be considered when evaluating network organization. A, anterior; P, posterior; D, dorsal; V, ventral. SNR maps for all participants are provided in the Supplementary Materials.

**Figure 2.**
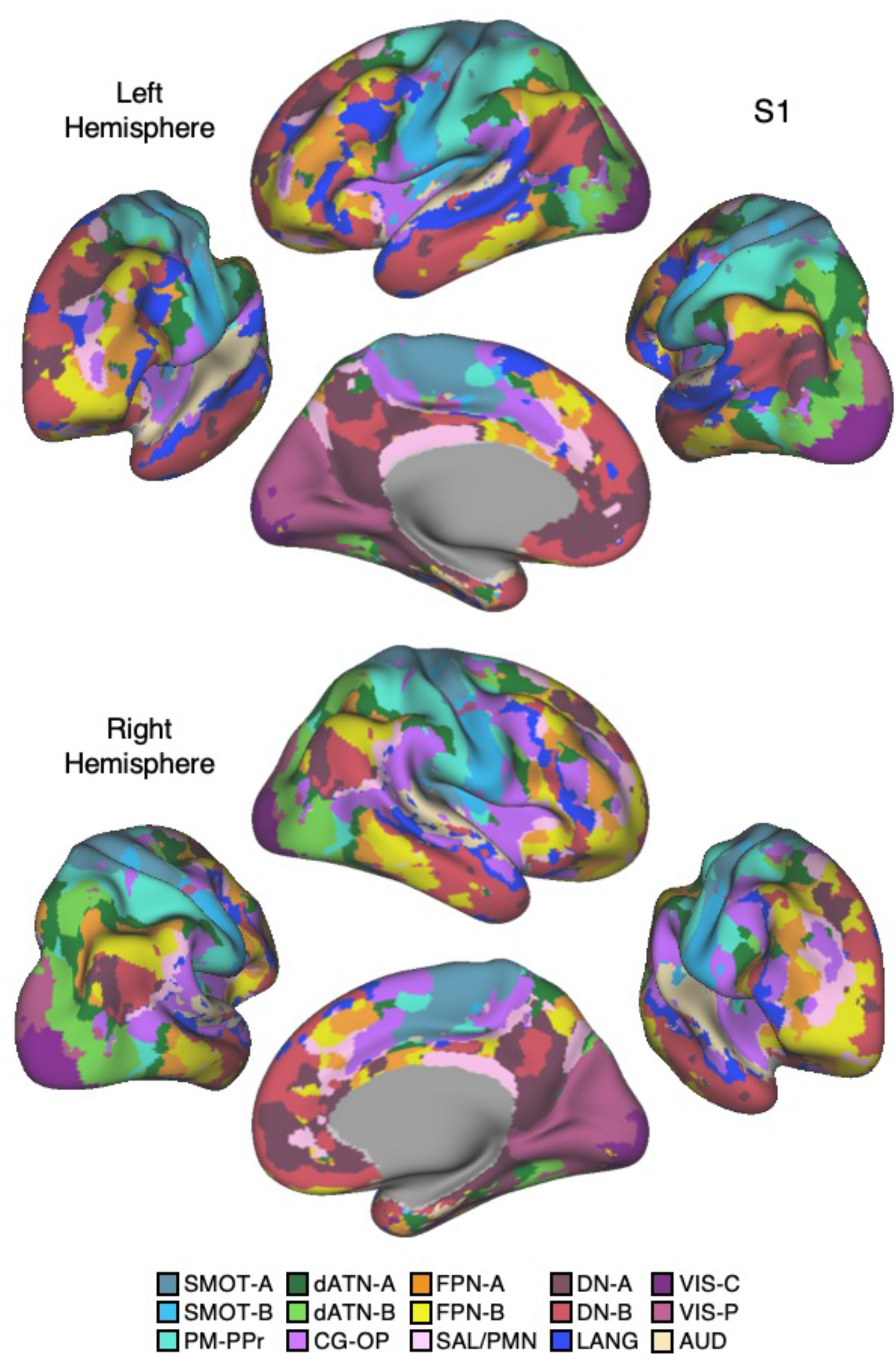
15-network cerebral cortical parcellation estimated for S1. Network estimates from the multi-session hierarchical Bayesian model (MS-HBM) are displayed across four views. The left hemisphere is on top and right hemisphere below. Each color represents a distinct network estimated by the model. Some networks possess primarily local organization (e.g., Somatomotor, Visual), while other networks possess widely distributed organization (e.g., those involving prefrontal, temporal, and parietal association zones). The network labels are used similarly throughout the figures. SMOT-A, Somatomotor-A; SMOT-B, Somatomotor-B; PM-PPr, Premotor-Posterior Parietal Rostral; CG-OP, Cingular-Opercular; SAL / PMN, Salience / Parietal Memory Network; dATN-A, Dorsal Attention-A; dATN-B, Dorsal Attention-B; FPN-A, Frontoparietal Network-A; FPN-B, Frontoparietal Network-B; DN-A, Default Network-A; DN-B, Default Network-B; LANG, Language; VIS-C, Visual Central; VIS-P, Visual Peripheral; AUD, Auditory.

**Figure 3.**
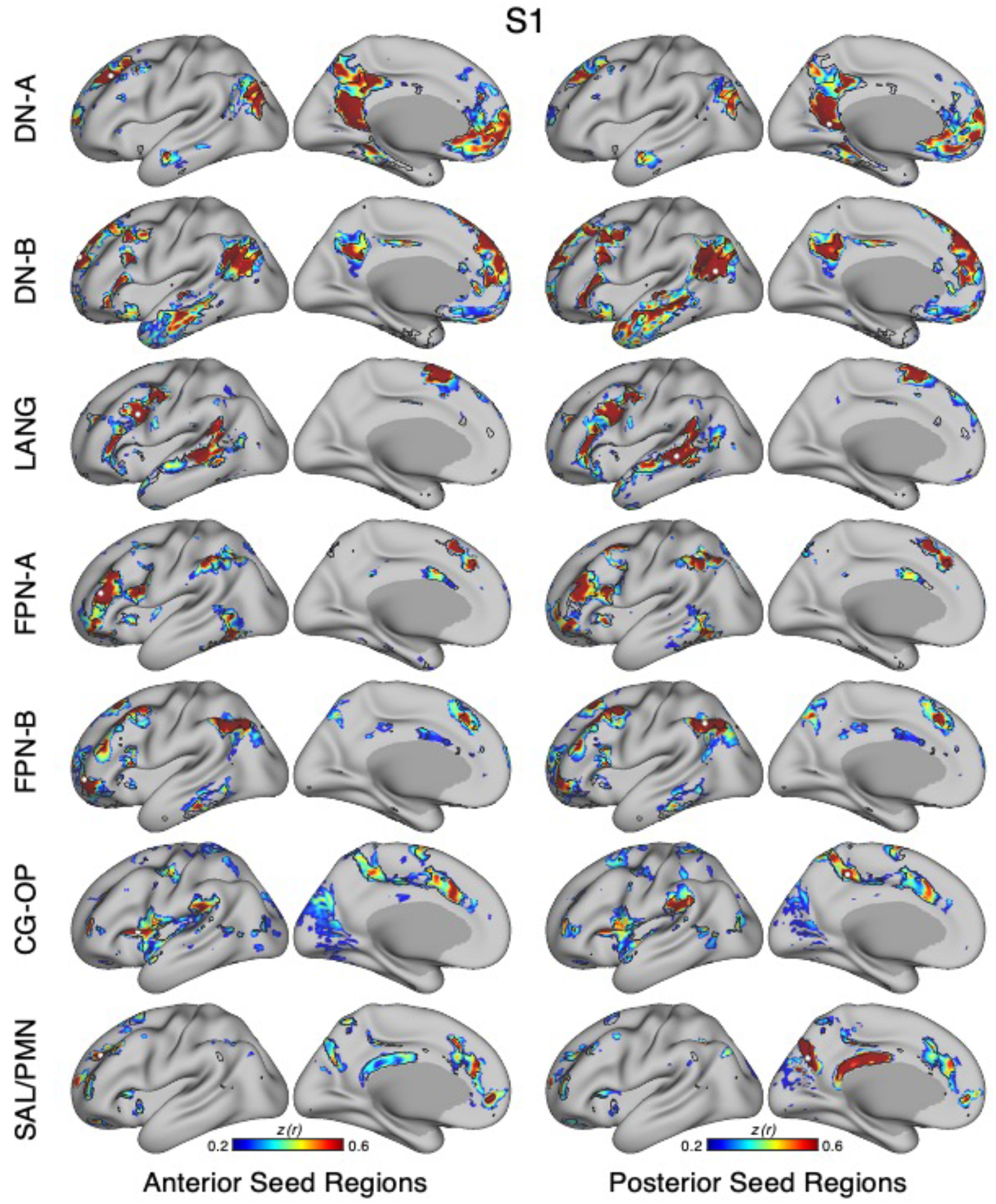
Model-free confirmation of networks using seed-region correlation for S1. The correlation patterns from individual seed regions placed within networks are displayed. In each row, a distinct network is targeted, labeled to the left. The two left columns display correlation maps using an anterior seed region of each network, while the two right columns display correlation maps using a posterior seed region. Lateral and medial views are displayed. White-filled circles display the seed region locations. Black outlines show the boundaries of individual-specific networks estimated from the MS-HBM as shown in Fig. 2. The correlation maps are plotted as z(r) with the color scale at the bottom. The correlation maps are not constrained to fall within the estimated network boundaries. Nonetheless, the network boundaries capture a great deal of the spatial correlational properties of the underlying data.

**Figure 4.**
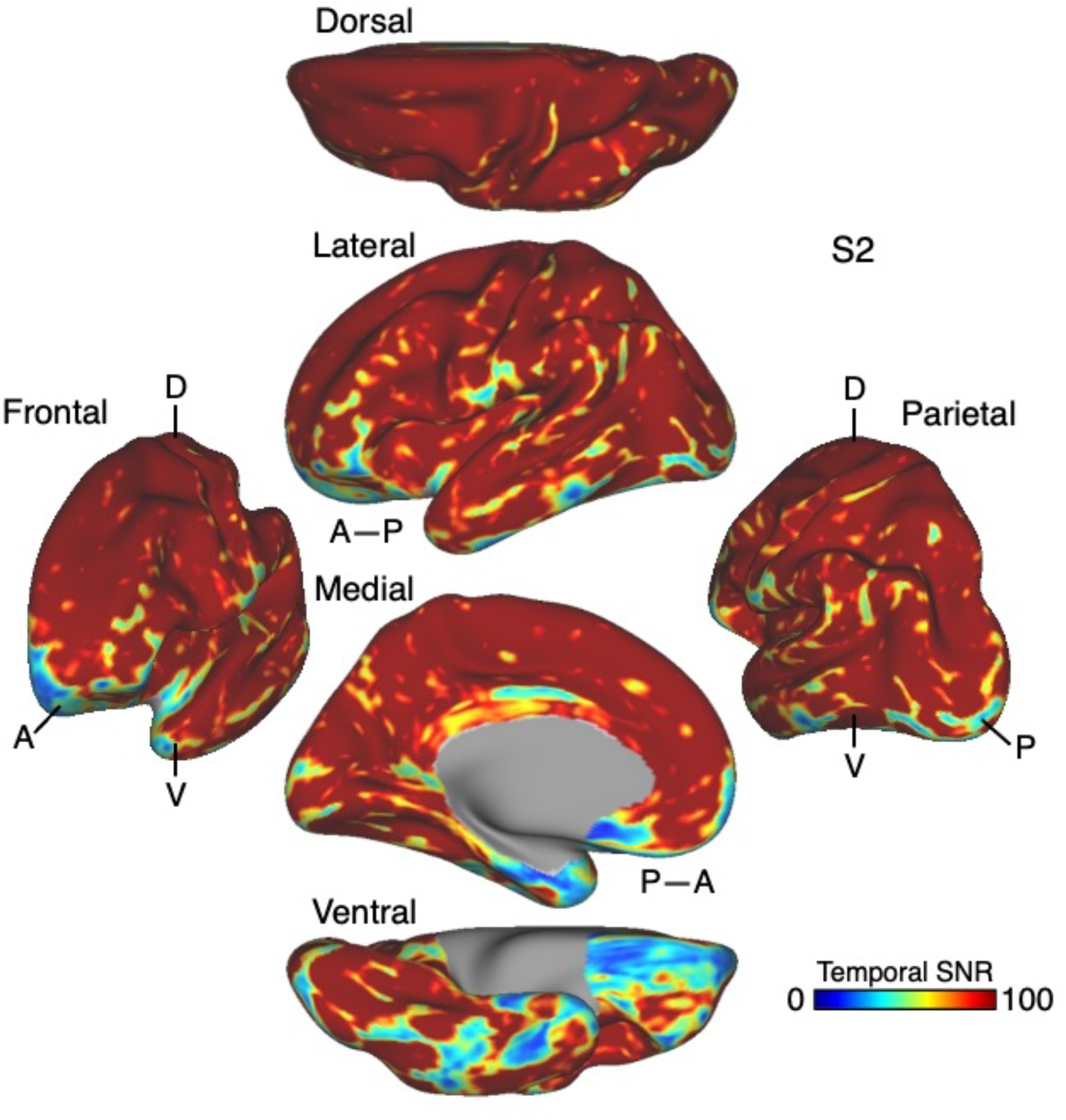
Temporal signal-to-noise ratio (SNR) map for S2. Paralleling Fig. 1, the mean estimate of temporal SNR for the fMRI data is illustrated for multiple views of the left hemisphere on the inflated cortical surface (from 61 runs collected over 31 days). A, anterior; P, posterior; D, dorsal; V, ventral.

**Figure 5.**
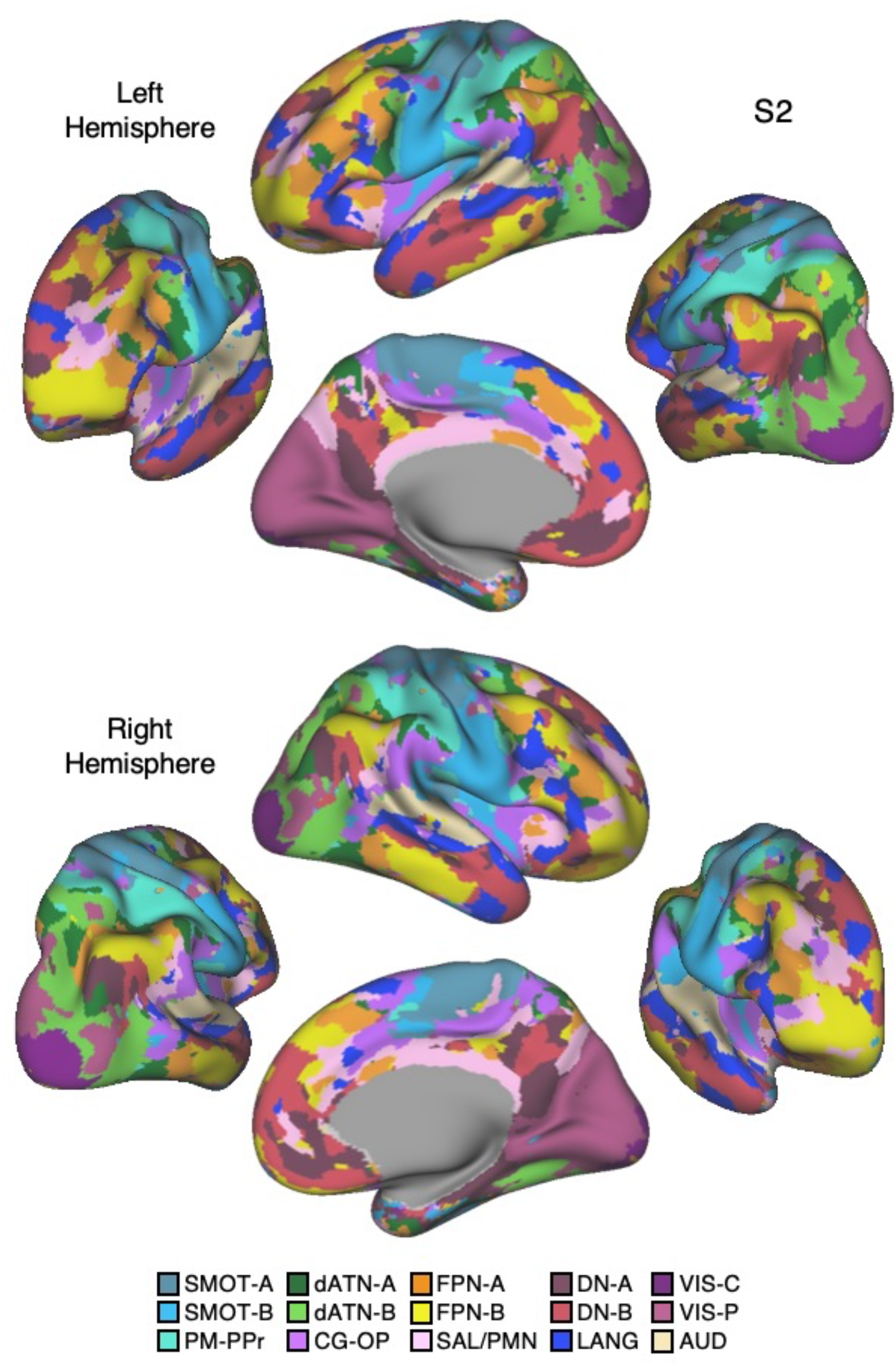
15-network cerebral cortical parcellation estimated for S2. Paralleling Fig. 2, network estimates from the MS-HBM are displayed across four views. The left hemisphere is on top and right hemisphere below. Each color represents a distinct network estimated by the model. The names of cortical networks are shown at the bottom.

**Figure 6.**
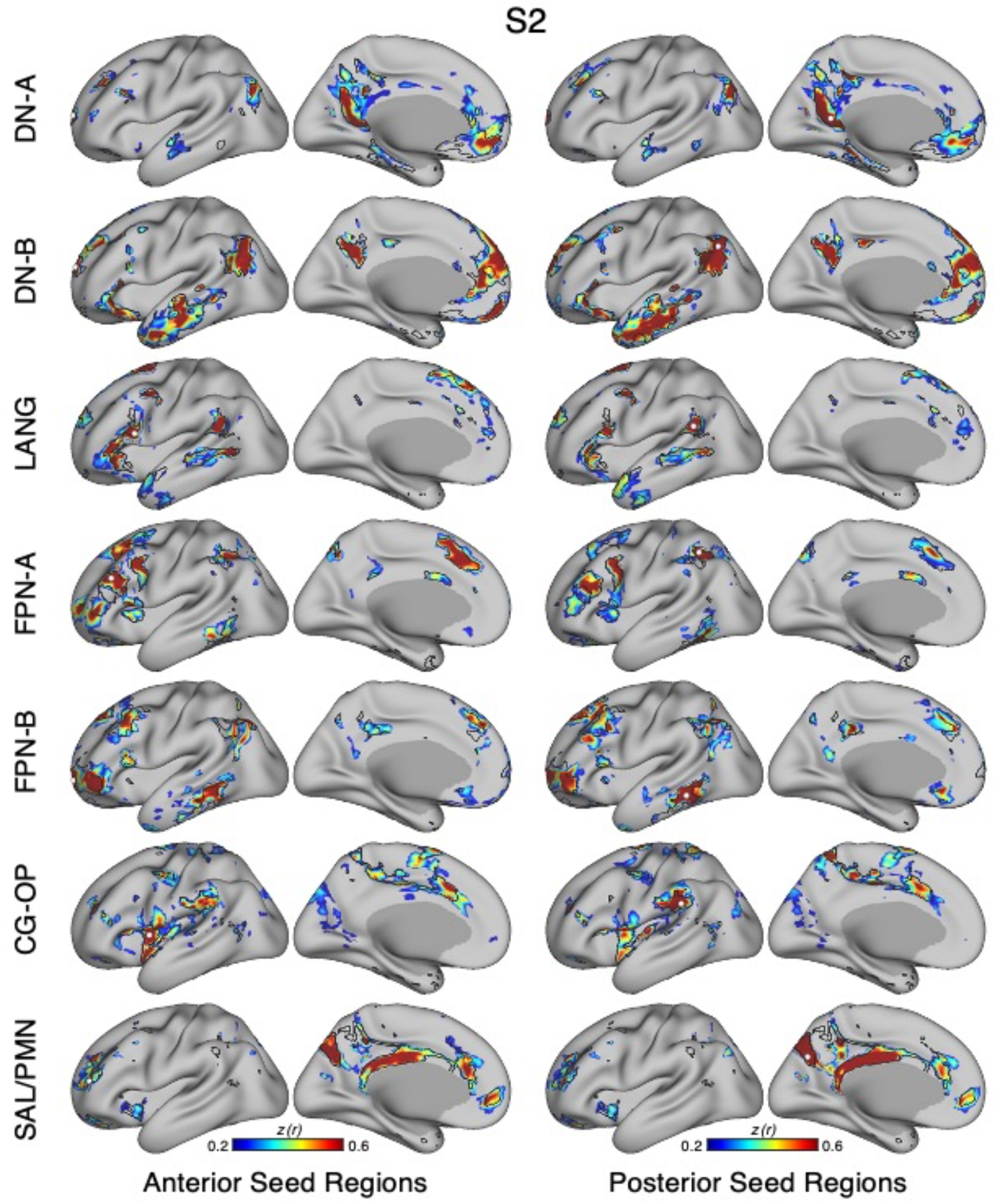
Model-free confirmation of networks using seed-region correlation for S2. Paralleling Fig. 3, the correlation patterns from individual seed regions placed within networks are displayed for S2. The two left columns display correlation maps using an anterior seed region for each network, while the two right columns display correlation maps using a posterior seed region. Lateral and medial views are displayed for each seed region. White-filled circles display the seed region locations. Black outlines indicate the boundaries of corresponding individual-specific parcellation-defined networks estimated from the MS-HBM as shown in Fig. 5. The correlation maps are plotted as z(r) with the color scale at the bottom.

The first results pertain to data quality. The SNR maps are displayed on the cortical surface (Figs. 1 and 4). Most of the cortical mantle possessed high SNR. As expected, given signal dropout near the sinuses and the inner ear (Ojemann et al. 1997), there is variability in SNR across the cortical surface with orbitofrontal cortex (OFC) and adjacent ventrolateral prefrontal cortex (VLPFC), rostral inferior temporal cortex, and the temporal pole showing low SNR (see Supplementary Materials for additional visualizations). Network assignments in low SNR regions should be interpreted cautiously.

The primary result of our procedures was an estimated parcellation into distinct candidate networks. Figs. 2 and 5 display the 15-network estimates for S1 and S2. All networks, including local sensory and motor networks, as well as distributed association networks, were identified in both participants. While the general organization was shared between the two participants, the spatial boundaries were idiosyncratic. These patterns will be elaborated upon in detail in the upcoming results of the novel 15 participants. For these first two individuals we focused on validating the methods.

### Model-Free Seed-Region Based Correlation Confirms the 15-Network Parcellation

The network estimates were based on a 15-network MS-HBM model. In addition to assuming a specific number of networks, the method also employed group priors to constrain the estimates (see Supplementary Materials). As such, it is possible that the resultant networks do not accurately reflect the underlying within-individual correlation patterns as one might expect. To explore this possibility and intuitively visualize the degree to which the model captures underlying correlation patterns, a model-free seed-region based correlation analysis was performed. A seed region was placed in an anterior position and separately a posterior position within each network within each individual. The resulting correlation maps are displayed in Figs. 3 and 6 in relation to the MS-HBM network boundaries.

The estimated networks captured features of the correlation patterns remarkably well including across small, distributed regions that might otherwise be overlooked. The alignments were not perfect. Specifically, the correlation patterns included most of the distributed regions in the MS-HBM solutions, and the patterns were largely selective to the estimated networks. Small deviations, in the form of extensions of the patterns beyond the network boundaries were common, likely in part because the network estimates forced a winner-take-all assignment, but also possibly because additional network details may be missed^5^. The consistency between the general correlational structure and the network estimates in one sense is unsurprising because the underlying correlation matrix was employed by the network model. However, it is not obligated, and deviation could be seen if the model forced assignments, or the model failed to capture the structure of the data.

### The 15-Network Parcellation Captures Features that Are Not Captured by a 10-Network Parcellation

We next sought to explore what is gained by adopting the 15-network parcellation rather than the simpler 10-network parcellation. Figs. 7 and 10 display the MS-HBM parcellation estimate for the 10-network and 15-network solutions for each participant. The first notable result is that, for most networks, there was little difference between the two models’ estimates. For example, the separation of DN-A and DN-B was well captured by both model solutions with the distributed spatial patterns and idiosyncratic features quite similar between models. That is, if the goal were to study DN-A and DN-B, there is little gained by utilizing the more complex 15-network model. In both S1 and S2, many of the other major networks were also similar between the two parcellations, including FPN-A, FPN-B, SMOT-A and SMOT-B. Thus, for networks well captured by the 10-network model, they appear to be roughly unchanged in the 15-network model. For other networks though, there were substantive differences.

**Figure 7.**
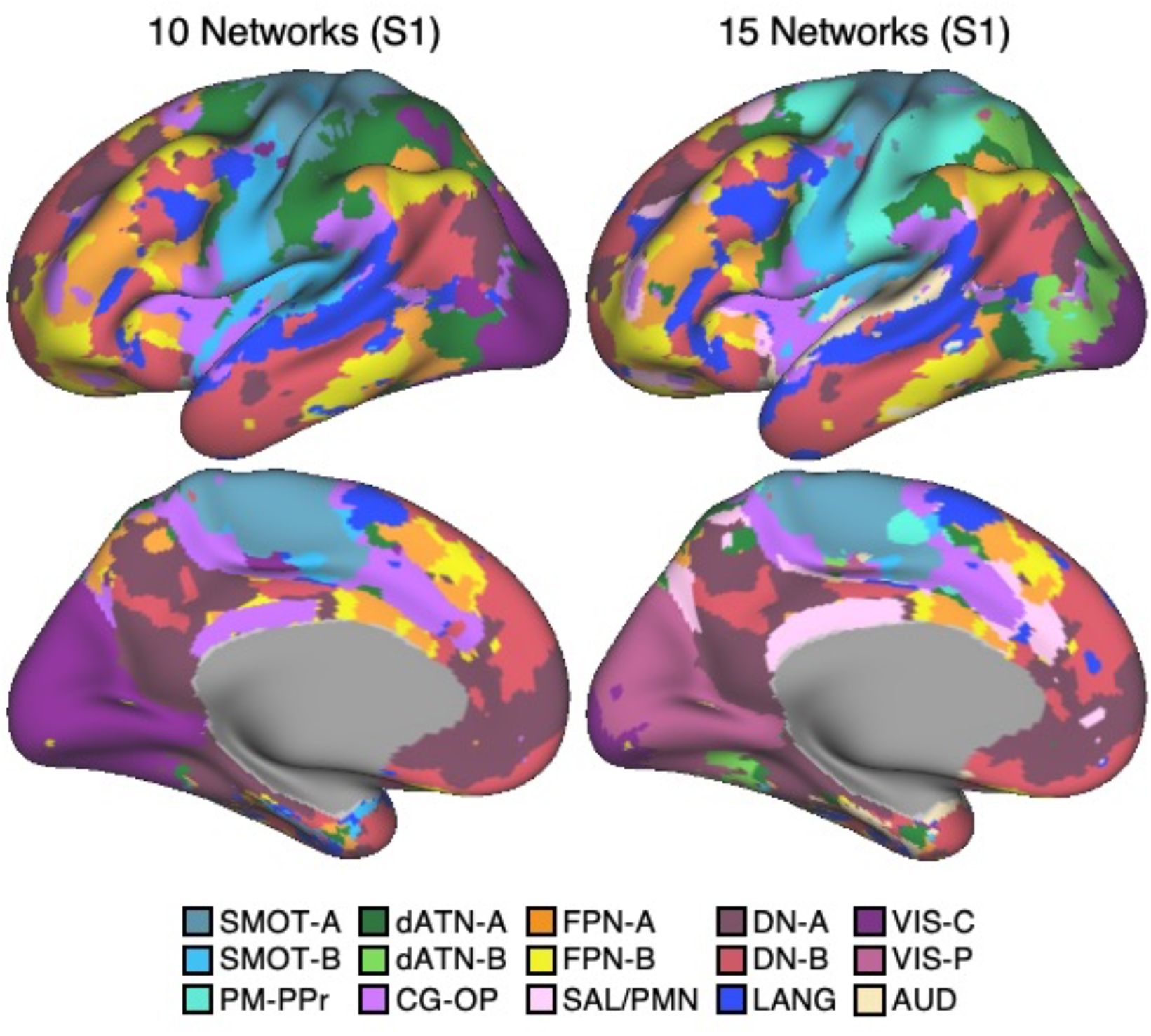
Direct comparison of 10-network and 15-network cerebral cortical parcellations for S1. The left displays the 10-network estimate and the right the 15-network estimate. Many of the major networks are similar between the two parcellations, including LANG, DN-A, DN-B, FPN-A, FPN-B, SMOT-A, SMOT-B. VIS in the 10-network estimate is differentiated into dATN-B, VIS-C and VIS-P in the 15-network estimate. A monolithic large network in the 10-network estimate is differentiated into SAL / PMN and CG-OP in the 15-network estimate. dATN in the 10-network estimate is differentiated into dATN-A and PM-PPr in the 15-network estimate, and a distinct AUD network emerges near to LANG and SMOT-B. The network labels are shown at the bottom.

**Figure 8.**
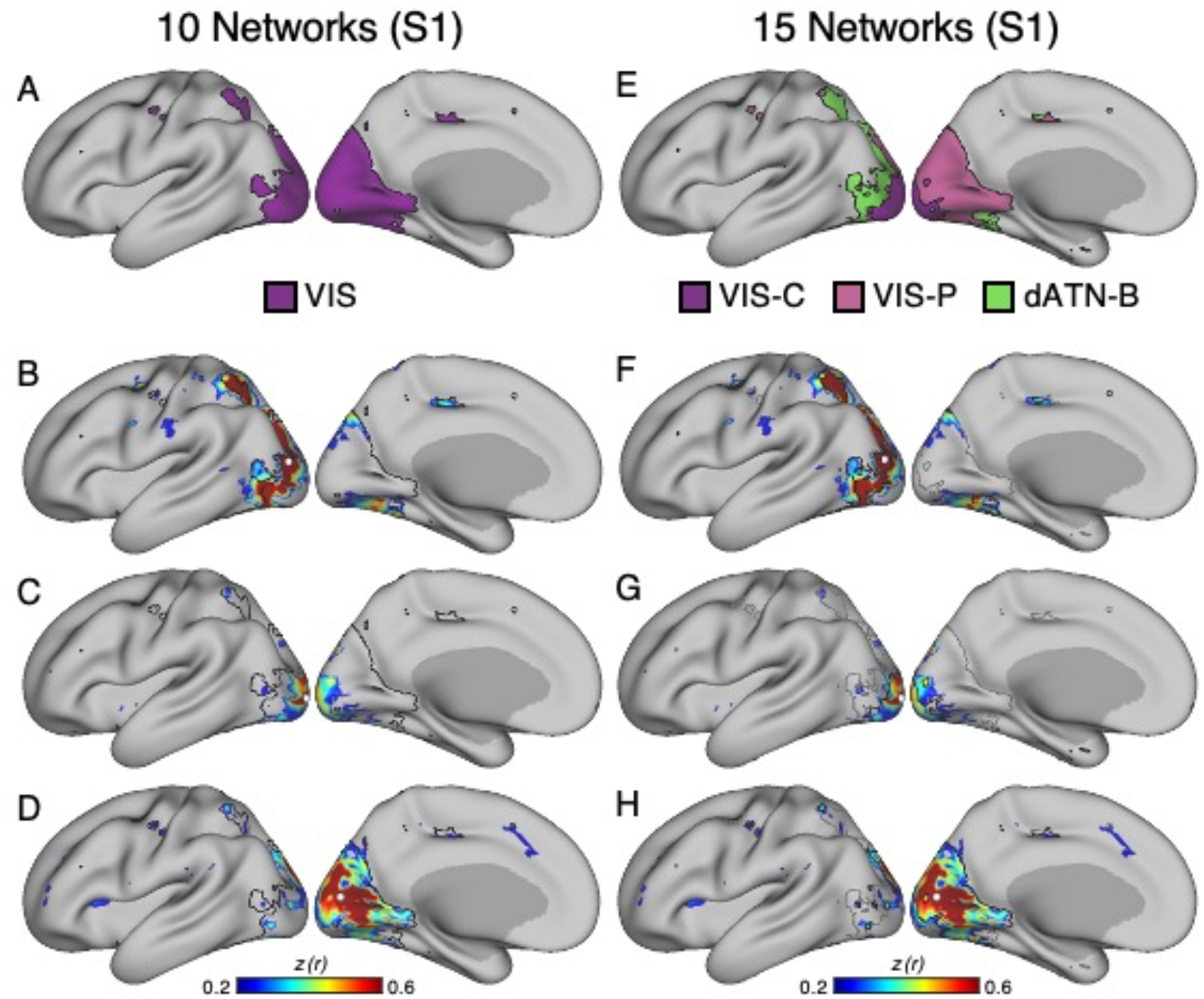
Model-free estimates illustrate the utility of the 15-network cerebral parcellation for visual networks for S1. Seed region correlation maps illustrate features captured by the 15-network estimate as contrast to the 10-network estimate. VIS in the 10-network estimate (**A**) is differentiated into dATN-B, VIS-C and VIS-P in the 15-network estimate (**E**). White-filled circles display the seed region locations. Black outlines indicate the boundaries of the networks above. The network labels are shown below. Correlation maps for three distinct seed regions in and around the vicinity of visual cortex are illustrated within the boundaries of the 10-network estimate (**B**, **C**, **D**) and the 15-network estimate (**F**, **G**, **H**). Note that the correlation patterns are well captured by the 15-network estimate. Black and gray outlines illustrate the networks from each parcellation estimate. The correlation maps are plotted as z(r) with the color scale at the bottom.

**Figure 9.**
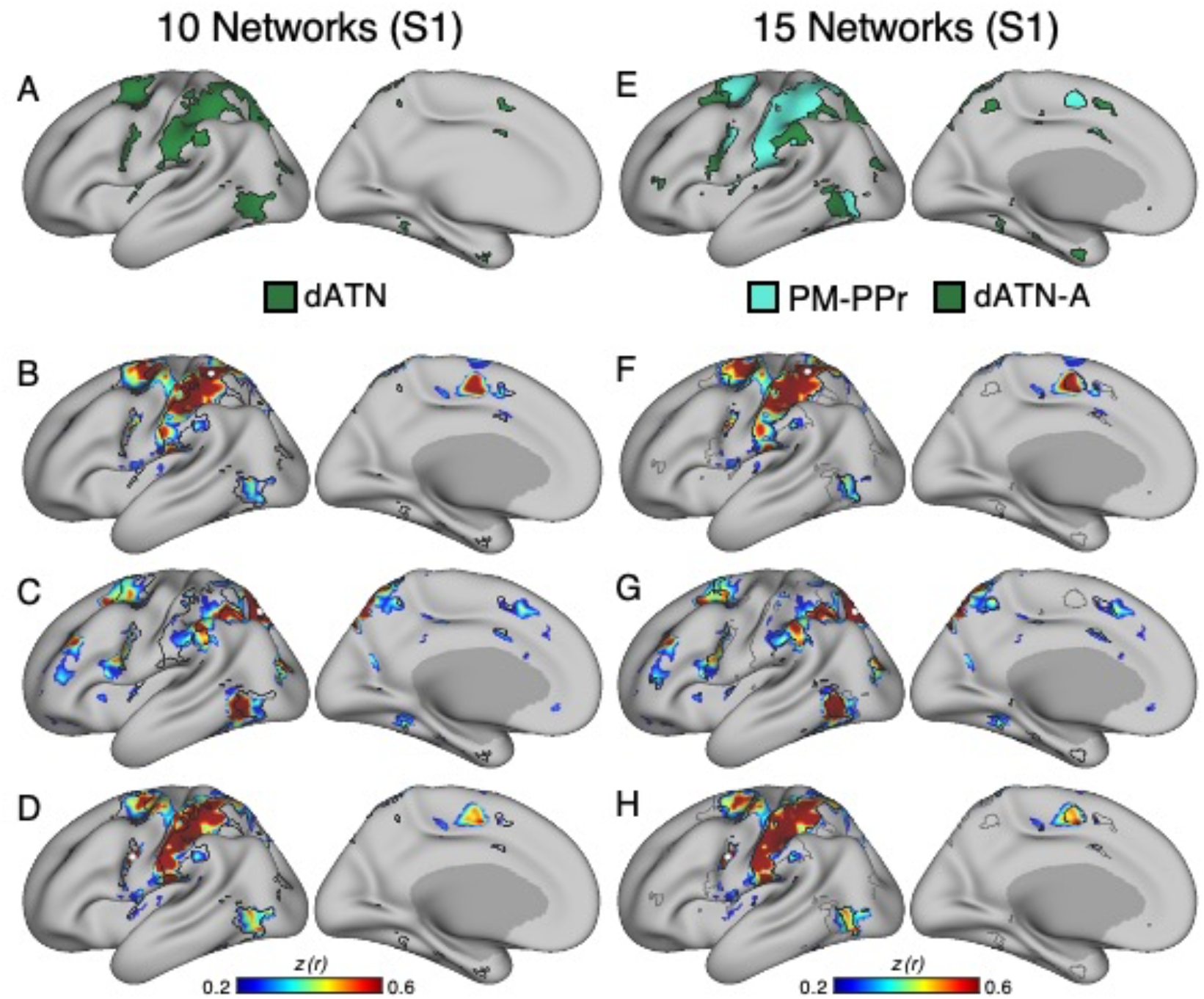
Model-free estimates illustrate the utility of the 15-network cerebral parcellation for networks surrounding somatomotor cortex for S1. Paralleling Fig. 8, seed region correlation maps illustrate features captured by the 15-network estimate as contrast to the 10-network estimate. dATN in the 10-network estimate (**A**) is differentiated into dATN-A and PM-PPr in the 15-network estimate (**E**). White-filled circles display the seed region locations. Black outlines indicate the boundaries of the networks above. The network labels are shown below. Correlation maps for three distinct seed regions surrounding somatomotor cortex are illustrated within the boundaries of the 10-network estimate (**B, C, D**) and the 15-network estimate (**F, G, H**). Black and gray outlines illustrate the networks from each parcellation estimate. The correlation maps are plotted as z(r) with the color scale at the bottom.

**Figure 10.**
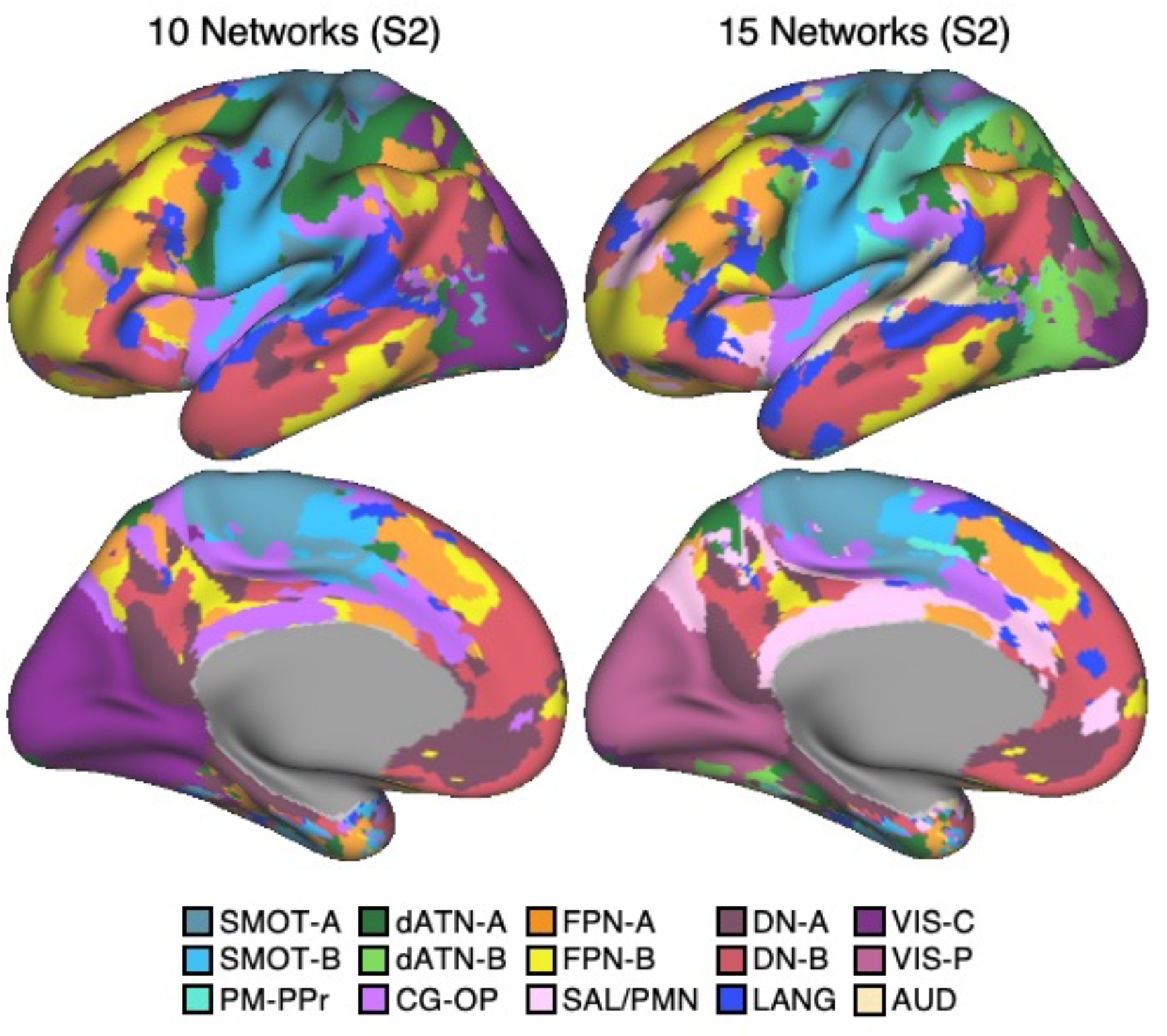
Direct comparison of 10-network and 15-network cerebral cortical parcellations for S2. Paralleling Fig. 7, the left displays the 10-network estimate and the right the 15-network estimate. The network labels are shown at the bottom.

One motivation for investigating a 15-network model was that certain networks did not differentiate established distinctions at or around somatomotor cortex and visual cortex, as well as between multiple networks within or adjacent to the insula including separation of a Cingular-Opercular Network from a Salience Network (Seeley et al. 2007; see Seeley, 2019 for discussion). These features were captured in the 15-network MS-HBM. Specifically, the single visual network in the 10-network estimate was differentiated among dATN-B, VIS-C and VIS-P in the 15-network solution (Fig. 8). The SAL network in the 10-network estimate was differentiated into two separate networks here labeled SAL / PMN and CG-OP (Fig. 12). The dATN in the 10-network estimate was differentiated into dATN-A and PM-PPr in the 15-network solution (Fig. 9), and a distinct AUD network emerged near to LANG and SMOT-B (Fig. 11). Critically, seed-region based correlation patterns suggested that this expansion of networks from 10 to 15 captured clear features of the underlying correlation patterns (Figs. 8, 9, 11, and 12).

**Figure 11.**
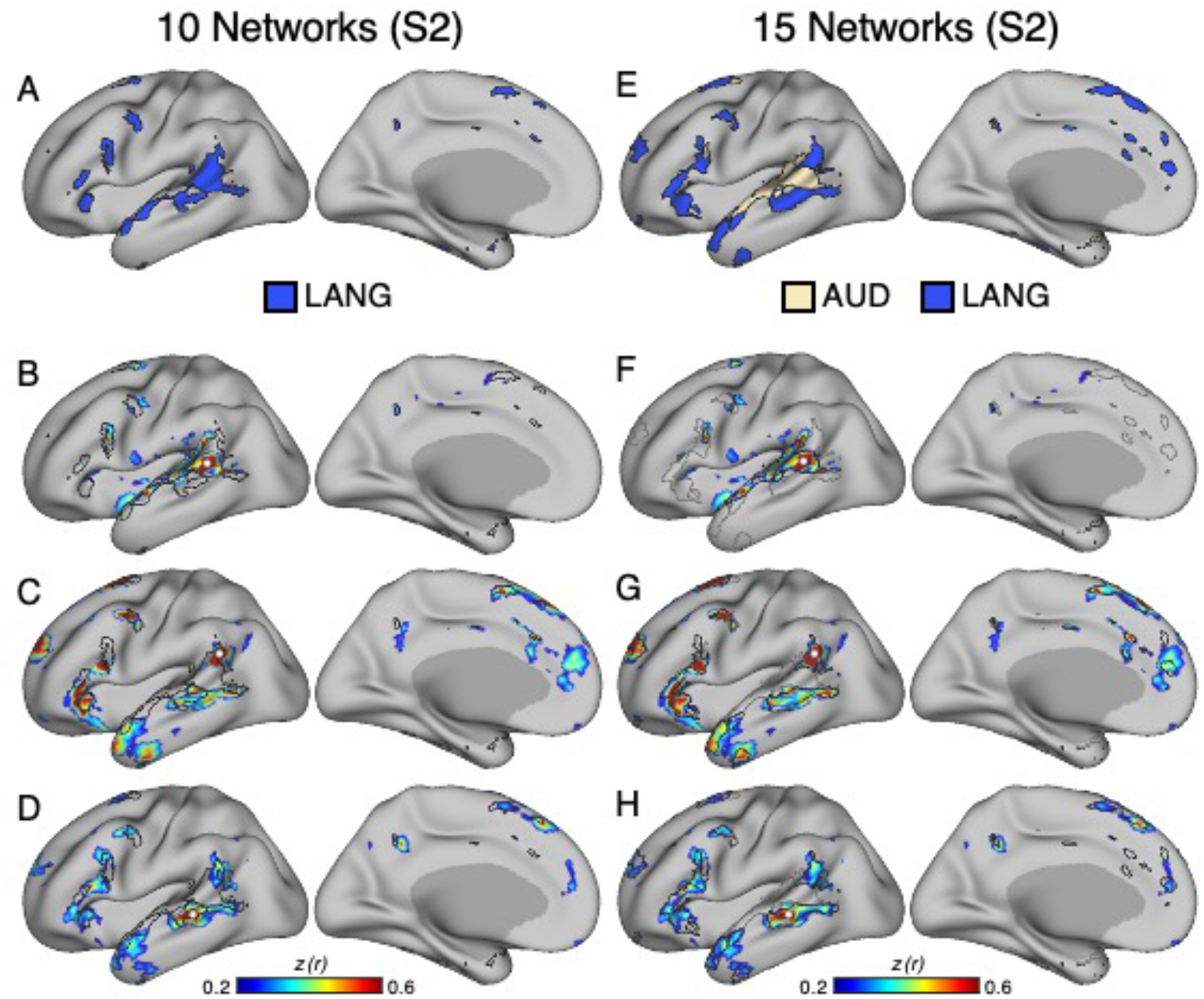
Model-free estimates illustrate the utility of the 15-network cerebral parcellation for auditory and language networks for S2. Seed region correlation maps illustrate features captured by the 15-network estimate as contrast to the 10-network estimate. LANG in the 10-network estimate (**A**) is differentiated into AUD and LANG in the 15-network estimate (**E**). White-filled circles display the seed region locations. Black outlines indicate the boundaries of the networks above. The network labels are shown below. Correlation maps for three distinct seed regions in and around the vicinity of auditory cortex are illustrated within the boundaries of the 10-network estimate (**B, C, D**) and the 15-network estimate (**F, G, H**). Black and gray outlines illustrate the networks from each parcellation estimate. The correlation maps are plotted as z(r) with the color scale at the bottom.

**Figure 12.**
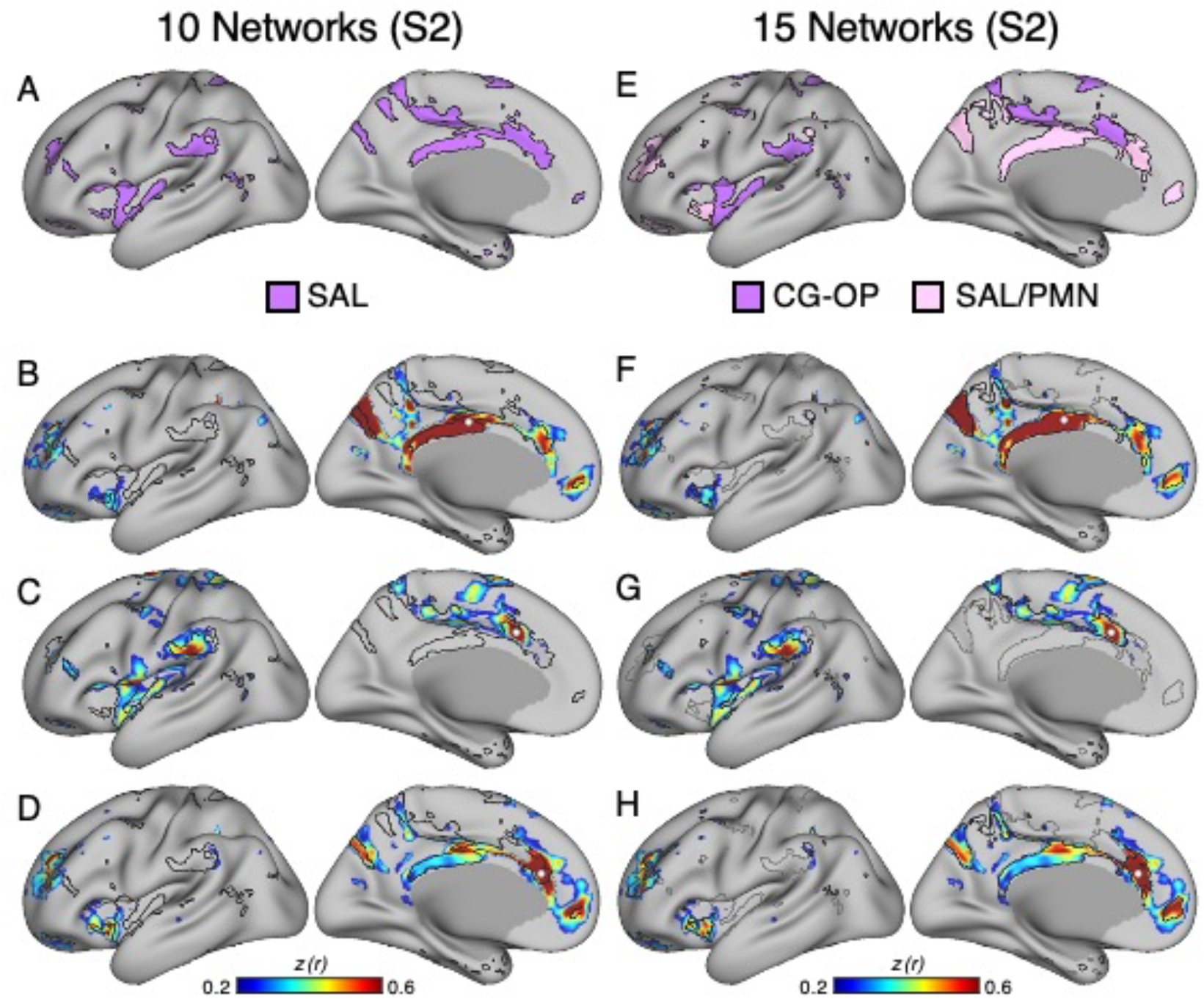
Model-free estimates illustrate the utility of the 15-network cerebral parcellation for networks at and around cingulate cortex for S2. Paralleling Fig. 11, seed region maps illustrate features captured by the present 15-network estimate as contrast to the 10-network estimate. SAL in the 10-network estimate (**A**) is differentiated into the SAL / PMN and the CG-OP networks in the 15-network estimate (**E**). White-filled circles display the seed region locations. Black outlines indicate the boundaries of the networks above. The network labels are shown below. Correlation maps for three seed regions around the cingulate are illustrated within the boundaries of the 10-network estimate (**B, C, D**) and the 15-network estimate (**F, G, H**). Black and gray outlines illustrate the networks from each parcellation estimate. The correlation maps are plotted as z(r) with the color scale at the bottom.

One unexpected result was that our 15-network parcellation included a single network that has been variably described in the literature. What has been called the “Parietal Memory Network” (Gilmore et al. 2015), with focus on the posterior midline, has often been discussed separately from the network referred to by Seeley and colleagues as the “Salience Network” (Seeley 2019). Here a single distributed network was identified that possessed the canonical features of both networks. The seed-region based correlation maps supported that the two networks discussed historically as distinct are likely a single network (Figs. 3, 6, 12), a result that will be further examined in the prospectively acquired and analyzed data.

### Network Estimates Are Reliable Within Individuals

We next sought to address two related questions. First, are the network estimates described above reliable within individuals? Second, can they be obtained with a lesser amount of data? The resting-state fixation runs of S1 and S2 were divided into three datasets with roughly equal amounts of runs contributing to each data subset (20/20/22 runs of data for S1 and 20/20/21 runs of data for S2). The 15-network MS-HBM was estimated independently for each data subset. Results are displayed in Fig. 13.

**Figure 13.**
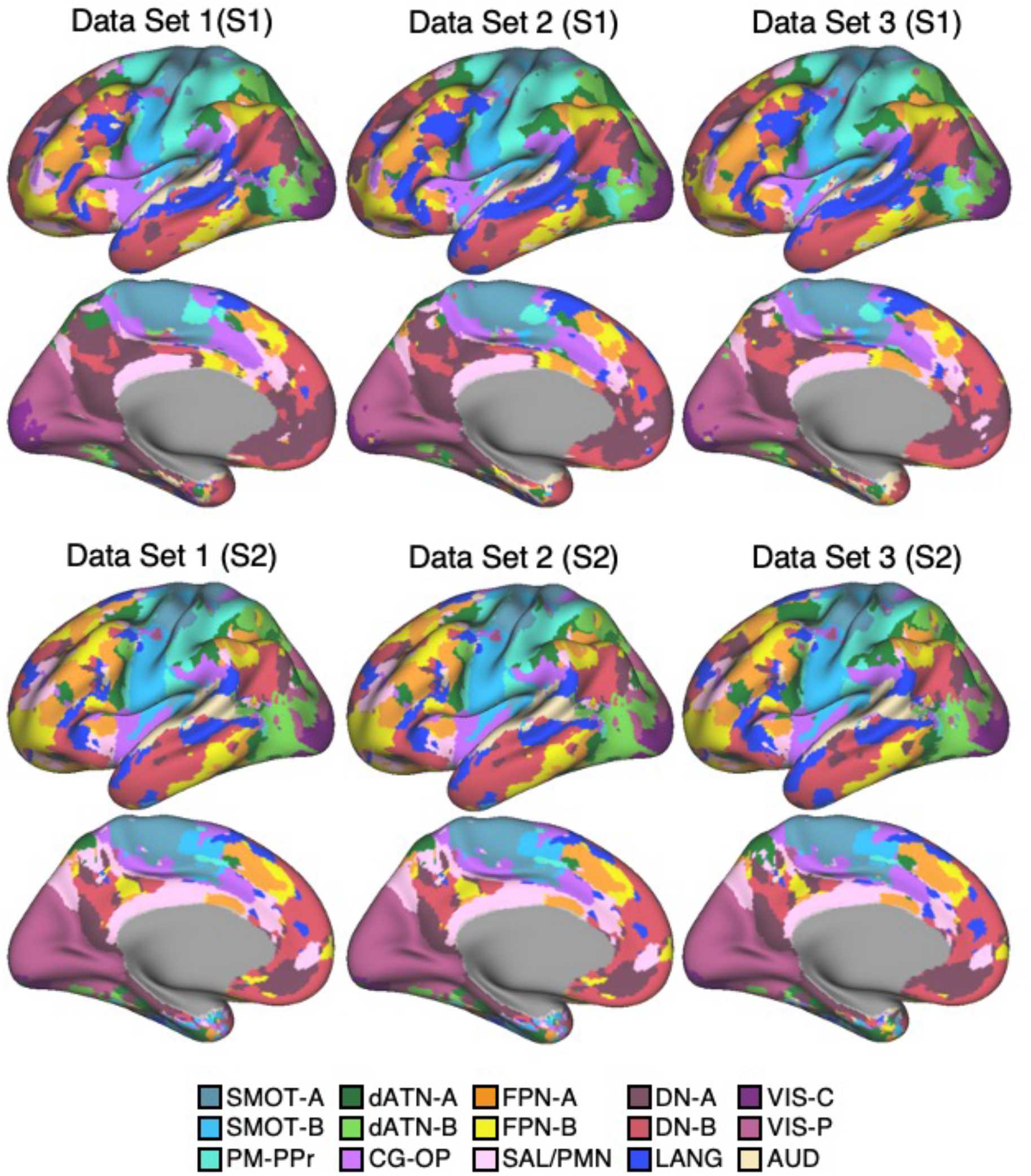
Cerebral cortical network estimates are reliable across independent datasets within individuals. Independently analyzed subsets of data from S1 (**Top**) and S2 (**Bottom**) illustrate the reliability of the network estimates. The resting-state fixation data of S1 and S2 were split into three datasets to estimate networks using the MS-HBM applied independently to each dataset. The individual-specific cortical parcellations are replicable within participants, critically for models based on ∼20 runs of resting-state fixation data as will be employed for the 15 new participants analyzed throughout the remainder of this paper. The network labels are shown at the bottom.

In S1, 84.2% of cortical vertices were assigned to the same networks across the three independent datasets from within the individual. In S2, 88.0% of cortical vertices were assigned to the same networks. By contrast, overlap between the separate parcellations of S1 and S2 were 58.3%, 58.9% and 59.2%, indicating that between-individual variability was substantially larger than within-individual variability.

These findings suggest that cortical parcellations of the resolution and within-individual detail targeted here are replicable for models based on ∼20 runs of data. Notably, this is the amount of data collected for the 15 new participants in the implementation stage dataset analyzed throughout the rest of this paper.

### Network Estimates in 15 New Participants Reveal Organizational Features

#### Discovery, Replication and Triplication in the Implementation Stage Data

15 cerebral networks were estimated for all new participants. The 15 individuals were analyzed within subsamples (each n = 5) intended to replicate the MS-HBM’s network estimates in prospective participants including novel discovery (P1-P5), replication (P6-P10) and triplication (P11-P15) datasets. Results were similar across all three subsamples, and the full parcellation for each individual is available in the Supplementary Materials on the surface and within the individual’s own native-space volume. Despite idiosyncratic spatial details of network organization, the broad properties were largely consistent. Three representative participants, one from each subsample, are displayed in Figs. 14 to 16.

**Figures 14-16.**
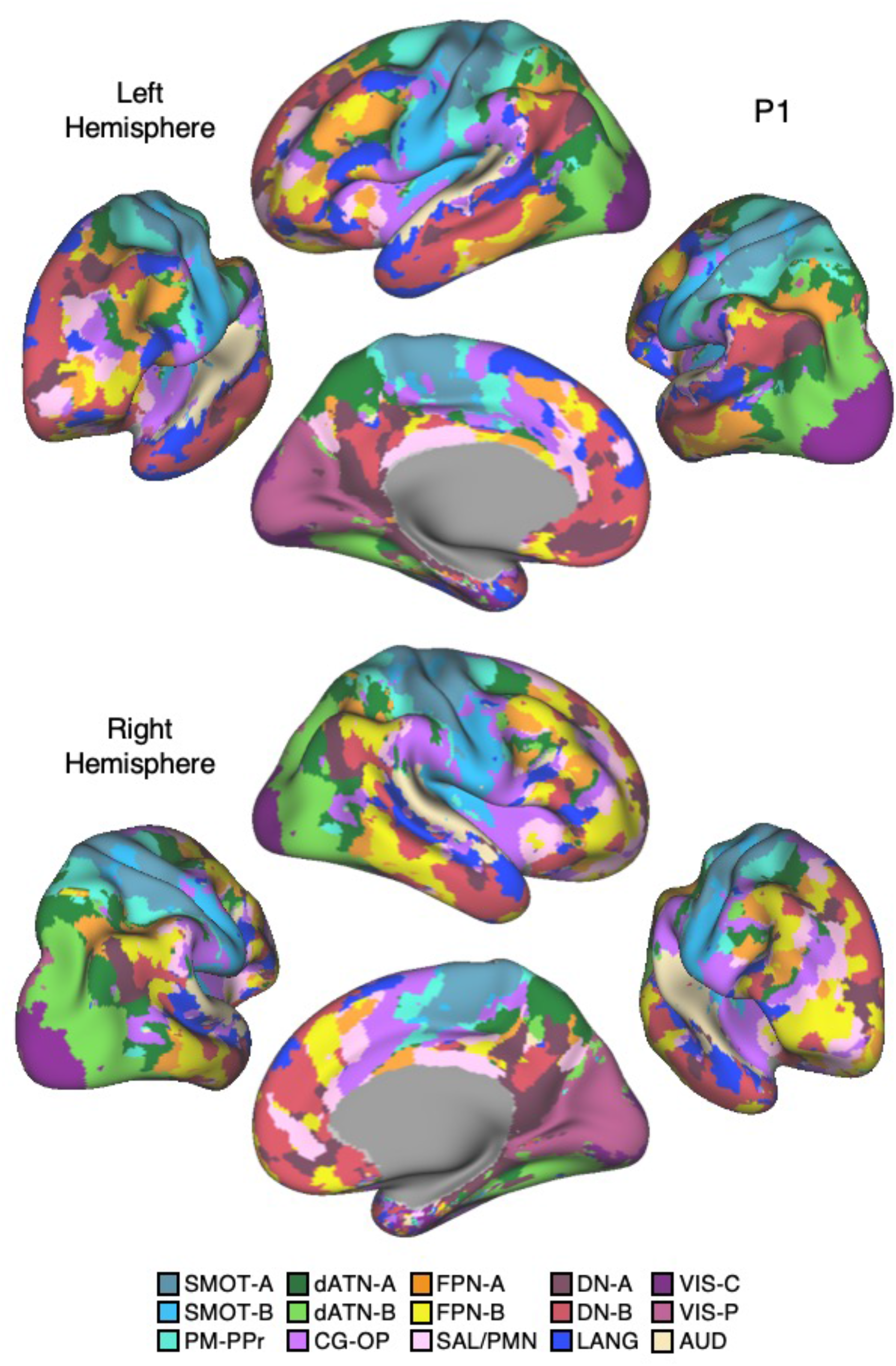
Network estimates for novel participants. Networks estimated for representative participants from the novel discovery (P1), replication (P6) and triplication (P11) datasets are displayed. The network estimates are from the 15-network MS-HBM. Four views for each hemisphere show details of cortical network organization, with lateral and medial views as well as rotated frontal and posterior views. The left hemisphere is on top and right hemisphere below. Each color represents a distinct network with the network labels shown at the bottom. Similar maps for all available participants are provided in the Supplementary Materials.

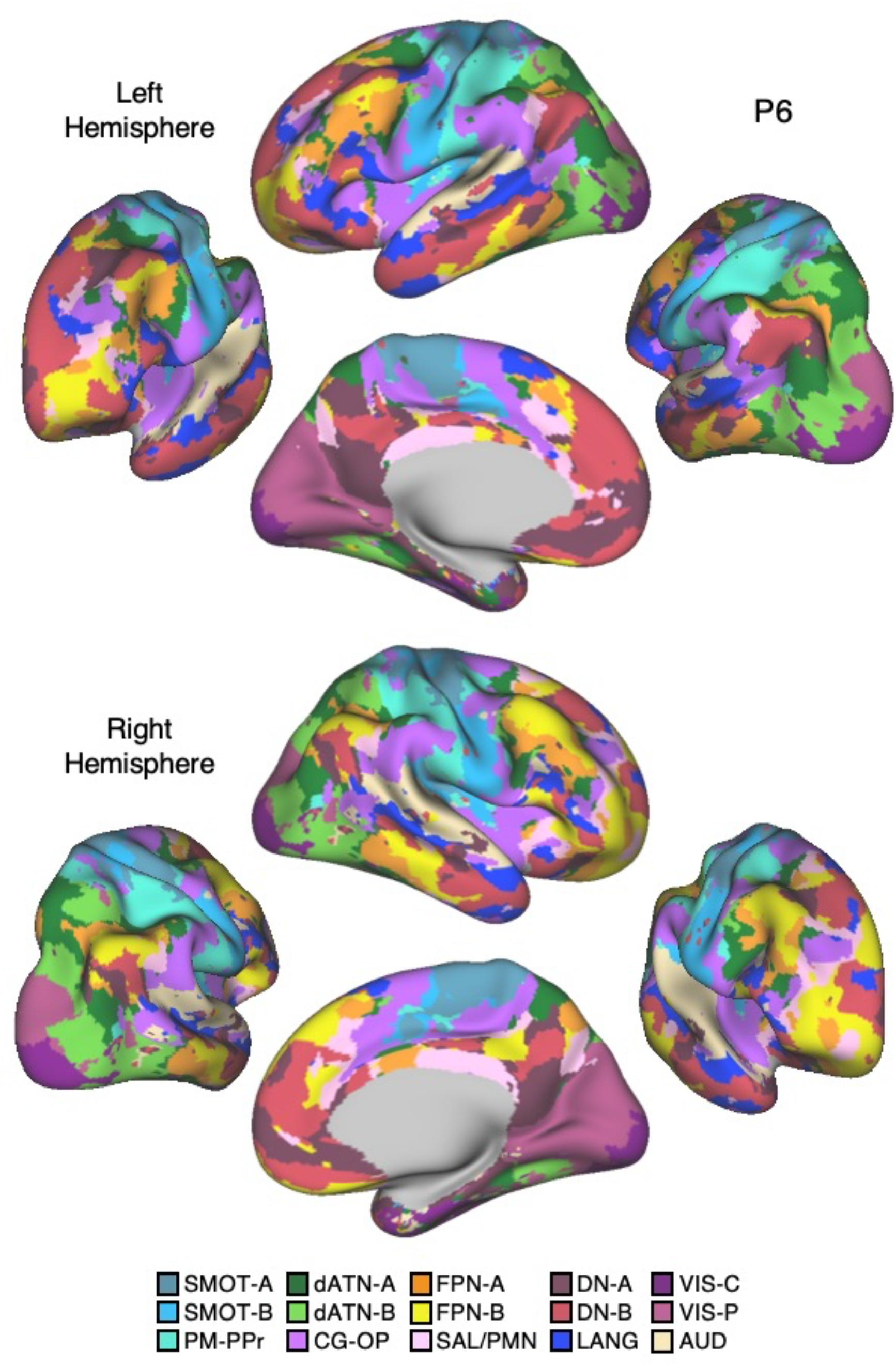

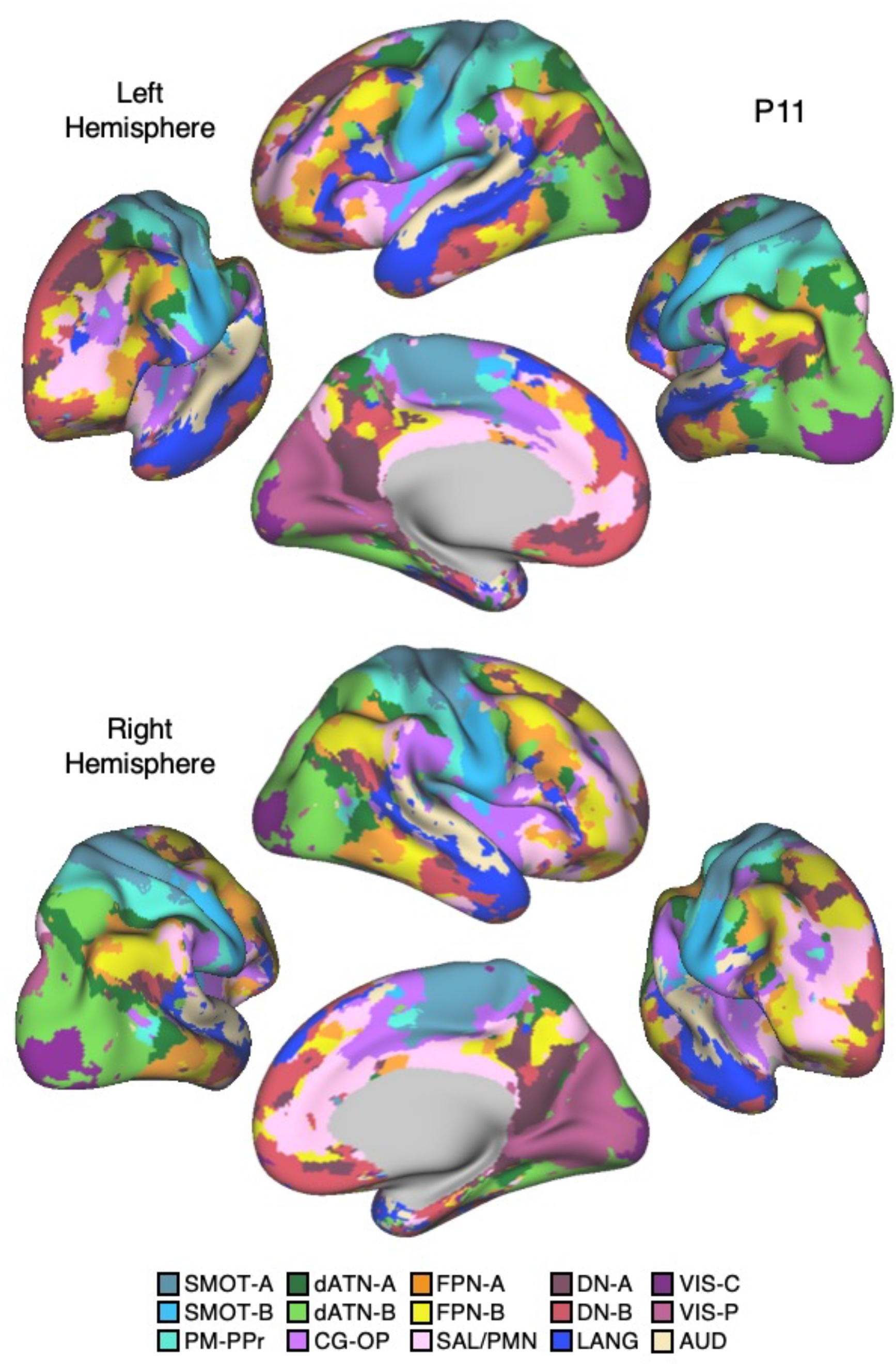

#### Network Estimates Reveal Predominantly Local Sensory and Motor Networks

VIS-C and VIS-P were identified in each participant extending from the calcarine sulcus on the midline to the lateral surface. The extent of the two networks across the occipital lobe did not align them with individual visual areas, but rather the expanded regions of the V1/V2/V3 retinotopic cluster, and likely adjacent retinotopic clusters (Wandell et al. 2005; Wandell, Domoulin, Brewer 2007). The multiple networks appeared to divide along the eccentricity gradient (Buckner and Yeo 2014). The VIS-C network overlapped regions likely aligned to the central portions of the V1/V2/V3 retinotopic representations, while VIS-P overlapped the peripheral retinotopic representations (see Yeo et al. 2011). The relation of VIS-C and VIS-P to task-elicited responses is directly explored in a later section.

While the VIS-C and VIS-P networks contained vertices that were mostly contiguous, there were exceptions. Discontinuous islands were sometimes found in occipital-temporal cortex, possibly a reflection of separate extrastriate retinotopic clusters (e.g., at or near the MT/V5 hemifield representation). VIS-P also occasionally contained small, punctate representations near to dorsolateral prefrontal cortex (DLPFC). These were the exceptions: the majority of the VIS-C and VIS-P networks’ included vertices were continuous and adjacent to one another, overlapping the expected location of early retinotopic visual cortex.

Similarly, SMOT-A and SMOT-B were identified reliably as spatially continuous networks along the central sulcus, extending onto the midline and into the posterior insula. These two somatomotor networks also do not likely align to individual architectonic areas, but rather extend across the pre- and post-central sulcus including primary motor as well as somatosensory areas. The extent along the midline and into the posterior insula further suggests the networks span multiple body maps, not simply the dominant inverted body map along the central sulcus. The separation into two networks is consistent with separation of distinct portions of the somatotopic map along the body axis, a functional hypothesis that will also be directly tested in a later section.

A final predominantly local sensory network, AUD, was consistently identified near to the superior temporal sulcus. This network extended across the full supratemporal plane including Heschl’s gyrus, and into adjacent regions.

#### Multiple Distributed Networks Lie Adjacent to the Local Sensory and Motor Networks

Multiple distributed networks were identified in each participant that were immediately adjacent to the local sensory and motor networks, with each network containing distributed regions that spanned multiple zones of cortex. dATN-A and dATN-B were adjacent to VIS-C and VIS-P but also with distant regions in the frontal cortex, likely at or near the frontal eye field (FEF) (Corbetta & Shulman 2002; Hutchinson et al. 2012). Similarly, CG-OP and PM-PPr radiated outwards from the early somatomotor networks SMOT-A and SMOT-B. CG-OP and PM-PPr sometimes contained small islands indenting or even within the SMOT network boundaries that may relate to interspersed inter-effector regions along the central sulcus (Gordon et al. 2023). CG-OP and PM-PPr also included regions abutting and within the Sylvian fissure. The relations among the networks will become even clearer in the upcoming flat map visualizations.

A final network, SAL / PMN, displayed a spatial pattern that was adjacent to CG-OP in many locations but also with differences. While SAL / PMN contained a prominent region in the anterior insula, the network’s positioning did not juxtapose the somatosensory networks. Rather, SAL / PMN was adjacent to a posterior midline cluster of association networks near to regions of the canonical “Default Network” (e.g., Shulman et al. 1997; Buckner et al. 2008; Power et al. 2011; Yeo et al. 2011). SAL / PMN consistently included a region within ACC anterior to CG-OP and a prominent set of regions along the mid-cingulate and the posterior midline. As noted for S1 and S2, the SAL / PMN network’s spatial pattern combined features described in prior work on the Salience Network (Seeley et al. 2007; see Seeley 2019 for discussion and Dosenbach et al. 2006 for related work) and the Parietal Memory Network (Gilmore et al. 2015).

#### Much of Association Cortex is Populated by Multiple Parallel Juxtaposed Networks

The remaining regions of association cortex -- that contain the majority of PFC, a large region of PPC extending into the temporoparietal junction (TPJ) and lateral temporal cortex (LTC) -- were populated by five distinct networks. With some exceptions, each of these five networks tended to possess regions in each of the distributed zones. The five networks were interwoven with local patterns of adjacencies that repeated across cortex.

Specifically, FPN-A and FPN-B were adjacent to one another throughout the cortical mantle. FPN-A and FPN-B displayed a distributed pattern consistent with the well-studied group-estimated Frontoparietal Control Network, also referred to as the Multiple-Demand System (Duncan et al. 2010; Power et al. 2011; Yeo et al. 2011). These two juxtaposed networks (FPN-A and FPN-B) consistently neighbored an additional clustered set of three networks – LANG, DN-B, and DN-A. These three additional networks were tightly juxtaposed among themselves on the lateral cortical surface including zones within PPC, LTC, and both DLPFC and VLPFC. DN-A and DN-B were interdigitated as well along the anterior and posterior midline, consistent with previous studies (Braga and Buckner 2017; Gordon et al. 2017; Braga et al. 2019).

Despite being adjacent in many locations, clear features distinguished the three networks. The LANG network surrounded the Sylvian fissure and included regions near to the AUD network and in VLPFC at or near historically defined “Broca’s area”. The LANG network was generally larger in the left hemisphere compared to the right (but see Braga et al. 2020 for an exception). DN-A showed a strong correlation with the posterior parahippocampal cortex (PHC) (see also Reznick et al. 2023 for further details). Additionally, DN-A occupied regions at or adjacent to the retrosplenial cortex (RSC) and ventral posterior cingulate cortex (PCC). DN-B prominently included anterior regions of the inferior parietal lobule extending into the TPJ (a region of particular focus, e.g., Saxe and Kanwisher 2003; Jacoby et al. 2016). The posterior midline region of DN-B fell between regions of DN-A and specifically did not extend into RSC. DN-B also included a larger region of the LTC than DN-A; DN-A tended to include a small region or a few discontinuous regions in anterior LTC. Of importance, the spatial arrangements of the five networks (FPN-A, FPN-B, LANG, DN-B, and DN-A) repeated multiple times across the cortical mantle, a discovery that will be expanded upon in the analyses of spatial juxtapositions on the flattened cortical surface.

### A Cautionary Note About Potential Artifacts

Certain aspects of the network estimates were impacted by signal loss. Low SNR regions were observed in the OFC, ventral regions of VLPFC, and anterior regions of the temporal lobe (see Figs. 1 and 4, and Supplementary Materials). When interpreting the network assignments, it is important to keep these spatially variable effects in mind. For example, a localized AUD network was observed across the supratemporal plane including Heschl’s gyrus. Inconsistent, discontinuous vertices were also labelled as part of the AUD network in the inferior temporal cortex and OFC, in the regions of the greatest signal dropout due to magnetic susceptibility differences. The network assignments in low SNR regions should not be trusted.

### Model-Free Seed-Region Based Correlations Again Confirm Network Estimates

To demonstrate that the correlation properties of the within-individual data were captured in the network assignments, seed-region based correlation maps were visualized. While there were minor differences between the MS-HBM network estimates and the seed-region based correlation maps, networks could be identified in all participants using both methods. Furthermore, the maps defined by anterior and posterior seed regions were similar, indicating that the seed-region based method was not dependent on a single vertex or one general region of cortex. Seed-region based confirmation for DN-A, DN-B, LANG, FPN-A, FPN-B, CG-OP, and SAL / PMN are displayed for three representative participants in Figs. 17 to 19, and for all participants in the Supplementary Materials.

**Figures 17.**
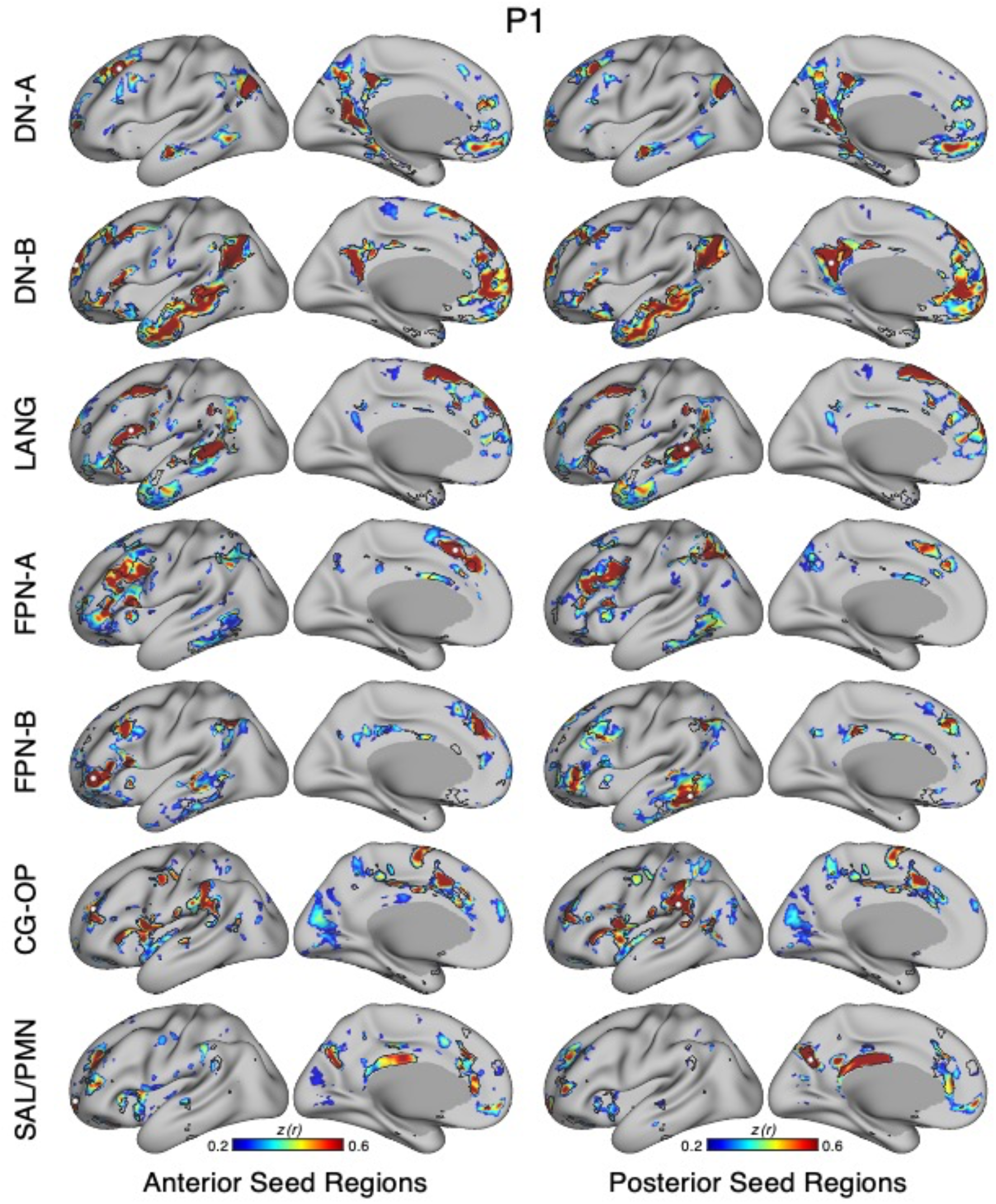
Model-free confirmation of networks using seed-region based correlation for the implementation stage participants. The correlation patterns from individual seed regions placed within networks are displayed for representative participants from the novel discovery (P1), replication (P6) and triplication (P11) datasets. The two left columns display correlation maps using an anterior seed region for each network, while the two right columns display correlation maps using a posterior seed region. Lateral and medial views are displayed for each seed region. Black outlines indicate the boundaries of corresponding individual-specific parcellation-defined networks estimated from the MS-HBM as shown in Figs. 14-16. The correlation maps are plotted as z(r) with the color scale at the bottom. Strong agreement is evident between the seed-region based correlation maps and the estimated network boundaries. Similar maps for all available participants are provided in the Supplementary Materials.

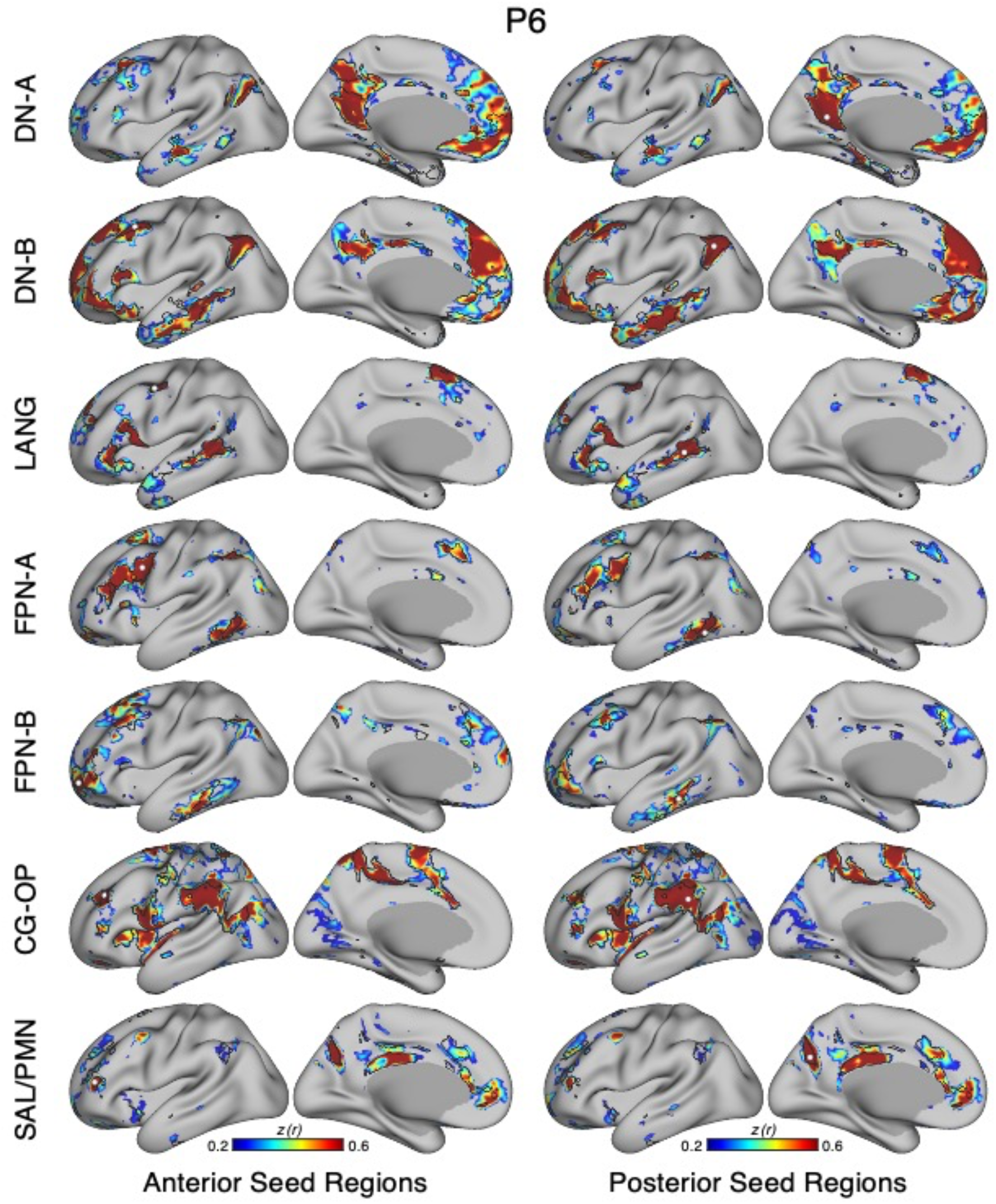

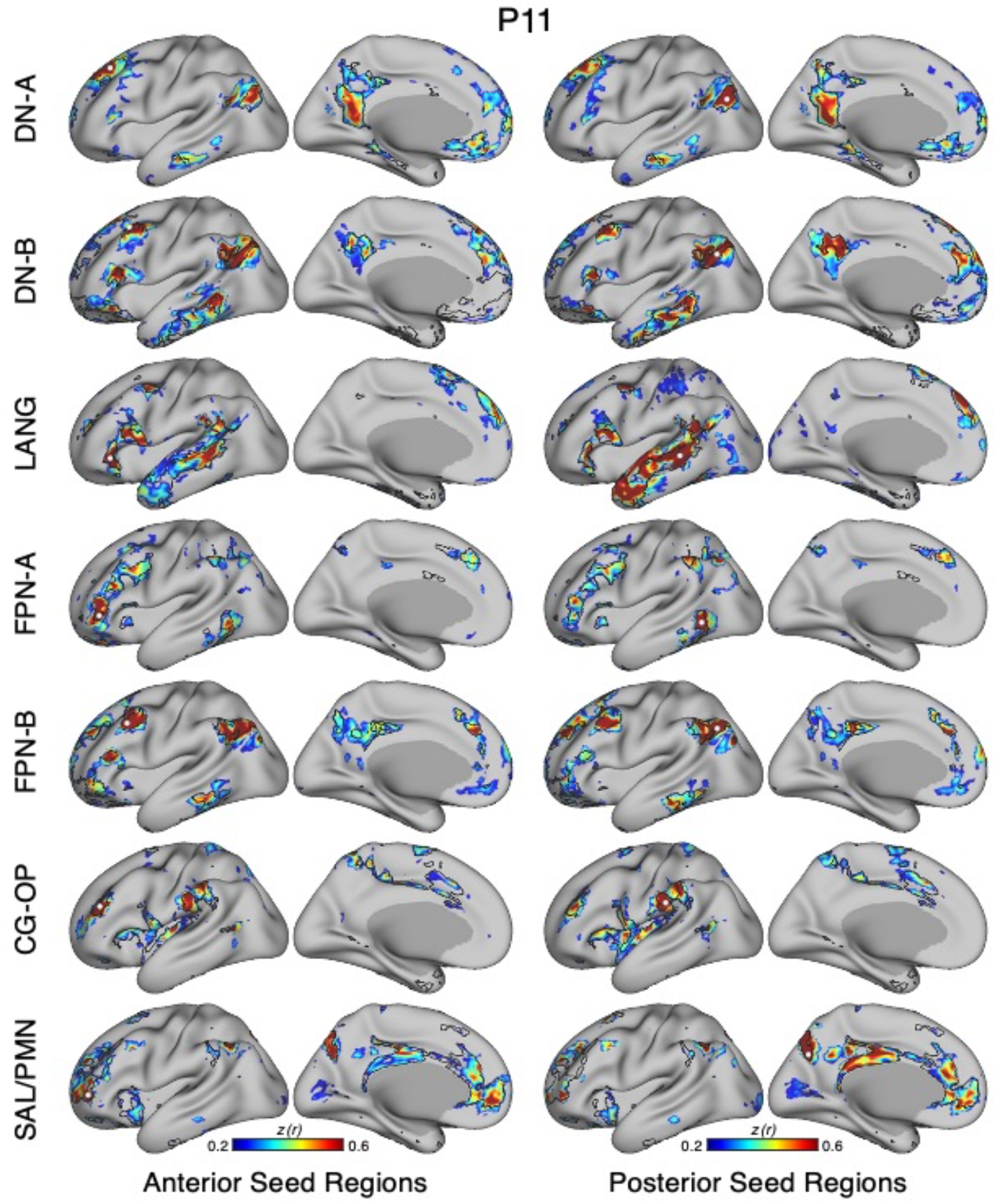

### Variability in Network Estimates Across Individuals

An overlap map of assignments for each network from the MS-HBM model for the 15 participants is displayed in Fig. 20. Results revealed that the general organization of the networks was highly conserved across individuals, but with differences in the idiosyncratic spatial positioning and extents of the networks.

**Figure 20.**
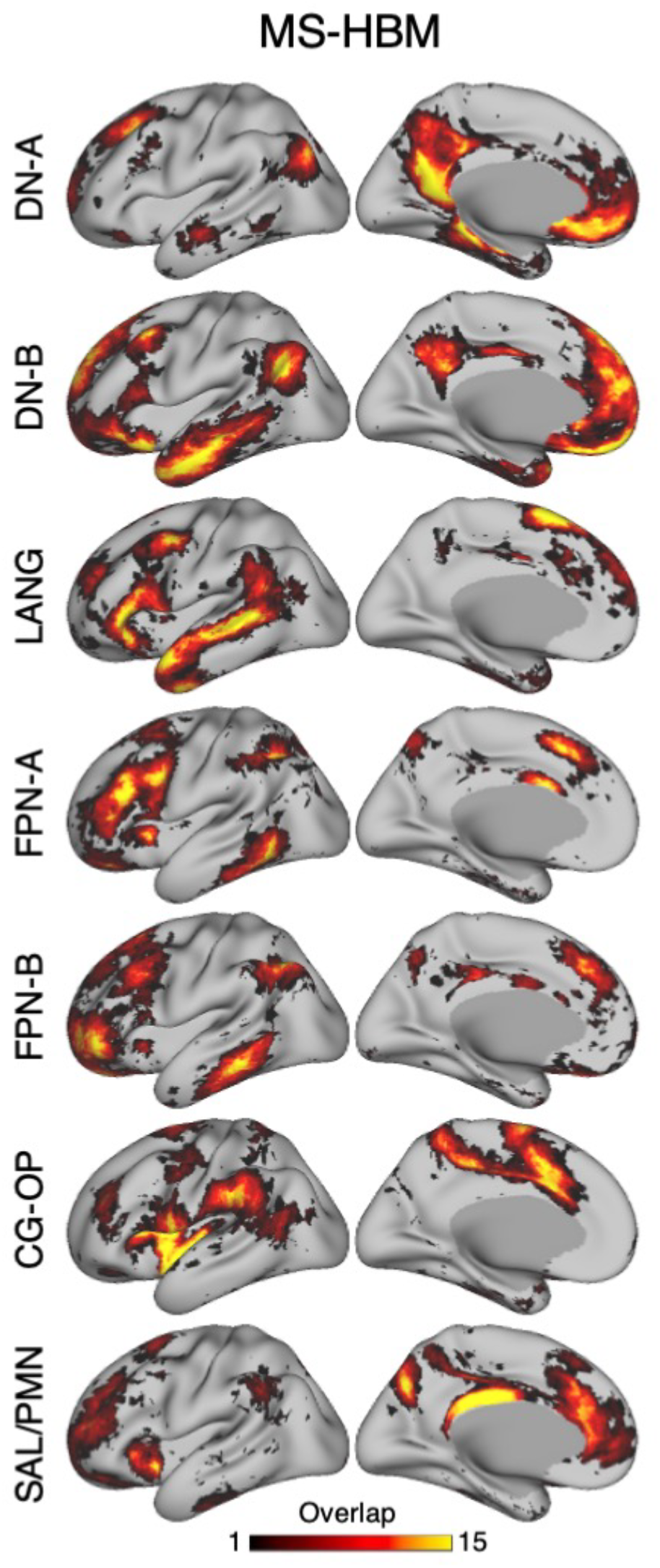
Overlap of network estimates derived from the MS-HBM model. Each row displays the overlap map from one target network for the full set of 15 novel participants using the estimates from the 15-network MS-HBM. The network targets are labeled to the left. DN-A, DN-B, LANG, FPN-A, FPN-B, CG-OP, and SAL / PMN networks are examined separately. The purpose of these maps is to illustrate the overlap of network organization across participants as well as illustrate how the separate networks are distinct from one another.

A challenge in examining spatial overlap is that there is circularity in network definition since the process initiates with the same 15-network group prior, which could bias the networks to show more overlap than is truly in the data. To mitigate this concern, we also examined overlap using the network estimates derived from the seed-region correlation maps. These maps are not constrained by the group prior and do not enforce a winner-take-all assumption, allowing deviations to emerge. Overlap maps of correlation patterns were obtained using anterior and posterior seed regions within each network for all 15 participants (Fig. 21).

**Figure 21.**
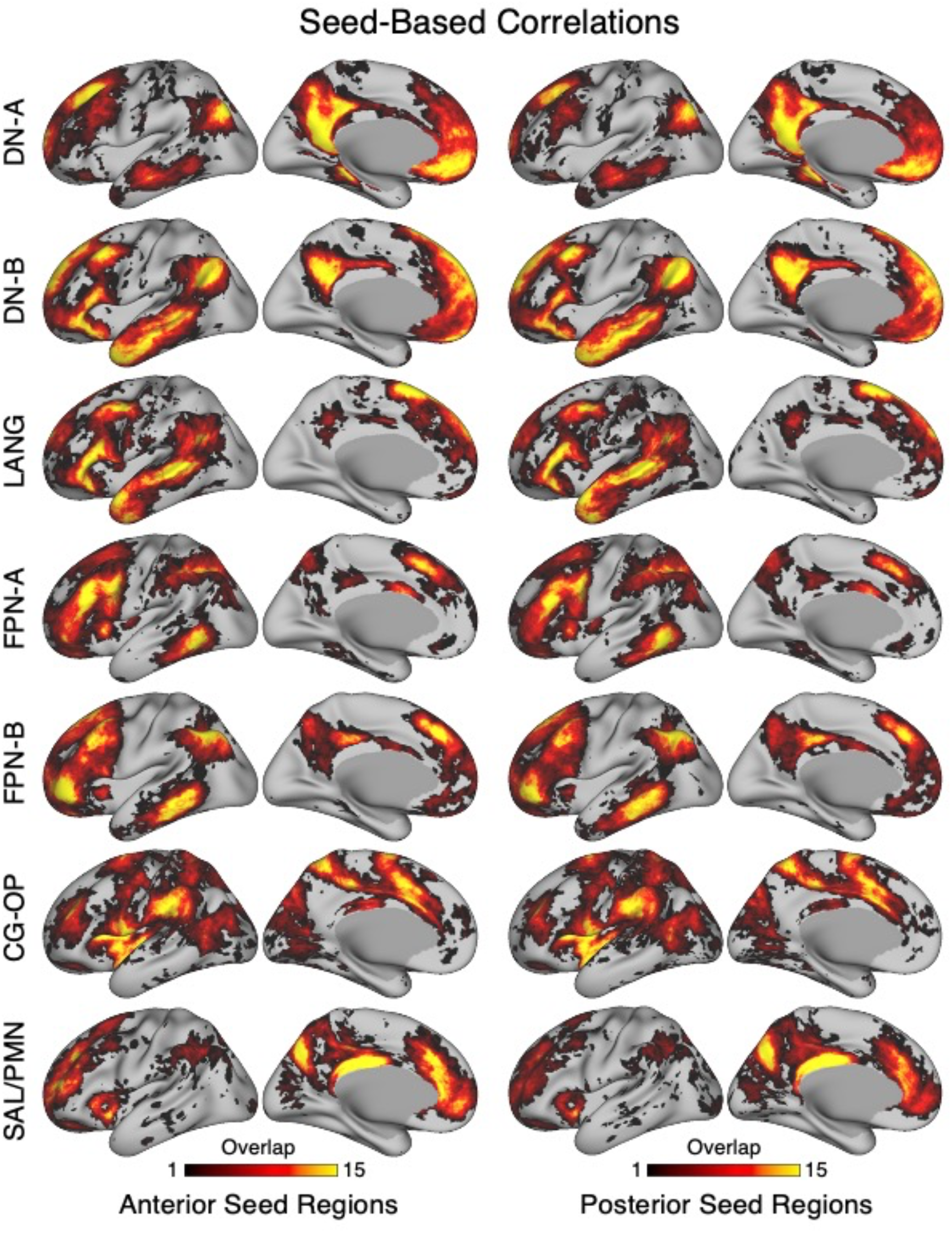
Overlap of network estimates derived from model-free seed-region correlation maps. Each row displays the overlap map from one target network for the full set of 15 novel participants using only seed-region based correlation estimates of the networks. In the left two columns, each row displays the overlap map of correlation patterns based on an anterior seed region. In the right two columns, each row displays the overlap map based on a posterior seed region. The network targets are labeled to the left. DN-A, DN-B, LANG, FPN-A, FPN-B, CG-OP, and SAL / PMN networks are examined separately. The purpose of these maps is to illustrate the overlap of network organization without strong model assumptions (priors) that might bias the degree of overlap.

**Figure 22.**
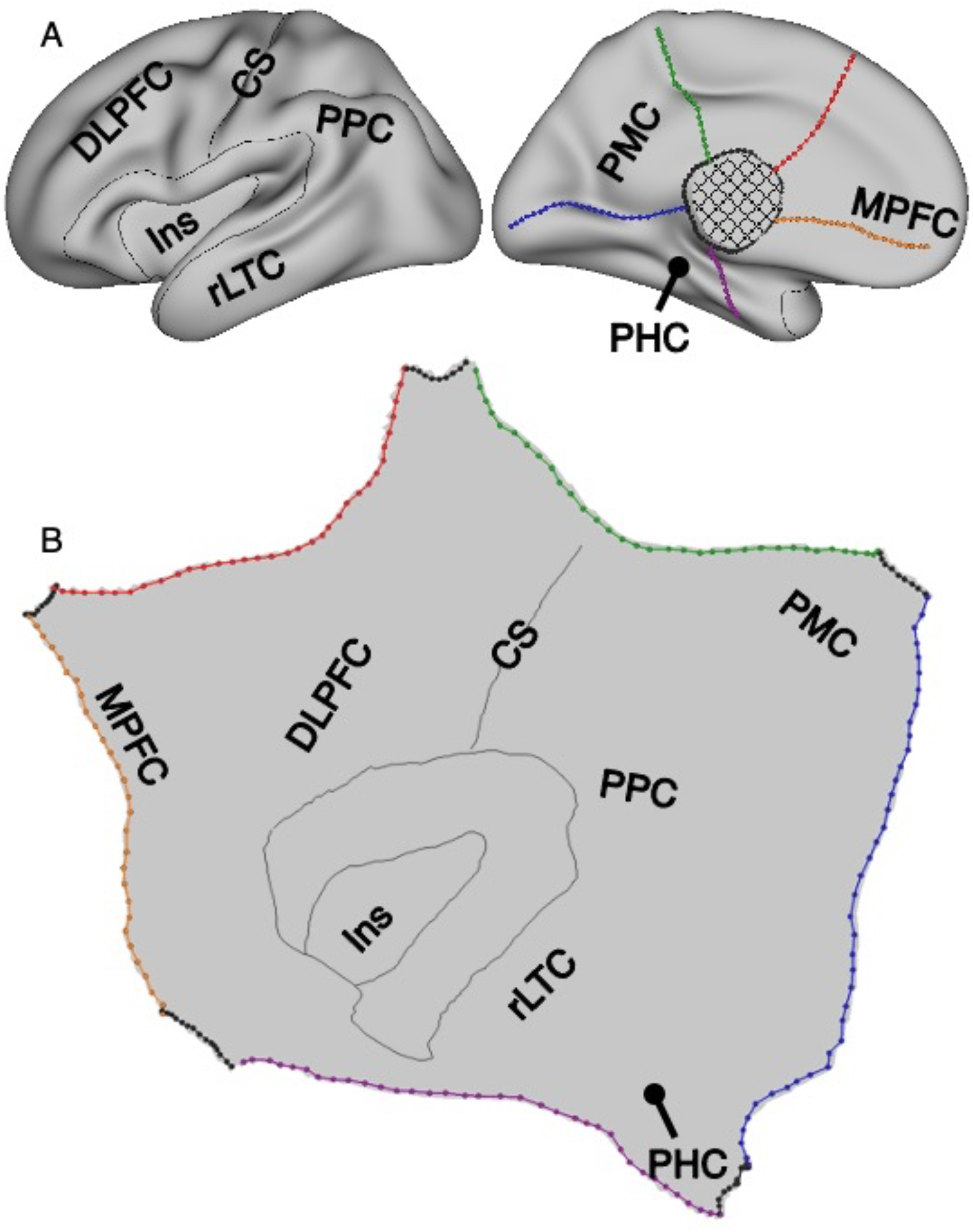
Visualization on the flattened cortical surface. A fully flattened cortical surface was constructed to better reveal topographic relations among networks. By applying five cuts along the colorful lines on the midline, the inflated cortical surface (**A**) was flattened (**B**). The five cuts included one cut along the calcarine sulcus (blue dotted line) and four additional cuts radiating outwards from the medial wall. The surface enclosed by the circular cut was removed. Reference lines illustrate the inner and outer boundaries of the insula (Ins) as well as along the central sulcus (CS). Additional landmarks are dorsolateral PFC (DLPFC), posterior parietal cortex (PPC), rostral lateral temporal cortex (rLTC), posteromedial cortex (PMC), parahippocampal cortex (PHC), and medial PFC (MPFC). The procedure was applied separately to the two hemispheres.

As a final exploration of variability, the individual networks were plotted next to one another for all 15 participants, allowing another means to identify shared and idiosyncratic features of each participant’s estimate. The results of this final analysis are available in the Supplementary Materials.

### Higher-Order Networks Nest Outwards from Primary Cortices

To better reveal spatial relations among networks, a flattened cortical surface was constructed (Fig. 22). The 15 networks are displayed in representative participants in Figs. 23 to 25, and for all participants in the Supplementary Materials. A broad observation was that higher-order networks nest outwards from sensory and motor cortices.

**Figures 23-25.**
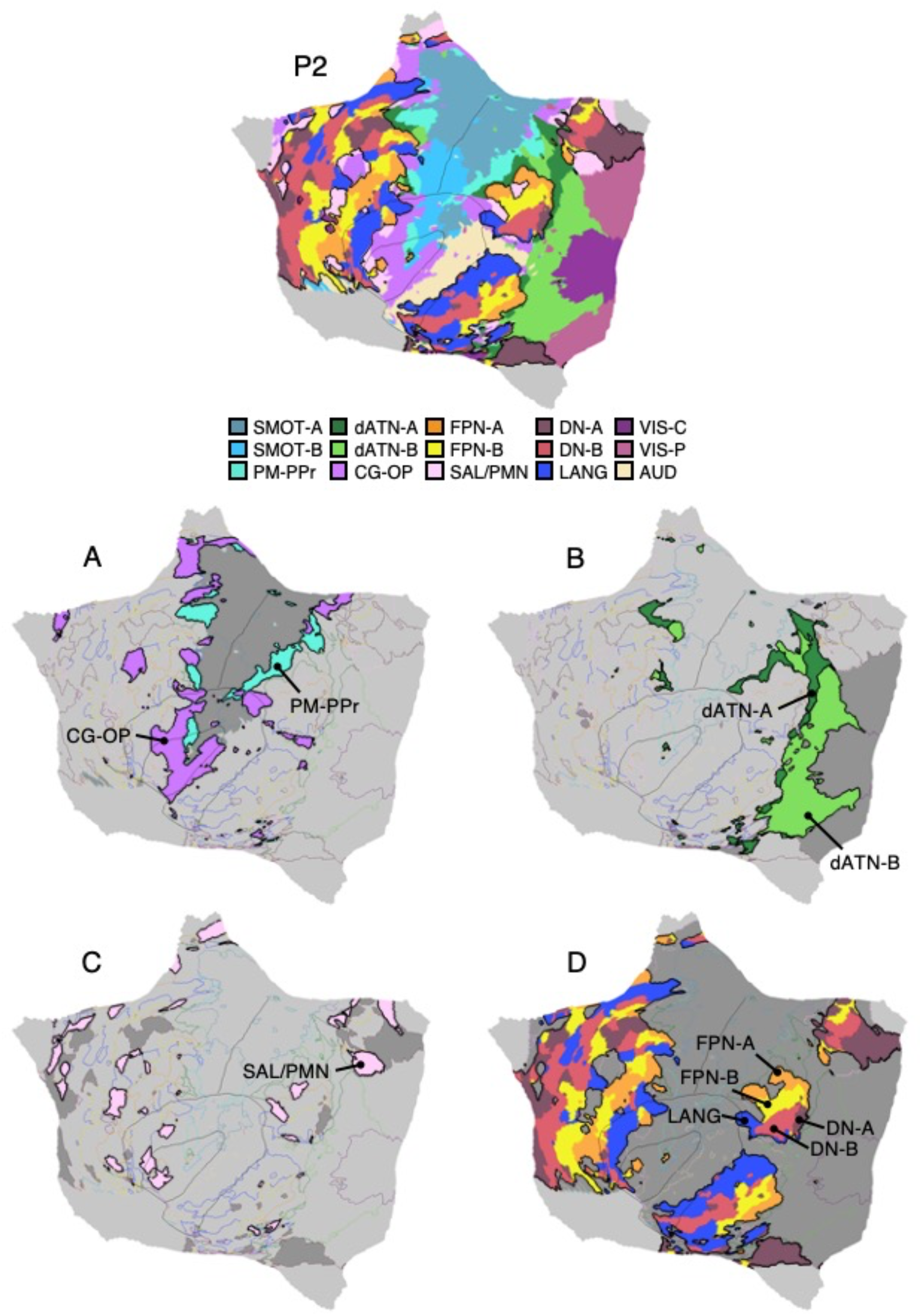
Higher-order networks nest outwards from sensory and motor cortices. Networks displayed on the flattened cortical surface reveal orderly spatial relations in representative participants from the novel discovery (P2), replication (P6) and triplication (P12) datasets. The top map displays all networks estimated using the MS-HBM. The maps below show subsets of networks to highlight spatial relations. (**A**) Somatomotor networks SMOT-A and SMOT-B, in dark gray, are surrounded by spatially adjacent second-order networks CG-OP and PM-PPr. The second-order networks are more distributed than the first-order SMOT-A and SMOT-B networks, which are primarily locally organized. (**B**) Visual networks VIS-C and VIS-P, in dark gray, are surrounded by spatially adjacent second-order networks dATN-A and dATN-B, that possess distributed organization. (**C**) The SAL/PMN network has a widely distributed organization, that includes adjacency to DN-A, shown in gray, especially along the posterior midline. (**D**) The distributed association zones that fall outside of the first- and second-order networks are illustrated. These zones are populated by five distinct networks (DN-A, DN-B, LANG, FPN-A and FPN-B) that possess repeating spatial adjacencies across the cortex, most clearly visible in posterior parietal association cortex and temporal association cortex. FPN-A and FPN-B are adjacent to one another, and together adjacent to the three other juxtaposed networks LANG, DN-B and DN-A. We call these repeating clusters of networks Supra-Areal Association Megaclusters (SAAMs) and explore them further in later analyses. The network labels in (**D**) are positioned around the SAAM in posterior parietal cortex. The network labels are defined in Fig. 2. Similar maps for all available participants are provided in the Supplementary Materials.

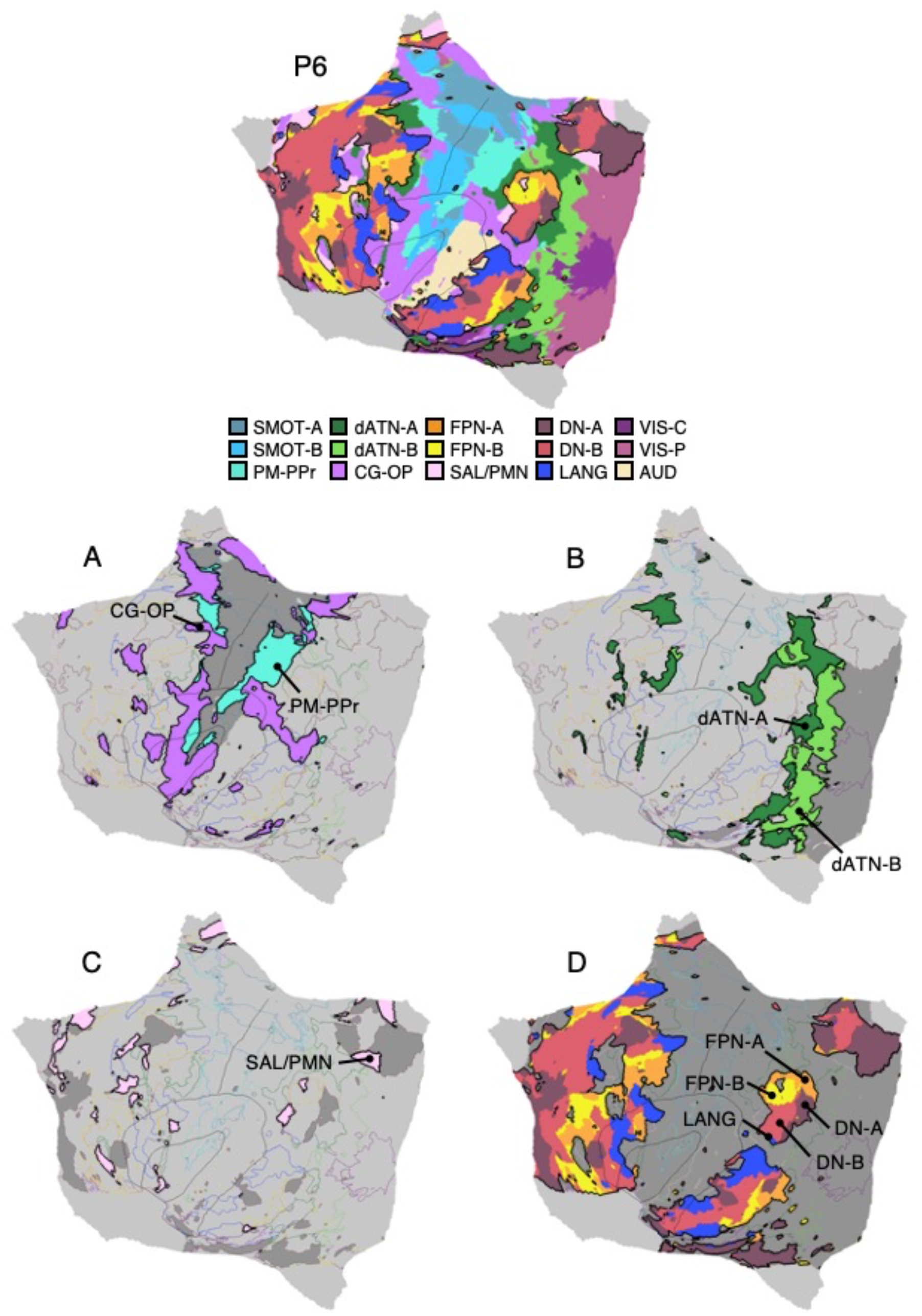

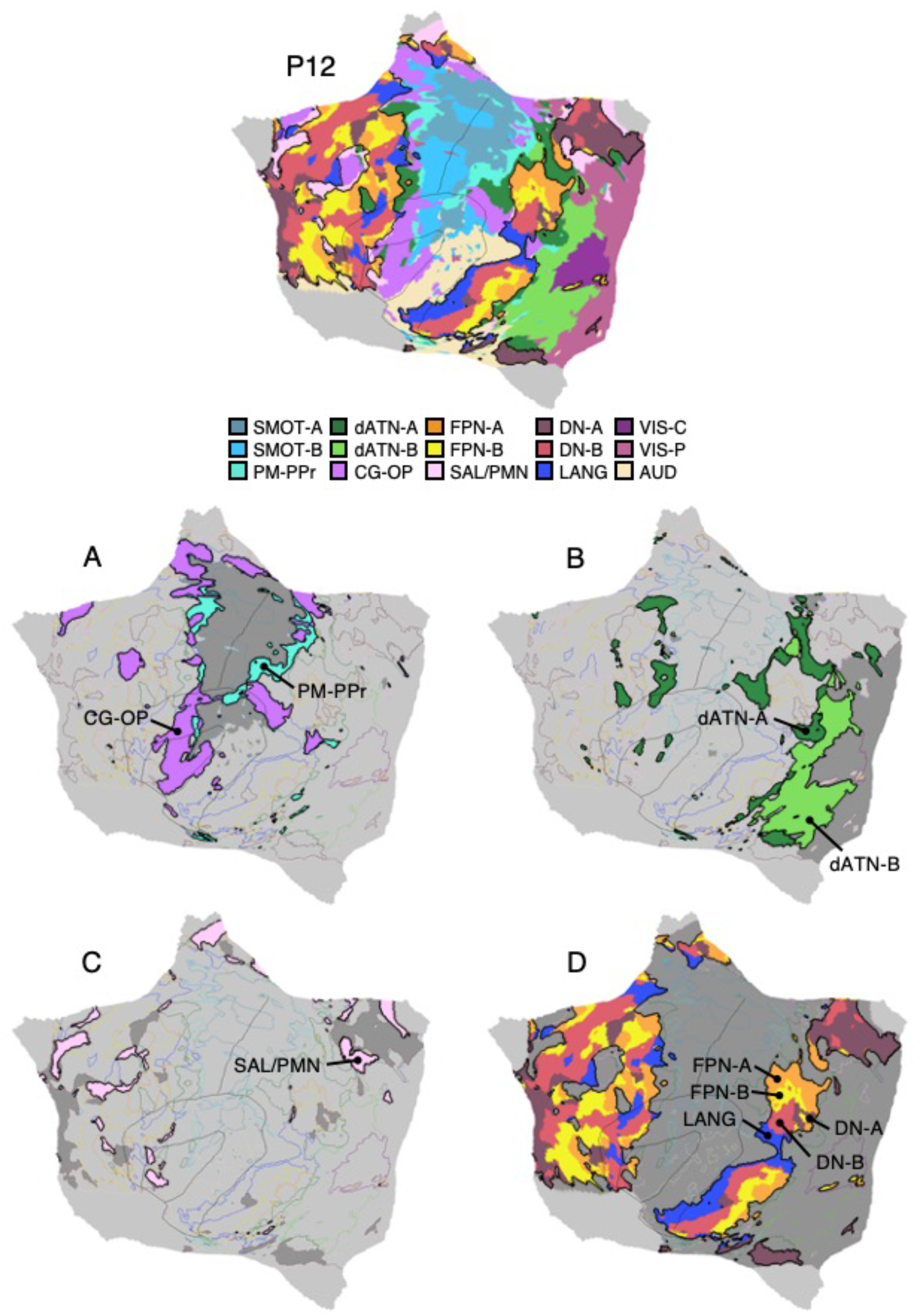

Specifically, the networks could be heuristically grouped into three levels beginning with first-order sensory and motor networks^6^. The first-order networks were primarily locally organized, spatially arranged along the central sulcus (for SMOT-A and SMOT-B) and near to the calcarine sulcus (for VIS-C and VIS-P). Surrounding these first-order networks were adjacent networks that radiated outwards. We refer these as second-order networks. CG-OP and PM-PPr surrounded SMOT-A and SMOT-B, and dATN-A and dATN-B were adjacent to VIS-C and VIS-P. In between these second-order networks were third-order networks (FPN-A and FPN-B, LANG, DN-B, DN-A) that populated the large, expanded zones of higher-order association cortex. The flattened representation allowed further features to be appreciated.

CG-OP and PM-PPr nearly fully surrounded both the anterior and posterior extents of the somatomotor networks, including the insular regions that are buried within the Sylvian fissure. While CG-OP and PM-PPr were generally interdigitated around the first-order somatomotor networks, in several locations CG-OP fell distal to PM-PPr (meaning PM-PPr directly juxtaposed SMOT-A and SMOT-B and CG-OP juxtaposed PM-PPr). Furthermore, while the PM-PPr network was adjacent to the somatomotor networks across its extent, CG-OP also involved distant regions in PFC and posterior association zones that were not adjacent to somatomotor networks. Thus, CG-OP possessed a partially distributed motif. Additional details of dATN-A and dATN-B were also evident. Of the two networks, dATN-B fell more proximal to the early visual networks and dATN-A more distal. dATN-A included distributed regions in frontal cortex at or near FEF, while dATN-B was more locally organized but not exclusively so.

Comparing dATN-A and dATN-B with CG-OP and PM-PPr, as highlighted in panels A and B of Figs. 23 to 25, revealed parallel features. The second-order networks were all spatially anchored near to the early (first-order) sensory and motor networks, appearing as if they grew out or formed from the earlier networks. And, despite being far apart in their major extents, both sets of networks had distributed components with adjacencies in frontal cortex. Thus, from the standpoint of a potential hierarchy among networks, these second-order networks possess a motif that anchors them to the early sensory and motor networks and simultaneously connects them to distributed zones of cortex.

We provisionally label SAL / PMN as a second-order network, but it has juxtapositions that differentiate it from the other second-order networks. Across much of its extent, SAL / PMN paralleled CG-OP with multiple juxtapositions. SAL / PMN differed in that it was not adjacent to early sensory and motor networks. Rather, SAL / PMN contained regions that were near to the network labeled DN-A, especially along the posterior midline, where its regions could easily be confused with the large DN-A and DN-B regions that occupied much of the posterior midline.

The zones farthest away from the sensory and motor regions were populated by five third-order association networks (FPN-A, FPN-B, LANG, DN-B, and DN-A). Each third-order network possessed regions distributed widely throughout association cortex. Moreover, regions of distinct third-order networks displayed side-by-side juxtapositions with a pattern that repeated similarly across multiple zones of cortex. We will focus on these repeating clusters of five networks extensively in later sections.

### Somatomotor and Visual Networks Respond to Body Movements and Visual Stimulation in a Topographic Manner

The spatial extent of task-elicited responses to body movements and visual stimulation were mapped in direct relation to the network boundaries. The goal of these analyses was to explore whether within-individual network estimates predict task responses. In all cases, the network boundaries were established before examination of the task responses. Fig. 26 illustrates the mapping strategy, and Fig. 27 displays the detailed body movement and visual stimulation maps for one representative participant on the inflated and flattened surfaces. Several results are notable.

**Figure 26.**
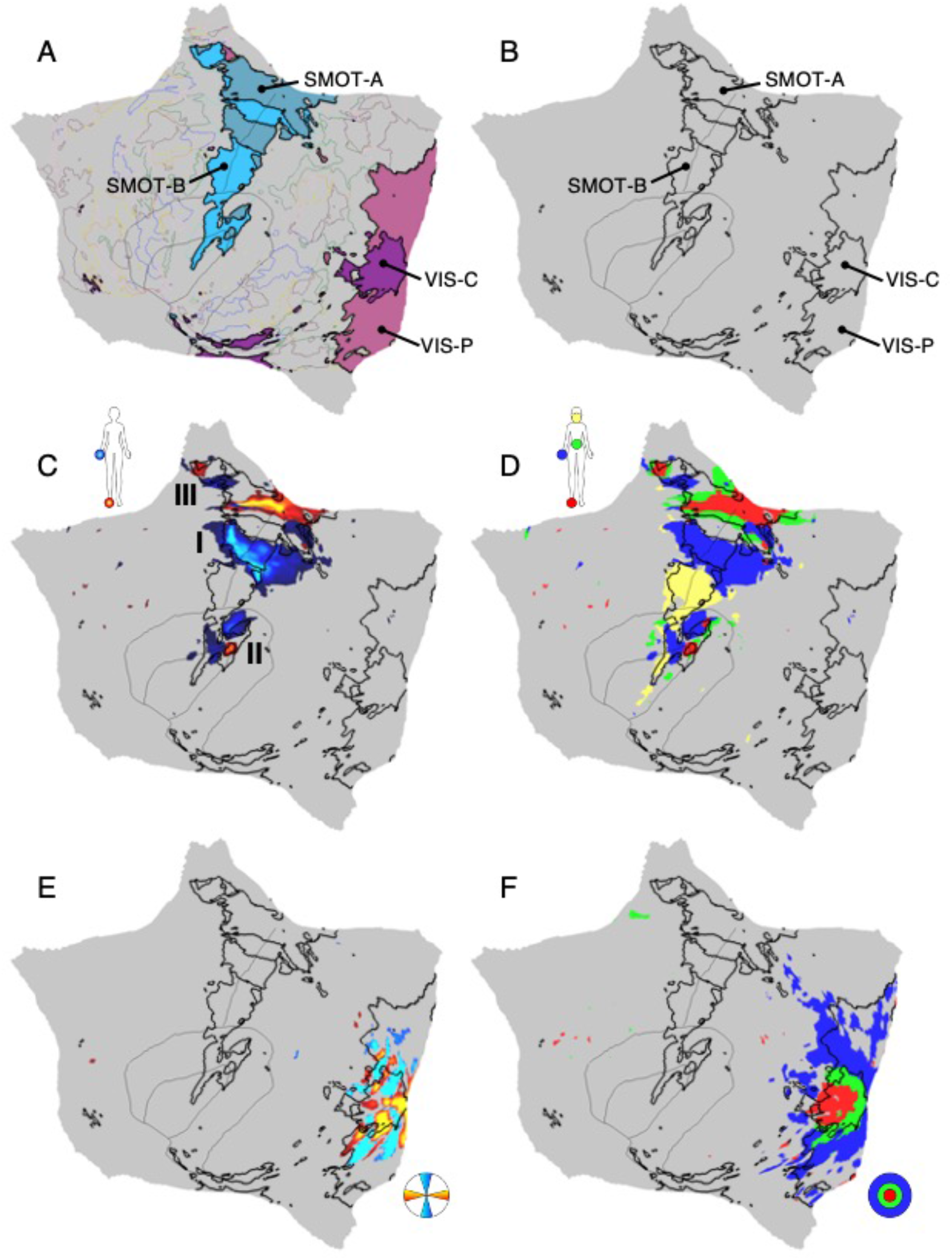
Strategy for exploring somatomotor and visual task responses in relation to networks. Steps employed to generate a combined motor movement and visual stimulation map for a representative participant (P6) are illustrated. (**A**) The within-individual *a priori*-defined somatomotor networks SMOT-A and SMOT-B (blue colors) and visual networks VIS-C and VIS-P (purple colors) are displayed on the flattened cortical surface. Thin colored outlines mark the boundaries of all other networks. (**B**) The borders of SMOT-A, SMOT-B, VIS-C and VIS-P are isolated as black outlines. (**C**) The task contrasts of right versus left foot movements (red) and right versus left hand movements (blue) are mapped in relation to the network boundaries. Presentation of the hand and foot representations in isolation allows visualization of three separate candidate body maps (labelled I, II, and III). The thresholds are z > 2.31 in all cases. (**D**) Binarized motor task contrast maps combine the foot (red), hand (blue), tongue (yellow) and glute (green) movements. Note how adding body parts fills in much of the remaining cortical regions within the somatomotor networks. The thresholds are z > 2.13 in all cases. (**E**) The task contrast of horizontal versus vertical meridian visual stimulation is mapped in relation to the network boundaries to illustrate that multiple areas fall within the VIS-C and VIS-P networks. The thresholds are z < −2.86 and z > 3.16. (**F**) Binarized visual task contrast maps combine the center versus the other apertures (red), middle versus other apertures (green), and peripheral versus other apertures (blue). The threshold is z > 4.15. For display purposes, the binarized maps from **D** and **F** were combined to yield a combined map of somatomotor topography along the body axis and visual topography along the eccentricity gradient.

**Figure 27.**
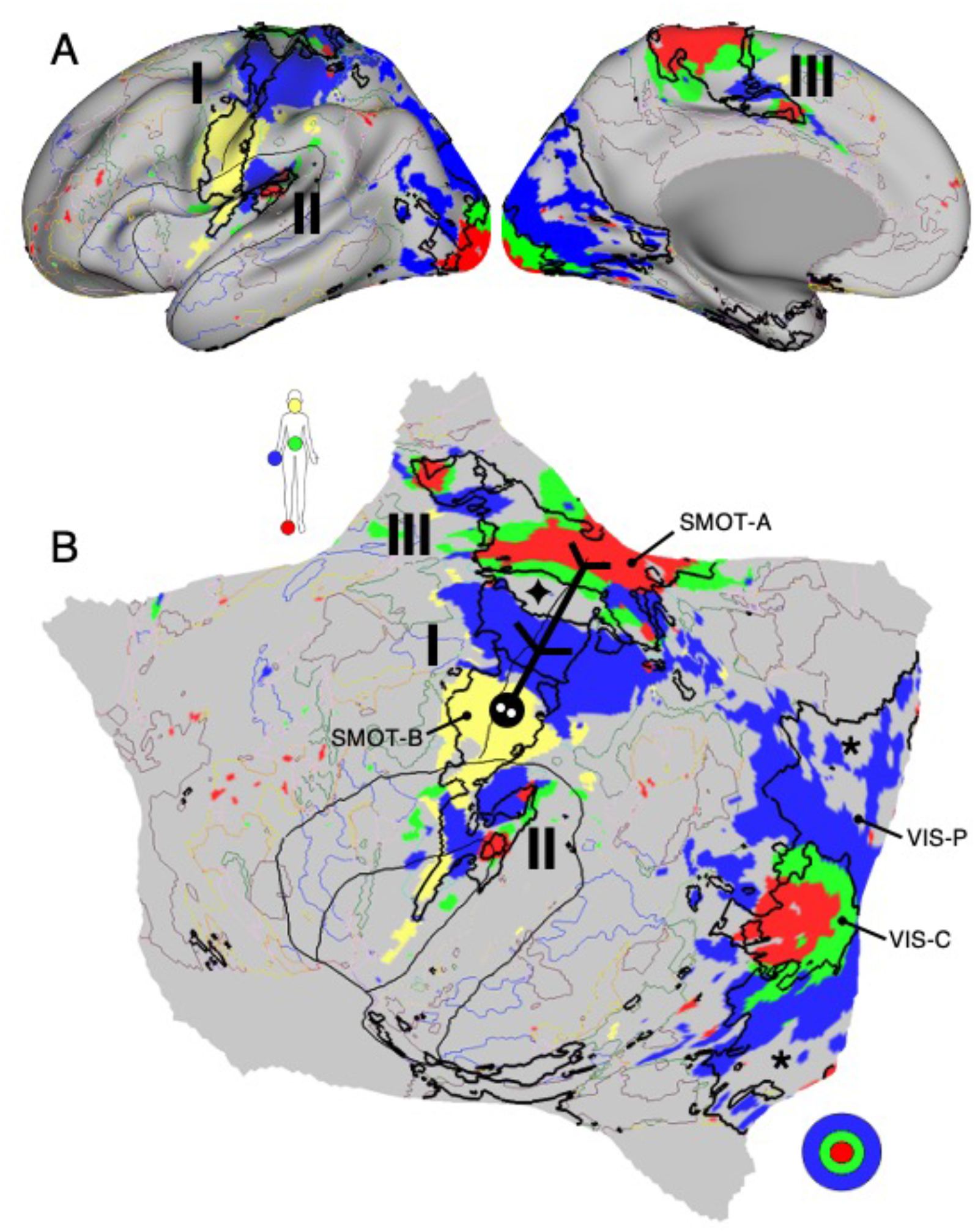
First-order somatomotor and visual networks respond to task stimulation in a topographically specific manner. A detailed view of the inflated (**A**) and flattened (**B**) surfaces display the somatomotor body axis and visual eccentricity maps for P6. The visualization combines panels **D** and **F** of Fig. 26, including binarized contrast maps targeting foot (red), hand (blue), tongue (yellow) and glute (green) movements, as well as central (red), middle (green), and peripheral (blue) visual stimulation. The black labeled outlines highlight networks SMOT-A, SMOT-B, VIS-C, and VIS-P. Thin colored outlines mark the boundaries of all other networks. At least three representations of body topography can be observed within the somatomotor networks SMOT-A and SMOT-B (labeled **I**, **II**, and **III**). The orientation of the main body map (**I**) along the central sulcus is shown by a stick figure. The second body map (**II**) is partially buried in the Sylvian fissure, and the third map (**III**) falls along the frontal midline. The visual gradient from central to peripheral eccentricity is mapped expanding from VIS-C to VIS-P subsuming the V1/V2/V3 cluster (as verified from the task contrast of meridian visual stimulation; see Fig. 26E). One exception is that the eccentricity map spares portions of VIS-P (marked by asterisks) likely due to the limited extent of peripheral stimulation (see methods). A second exception is the gap in the body topography (marked by a diamond) that may be an inter-effector region.

First, while not without exceptions, body movements and visual stimulation elicited responses that were aligned to, and *generally* filled in, the first-order network estimates (SMOT-A, SMOT-B, VIS-C, and VIS-P). That is, the idiosyncratic network estimates in each individual predicted the localization of the movement and visual stimulation responses. The visual responses often extended beyond the anterior boundaries of VIS-C and VIS-P, including visually responsive portions of dATN-B.

Second, the main body map along the central sulcus extended across networks (SMOT-A and SMOT-B) as did the retinotopic eccentricity gradient (VIS-C and VIS-P). Within the visual system, there was a clear correspondence between the two visual networks and eccentricity. VIS-C, as anticipated given its anatomical position, tracked the central representation. VIS-P covered the peripheral representation. A gap emerged for the most peripheral regions of the dorsal and ventral eccentricity portions of VIS-P, likely because the visual stimulation did not extend fully to the periphery (see Park et al. 2023). Within the somatosensory and motor systems, there was a distinct gap between the representations of the hand and glutes which may be an inter-effector region (Gordon et al. 2023).

Third, the response patterns, like the networks, did not align to expected boundaries of individual brain areas (i.e., V1, S1). The body movements activated regions pre- and post-central sulcus, spanning multiple motor and somatosensory areas. Examined in detail, the body movement responses suggest at least three distinct maps of the main body axis, labeled I, II, and III in Fig. 27. The largest distinct body map was found aligned to primary somatomotor cortex, exhibiting a medial-to-lateral progression from foot to hand to tongue (Fig. 27, labeled I). In the posterior insula, the body map was buried with a posterior to anterior orientation (Fig. 27, labeled II). On the medial wall, the body map progressed from anterior to posterior (Fig. 27, labeled III). Similarly, the visual responses spanned the extent of at least the V1/V2/V3 retinotopic cluster, with networks cutting across areas (verified through polar mapping as illustrated in Fig. 26E). Thus, the response patterns confirm that early somatomotor and visual networks group multiple areas together and are separated one from another along topographic gradients (e.g., VIS-C versus VIS-P).

The features described above can be observed in additional participants (Fig. 28), and in all participants with available task data as shown in the Supplemental Materials.

**Figure 28.**
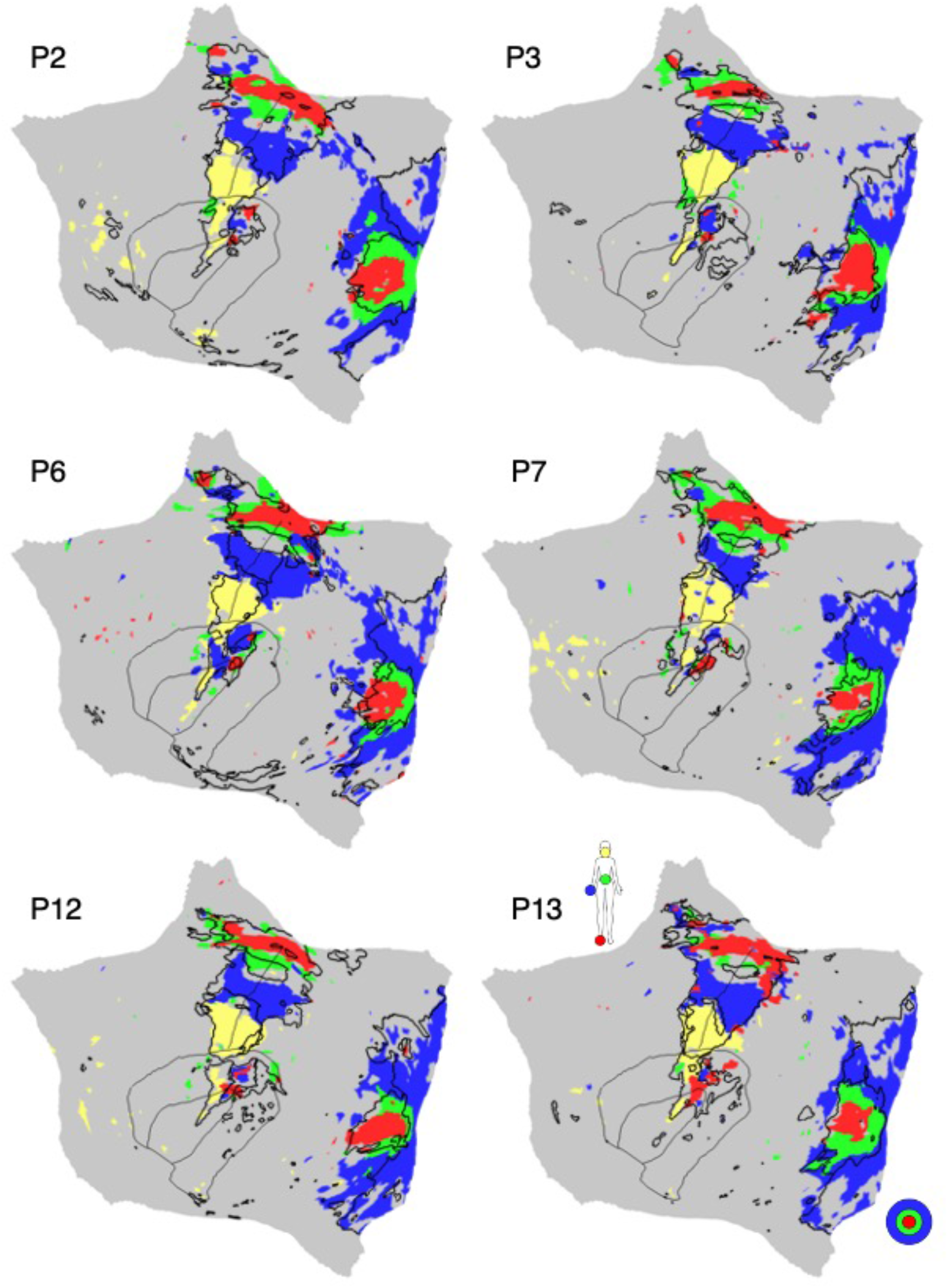
Somatomotor and visual topographic maps are aligned to first-order networks across multiple participants. Flattened surfaces display the somatomotor body axis and visual eccentricity maps in representative participants from the discovery (P2, P3), replication (P6, P7) and triplication (P12, P13) datasets. A body axis topography is evident in each individual by the ordering of tongue-hand-glute-foot along the central sulcus. A visual eccentricity gradient is evident along the calcarine sulcus. While the idiosyncratic spatial details vary between individuals, the somatomotor and visual maps show substantial overlap in each instance with the first-order networks SMOT-A, SMOT-B, VIS-C, and VIS-P. Similar maps from all available participants are included in the Supplementary Materials.

### CG-OP and SAL / PMN Respond to Salient Transients

The oddball task was designed to measure the transient response to uncommon visually salient targets that require participant response. The mapping strategy is illustrated in Fig. 29. On the flattened cortical surface, the within-individual *a priori*-defined networks CG-OP and SAL / PMN are displayed in relation to the Oddball Effect task contrast. The details of the Oddball Effect task contrast are shown for one representative participant in Fig. 30. Fig. 31 illustrates that the features can be observed in additional participants, and in all participants as shown in the Supplemental Materials.

**Figure 29.**
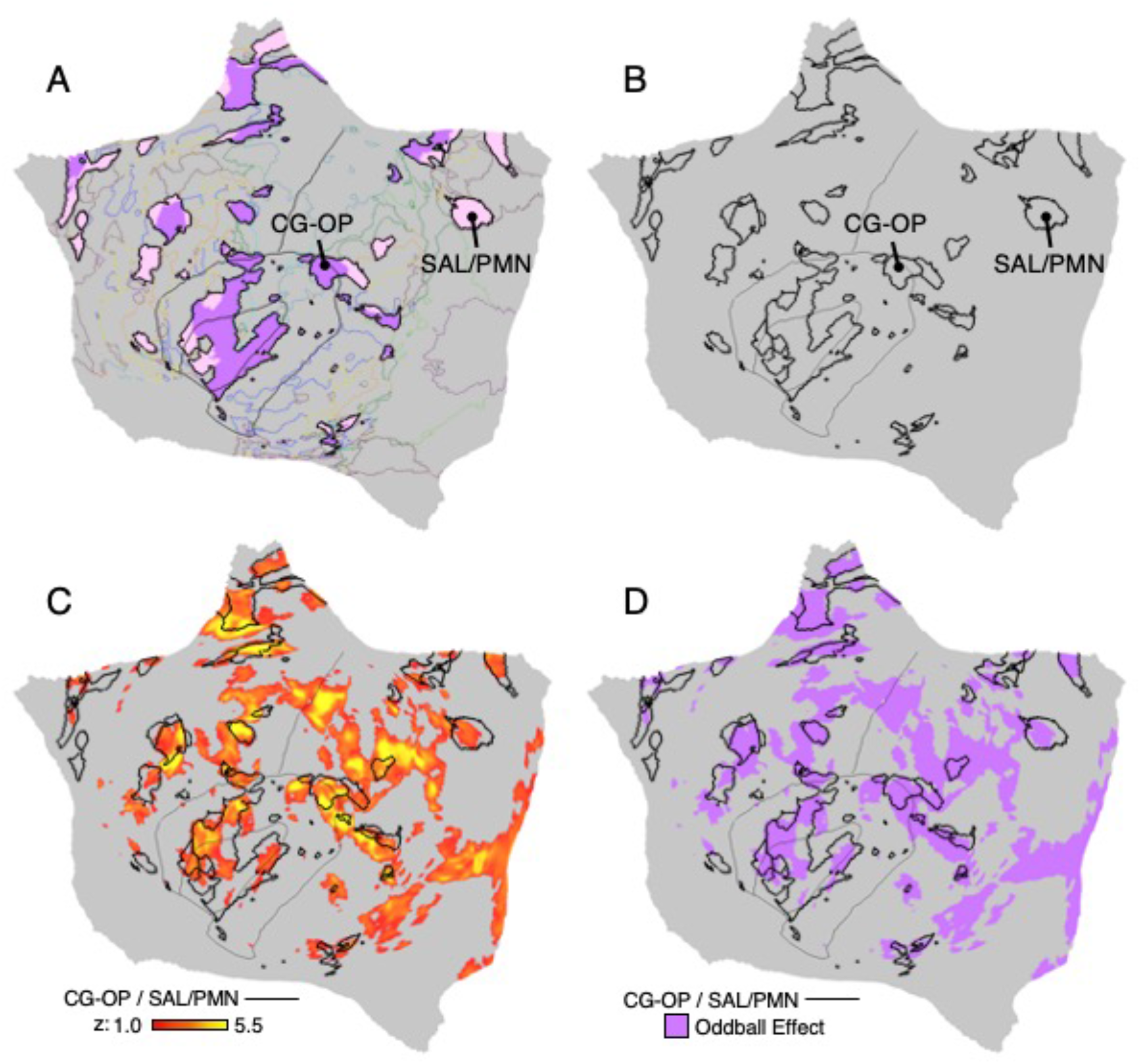
Strategy for exploring responses to oddball detection in relation to networks. Steps employed to generate a map of the Oddball Effect for a representative participant (P6) are illustrated. (**A**) The within-individual *a priori*-defined networks CG-OP and SAL / PMN are displayed on the flattened cortical surface. Thin colored outlines mark the boundaries of all other networks. (**B**) The borders of CG-OP and SAL/PMN are isolated as black outlines. (**C**) The task contrast of oddball event detection versus non-targets, labeled the Oddball Effect, is mapped in relation to the network boundaries. (**D**) The binarized Oddball Effect task contrast map is shown in pink. The threshold is z > 1.00.

**Figure 30.**
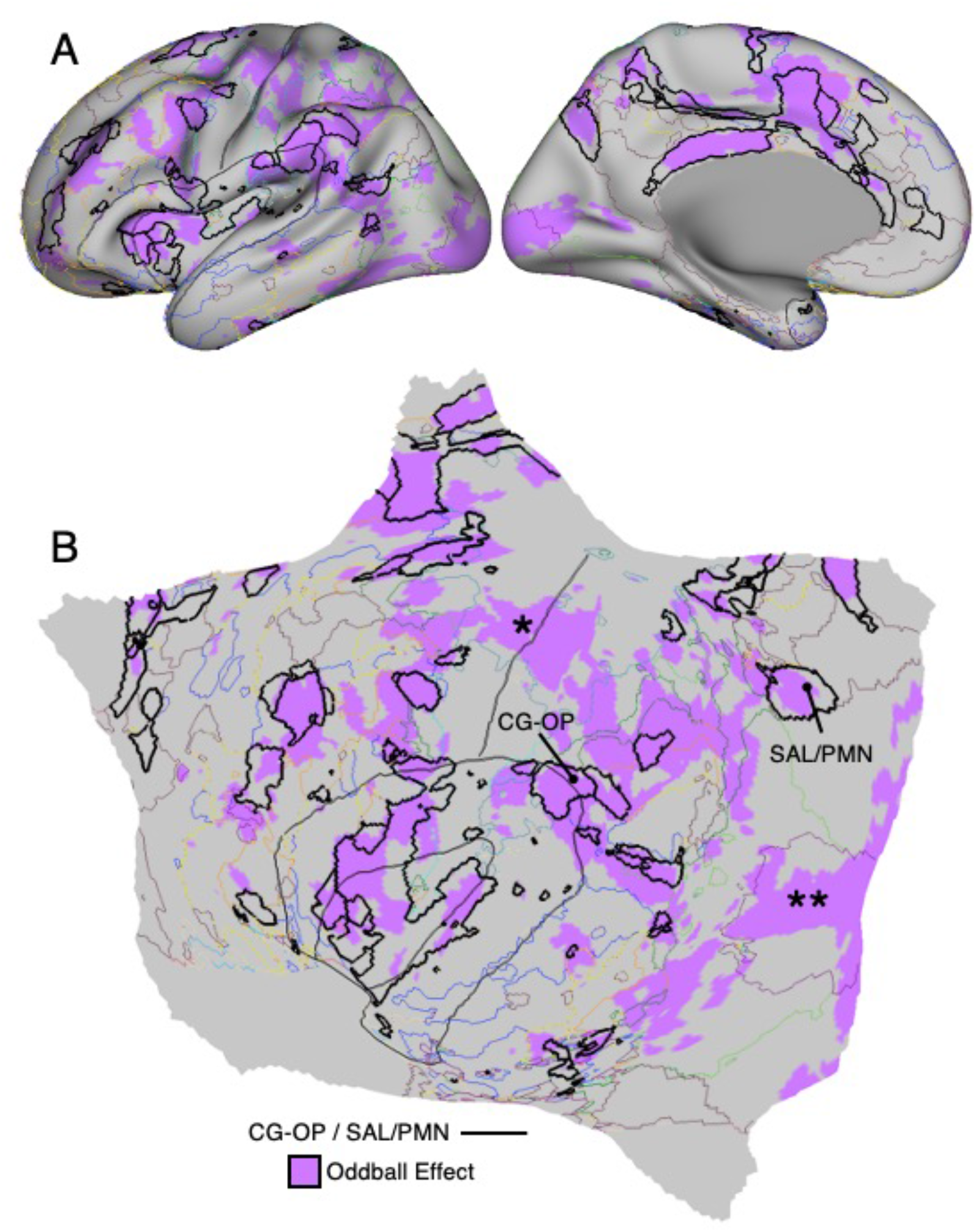
Second-order networks CG-OP and SAL / PMN respond to transients associated with oddball detection. A detailed view of the inflated (**A**) and flattened (**B**) surfaces display the Oddball Effect task contrast map for P6. The black labelled outlines highlight networks CG-OP and SAL / PMN. Thin colored outlines mark the boundaries of all other networks. The Oddball Effect is a distributed with prominent response in the frontal insula, as well as along the posterior and anterior midline. The full response pattern involves many distributed regions of the CG-OP and SAL / PMN networks including posterior midline zones. The effect is not selective to these two networks with a robust response in the hand region of left somatomotor cortex along the central sulcus (marked by asterisk) and the foveal region of visual cortex along the calcarine sulcus (marked by a double asterisk), presumably due to the oddball target response demanding a key press and enhanced attention to the visual cue. The response in the motor region is strongly lateralized (not shown) as expected given the right-handed response.

**Figure 31.**
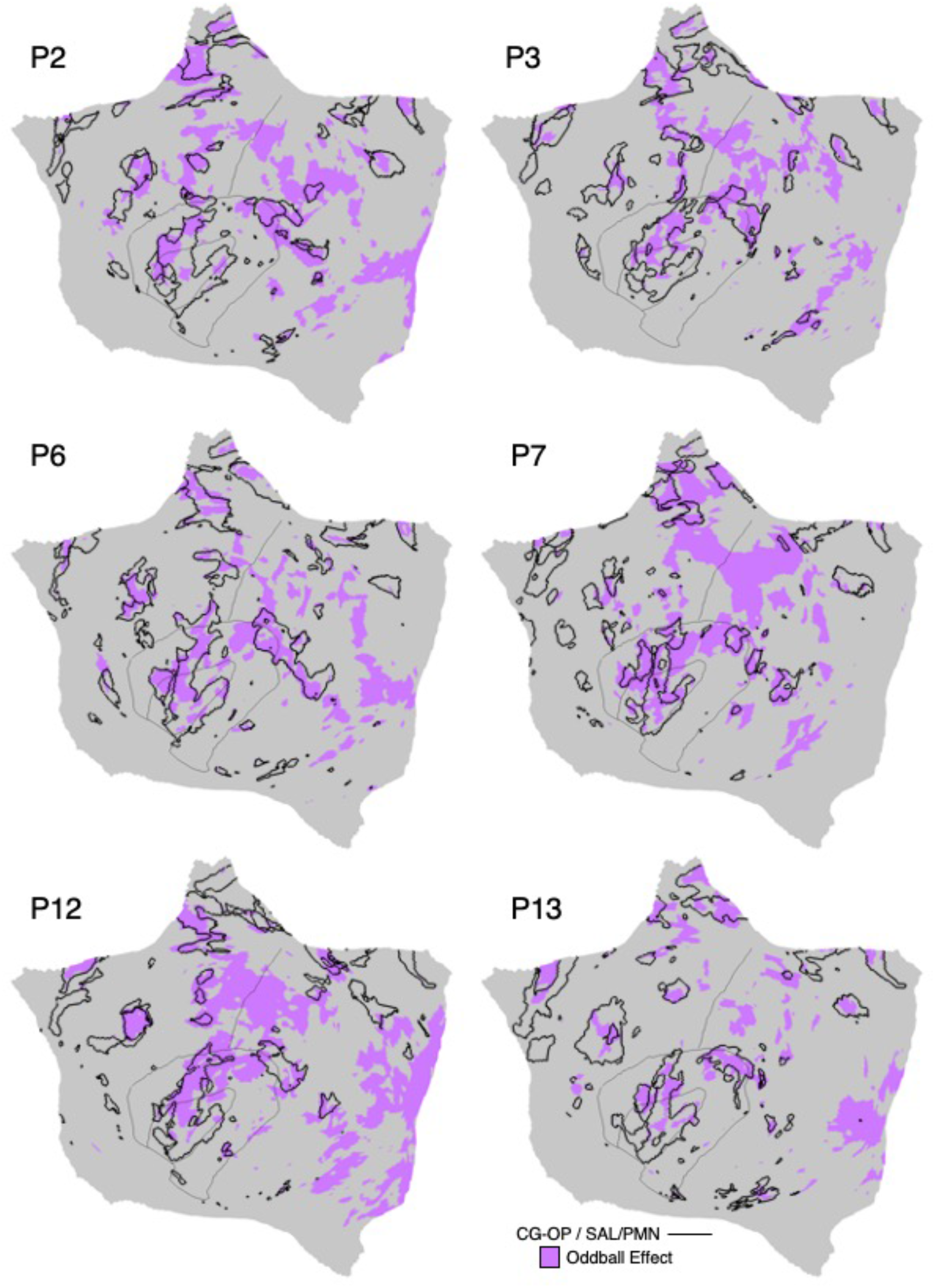
The Oddball Effect is aligned to CG-OP and SAL / PMN across multiple participants. Flattened surfaces display maps of the binarized Oddball Effect in representative participants from the discovery (P2, P3), replication (P6, P7) and triplication (P12, P13) datasets. While the spatial details vary between individuals, the Oddball Effect is broadly localized to the CG-OP and SAL / PMN networks and less so in regions of adjacent association networks, a qualitative impression that is formally quantified in the next figure. Similar maps from all available participants are included in the Supplementary Materials.

The Oddball Effect task contrast response was widely distributed across the cortex. The response prominently involved the distributed regions of the CG-OP and SAL / PMN networks, including regions in the anterior insula as well as along the posterior midline. These collective regions revealing an Oddball Effect task contrast response have been the emphasis of prior studies separately focused on the Salience Network and Parietal Memory Network. Thus, as predicted by the hypothesis that SAL / PMN is a single network, the response pattern observed here extended across the full distributed extent of the network.

In addition to the consistent responses across the distributed regions of CG-OP and SAL / PMN, additional responses were reliably observed – a response along the central sulcus in the left hemisphere near the estimated location of the hand representation and along the calcarine sulcus near the central representation of the visual field (contrast Fig. 30 with Fig. 27). The response in the hand region of the central sulcus was exclusively in the left hemisphere consistent with the right-handed response.

To quantify the selectivity of the task response, the mean *z*-values for the Oddball Effect task contrast were calculated separately for each association network. The estimates were obtained within the bounds of each individual’s *a priori* defined networks and then averaged across participants (N = 14). Quantification at the group level showed a strong, significant positive response to oddball targets in both the CG-OP (*t*(13) = 7.97, *p* < 0.001) and SAL / PMN networks (*t*(13) = 6.21, *p* < 0.001). By contrast, for many of the third-order association networks, the response was significantly negative (DN-A: *t*(13) = −11.76, *p* < 0.001, DN-B: *t*(13) = −8.81, *p* < 0.001, LANG: *t*(13) = −3.82, *p* < 0.01, FPN-B: *t*(13) = −3.02, *p* < 0.01), with FPN-A being the exception. FPN-A showed a weak, non-significant positive response (*t*(13) = 1.82, *p* = 0.09). These observations suggest that the CG-OP and SAL / PMN networks are recruited during the Oddball Effect task contrast.

Given the historical focus on the Salience Network and Parietal Memory Network as separate networks, and their proximity along the posterior midline to the Default Network, we replotted the Oddball Effect task contrast on the inflated surface (Fig. 33). For this visualization, the task map threshold was reduced to zero. Much of the full extent of the CG-OP and SAL / PMN networks was strongly activated. The positive response included the posterior midline regions that have been the focus of the Parietal Memory Network (Gilmore et al. 2015) as well as the anterior insula region that has been a focus of the Salience Network (Seeley et al. 2007; 2019). An interesting feature is that islands of the CG-OP network that fell within the frontal midline showed positive responses in the within-individual maps (Fig. 33). These small responses, which were adjacent to large regions with an opposite response pattern, were absent in the group-averaged response (Fig. 33, bottom). The positive response was not selective to these two specific networks, with motor and visual responses as noted earlier. The positive response also extended into the region of the visual second-order networks (e.g., dATN-B).

**Figure 32.**
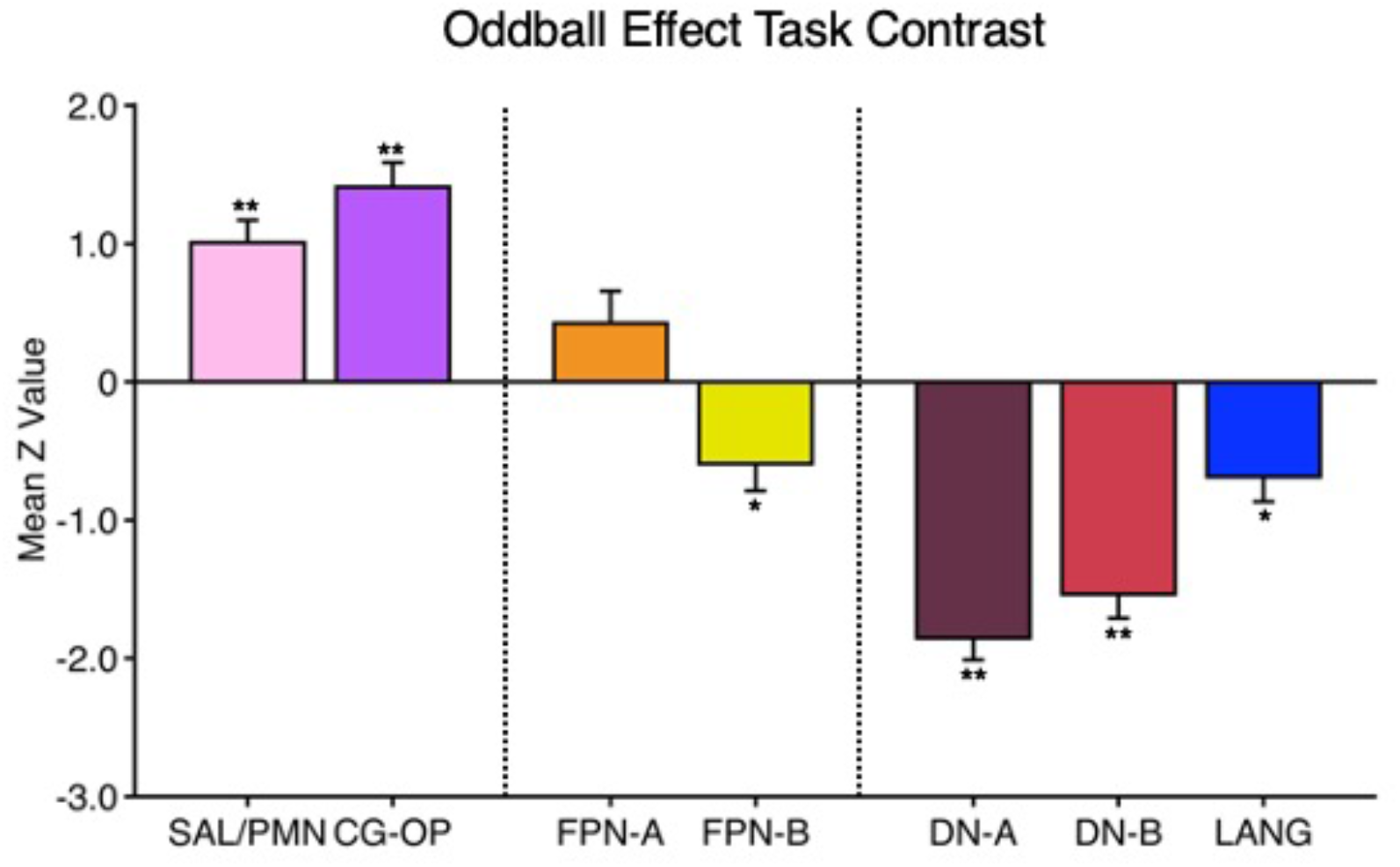
CG-OP and SAL / PMN respond preferentially to transients associated with oddball detection. Bar graphs quantify the Oddball Effect as mean z-values (N = 14) across the multiple *a priori*-defined networks. A strong positive response was observed in the CG-OP and SAL/PMN networks, while adjacent networks displayed lesser (and most often significantly negative) response. Asterisks indicate a value is significantly different from zero (* = *p* < 0.05, ** = *p* < 0.001). Error bars are the standard error of the mean. Note that the CG-OP and SAL / PMN networks are each more active than the other five networks (10 of 10 tests significant *p* < 0.05).

**Figure 33.**
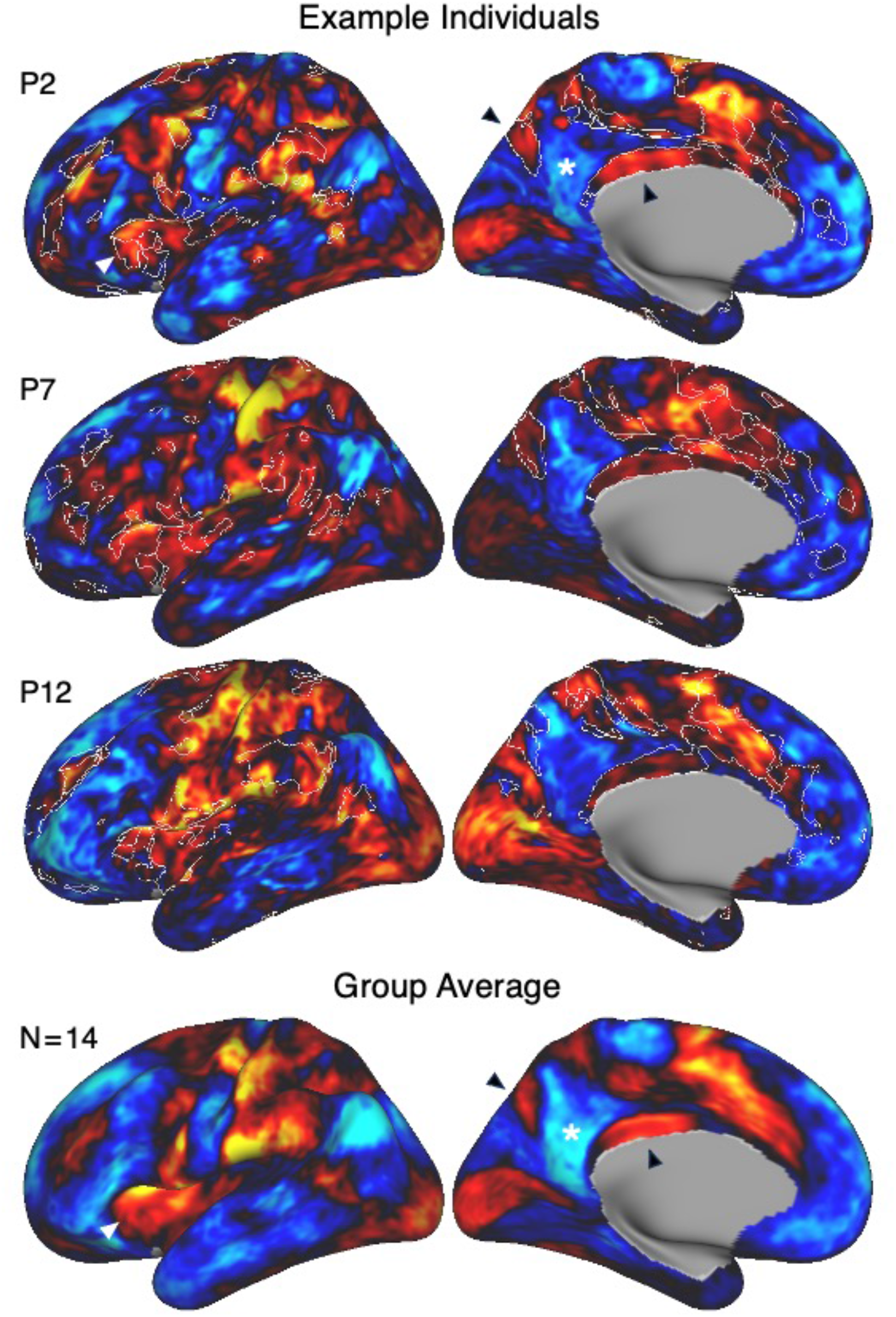
The Oddball Effect robustly dissociates CG-OP and SAL / PMN from regions traditionally associated with the default network. Inflated surfaces display maps of the increases (red/yellow) and decreases (blue) in response for the Oddball Effect task contrast. No threshold is applied to allow full visualization of the effect in both directions. Images in the first three rows are from representative participants from the discovery (P2), replication (P7) and triplication (P12) datasets, and the bottom row displays the group average (N = 14). The white outlines for the individual participants are the outline for the *a priori*-defined CG-OP and SAL / PMN networks. Notice that the Oddball Effect task contrast increases response broadly across the CG-OP and SAL / PMN networks, while there are simultaneously distributed decreases that span multiple networks including DN-A and DN-B. In the top and bottom images, arrowheads highlight the increases in response along the posterior midline (black arrowheads) that surround the canonical Default Network regional decreases (noted by a white asterisk), as well as increases in the anterior insula (white arrowhead). Similar maps from all available participants are included in the Supplementary Materials.

Critically, the networks at or near the historical Default Network, here estimated within-individuals as encompassing at least DN-A and DN-B, were all strongly ‘deactivated’ meaning more active during the implicitly coded baseline reference than during the salient targets. That is, the contrast replicated the task deactivation pattern that originally generated interest in the Default Network (Shulman et al. 1997; Mazoyer et al. 2001; Raichle et al. 2001) in the presence of a robust positive response across the distributed extent of the SAL / PMN network. Thus, the separation of the effects along the posterior midline revealed a spatial dissociation between the second-order network SAL / PMN and the third-order networks DN-A and DN-B.

### Higher-Order Zones of Association Cortex Possess a Repeating Motif

Distributed throughout association cortex, in the zones roughly^7^ between the second-order networks, were the five association networks FPN-A, FPN-B, LANG, DN-B, and DN-A (Fig. 34). Among these networks, side-by-side juxtapositions repeated across multiple cortical zones (refer to I, II, III and IV in Fig. 34). FPN-A and FPN-B were reliably positioned adjacent to one another and, as a pair, were adjacent to a repeating group of the three other networks: LANG, DN-B and DN-A. We call these repeating clusters of five networks Supra-Areal Association Megaclusters or SAAMs. The reproducibility of the SAAMs across participants was striking and is illustrated for the posterior association zones in all 15 participants in Fig. 35. While the idiosyncratic spatial details varied, multiple SAAMs were consistently observed. The remaining task analyses explored functional response properties of the association networks embedded within the SAAMs.

**Figure 34.**
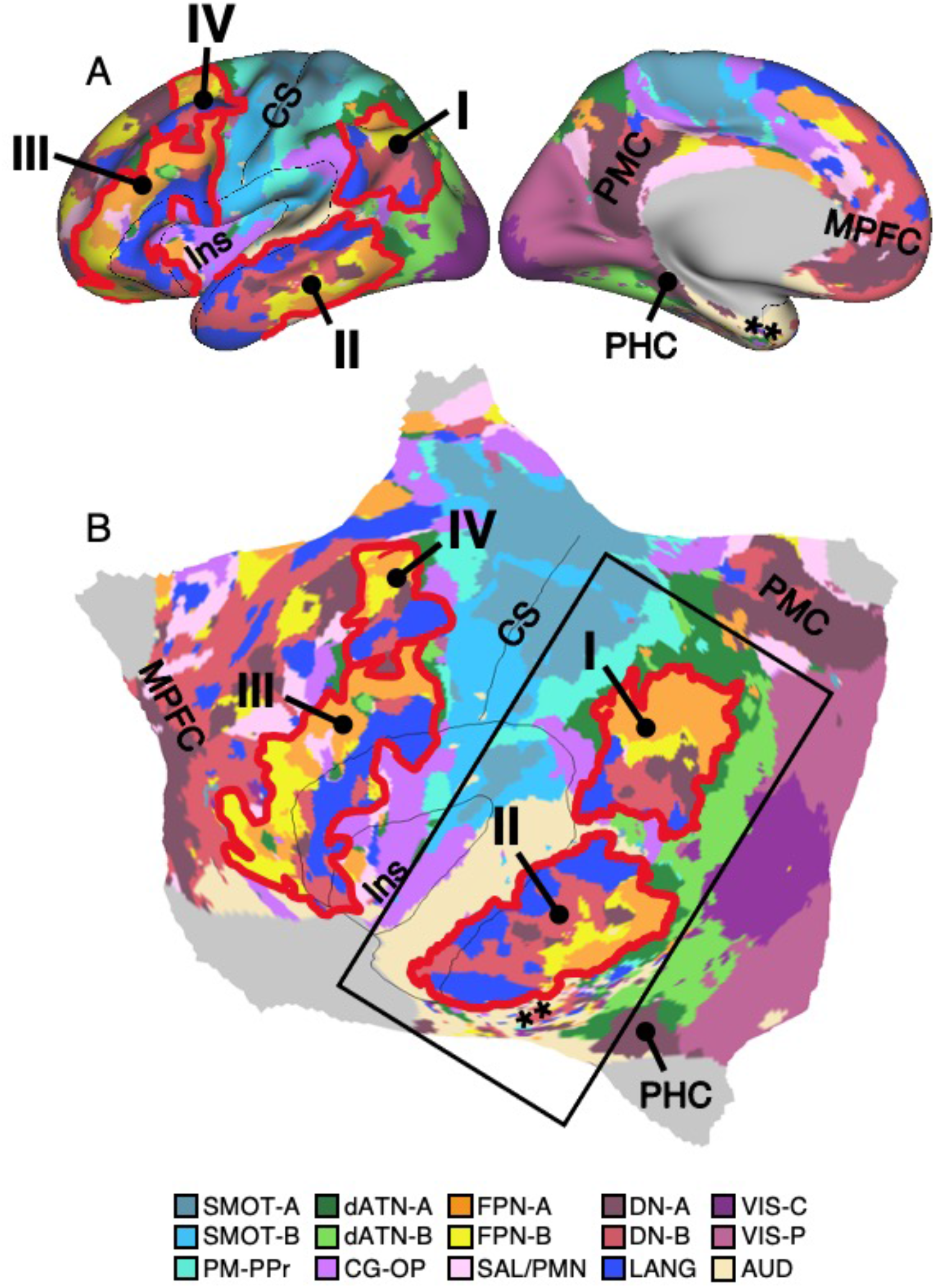
Supra-Areal Association Megaclusters (SAAMs). A detailed view of the inflated (**A**) and flattened (**B**) surfaces display the full set of networks for P4 to visualize an interesting topographic feature of association cortex: a cluster of networks repeats across multiple zones, including within posterior parietal cortex (PPC, **I**), lateral temporal cortex (LTC, **II**), and multiple times throughout PFC (**III**, **IV**). We refer to these repeating clusters as Supra-Areal Association Megaclusters or SAAMs. Within each SAAM, FPN-A and FPN-B are adjacent to one another, and together are adjacent to DN-A, DN-B, and LANG. Thick red outlines mark four SAAMs. The repeating motif is most clear for PPC (**I**) where the cluster has a “north-to-south” orientation and LTC (**II**) where a similar set of juxtapositions display an “east-to-west” orientation. Within PFC, the pattern is present but more ambiguous. Two candidate SAAMs in ventrolateral PFC (VLPFC, **III**) and dorsolateral PFC (DLPFC, **IV**) are highlighted. Reference landmarks include the insula (Ins), central sulcus (CS), posteromedial cortex (PMC), parahippocampal cortex (PHC), and medial PFC (MPFC). Regions of poor SNR that do not allow for confident network assignment are noted by a double asterisk. The rectangle in **B** indicates the portion of the surface that is extracted and displayed for all participants in Fig. 35.

**Figure 35.**
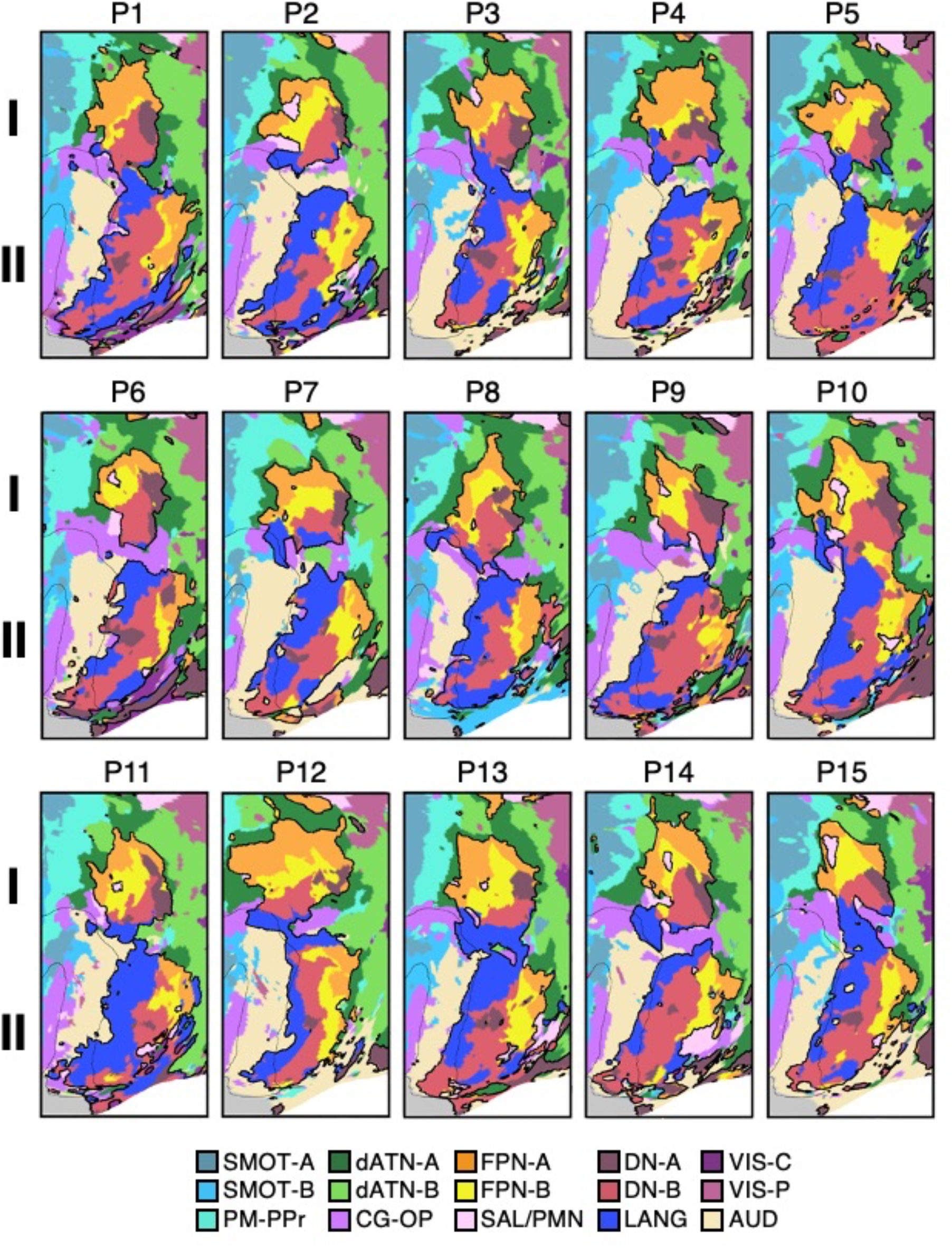
Supra-Areal Association Megaclusters (SAAMs) are reliably observed across multiple participants. Panels display a rotated portion of the flattened surface for 15 individuals (P1 to P15). The displayed portion includes the two SAAMS within PPC (**I**) and LTC (**II**) as illustrated in Fig. 34B. Black outlines illustrate the boundaries of the five networks in each SAAM, including FPN-A, FPN-B, DN-A, DN-B, and LANG. While the idiosyncratic spatial details vary, in most individuals, the separate SAAMs are clear and distinct. Within each SAAM, FPN-A falls at one end juxtaposed with FPN-B. The three side-by-side networks DN-A, DN-B, and LANG fall at the other end of the SAAM with the LANG network most closely juxtaposed to DN-B.

### FPN-A and FPN-B Respond to Domain-Flexible Working Memory Demands

The functional properties of the association networks comprising the SAAMs (FPN-A, FPN-B, LANG, DN-B, DN-A) were explored first in relation to domain-flexible demands on working memory and, in the next section, in relation to domain-specialized processing functions. The hypothesis was that FPN-A and FPN-B would modulate their response in relation to increasing working memory load across multiple verbal and non-verbal stimulus conditions. The mapping strategy is illustrated in Fig. 36. On the flattened cortical surface, the within-individual *a priori*-defined networks FPN-A and FPN-B are displayed in relation to the N-Back Load Effect task contrast (collapsed across stimulus conditions). The N-Back Load Effect task contrast is shown in detail in Fig. 37 for one representative participant. Fig. 38 illustrates that the features can be observed in additional participants, and in all participants with available task data in the Supplemental Materials.

**Figure 36.**
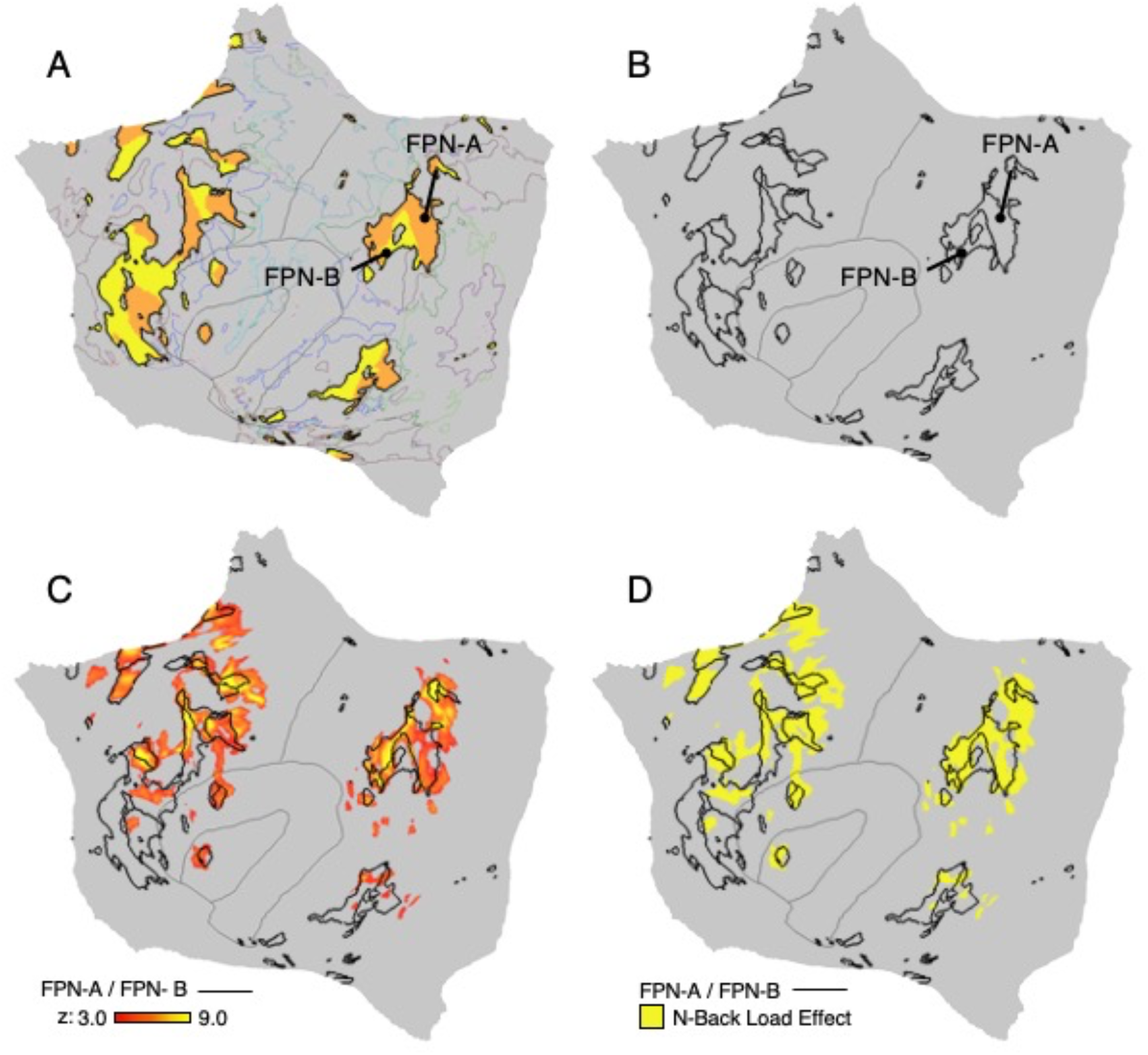
Strategy for exploring responses to high working memory load in relation to networks. Steps employed to generate a map of the N-Back Load Effect for a representative participant (P6) are illustrated. (**A**) The within-individual *a priori*-defined networks FPN-A and FPN-B (orange and yellow colors) are displayed on the flattened cortical surface. Thin colored outlines mark the boundaries of all other networks. (**B**) The borders of FPN-A and FPN-B are isolated as black outlines. (**C**) The task contrast of 2-Back (High Load) versus 0-Back (0-Back), labeled the N-Back Load Effect (red/yellow), is mapped in relation to the network boundaries. (**D**) The binarized N-Back Load Effect task contrast map is shown in yellow. The threshold is z > 3.00.

**Figure 37.**
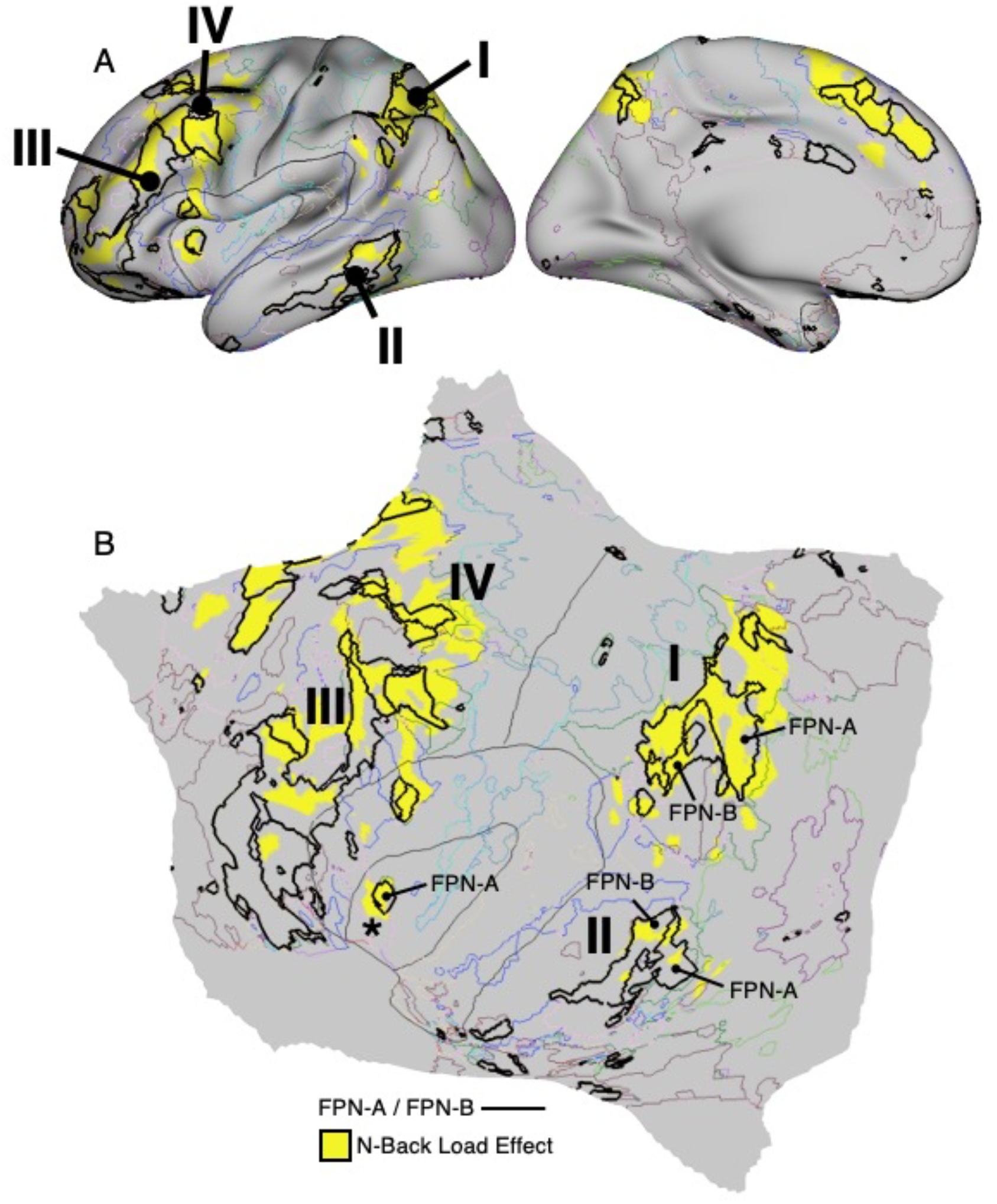
Networks FPN-A and FPN-B respond to high working memory load. A detailed view of the inflated (**A**) and flattened (**B**) surfaces display the N-Back Load Effect task contrast map for P6. The black labeled outlines highlight the FPN-A and FPN-B networks. Thin colored outlines mark the boundaries of all other networks. The N-Back Load Effect shows prominent response across the multiple, distributed association zones preferentially within the FPN-A / FPN-B networks, including the relevant portions of the SAAMs. The zones are labeled **I** to **IV** to orient to the corresponding labels of the SAAMs as displayed in Fig. 34. The response also consistently includes a small subregion of the anterior insula that is associated with FPN-A (labeled with an asterisk).

As hypothesized, the N-Back Load Effect task contrast increased activation within and near the boundaries of the FPN-A and FPN-B networks (Figs. 36 and 37). The widely distributed response included extensive regions of PFC, as well as regions of PPC and the dorsal ACC – all canonical regions associated with domain-flexible cognitive control (e.g., Duncan 2001; Cromer et al. 2010; Duncan 2013; Fedorenko et al. 2012). As predicted by the network estimates, there was also a response in LTC and a small subregion of the anterior insula that is spatially distinct from that of other networks.

Of equal importance was the consistent absence of response in the distributed association regions linked to the LANG, DN-B, and DN-A networks, including within the PPC and LTC. In essence, the N-Back Load Effect task contrast split the SAAMs and activated the portions linked to the FPN-A and FPN-B networks selectively with minimal or no response in the juxtaposed portions associated with the LANG, DN-B, and DN-A networks.

To quantify the selectivity of the task response, the mean *z*-values for the N-Back Load Effect task contrast were calculated separately for each association network. The estimates were obtained within the bounds of each individual’s independent *a priori* defined networks and then averaged (N = 15). Results plotted in Fig. 39 reveal a positive N-Back Load Effect response that was strongest in FPN-A (*t*(14) = 21.67, *p* < 0.001) and also quite strong in FPN-B (*t*(14) = 6.45, *p* < 0.001). SAL / PMN unexpectedly showed a significant positive response (*t*(14) = 7.91, *p* < 0.001) that was significantly weaker than either FPN-A (*t*(14) = −15.09, *p* < 0.001) or FPN-B (*t*(14) = −2.91, *p* < 0.01). Thus, while SAL / PMN showed a response, the functional response was less relative to FPN-A and FPN-B, opposite to the pattern found earlier (contrast Fig. 39 with Fig. 32). The remaining networks, including the three additional networks that were adjacent within the SAAMs, showed a negative N-Back Load Effect. The effect was significantly negative for DN-A (*t*(14) = −4.85, *p* < 0.001) and DN-B (*t*(14) = −7.14, *p* < 0.001) but not LANG (*t*(14) = −0.81, *p* = 0.43). These results provide evidence that two parallel networks – FPN-A, FPN-B – are involved in processes enhanced by increasing working memory demands, while other juxtaposed networks – LANG, DN-B and DN-A – are functionally dissociated, consistent with the qualitative patterns visualized in the activation maps.

**Figure 38.**
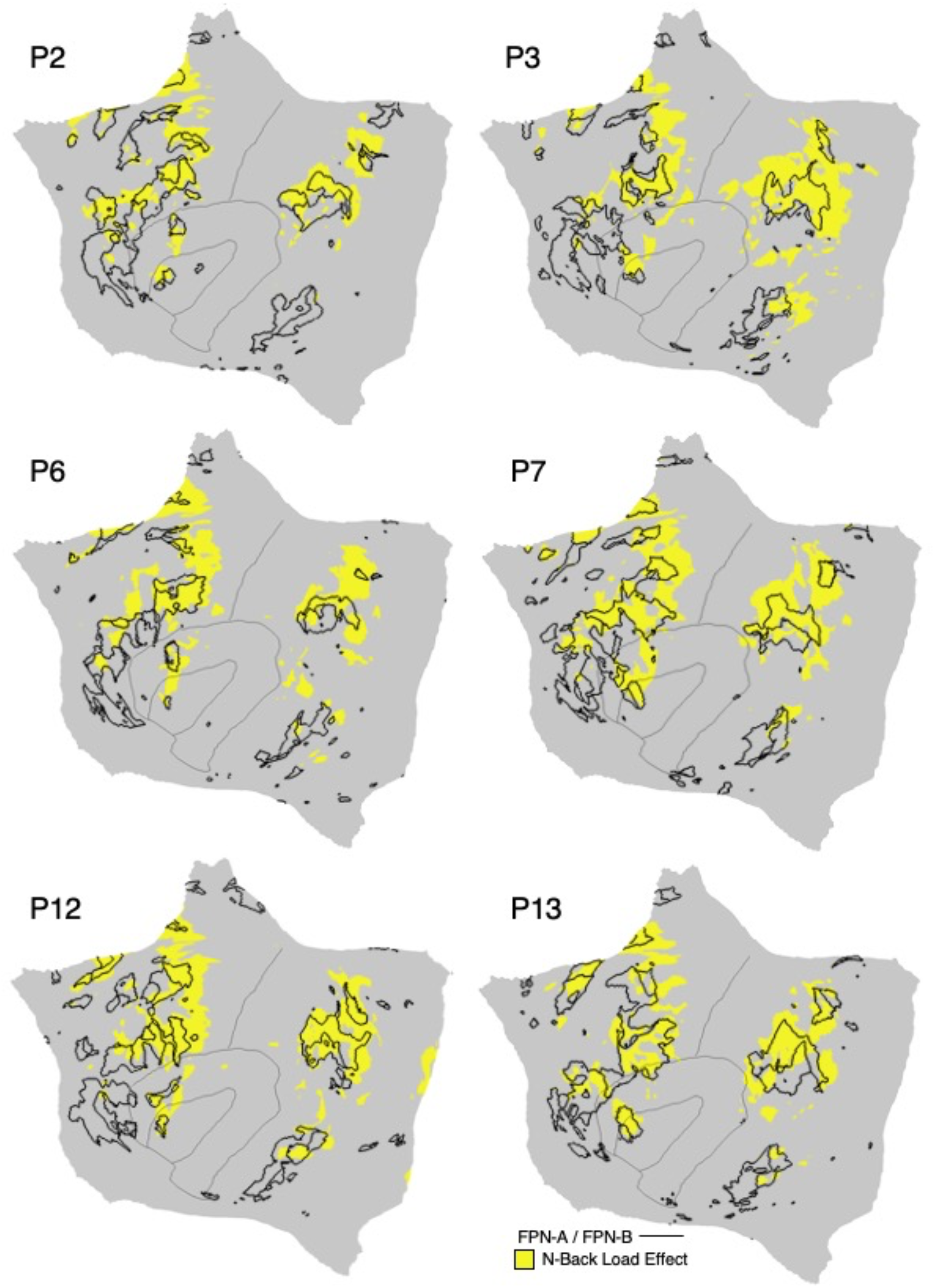
The N-Back Load Effect is aligned to FPN-A and FPN-B across multiple participants. Flattened surfaces display the binarized N-Back Load Effect maps for multiple participants from the discovery (P2, P3), replication (P6, P7) and triplication (P12, P13) datasets. While individuals vary in anatomical details, the N-Back Load Effect is generally localized to the FPN-A and FPN-B networks. Similar maps from all available participants are included in the Supplementary Materials.

**Figure 39.**
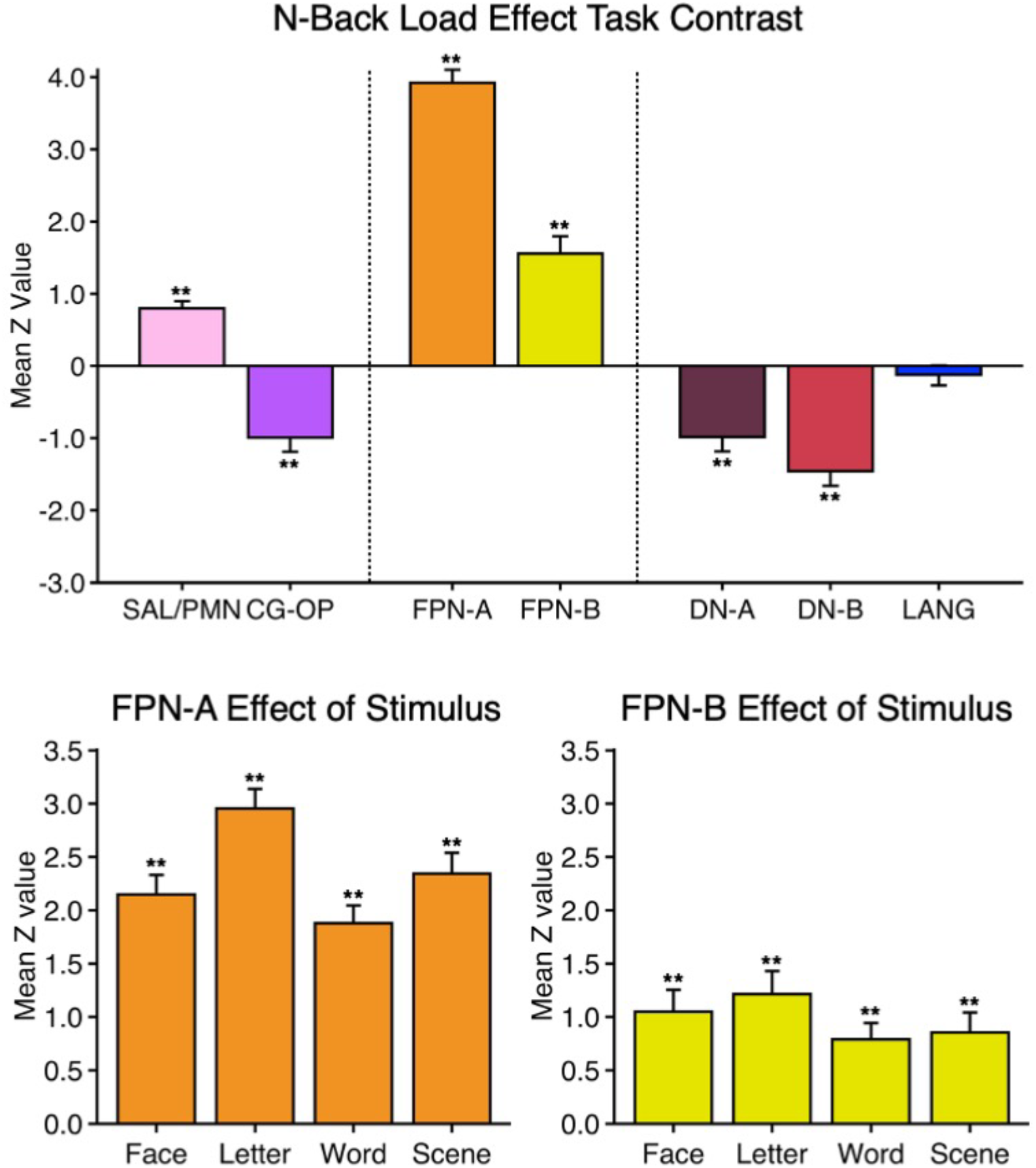
FPN-A and FPN-B respond preferentially to high working memory load in a domain-flexible manner. Bar graphs quantify the N-Back Load Effect as mean z-values (N = 15) across the multiple *a priori*-defined networks. (**Top**) A strong positive response was observed in the FPN-A and FPN-B networks, while other association networks displayed minimal or no response, with the exception of the SAL / PMN network which also displayed a significant, positive response. Error bars are the standard error of the mean. Note that FPN-A and FPN-B are each more active than all five of the other networks (10 of 10 tests were significant *p* < 0.05). (**Bottom Left**) The N-Back Load Effect is quantified separately for each stimulus domain (Face, Letter, Word, and Scene) within FPN-A. Note that the effect is robust and significant across domains. (**Bottom Right**) The N-Back Load Effect is quantified separately for each stimulus domain within FPN-B. Note again that the effect is positive and significant across domains. Asterisks indicate a value is significantly different from zero (* = *p* < 0.05, ** = *p* < 0.001).

To further investigate the domain flexibility of FPN-A and FPN-B, the mean *z*-values for each of the four stimulus conditions of the N-Back Load Effect (Face, Letter, Word, and Scene) were separately plotted (Fig. 39). Both FPN-A (Face: *t*(14) = 11.74; Letter: *t*(14) = 16.03; Word: *t*(14) = 11.30; Scene: *t*(14) = 12.05, all *p* < 0.001) and FPN-B (Face: *t*(14) = 5.13; Letter: *t*(14) = 5.60; Word: *t*(14) = 5.15; Scene: *t*(14) = 5.54, all *p* < 0.001) exhibited a significant response across all conditions of the N-Back Load Effect task contrast, supporting that their processing role generalizes across both verbal and nonverbal domains. That is, FPN-A and FPN-B responded robustly to working memory demands, more so than adjacent networks and did so in a do’ain-flexible manner.

### LANG, DN-B, and DN-A Respond Differentially to Distinct Cognitive Domains

Among the networks that populate the distributed zones of higher-order association cortex, FPN-A and FPN-B responded in a domain-flexible manner to increasing working memory load. The adjacent trio of networks – LANG, DN-B, and DN-A – did not. In our final analyses, we explored the functional specialization of these additional three networks by examining Episodic Projection, Theory-of-Mind and Sentence Processing task contrasts designed to emphasize distinct specialized domains of higher-order cognitive processing.

The mapping strategy is illustrated in Fig. 40. On the flattened cortical surface, the within-individual *a priori*-defined networks LANG, DN-B, and DN-A are displayed in relation to the three separate task contrasts simultaneously, to illustrate the adjacency of the responses in relation to each other and to the network boundaries. The details of one composite task contrast map are displayed for a representative participant in Fig. 41. Fig. 42 illustrates additional participants, and all participants with available data are shown in the Supplemental Materials. Several results are notable.

**Figure 40.**
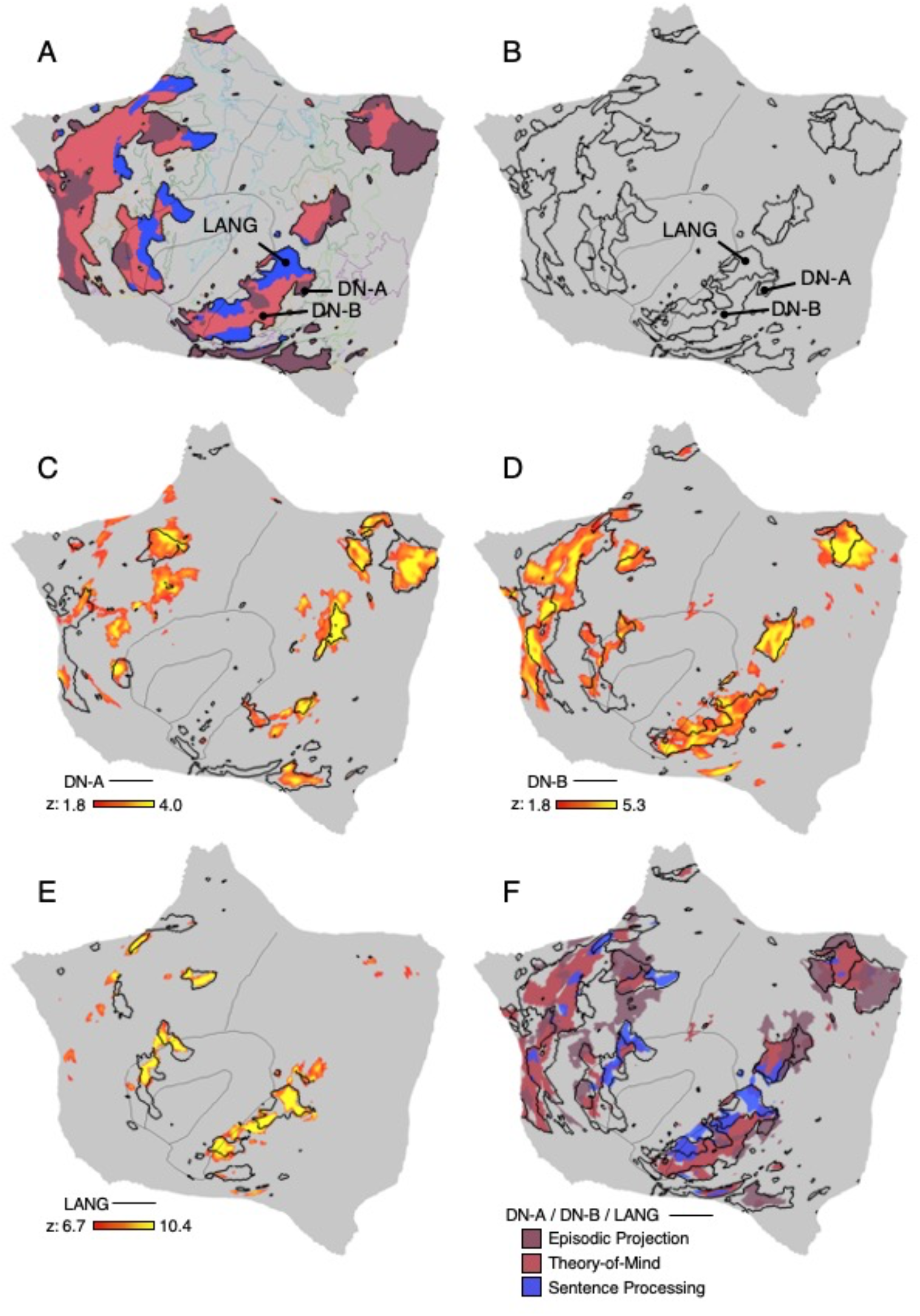
Strategy for exploring domain-preferential higher-order responses in relation to networks. Steps employed to generate a combined map revealing domain-selective responses for a representative participant (P6) are illustrated. (**A**) The within-individual *a priori*-defined networks DN-A (dark red), DN-B (light red) and LANG (blue) are displayed on the flattened cortical surface. Thin colored outlines mark the boundaries of all other networks. (**B**) The borders of DN-A, BN-B and LANG are isolated as black outlines. (**C**) The Episodic Projection task contrast (red/yellow) is mapped on its own in relation to the DN-A network boundary. (**D**) The Theory-of-Mind task contrast (red/yellow) is mapped on its own in relation to the DN-B network boundary. (**E**) The Sentence Processing task contrast (red/yellow) is mapped on its own in relation to the LANG network boundary. (**F**) Binarized task contrast maps are shown together (dark red, Episodic Projection; light red, Theory-of-Mind; blue, Sentence Processing). The threshold is z >1.80. The combined, binarized map allows visualization of the multiple functional domains in the same view.

**Figure 41.**
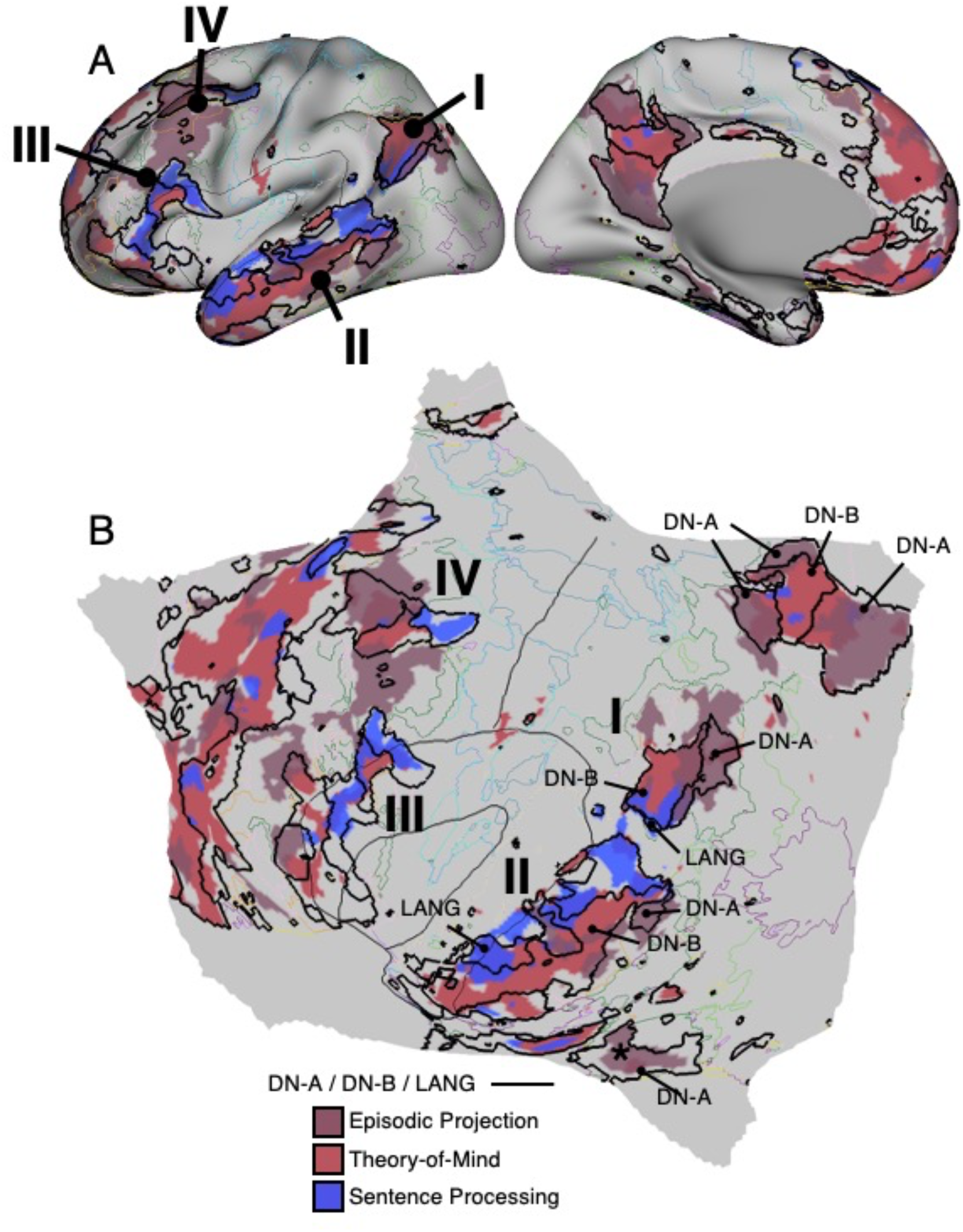
DN-A, DN-B, and LANG respond in a domain-selective manner. A detailed view of the inflated (**A**) and flattened (**B**) surfaces display the Episodic Projection (dark red), Theory-of-Mind (light red), and Sentence Processing (blue) task contrast maps for P6. The black labeled outlines highlight the DN-A, DN-B, and LANG networks. Thin colored outlines mark the boundaries of all other networks. The task contrasts reveal clear spatial separation across the multiple, distributed association zones preferentially within the DN-A, DN-B, and LANG networks, including the relevant portions of the SAAMs. The zones are labeled **I** to **IV** to orient to the corresponding labels of the SAAMs as displayed in Figs. 34 and 37. The parahippocampal cortex (labeled with an asterisk) responds preferentially to the Episodic Projection task contrast without juxtaposed response from other domains, unlike the SAAMs which each have representation of all three domains, separate from (but adjacent to) zones responding in a domain-flexible manner to working memory load (see Fig. 38).

**Figure 42.**
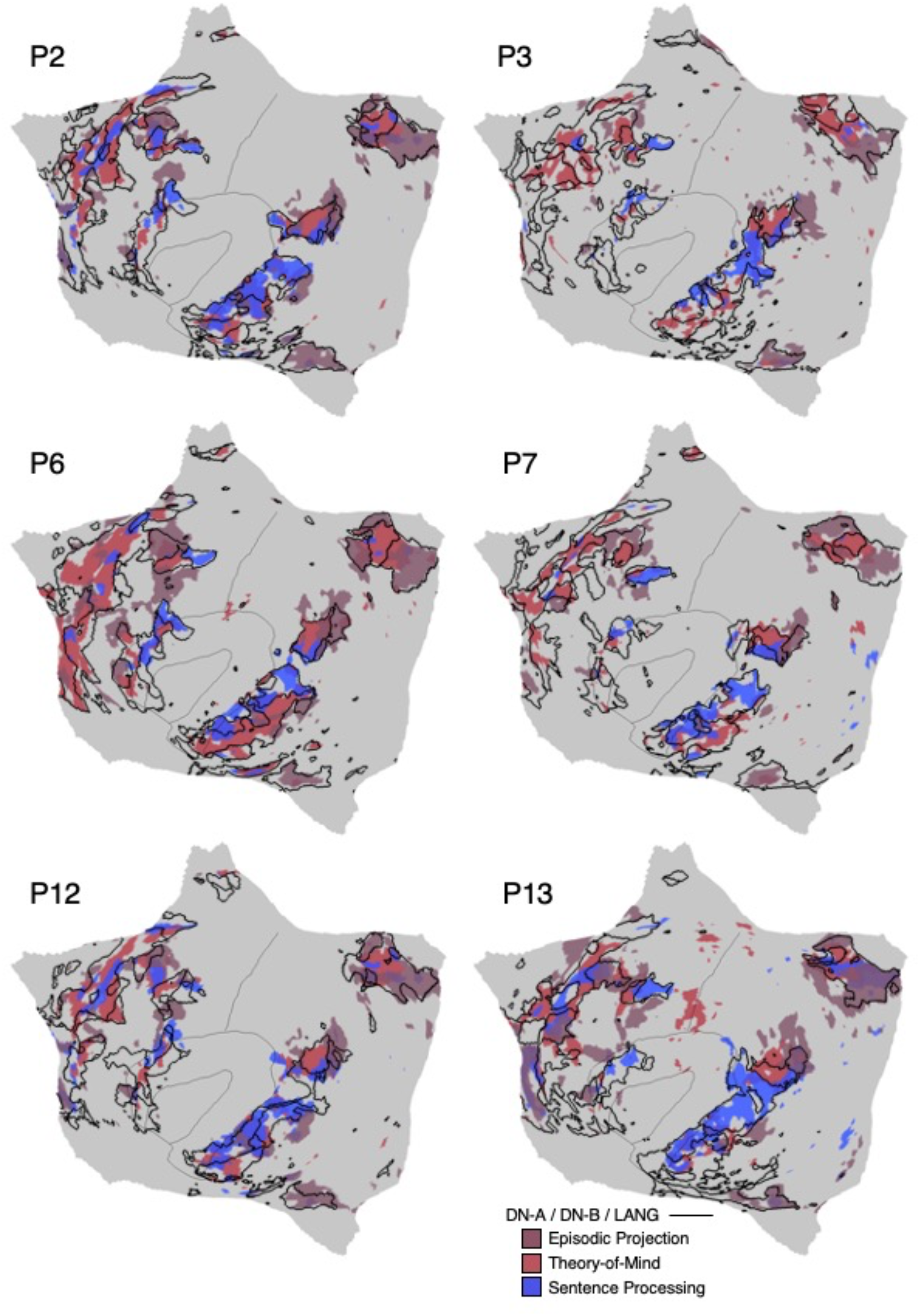
Domain-selective responses are aligned to DN-A, DN-B, and LANG across multiple participants. Flattened surfaces display maps of the binarized Episodic Projection, Theory-of-Mind, and Sentence Processing task contrast maps for multiple participants from the discovery (P2, P3), replication (P6, P7) and triplication (P12, P13) datasets. The domain-preferential effects are generally localized to corresponding DN-A, DN-B, and LANG networks and separate from the adjacent zones that respond to working memory load (contrast the present maps with those of Fig. 38). Similar maps from all available participants are included in the Supplementary Materials.

First, the composite activation patterns across the three task contrasts filled in the remaining zones of association cortex. Strikingly, the domain-specialized task responses are situated adjacent to, but separate from, the regions activated by domain-flexible working memory demands (contrast Fig. 41 with Fig. 37). This separation can be seen in many locations, with a particularly clear example visualized within the PPC where the N-Back Load Effect task contrast showed a posterodorsal response relative to the three current task contrasts. The side-by-side juxtaposition of domain-specialized and domain-flexible regions was also observed within LTC and multiple locations throughout PFC.

Second, within each juxtaposed cluster of domain-specialized regions, the region preferentially responding to the Sentence Processing task contrast abutted the region preferentially responding to the Theory-of-Mind task contrast, and these abutted the region preferentially responding to the Episodic Project task contrast. While overlap and exceptions were found, the differential response patterns generally tracked the network separations between LANG, DN-B, and DN-A. The idiosyncratic positions and boundaries of the three networks in any given individual – LANG, DN-B, and DN-A – predicted the positions of the domain-specialized activation responses (Fig. 42).

Thus, within each local zone the regions associated with the separate networks responded to their distinct specialized cognitive domains. Moreover, the spatially differentiated response patterns repeated across the multiple SAAMs (refer to I, II, III and IV in Fig. 41). There were exceptions. For example, regions of task activation in VLPFC did not overlap well with the estimated networks in P12. The discrepancies tended to fall within anterior temporal regions and PFC regions where SNR is low, raising the possibility that technical variance played a role. To reveal the details of the task maps more fully, the Supplementary Materials include task maps for each task contrast separately in addition to the composite maps for all available participants.

The response was quantified for each of the three task contrasts for each network to formally test for the hypothesized interaction. For each domain-specialized task contrast, the *z*-values within the bounds of each individual’s three independent *a priori* defined networks (LANG, DN-B and DN-A) were obtained and then averaged (N = 13). The resulting mean *z*-values are plotted in Fig. 43. A repeated measures ANOVA on network-level task response revealed a significant 3 x 3 interaction between the effect of task contrast and network (*F*(4, 48) = 77.82, *p* < 0.001). Paired *t*-tests then tested the individual contrasts, with the hypothesis that each network’s within-domain response would be significantly greater than either of the other two networks. All six of these planned comparisons were significant. The Episodic Projection task contrast recruited DN-A regions over those of DN-B (*t*(12) = 16.38, *p* < 0.001) and LANG (*t*(12) = 14.49, *p* < 0.001). The Theory-of-Mind task recruited DN-B regions over those of DN-A (*t*(12) = 5.27, *p* < 0.001) and LANG (*t*(12) = 10.09, *p* < 0.001), and the Sentence Processing task contrast recruited the LANG regions over those of DN-A (*t*(12) = 6.55, *p* < 0.001) and DN-B (*t*(12) = 5.42, *p* < 0.001).

**Figure 43.**
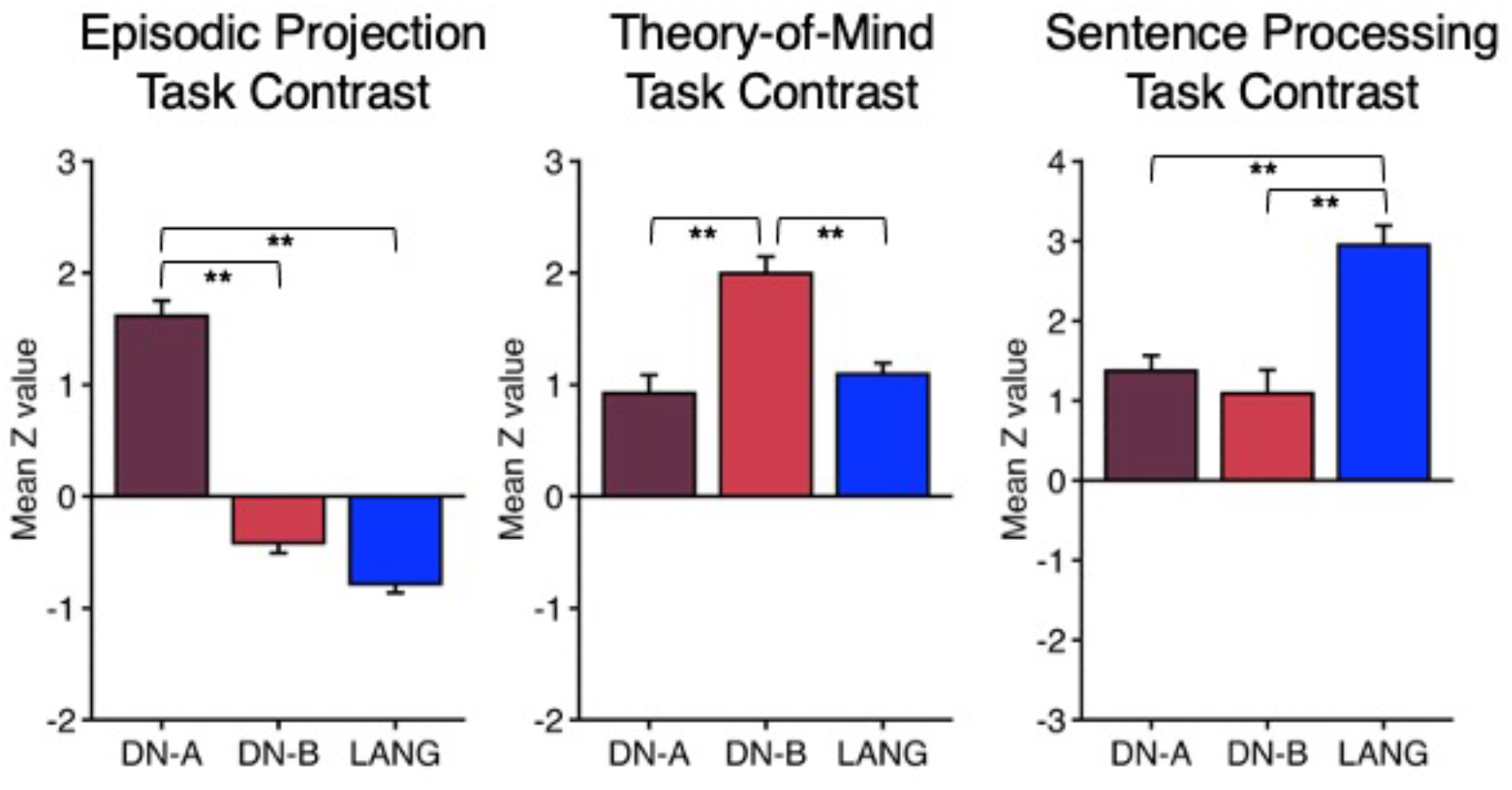
DN-A, DN-B, and LANG respond in a domain-selective manner. Bar graphs quantify the Episodic Projection, Theory-of-Mind, and Sentence Processing task contrasts as mean z-values (N = 13) across the multiple *a priori*-defined networks. Each plot displays data from a distinct task contrast; each bar represents a distinct network. The full 3×3 interaction (network by task contrast) is significant (*p* < 0.001). DN-A is robustly and preferentially activated for the Episodic Projection task contrast; DN-B is robustly and preferentially activated for the Theory-of-Mind task contrast; and LANG is robustly and preferentially activated for the Sentence Processing task contrast. All planned pairwise comparisons are significant confirming the full triple dissociation. Asterisks indicate a value is significantly different from zero (** = *p* < 0.001).

Thus, in addition to the qualitative impressions (Figs. 40-42), statistical tests revealed the full interaction was significant with all pairwise tests also significant in support of a triple functional dissociation across the three networks. These observations suggest that the parallel networks LANG, DN-B, and DN-A, with adjacent regions across multiple cortical zones, are specialized to support distinct higher-order cognitive domains.

## Discussion

Detailed network estimates reveal a global organization that can be conceptualized as three levels of cortical hierarchy: locally-organized first-order sensory and motor networks, spatially adjacent second-order networks that link to distant regions, and third-order networks that populate and connect widely distributed zones of higher-order association cortex. Repeating side-by-side spatial juxtapositions among the third-order association networks form organized motifs that we call Supra-Areal Association Megaclusters or SAAMs. Within each SAAM, the regions linked to distinct association networks demonstrate differential task response properties. Certain networks contribute to domain-flexible cognitive control and others to domain-specialized processes involved in language, social, and spatial / episodic functions. We discuss the practical and conceptual implications of these findings including how repeating organizational motifs might arise during development.

### Within-Individual Network Estimates

In the present work, we explored the utility of a 15-network MS-HBM estimate of cerebral cortical organization that allowed the idiosyncratic details of each individual’s own anatomy to guide the solutions (e.g., Figs. 2 and 4). The method yielded robust, stable network estimates that were confirmed using analyses of seed-region based correlation (e.g., Figs. 3 and 5). All quantitative analyses and visual inspections of the data reinforced that the present 15-network estimate captured a great deal of the structured correlations present in the underlying data. From a methodological standpoint, the present results indicate that a MS-HBM can be used to estimate networks automatically and robustly within individuals (Kong et al. 2019; Xue et al. 2021). Several features of our network estimates revise or expand earlier ideas.

First, the current network parcellation falls into a class of within-individual network estimates that refine group-based estimates. In group-based estimates, including multiple estimates from our laboratory, large monolithic networks have been identified that encompass extensive regions of association cortex (e.g., Damoiseaux et al. 2006; Yeo et al. 2011; Power et al. 2011; Doucet et al. 2011; Smith et al. 2013). Our network estimates are broadly similar but separate the large group-based networks into multiple distinct parallel networks. For example, the network historically known as the Default Network overlaps four separate networks in the present parcellation including networks LANG, DN-B, DN-A, and SAL / PMN. Each of these four distinct networks can be identified in every individual in the present study. The multiple networks are not estimated to be “sub-networks” with shared regions or anatomical convergence, but rather are distinct networks that are near to one another and often blurred in group-averaged data (see also Fedorenko et al. 2010; 2012; Laumann et al. 2015; Michalka et al. 2015; Braga et al. 2017; 2019; Gordon et al. 2017; Buckner and DiNicola 2019; Smith et al. 2021). Thus, an advance of within-individual network estimates, including the present contribution, is to fully resolve adjacent networks that are difficult to separate through approaches that average over people.

Second, among within-individual parcellation estimates, we settled on a 15-network solution because of our goal to separate nearby networks within the anterior insula (Seeley 2019), as well as to better separate early sensory and adjacent networks. Our analyses confirmed that the newly proposed 15-network parcellation could capture correlational features absent in simpler network solutions, including our own 10-network solution previously estimated in Xue et al. (2021; see current Figs. 8, 9, 11, and 12). In addition to detecting distinctions among networks that have close juxtapositions in the anterior insula, the present 15-network parcellation also revealed clear separation of the estimated AUD network from the nearby LANG network. Multiple networks identified in the simpler network solutions remained in the 15-network estimates, indicating the refinements did not come at the expense of the established networks.

Third, the present parcellation identifies a single distributed network, labeled SAL / PMN, that includes regions that have historically been studied separately as components of the Salience Network (Seeley et al. 2007) and the Parietal Memory Network (Gilmore et al. 2015). Note that we do not say “joins” two previously described networks, as we suspect there have never been two separate networks. Rather, different research lineages may have focused on distinct regional components of what is ultimately the same network. This hypothesis will require further testing, but several lines of evidence lead to the present proposal that SAL / PMN is a single, coherent network. In every individual, the estimated SAL / PMN network included regions along the posterior midline and within the anterior insula (Fig. 20). Seed-region based correlation patterns recapitulated the automated network estimates: seed regions placed in PFC and posterior cortex revealed clear regional correlation in the anterior insula as well as multiple distinct posterior midline regions (Fig. 21). Furthermore, independent task data focused on salience processing, via an oddball detection task, elicited robust responses in the distributed regions of the SAL / PMN network including the posterior midline and anterior insula (Figs. 30, 31, and 33).

Finally, it is important to note that the present estimates assume (and are optimized to detect) large-scale distributed networks. For this reason, our resultant parcellation is different from parcellations that are optimized to detect local gradients of change and / or directly estimate “area” boundaries (e.g., Cohen et al. 2008; Gordon et al. 2016; Glasser et al. 2016; for discussion see Buckner and Yeo 2014; Eickhoff et al. 2018). While there is some convergence between approaches, and it is possible to apply mutual constraints (Schaefer et al. 2018), our present parcellation is weighted to estimate networks based on long-range correlational properties, without weighting local gradients.

### Supra-Areal Association Megaclusters (SAAMs)

A striking observation that is apparent in the flat map visualizations is the recurrent spatial grouping of the same five higher-order networks throughout association cortex (FPN-A, FPN-B, LANG, DN-B, DN-A). The clearest examples are found in PPC and LTC (Fig. 35), but the adjacencies are also present in multiple PFC zones (Fig. 34), as if a shared organizing force plays out repeatedly across different cortical territories. Each grouping of regions possesses similar spatial relations among the five networks: networks FPN-A and FPN-B are next to one another, and that pair of networks is adjacent to the trio of networks LANG, DN-B, and DN-A.

SAAMs possess several additional features. While their global patterning – meaning spatial adjacencies between networks – is identifiable for multiple SAAMs within and across individuals, the orientations shift, and the exact spatial positions vary. For example, within the PPC the axis that begins with the FPN-A / FPN-B pairing and ends with the LANG / DN-B / DN-A triad is oriented dorsal-to-ventral. Within the LTC, the axis is oriented ventral-to-dorsal (Fig. 34). Moreover, while the SAAMs are readily identified in every person in the PPC and LTC, usually with a discontinuity between the two SAAMs, the idiosyncratic spatial details vary from one person to the next. In some individuals the two zones appear fused (Fig. 35). It is thus unsurprising that group-averaged data, while revealing certain spatial features apparent in the SAAMs, blur over the fine spatial details that are apparent in the within-individual maps.

The spatial juxtapositions that define the SAAMs in PPC and LTC are also present in multiple zones of the PFC. However, there is not always spatial separation. The boundaries of individual SAAMs in PFC are thus ambiguous. In Fig. 34 we note candidate SAAMs in VLPFC (labeled III) and DLPFC (labeled IV), recognizing these are hypotheses. A future endeavor might explore how a repeating pattern could parsimoniously explain the juxtapositions in PFC with the assumption that multiple SAAMs are present like those observed in PPC and LTC, but with the additional complication that there are multiple adjacent SAAMs that collide into one another, perhaps as a consequence of their formation during development.

A final detail regarding the SAAMs is subtle but potentially informative. While the presence of five regions linked to the distinct networks is a consistent feature of PPC, LTC, VLPFC, and DLPFC, there are also partial sets of the network juxtapositions in other cortical zones. For example, along the midline there is clear representation of networks DN-A and DN-B in PMC and MPFC, but not consistently the other networks (Fig. 34). The partial SAAMs may provide an insight into the origins of the patterning. DN-A is a putative hippocampal-cortical network that has been extensively studied in humans (e.g., Grecius et al. 2004; Vincent et al. 2007; Braga and Buckner 2017; Braga et al. 2019; Zheng et al. 2021; Reznik et al. 2023) and monkeys (e.g., Buckner et al. 2008; Binder et al. 2009; Margulies et al. 2009; Buckner and Margulies 2019; Buckner and DiNicola 2019). The hippocampal formation, via polysynaptic projections through entorhinal cortex and PHC, projects heavily to RSC and ventral PCC along the posterior midline, and also to MPFC (Suzuki and Amaral 1994; Lavenex et al. 2002; Blatt et al. 2003). The exclusive assignment of PHC to DN-A and the predominance of DN-A along the midline may thus reflect connectivity to the hippocampal formation. The interdigitation of DN-A with other higher-order networks might emerge as the hippocampal-predominant projections intermix with other anatomical projection gradients in the apex association zones where the fully formed SAAMs are present.

### The Relation of the Present Network Estimates with the Historical Default Network

The Default Network, or Default Mode Network, has received considerable attention among investigations of cerebral networks (Gusnard and Raichle 2001; Buckner et al. 2008; Smallwood et al. 2021). In relation to estimating networks using resting-state functional connectivity, after the seminal description of the method (Biswal et al. 1995), the Default Network was the first distributed association network to be characterized in humans (Greicius et al. 2003; 2004) and in monkeys (Vincent et al. 2007). All group-based network estimates, even low-dimensional solutions that identify as few as seven networks, find a large, distributed network that has the spatial pattern of the Default Network (e.g., Beckmann et al. 2005; Damoiseaux et al. 2006; Yeo et al. 2011; Power et al. 2011; Doucet et al. 2011). Thus, a critical issue to address, given the historical emphasis on the Default Network, is how the present network estimates relate to these earlier descriptions.

Our current hypothesis is that the large monolithic (or core-subnetwork) descriptions of the Default Network based on group-averaged data, including our own contributions (e.g., Buckner et al. 2008; Andrews-Hanna et al. 2010), are inaccurate and reflect an artifact of spatial blurring. As noted above, the canonical group-averaged Default Network overlaps fully or partially four distinct networks: LANG, DN-B, DN-A, and SAL / PMN. The separation of these networks is anticipated in some prior group-based analyses. For example, Andrews-Hanna and colleagues (2014) noted that regions within PPC responding to social inference (theory-of-mind) tasks tended to activate an anterior region relative to tasks targeting remembering. This distinction likely captures the separation of DN-B and DN-A in PPC. Similarly, in a thorough analysis of functional connectivity in group data, the network identified here as LANG was separated from the canonical Default Network (Lee et al. 2012; for discussion see Braga et al. 2020). However, the blurring induced by between-subject averaging, to date, has negated the ability to resolve the spatial details that fully distinguish the four nearby networks that are described here.

A further observation emerges from our task-based results. In addition to the challenge of identifying the multiple, juxtaposed networks due to spatial blurring, there is a separate functional property that has anchored study of the Default Network. The Default Network was originally described based on task-induced deactivations, referring to the observation that the distributed association regions that comprise the Default Network are more active in passive tasks and fixation than active, externally-orientated tasks (Shulman et al. 1997; Gusnard and Raichle 2001; Mazoyer et al. 2001; see Buckner and DiNicola 2019 for review). When a contrast is made between active and passive tasks, a distributed pattern of “deactivations” emerges that is robust and overlaps with group-based estimates of the Default Network (Buckner et al. 2008; Buckner 2014; Smallwood et al. 2021). What is surprising and interesting is that, even with the present high-resolution within-individual estimates, the task-induced pattern of deactivation remains broad and spans multiple networks.

Specifically, the Oddball Effect task contrast reveals a broad task-induced deactivation pattern within individual participants (Fig. 33). That is, the regions deactivated by attending and responding to external stimuli span multiple association networks even when group averaging is not a factor. Fig. 32 quantifies this effect: DN-A, DN-B, and LANG all show significant “deactivation,” with DN-A and DN-B being almost indistinguishable from one another, despite clear functional double dissociation during domain-relevant active tasks (e.g., Fig. 43; see also DiNicola et al. 2020). One possibility is that, while DN-A and DN-B are anatomically and functionally distinct networks, they may collectively be suppressed during certain externally oriented task events, perhaps as a result of a broad antagonistic process between externally versus internally oriented processing modes (Buckner and DiNicola 2019; see also Nyberg et al. 1996; Fransson 2005; Fox et al. 2005; Miller, Weaver, and Ojemann 2009; Anticevic et al. 2012). Thus, the phenomenon of task-induced deactivation, which is not selective to specific networks, may have reinforced that there is a coherent monolithic function across large swaths of association cortex, a possibility refuted by a growing number of robust functional dissociations (e.g., Peer et al. 2015; Silson et al. 2019; DiNicola et al. 2020; Deen and Friewald 2022; DiNicola et al. 2023).

Another relevant observation surrounds the relation between the Default Network and the present estimate of network SAL / PMN. The SAL / PMN network possesses regions distributed across the cortex, including multiple distinct regions along the posterior midline side-by-side with DN-A and DN-B network regions. The adjacencies make the regions easy to confuse. Despite their spatial proximity, Zheng et al. (2021; see their Fig. 6) noted that “deactivations” are restricted to the Default Network and separate from their estimate of SAL / PMN (labeled as the Parietal Memory Network in their paper). The transient positive response in SAL / PMN observed here to salient oddballs is robust including the regions along the posterior midline, separate from the juxtaposed DN-A and DN-B regions showing deactivation (Fig. 33). Moreover, SAL / PMN has small, focal regions of response in MPFC, which are also surrounded by DN-A and DN-B network regions. In the group-averaged map displayed in the bottom of Fig. 33, there is no detectable positive response in MPFC. Each individual shows a response but in slightly different spatial positions from one person to the next. The positive task response in MPFC is likely lost in the process of spatial averaging.

Our results thus converge with Zheng et al. (2021) to suggest that SAL / PMN is spatially and functionally distinct from the network historically described as the Default Network. The SAL / PMN network does not exhibit task-induced deactivation; rather, it displays an opposite functional response pattern – transiently activating to salient external task events, including in both posterior and anterior regions along the midline.

### Hierarchical Organization of the Cerebral Cortex

Paul Flechsig (1901; 1904; 1920) contributed the powerful but simple idea that the cerebral cortex develops sequentially radiating outwards from motor and sensory cortex (see Bailey and von Bonin 1951; Meyer 1981; Clarke and O’Malley 1996; Mesulam 2015 for translations and discussion; see Zilles 2018 for further context). The basis of Flechsig’s hierarchy was the developmental timing of myelination of the fibers reaching the cortex. By his account “in the cerebral convolutions, as in all other parts of the central nervous system, the nerve-fibers do not develop everywhere simultaneously, but step by step in a definitive succession” (translated in Clarke and O’Malley 1996, p. 548). The cortical motor and sensory (and certain limbic) zones myelinate first. Next are the intermediate zones that surround the motor and sensory zones. The terminal zones myelinate in the final stage, beginning approximately four months after birth, and encompass prefrontal, temporal, and parietal regions thought of today as higher-order association cortex. The prescient lens of hierarchical cortical organization provides a framework to understand our findings.

Specifically, the candidate assignments of first-, second-, and third-order networks are motivated by (and agree well with) Flechsig’s reference maps of sequential myelination (Fig. 44). In particular, the distributed regions late to myelinate (the terminal zones) are positionally similar to our estimated association zones containing the five higher-order networks that make up the SAAMs. These same general zones were emphasized more than a century ago as the regions distinguishing human and ape brains from the those of smaller monkeys (Mesulam 2015) and have been supported, based on modern comparative anatomical approaches, to be disproportionately expanded in humans relative to monkeys (Hill et al. 2010; Chaplin et al. 2013; Amlien et al. 2016; DiNicola et al. 2021). Taken together, the global spatial relations among networks (Figs. 23 to 25) and the repeating fractionation of the higher-order associations zones into five networks (Figs. 34 to 35) are consistent with processes that organize the cortex through distinct developmental stages.

**Figure 44.**
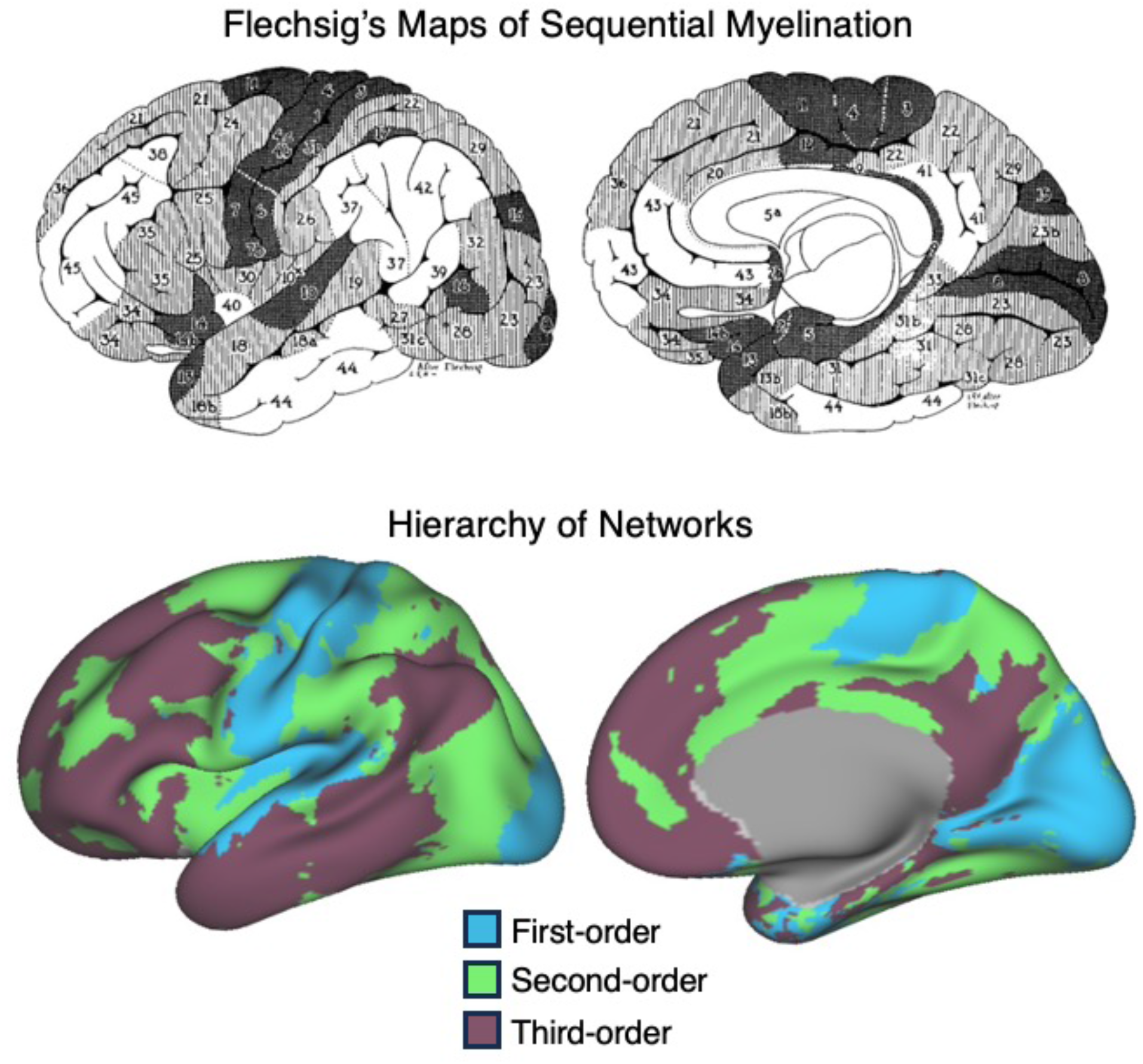
Hierarchical development might give rise to network patterning. (**Top**) The panel displays a combined rendition of Paul Flechsig’s maps of sequential myelination. Dark areas receive projections that myelinate first (before birth), gray striped areas next (during the first months of after birth), and the white areas last (starting several months after birth). Adapted from Bailey and von Bonin (1951). (**Bottom**) The present network estimates from a representative participant (P1) are recolored and grouped into first-, second-, and third-order networks to align to Flechsig’s maps. Note the similarity between the global spatial patterns and the locations of the distributed association third-order network zones and Flechsig’s zones of late myelinating, terminal fibers.

In a hypothesized first stage, cortical networks might progressively organize outwards from the early sensory and motor areas that themselves are patterned through structured inputs. For example, retinotopic organization is imparted on early visual cortex via spontaneous retinal activity waves that are present before birth and carried to the cortex through the thalamic nuclei (Katz and Shatz 1996; Ackman et al. 2012). These early organizing events may anchor the formation of the retinotopic clusters (Rosa and Tweedale 2005; Wandell et al. 2007) captured by our estimates of the VIS-C and VIS-P networks. The second-order networks may then organize tethered to these first-order networks, but with progressively more distributed regions, corresponding to Flechsig’s intermediate (or border) zones. Averbeck and colleagues have also proposed a similar nesting of networks outwards from the primary motor and somatosensory areas (S1-M1) based on extensive analyses on anatomical connectivity patterns (Giarrocco and Averbeck 2021; 2023; see also Vijayakumar et al. 2019; Du and Buckner 2021). The zones that generally fall between the regions of the second-order networks include large swaths of prefrontal, temporal, and posterior parietal association cortex that correspond to Flechsig’s terminal (or central) zones and are hypothesized to be the last to develop, forming our hypothesized third-order networks (Fig. 44). Thus, much of the cortical mantle may be patterned by a series of networks that nest outwards from the primary cortical areas (Margulies et al. 2016; Huntenburg, Bazin, and Margulies 2018; Smallwood et al. 2021).

In a second developmental stage, we hypothesize that, as the networks sequentially form, they may undergo a second process of fractionation and specialization (DiNicola and Buckner 2021). Our proposal of a distinct second process is specifically put forth to explain how juxtapositions might arise similarly across widely distributed (non-contiguous) zones of cortex, such as observed for the distinct SAAMs in LTC and PPC (Fig. 35). A specific prediction of this hypothesis is that, as development progresses, activity-dependent processes may eliminate and / or stabilize synapses that support specialization evident in the adult (for discussion see Schwartz and Goldman-Rakic 1990; Bourgeois et al. 1994; Price et al. 2006; Cadwell et al. 2019). In the cortical mantle of humans, the expanded associations zones may fractionate and specialize into the multiple juxtaposed networks that support higher-order cognition.

### Functional Specialization of Higher-Order Association Networks

By combining network estimation and task-based fMRI within the same individuals, the present results provide insight into the functional specialization of the networks. A broad observation was that the second-order distributed networks SAL / PMN and CG-OP were dissociated from the third-order association networks via their robust, transient response to oddballs (Fig. 32) consistent with prior studies (Seeley et al. 2007; see also Dosenbach et al. 2006; Seeley 2019). None of the third-order networks that populate the

SAAMs displayed a robust transient positive response. In fact, four of the five networks within the SAAMs (FPN-B, LANG, DN-B, and DN-A) showed a significant *negative* response (network FPN-A was equivocal). By contrast, all of the third-order association networks responded robustly to ongoing task demands with distinct forms of functional specialization as described below.

A first robust dissociation among the third-order networks came in their differential response to working memory demands. FPN-A and FPN-B responded to high memory load in the N-Back Load Effect task contrast and did so similarly across verbal and non-verbal materials (Fig. 39). FPN-B’s response was quantitatively lower but both FPN-A and FPN-B responded robustly across all conditions^8^. Further, FPN-A and FPN-B display the same general spatial pattern as the previously described multiple-demand network (Duncan and Owen 2000; Duncan 2010; Fedorenko, Duncan, and Kanwisher 2013; Assem et al. 2022; see also Badre and Nee 2018; Friedman and Robbins 2022). Our data are thus convergent with the existing literature to suggest there is a distributed frontal-parietal network (or networks) that responds when tasks become more effortful, perhaps related to processing functions associated with cognitive control (e.g., Miller and Cohen 2001; Badre and Nee 2018). The within-individual precision mapping allowed spatially precise network estimates to be made of FPN-A and FPN-B that predicted the idiosyncratic response patterns across participants.

A few further details are of interest. First, across most individuals, FPN-A included a small region in the anterior insula (labeled in Fig. 37). This small region showed a N-Back Load Effect response surrounded by spatially distinct components of the CG-OP and SAL / PMN networks. It would be easy to blur over or miss this buried insular region in group analysis. Our current estimates suggest that the anterior insula is a particularly challenging region of the cortex to study because multiple, distinct networks are spatially juxtaposed near to where the cortex folds onto itself in the volume. Examinations of group data, especially data that averages across participants within the volume, may be particularly vulnerable to distorting functional properties of the region.

Second, the spatially circumscribed regions of each SAAM that aligned to FPN-A and FPN-B, with some exceptions in low SNR regions, tended to show a robust N-Back Load Effect (Figs. 37 and 38). The adjacent regions of LANG, DN-B, and DN-A did not. Thus, the N-Back Load Effect functionally dissociated the FPN-A / FPN-B cluster from the LANG, DN-B, DN-A cluster multiple times across the distributed association zones including PPC, LTC, VLPFC and DLPFC.

Fedorenko and colleagues (2010; 2012) have previously noted that regions of the multiple-demand network lay side-by-side with functionally distinct domain-selective regions (specifically in the language domain). Our present results are consistent with their observations and reinforce that the functional dissociation is a general property of the association cortex including close spatial juxtapositions in temporal and parietal cortex, not only within PFC. We interpret the repeating pattern across the cortical mantle to reflect that functional specialization is a property of the networks, including all their distributed regions (see also Blank, Kanwisher, and Fedorenko 2013). Furthermore, robust functional dissociations were present for higher-order cognitive domains beyond language (see also DiNicola et al. submitted). That is, while networks LANG, DN-B, and DN-A did not modulate in a domain-flexible manner to working memory demands, each network responded robustly and selectively to a distinct specialized domain of higher-order cognition.

The most striking functional observation of the present study was the robust triple dissociation across networks LANG, DN-B, and DN-A. The LANG network responded when participant processed meaningful sentences; the DN-B network when participants engaged theory-of-mind tasks; and the DN-A network when participants remembered from their past or contemplated a future scenario. The triple dissociation was carried by a formal statistical interaction (Fig. 43) and could be visualized qualitatively on the flat maps of individual participants (Figs. 41 and 42; see also DiNicola et al. 2020; Braga et al. 2020). Considering that until recently, we and others conceptualized these zones of association cortex as being deployed flexibly across a range of higher-order cognitive domains (e.g., Buckner and Carroll 2007; Buckner et al. 2008; Spreng et al. 2009), this is a major revision to our understanding.

Our composite results suggest higher-order association cortex possesses at least three domain-specialized parallel networks supporting language, social behaviors and remembering the past and future. These domain-specialized networks are themselves separate from domain-flexible networks that participate in cognitive control. We do not know how these networks interact or whether they remain functionally separate across multiple task classes, but the robust dissociations among juxtaposed regions demonstrated here suggest that there is more modularity in association cortex, including PFC, than has typically been considered.

### Limitations and Future Directions

A key limitation of the present work is the reliance on correlational, indirect methods to infer network organization. The caveats surrounding interpreting such network estimates, and the empirical tests of their utility despite known limitations, are discussed elsewhere (Fox and Raichle 2007; Van Dijk et al. 2010; Buckner et al. 2013; Murphy et al. 2013; Smith et al. 2013; Power et al. 2014; Xue et al. 2021). Specific to the present work, it is notable that the boundaries in networks generally predicted task response patterns, bolstering confidence that valid organizational features are being described. However, there were exceptions and regions of mismatch, consistent with poor signal quality around the sinuses and inner ear (see Figs 1 and 4). Network estimates in these poorly sampled regions of cortex may be distorted.

There are also limitations to our modeling approach. In choosing the present parameters of the MS-HBM used to estimate networks, decisions were made that influence the estimates. Specifically, we choose to model 15 networks and initiated the model with a prior that arose from a group-averaged data set. As the seed-region analyses verified, the model captured within-individual correlational properties well, but not perfectly. Thus, a limitation in our current model is knowing whether one could do better and whether our specific decisions imparted bias. We assume the answer is yes to both questions. As our own work has evolved from a relatively crude 7-network estimate in average participant groups (Yeo et al. 2011) to a 10-network estimate within individuals (Xue et al. 2021), we expect the current network estimates will be refined further and eventually replaced. As a specific example, it is unclear that the present model fully captures the details of the recently described inter-effector connectivity pattern (Gordon et al. 2023)5. The structured correlations they observed, and we also find, are partially incorporated (perhaps erroneously) in our estimate of the CG-OP network (Figs. 20 and 21) but not entirely (e.g., see diamond in Fig. 27). With the expectation of further improvements in mind, we are struck by how the present network parcellation captures multiple functional dissociations prospectively in task data, including idiosyncratic and small regions of response.

Another limitation is that, while the task contrasts allowed for robust functional dissociations, the tasks were designed and implemented to differentiate networks, which is a different goal than interrogating in detail a hypothesized cognitive operation. That is, the limited task data we collected falls far short of systematically manipulating variables to clarify the component computations performed by each of the networks. In this sense, the robust empirical dissociations found here are positive evidence that networks perform distinct functions, but further work is needed to understand the nature of the processes.

Another future direction pertains to the need to better understand the relation of traditional area estimates with the present network estimates. By “area” we mean the demarcation of regions of cortex as separate, defined zones using functional, architectonic, connectivity, and topographic constraints (Kaas 1987; Felleman and Van Essen 1991). We previously noted discrepancies between functional connectivity patterns and areal boundaries (e.g., Yeo et al. 2011; Buckner and Yeo 2014; Buckner and DiNicola 2019) as have others (e.g., Van Essen and Glasser 2014). There are two topics to be considered.

First, for regions of cortex that have well recognized areas, our network borders do not align with the areal borders (e.g., Fig. 26E). For example, networks VIS-C and VIS-P group together V1, V2, and V3 and split them roughly along the 5° eccentricity line. The estimated networks likely reflect the dominant anatomical connectivity gradient that, within early visual areas, progresses along eccentricity (Maunsell and Van Essen 1983; Ungerleider et al. 2014; see also Falchier et al. 2002). The V1/V2/V3 areal boundaries are distinguished by a local reversal in polar angle along the horizontal meridian. Thus, connectivity transitions between early visual areas are relatively subtle (for further discussion of this issue see Buckner and Yeo

2014). The somatomotor networks similarly group M1 / S1 and multiple body maps that span architectonically distinct areas (Hatanaka et al. 2001). One future direction is to understand the relation of the networks estimated here and the finer-scale anatomical differences that demarcate adjacent areas.

The second related topic is the relation between the present network estimates and architectonic features in higher-order association cortex. This is a trickier topic. Varied perspectives have been put forth on whether association cortex possesses sharp areal boundaries that parallel those found in sensory systems (for discussion see Bailey and von Bonin 1951; Rosa 2002; Rosa et al. 2005; Buckner and DiNicola 2019). There is also an open question of whether, in practice, there are known stable features that can define areal borders in association cortex, especially when architectonics are considered in isolation (e.g., Lashley and Cark 1946; Bailey and von Bonin 1951). We will not resolve the debate here, but some of our observations are relevant to the discussion.

Most critically, the extent and complexity of the network juxtapositions encompassed within the SAAMs are of such a spatial scale that they seem unlikely to align to traditional architectonic borders, at least those reflected in any of the commonly used atlases. In the spirit of supra-areal clusters reported in the visual system (Buckner and Yeo 2014; Arcaro et al. 2015; see also Rosa and Tweedale 2005; Wandell et al. 2007), we refer to the repeated groupings of multiple association networks as Supra-Areal Association Megaclusters specifically to reinforce the possibility that they might span and split traditional architectonic patterns. One possibility is that future advances will find architectonic features that align to the transitions between SAAMs as well as between the multiple network regions within the SAMMs (perhaps via spatial transcriptomics; Nano et al. 2021). Alternatively, there may be broad patterning forces during development, such as captured in Flechsig’s maps of sequential myelination, that reflect processes that guide where SAAMs develop, but that do not specify the details of the borders and regional specializations within the SAAMs. The local spatial arrangements might be carried by extrinsic anatomical connectivity differences that refine relatively late in development through activity-dependent processes, without rigid alignment to architectonic features (Buckner and DiNicola 2019; DiNicola and Buckner 2021).

Data to inform these and other possibilities will emerge as the field charts development of association networks in non-human primates with direct anatomical techniques and in human infants using non-invasive approaches.

## Conclusions

The present study examined the organization of cerebral networks within intensively sampled individual participants. We provide the resulting network estimates and the raw data used to derive them as open resources for the community. Our initial explorations on the data uncovered a hierarchical organization which distinguishes three levels of cortical hierarchy: first-, second-, and third-order networks. Notably, regions of distinct third-order association networks consistently displayed side-by-side juxtapositions that repeated across multiple cortical zones, with clear and robust functional specialization among the embedded regions.

## Acknowledgments

We thank the Harvard Center for Brain Science neuroimaging core and FAS Division of Research Computing. We thank T. O’Keefe for assisting in optimization of data processing and R. Mair for MRI physics support. H. Kosakowski provided valuable discussion. K. Ntoh and F. Davy-Falconi assisted with data acquisition. E. Fedorenko, T. Konkle, and R. Saxe generously provided task stimuli. The multi-band EPI sequence was generously provided by the Center for Magnetic Resonance Research (CMRR) at the University of Minnesota.

## Footnotes

1 We term these networks somatomotor (SMOT) networks because each comprises both regions of somatosensory cortex and regions of motor cortex.

2 Network labels use conventions that often reflect historical origins and diverge from current understanding. For example, the canonical Default Network, originally identified in group-based positron emission tomography (PET) data, is now postulated to comprise multiple, distinct networks (see Buckner and DiNicola 2019 for review). The names, DN-A and DN-B, reflect the historical naming convention modified to the current understanding of multiple networks. As another example, the network labeled here as SAL / PMN has two distinct origins. Seeley et al. (2007) referred to the network as the Salience Network and Gilmore et al. (2015) as the Parietal Memory Network. Ideas about network organization and function are continuously evolving, while the labels often reflect historical (not contemporary) understanding.

3 The Frontoparietal Network (FPN) has been fractionated into distinct, parallel networks in multiple prior studies, but has not been consistently named. Here we label FPN-A and FPN-B to be consistent with the order (A/B naming convention) of Kong et al. (2019) and Xue et al. (2021) who also applied an MS-HBM to estimate networks. We caution the reader that other studies have used the reverse convention, which could lead to confusion (e.g., Braga et al. 2020; DiNicola et al. 2023).

4 Pilot analyses were conducted to test whether an individual’s network estimate was influenced by the group in which the participant was analyzed. In our explorations, the individual’s parcellation was nearly identical whether the participant was grouped with one set of other individuals or another set. We do not assume this will always be the case, as our analyses were conducted for a group of healthy young adult participants with large amounts of data.

5 Gordon et al. (2023) recently described a set of inter-effector regions along the central sulcus, which was not explicitly incorporated into our group prior or model. Examining the details of Figs. 3 and 6 shows that the correlation patterns from the seed regions placed within the CG-OP extend farther into the motor strip than the MS-HBM defined CG-OP network regions themselves. This could be due to blurring or, alternatively, that inter-effector regions are not properly distinguished by the present model.

6 Any heuristic framework will necessarily emphasize certain features of organization and deemphasize others. A three-level hierarchy, while not capturing local features of organization, is a useful framework to emphasize aspects of global organization that are the focus of this paper and, as will be discussed, converges with similar frameworks that have arisen from direct anatomical description (e.g., Flechsig 1904; 1920). Alternative organizational schemes are possible, and the three-level hierarchy proposed should viewed as an orienting framework.

7 While the higher-order association networks were generally positioned between the second-order networks, an exception to that pattern is that the LANG network juxtaposes the AUD network, without second-order networks interdigitated. It is unclear whether this is a true exception, or there are local organizational details that are not resolved by our current methods.

8 We do not yet interpret the differential response between FPN-A and FPN-B as there was no condition where FPN-B responded more than FPN-A. Main effects in BOLD response magnitude between regions can come from any number of technical reasons including the regional vasculature sampled and the inclusion of voxels with susceptibility artifact. We thus conservatively interpret differential responses when there is direct evidence for a double dissociation within or across task contrasts (following the logic of Shallice 1988).

