## Supplemental Materials for "Within-Individual Organization of the Human Cerebral Cortex: Networks, Global Topography, and Function"

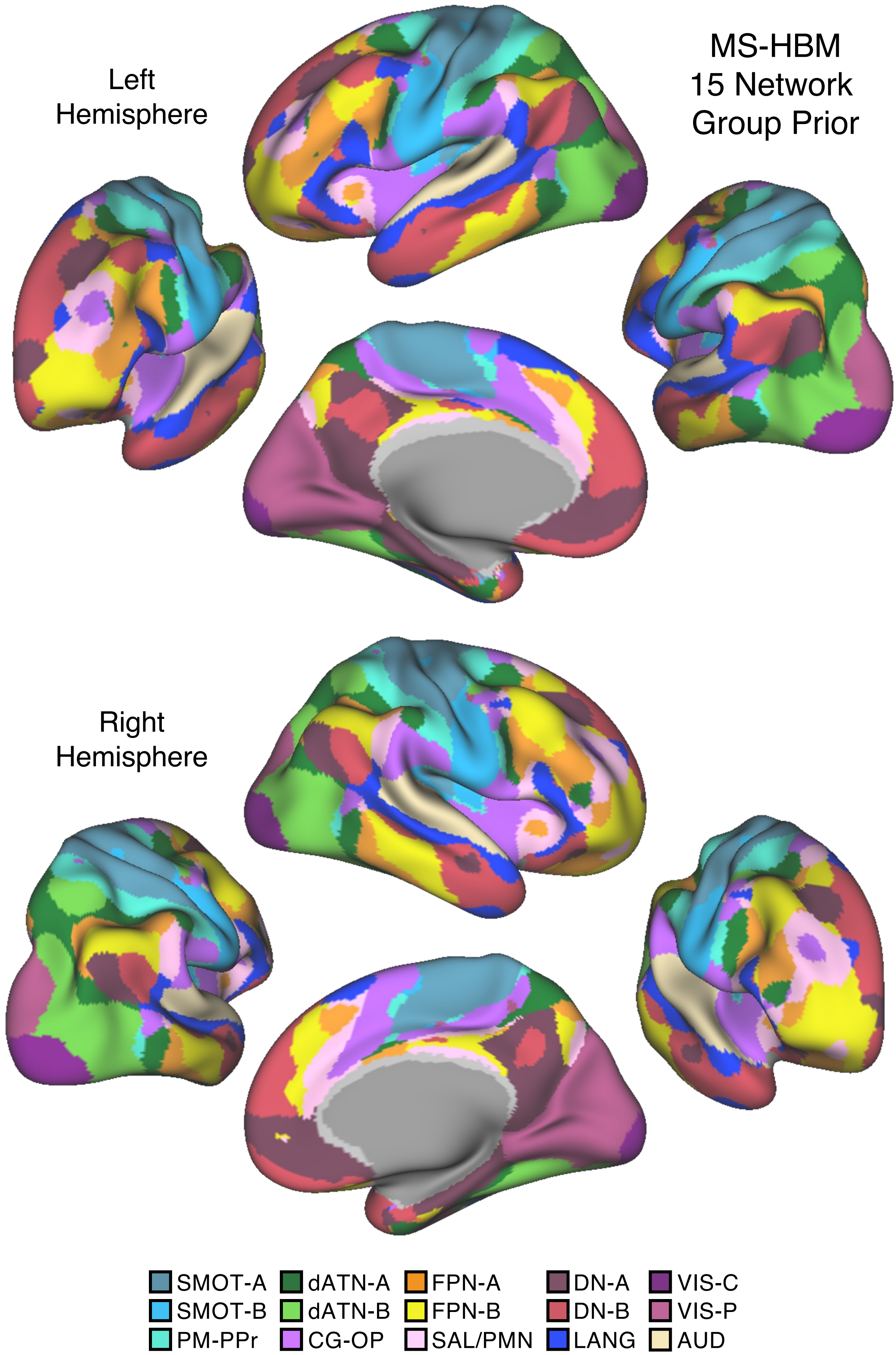
Left
Hemisphere
MS-HBM
15 Network
Group Prior
Right
Hemisphere
SMOT-A
SMOT-B
PM-PPr
dATN-A
dATN-B
CG-OP
FPN-A
FPN-B
SAL/PMN
DN-A
DN-B
LANG
VIS-C
VIS-P
AUD

Figure S1

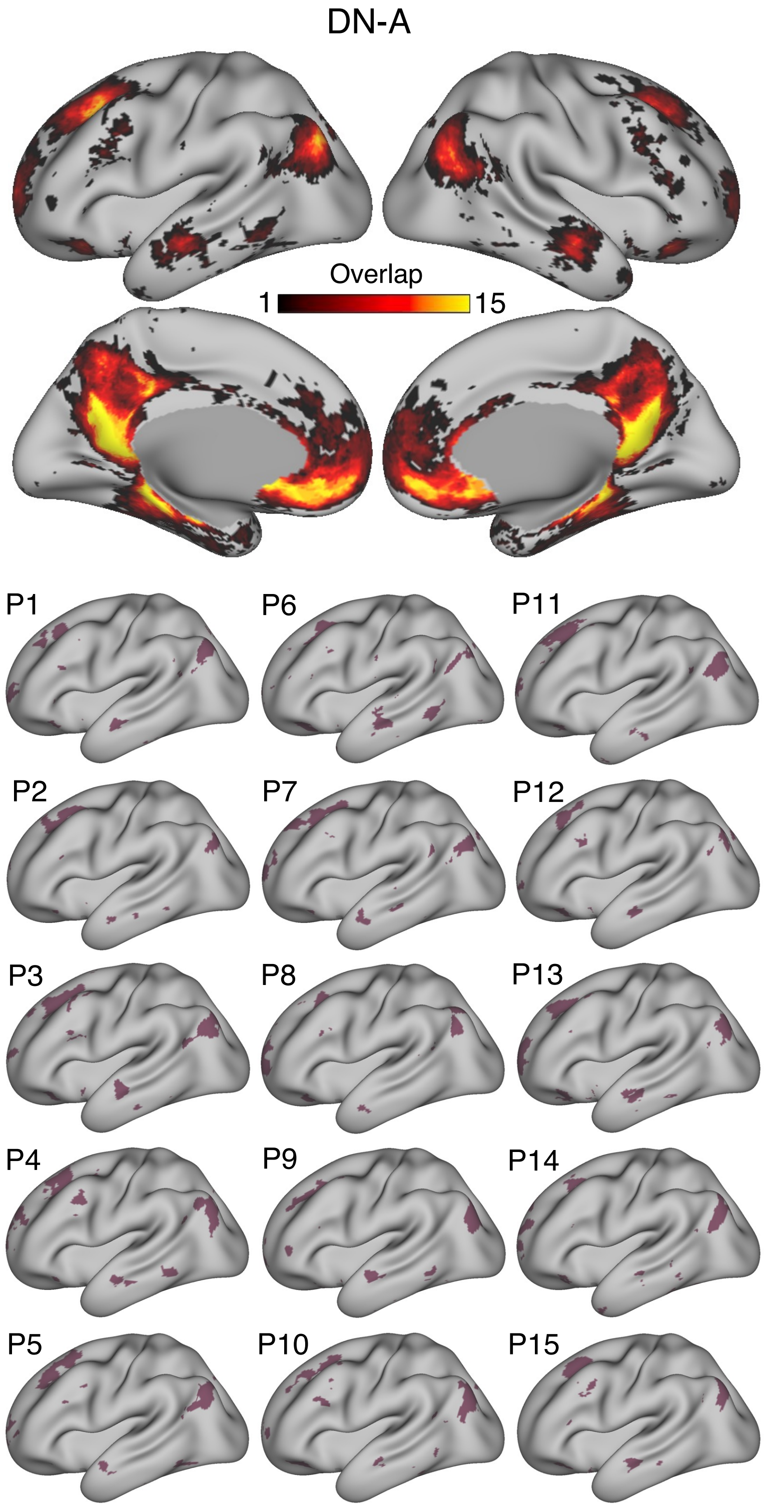


Figure S2

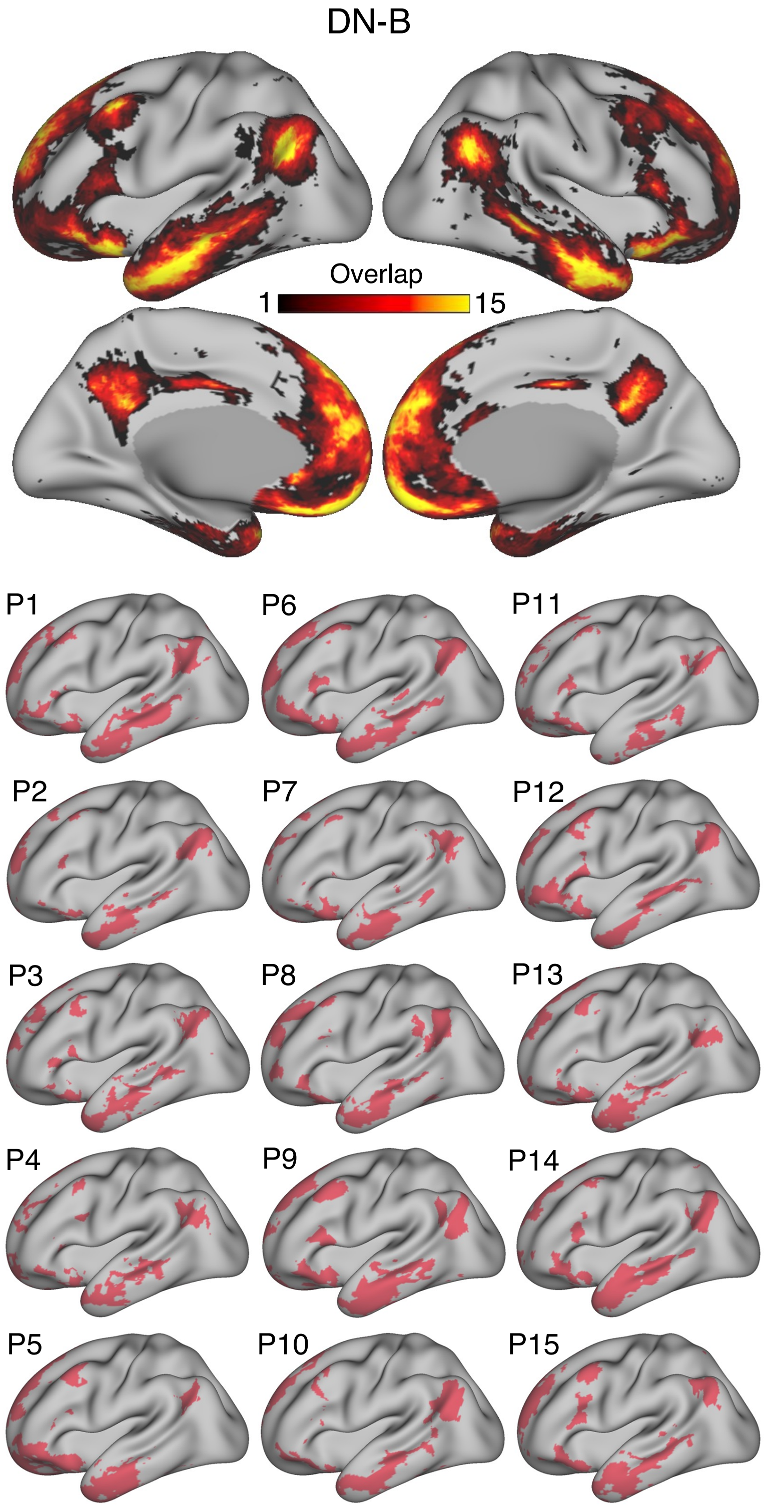


Figure S3

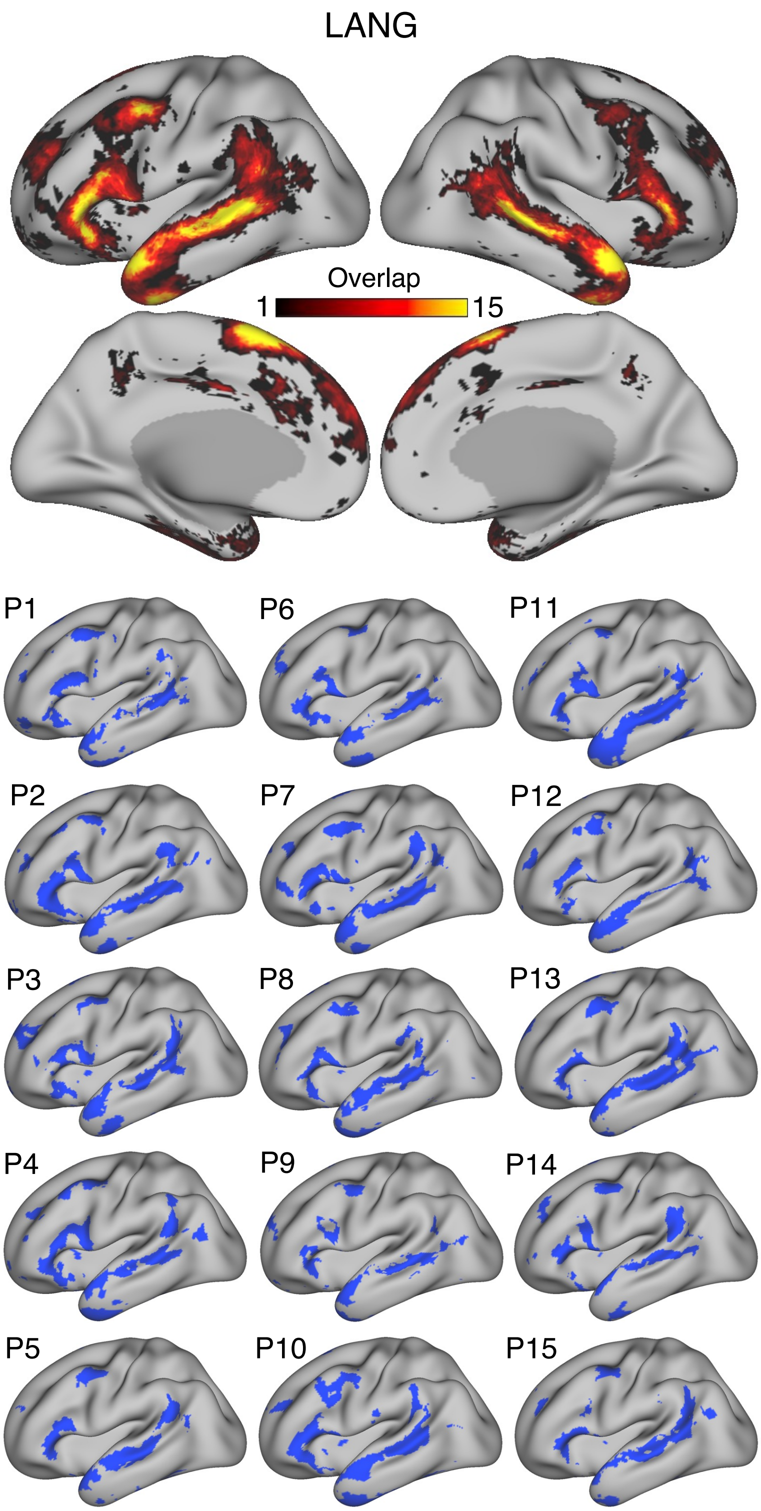

LANG
Overlap
1
15
P1
P6
P11
P2
P7
P12
P3
P8
P13
P4
P9
P14
P5
P10
P15


Figure S4

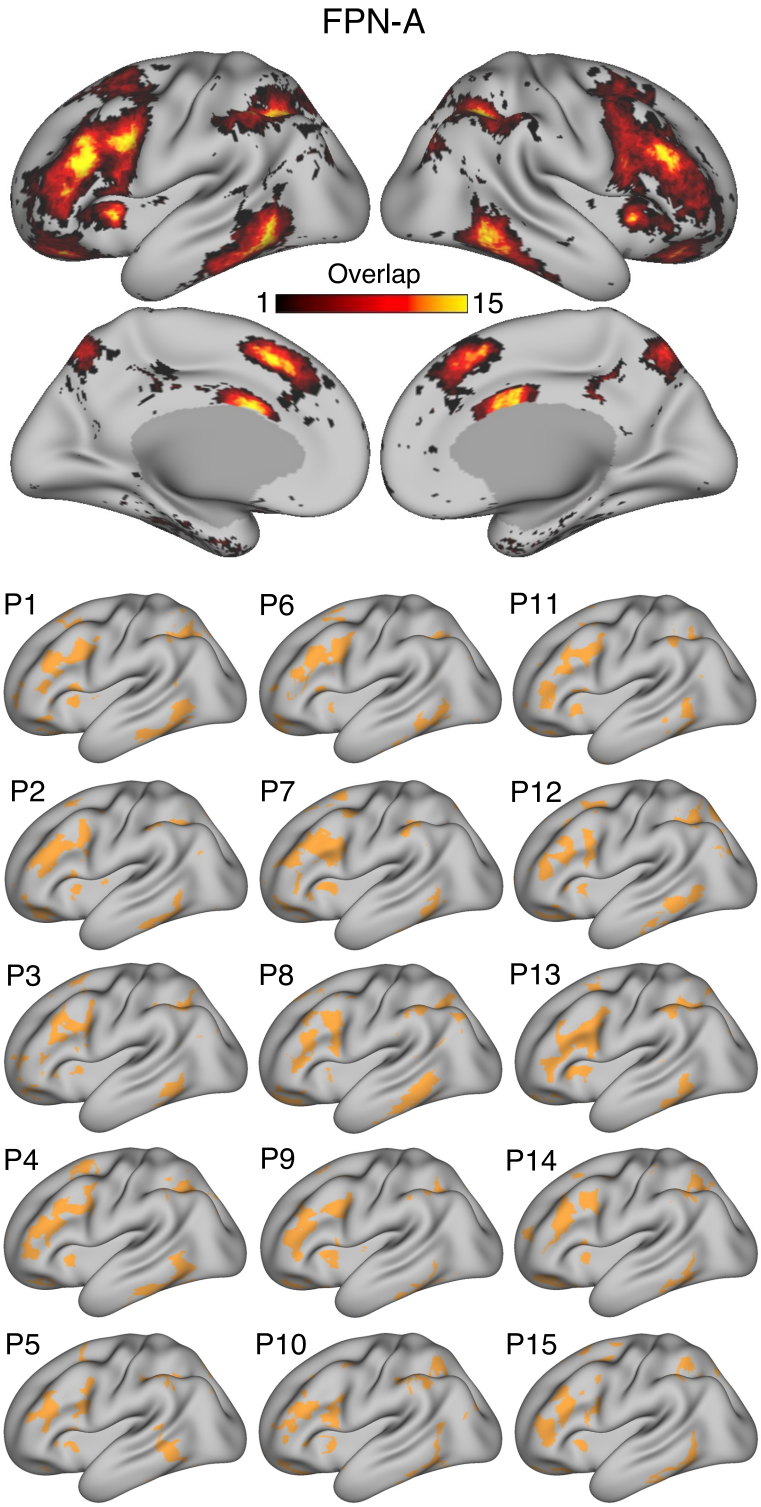
FPN-A
Overlap
1
15
P1
P6
P11
P2
P7
P12
P3
P8
P13
P4
P9
P14
P5
P10
P15

Figure S5

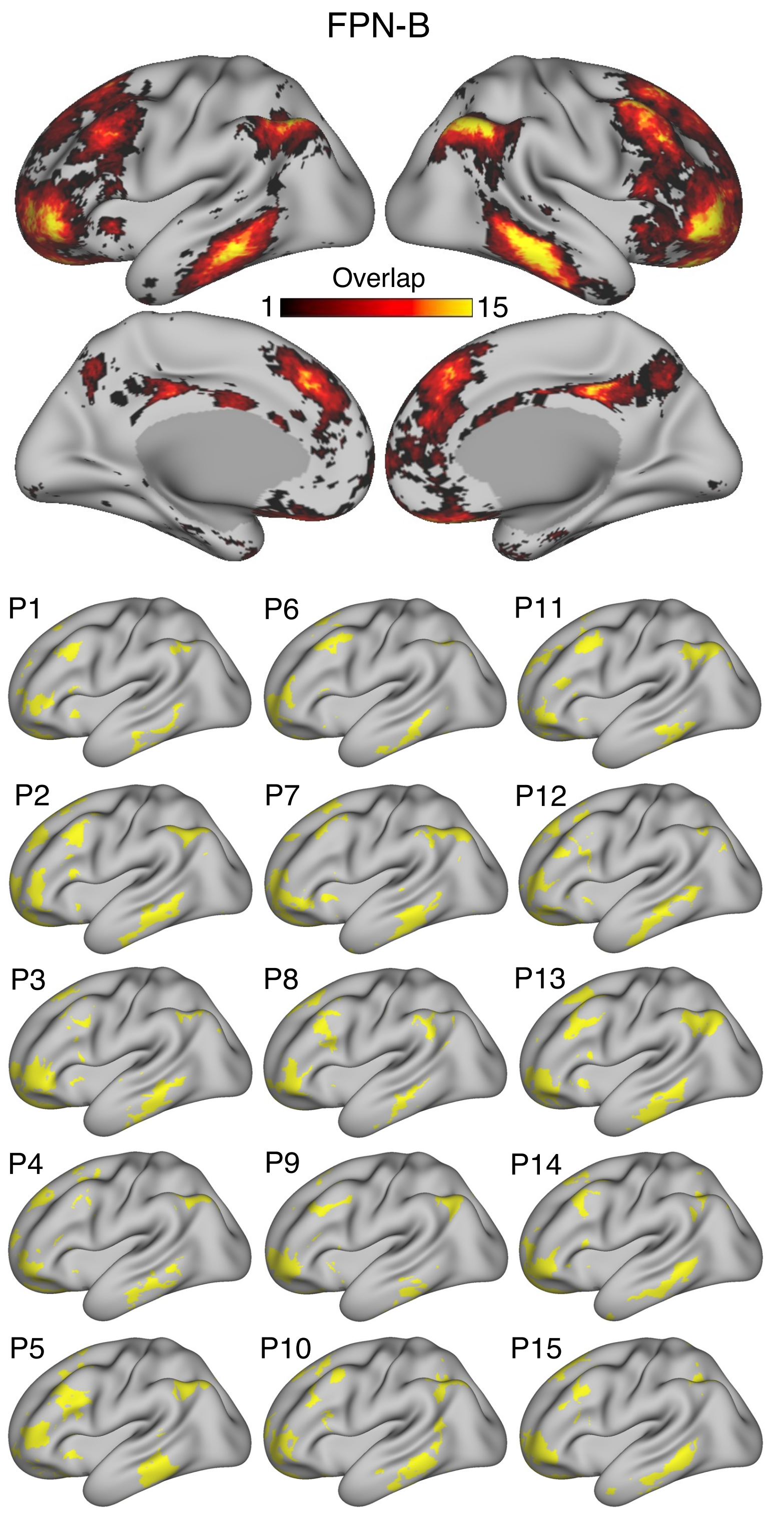
FPN-B
Overlap
1
15
P1
P6
P11
P2
P7
P12
P3
P8
P13
P4
P9
P14
P5
P10
P15

Figure S6

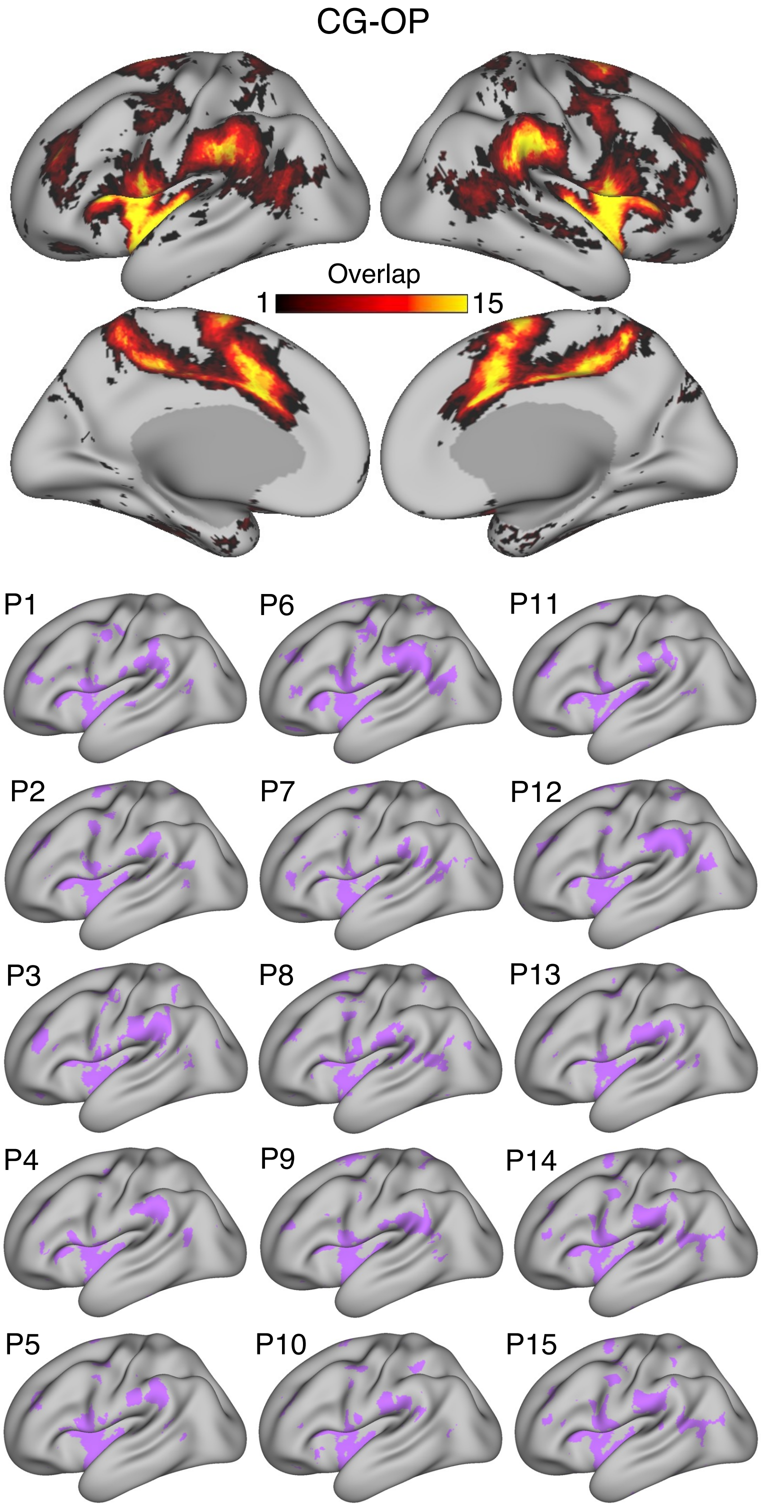
CG-OP
Overlap
1
15
P1
P6
P11
P2
P7
P12
P3
P8
P13
P4
P9
P14
P5
P10
P15

Figure S7

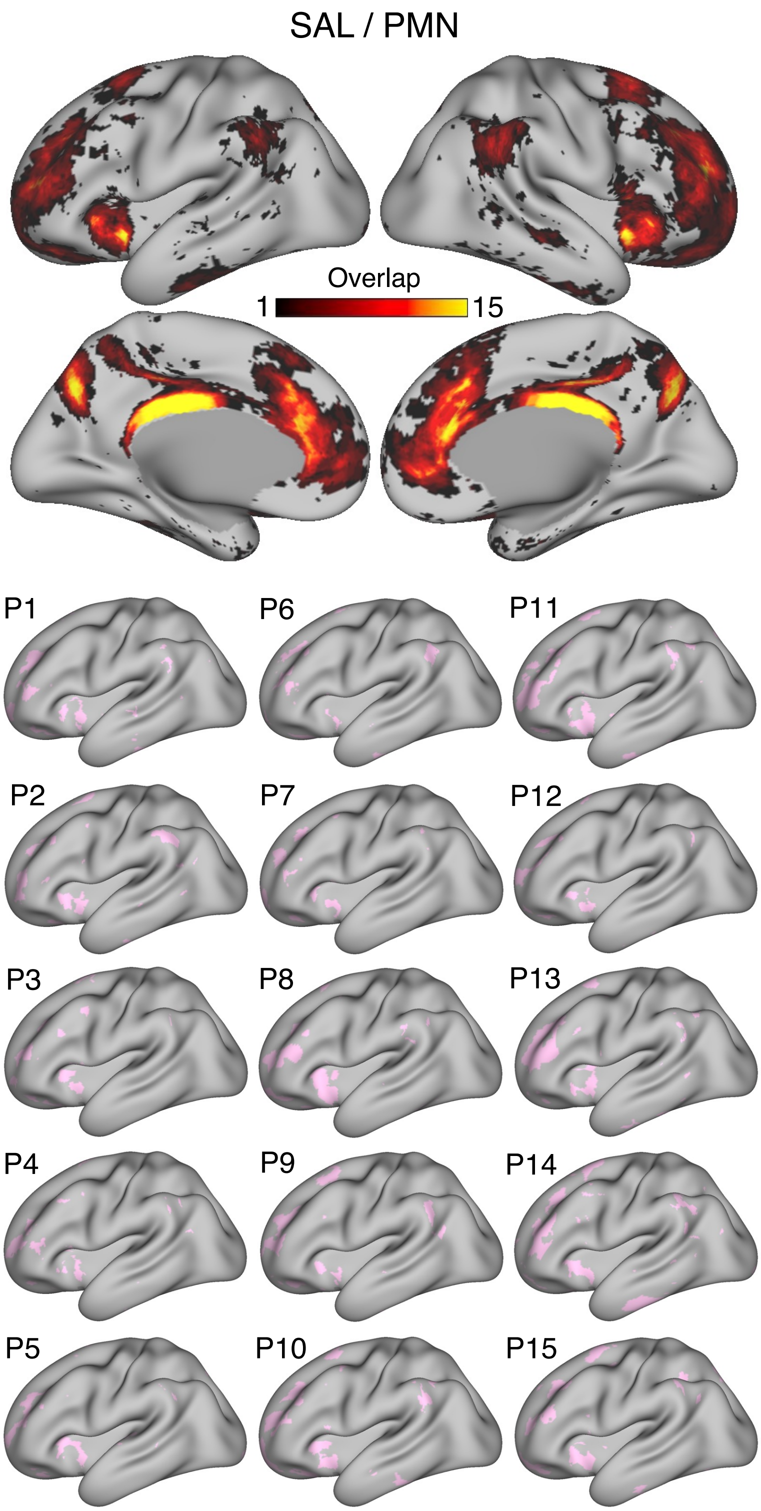
SAL / PMN
Overlap
1
15
P1
P6
P11
P2
P7
P12
P3
P8
P13
P4
P9
P14
P5
P10
P15

Figure S8

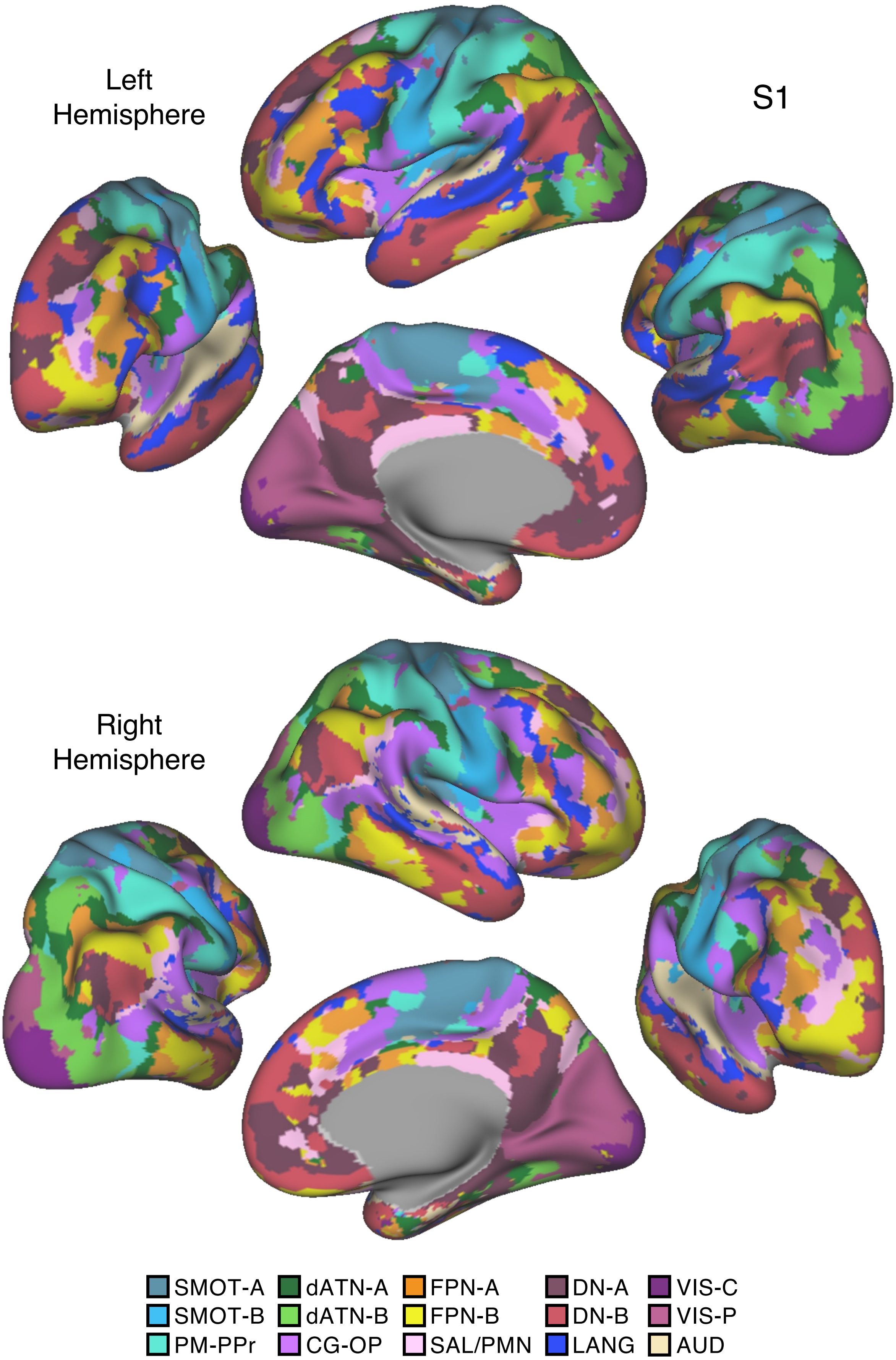

Left
Hemisphere
S1
Right
Hemisphere
SMOT-A
SMOT-B
PM-PPr
dATN-A
dATN-B
CG-OP
FPN-A
FPN-B
SAL/PMN
DN-A
DN-B
LANG
VIS-C
VIS-P
AUD


Figure S9

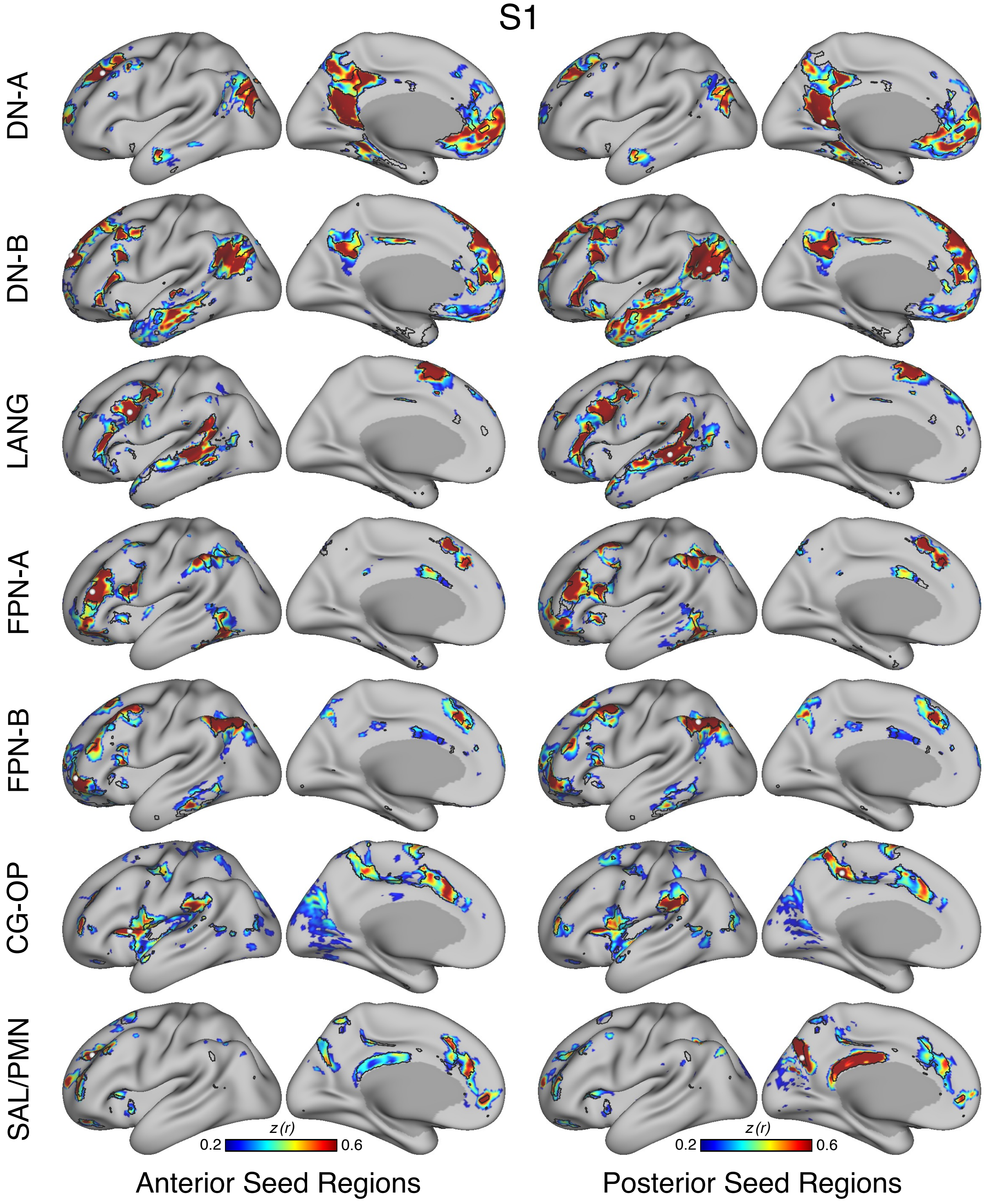
S1
DN-A
DN-B
LANG
FPN-A
FPN-B
CG-OP
SAL/PMN
z (r)
0.2
0.6
z (r)
0.2
0.6
Anterior Seed Regions
Posterior Seed Regions

Figure S10

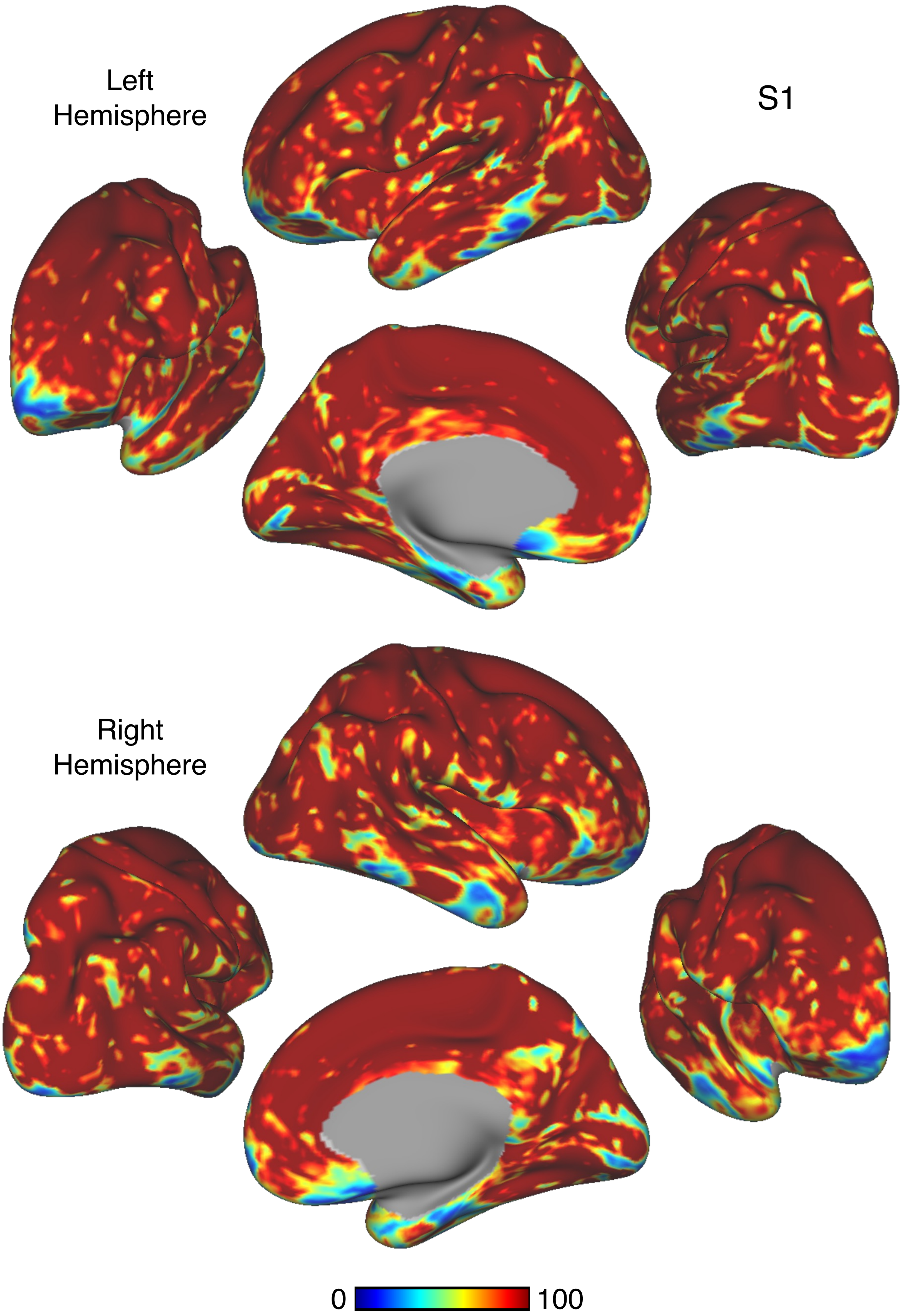


Figure S11

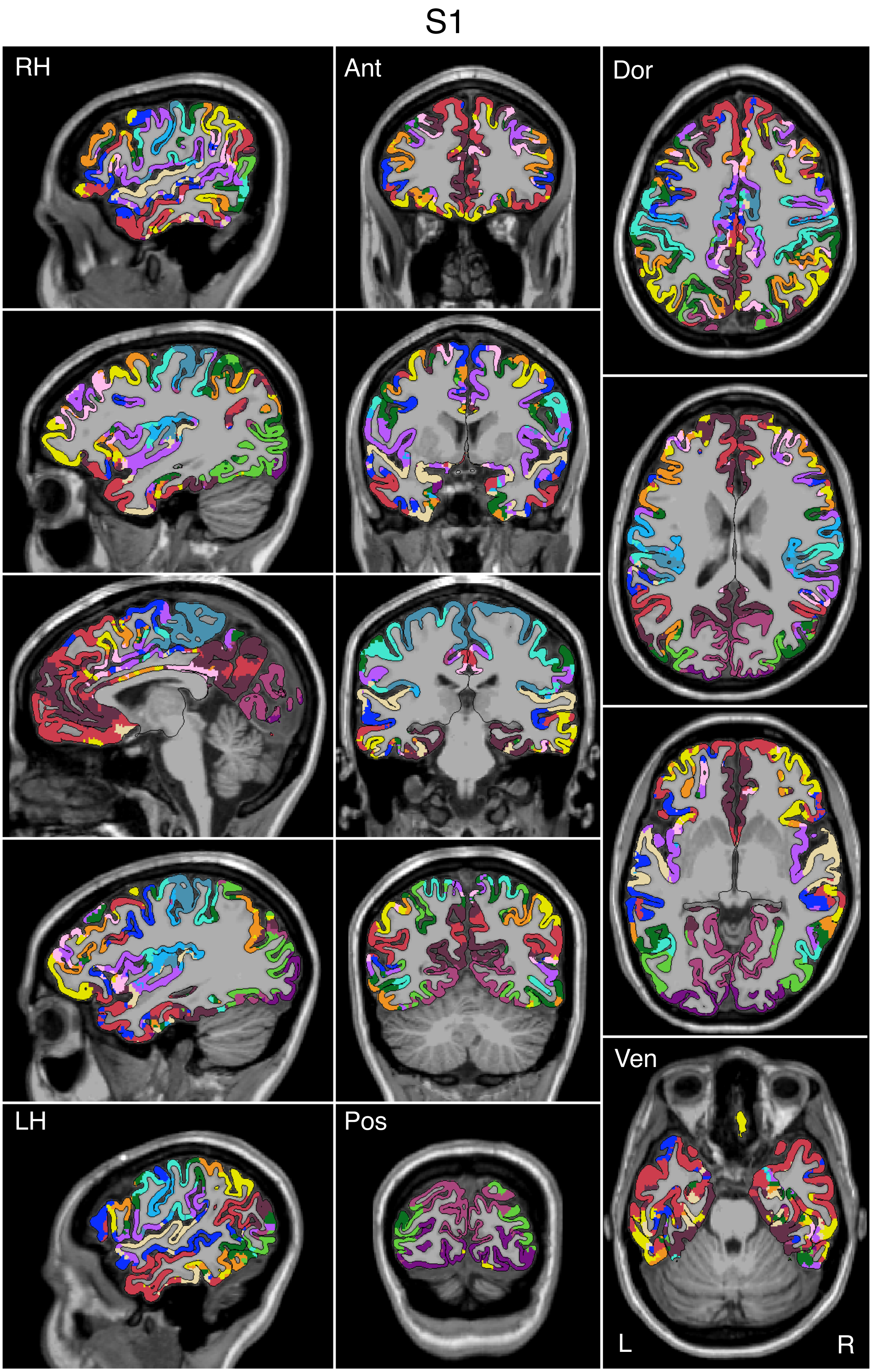


Figure S12

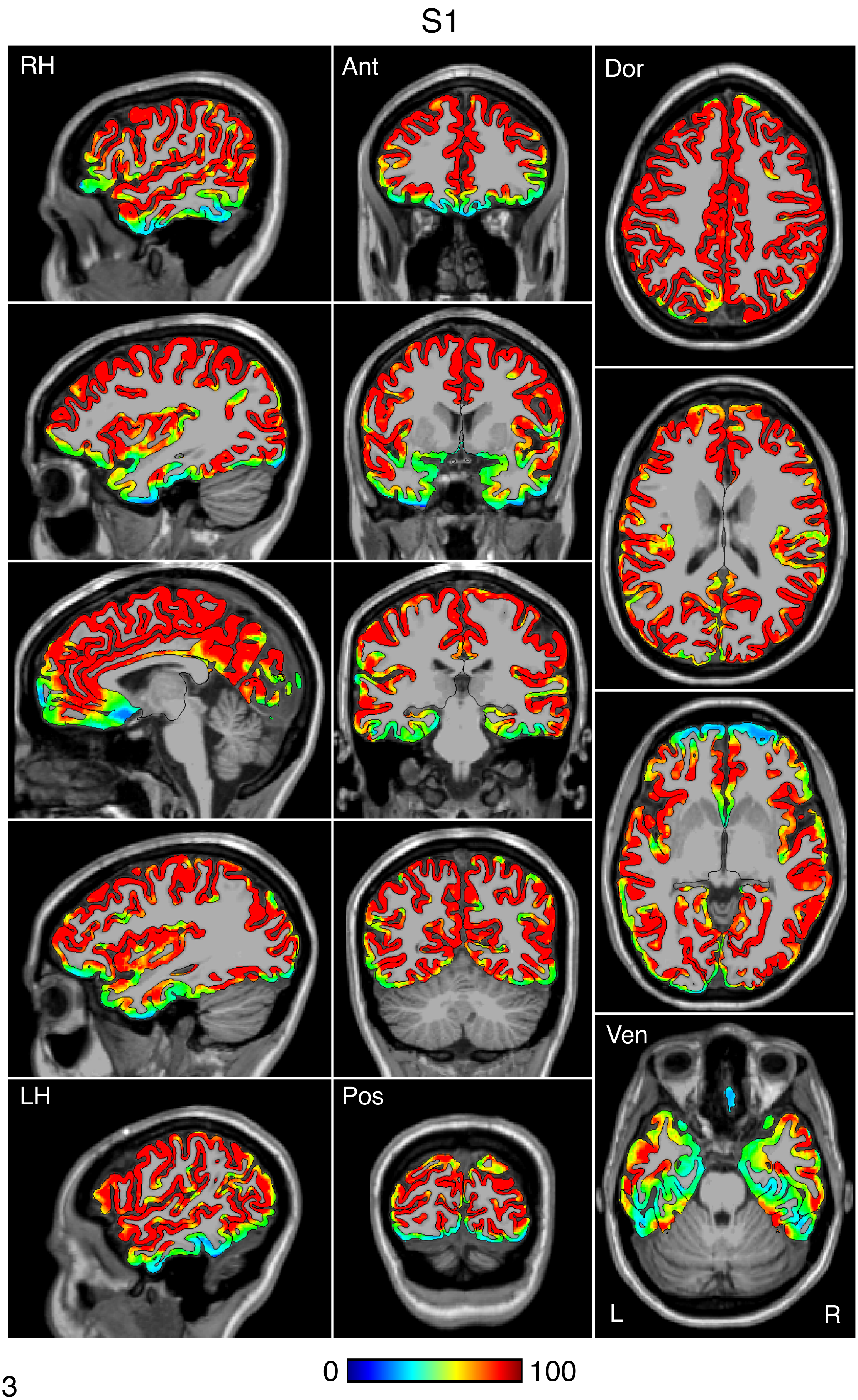


Figure S13

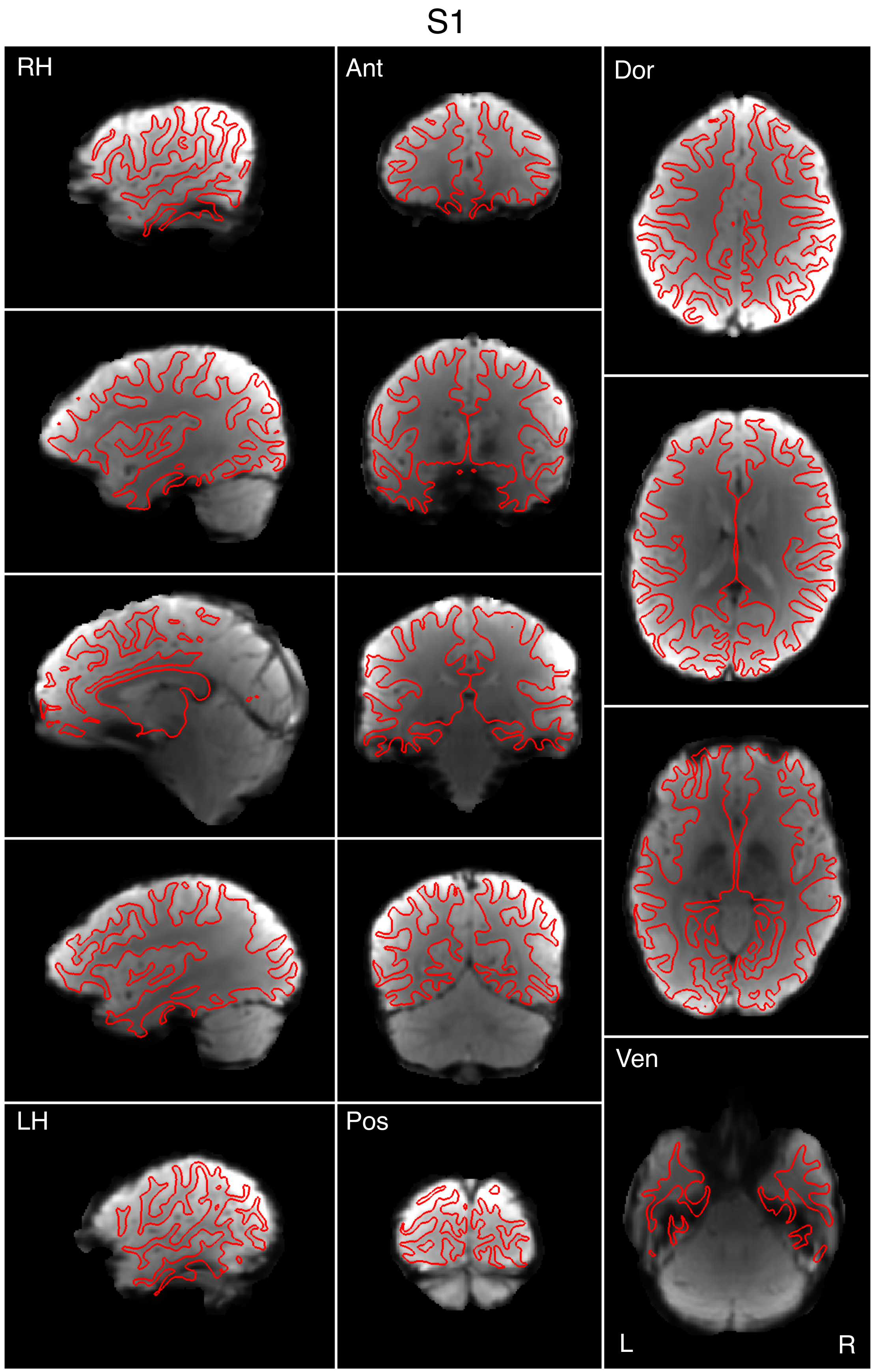


Figure S14

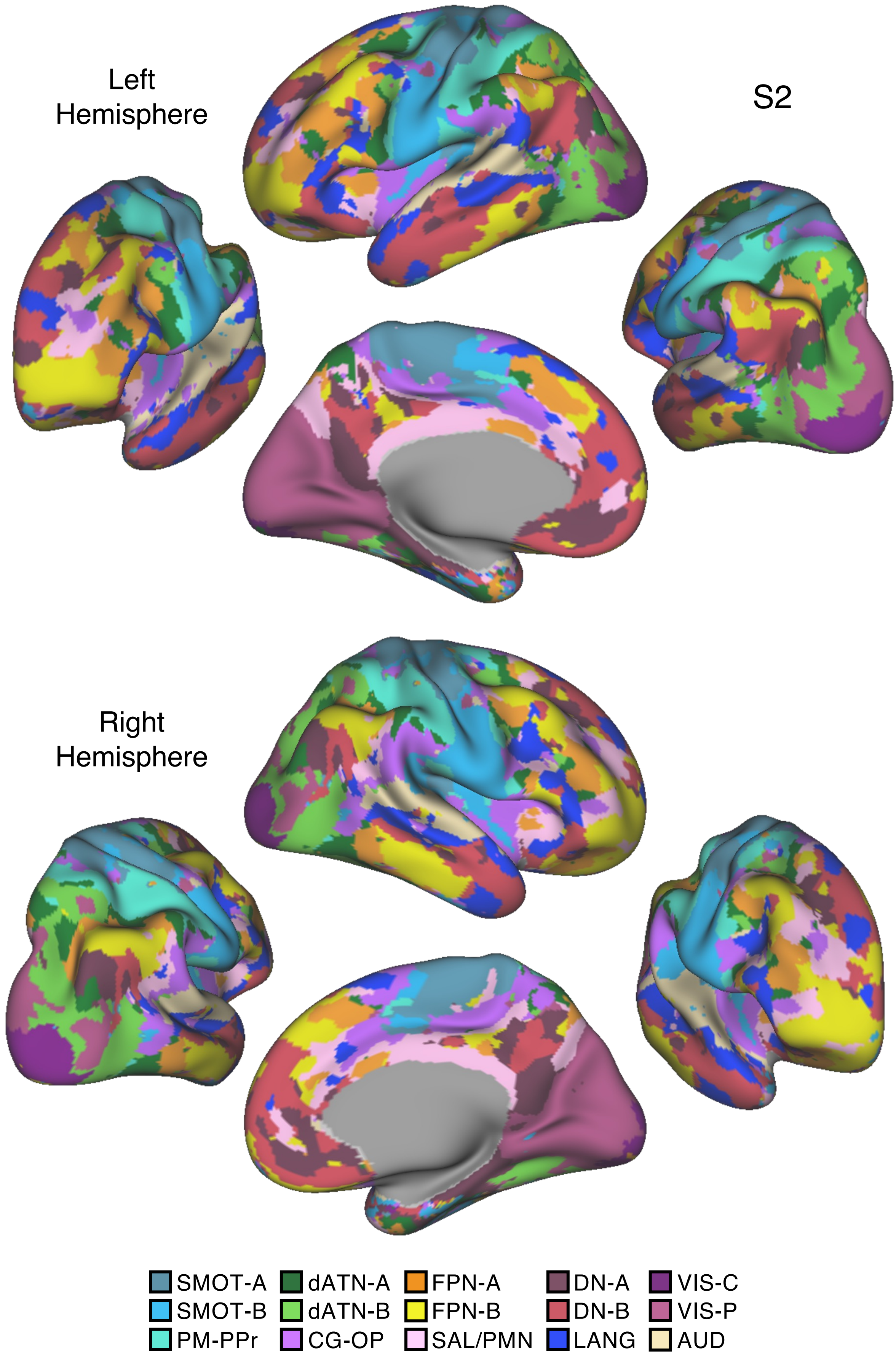
Left Hemisphere
S2
Right Hemisphere
SMOT-A
SMOT-B
PM-PPr
dATN-A
dATN-B
CG-OP
FPN-A
FPN-B
SAL/PMN
DN-A
DN-B
LANG
VIS-C
VIS-P
AUD

Figure S15

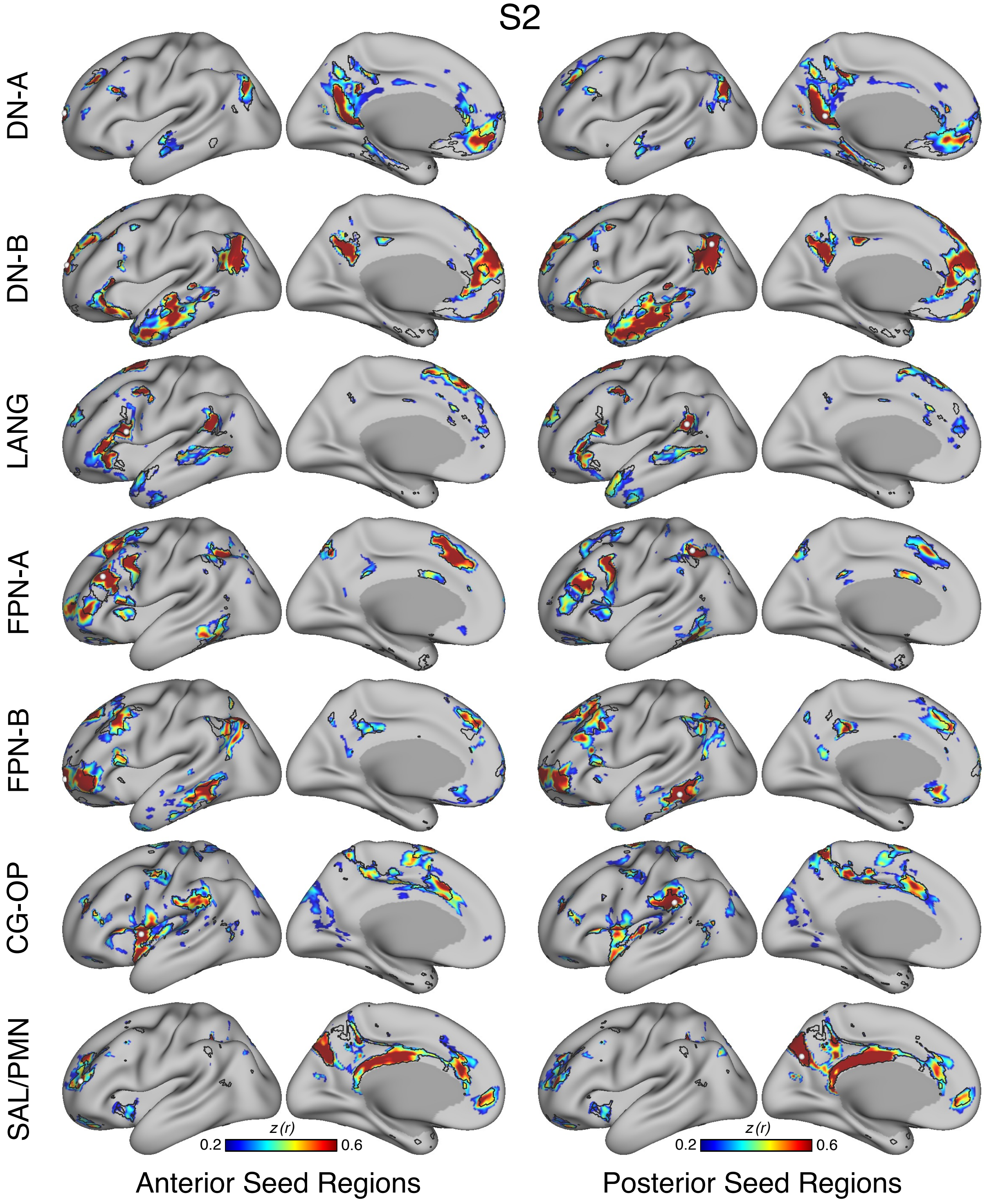
S2
DN-A
DN-B
LANG
FPN-A
FPN-B
CG-OP
SAL/PMN
z (r)
0.2
0.6
z (r)
0.2
0.6
Anterior Seed Regions
Posterior Seed Regions

Figure S16

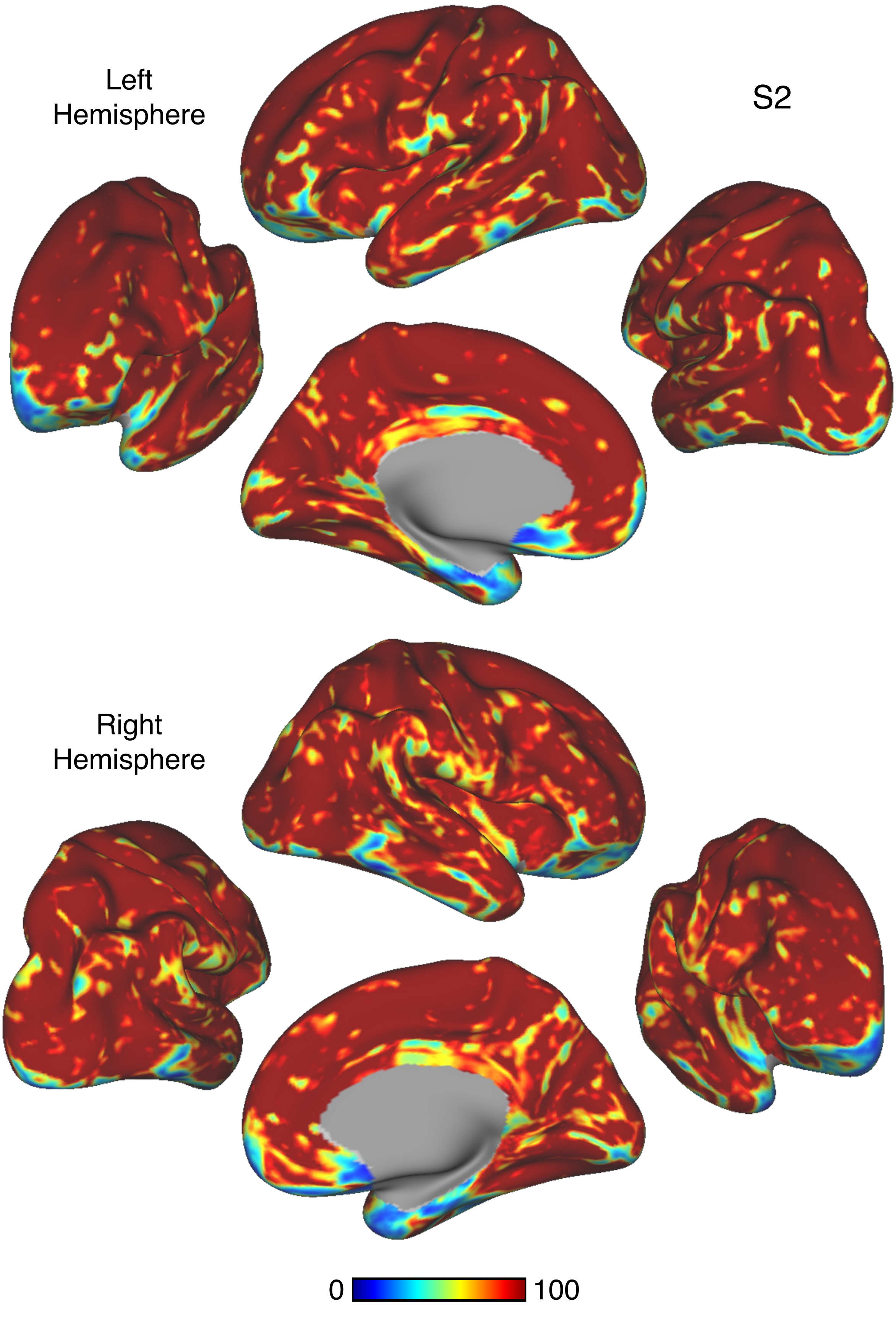


Figure S17

S2

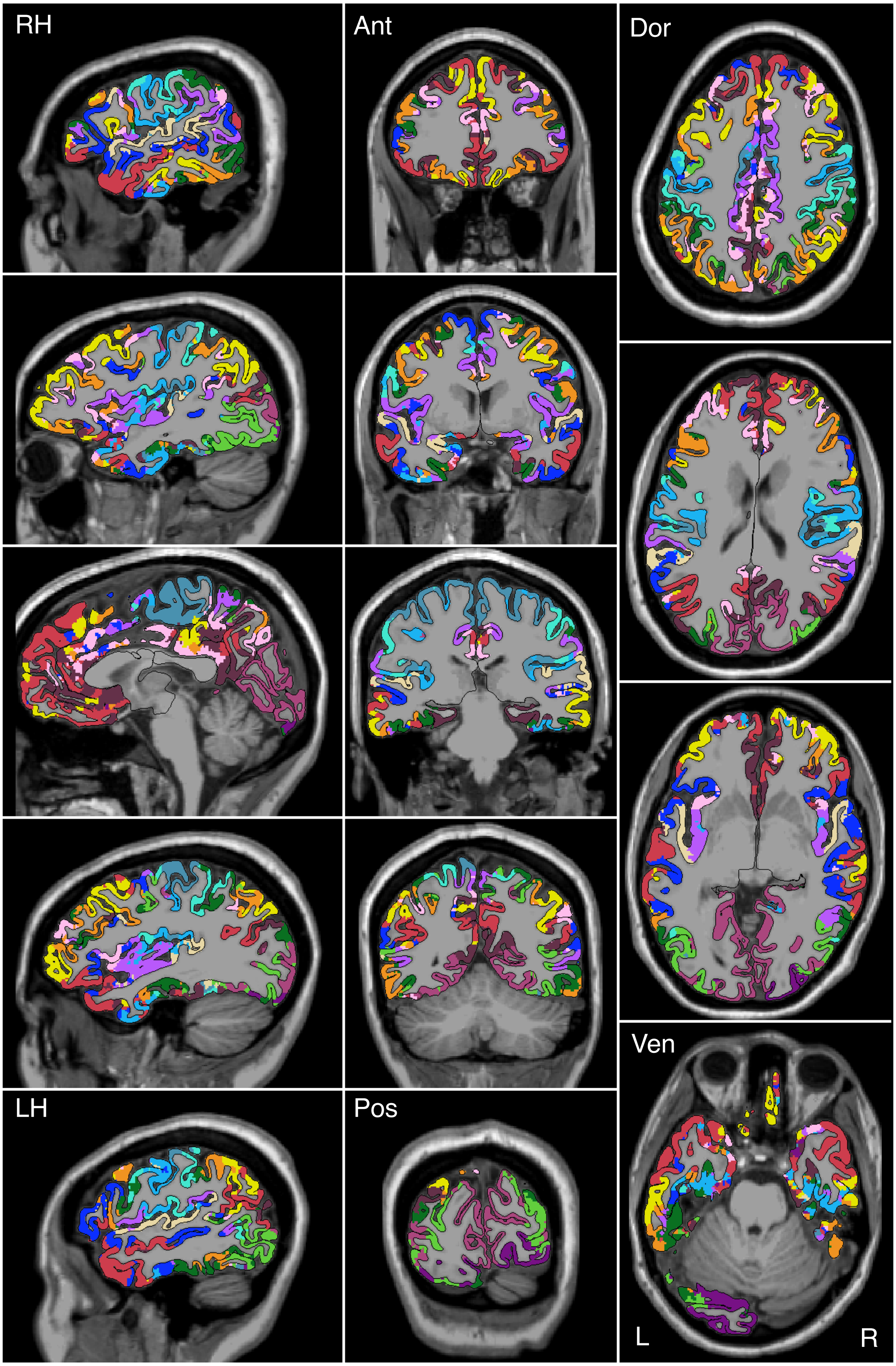


Figure S18

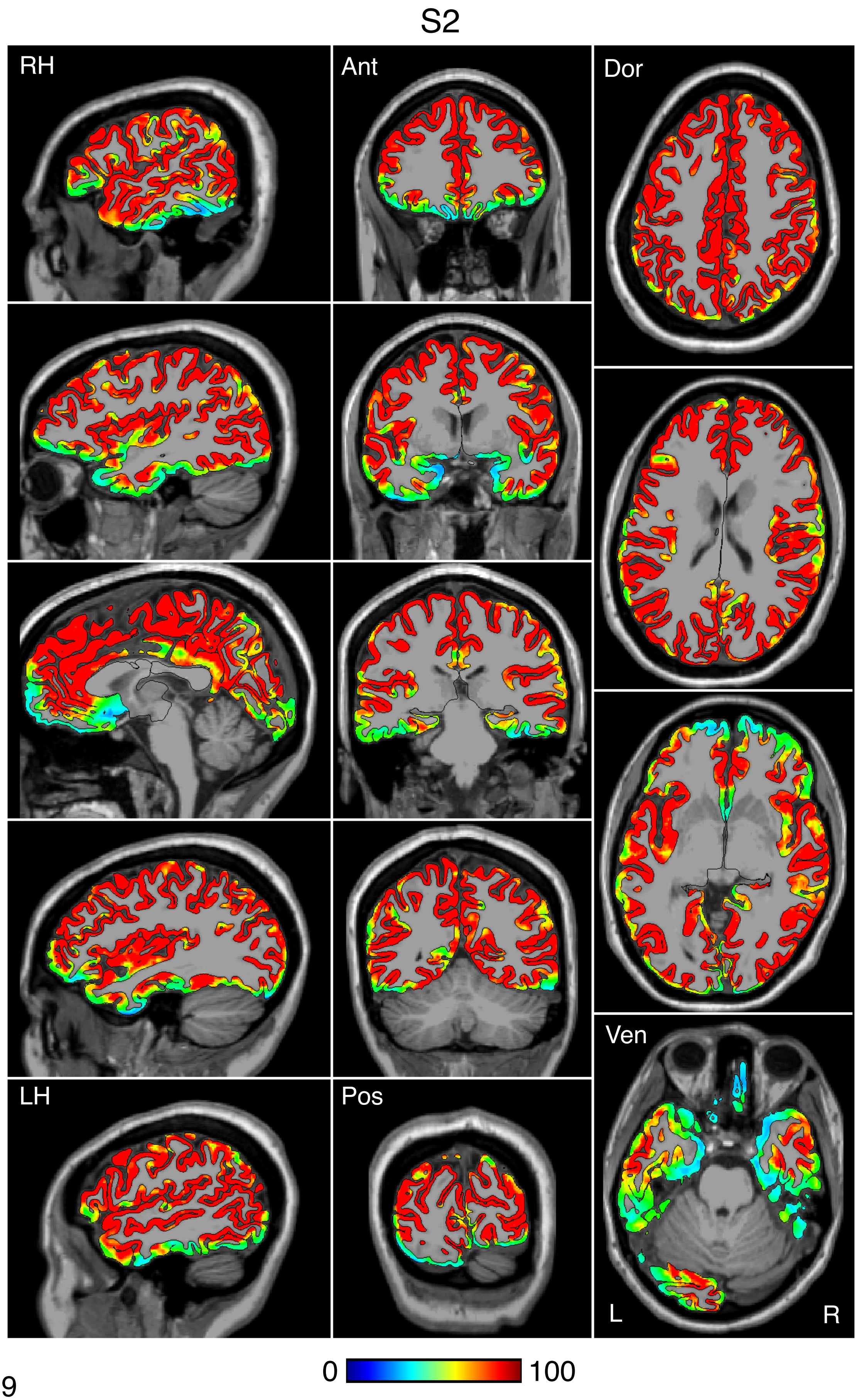


Figure S19

S2

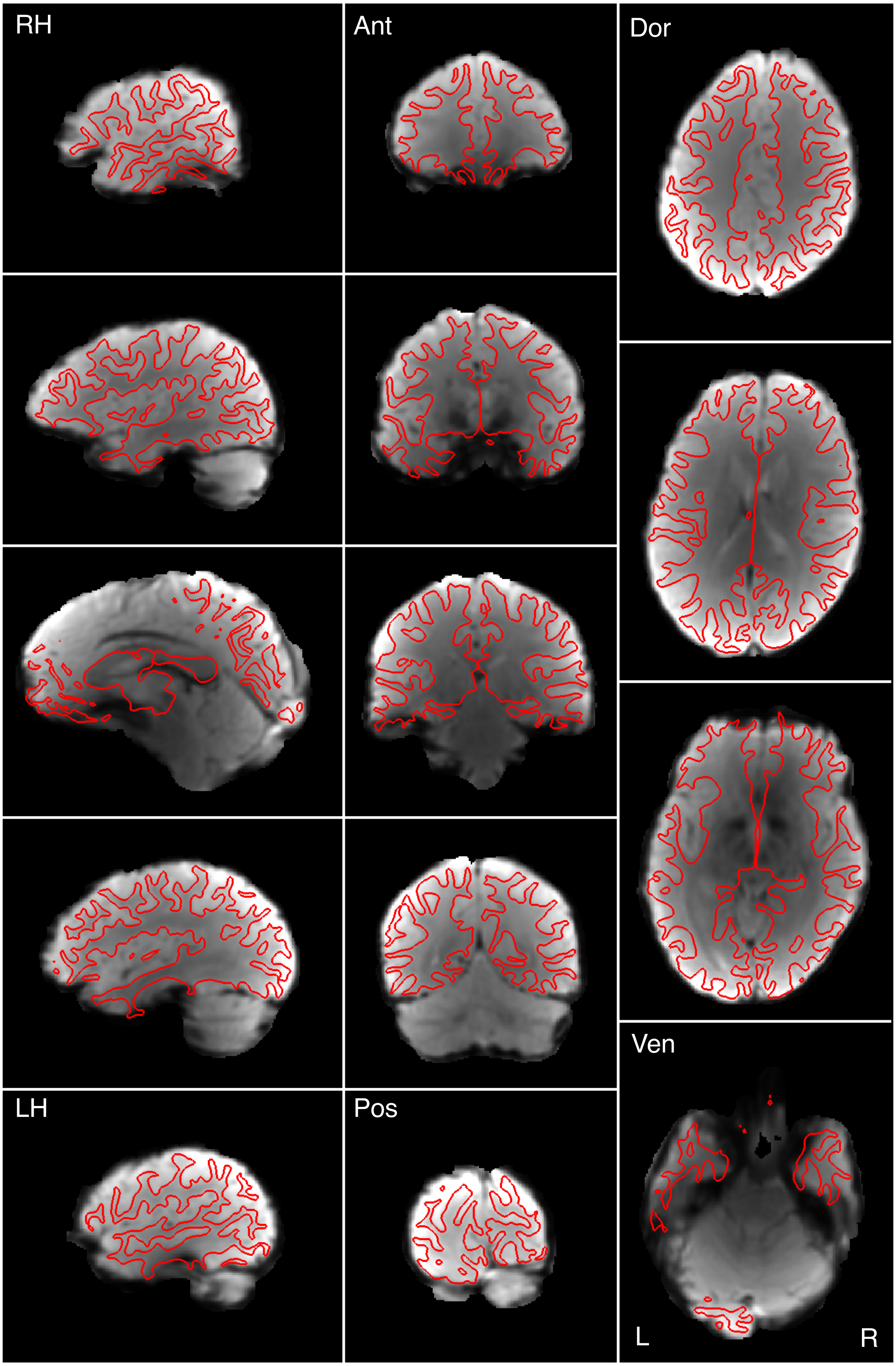


Figure S20

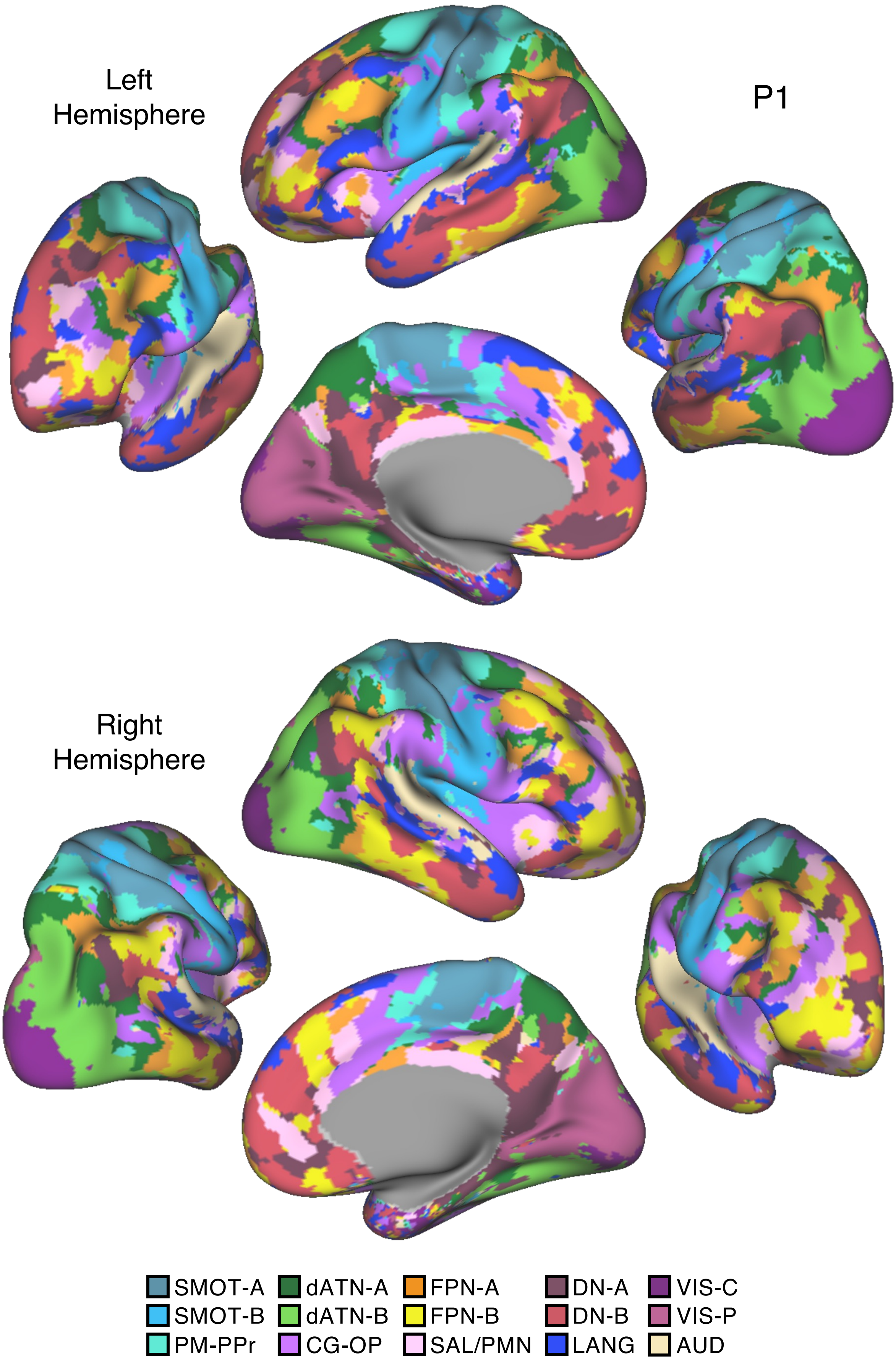

Left
Hemisphere
P1
Right
Hemisphere
SMOT-A
SMOT-B
PM-PPr
dATN-A
dATN-B
CG-OP
FPN-A
FPN-B
SAL/PMN
DN-A
DN-B
LANG
VIS-C
VIS-P
AUD


Figure S21

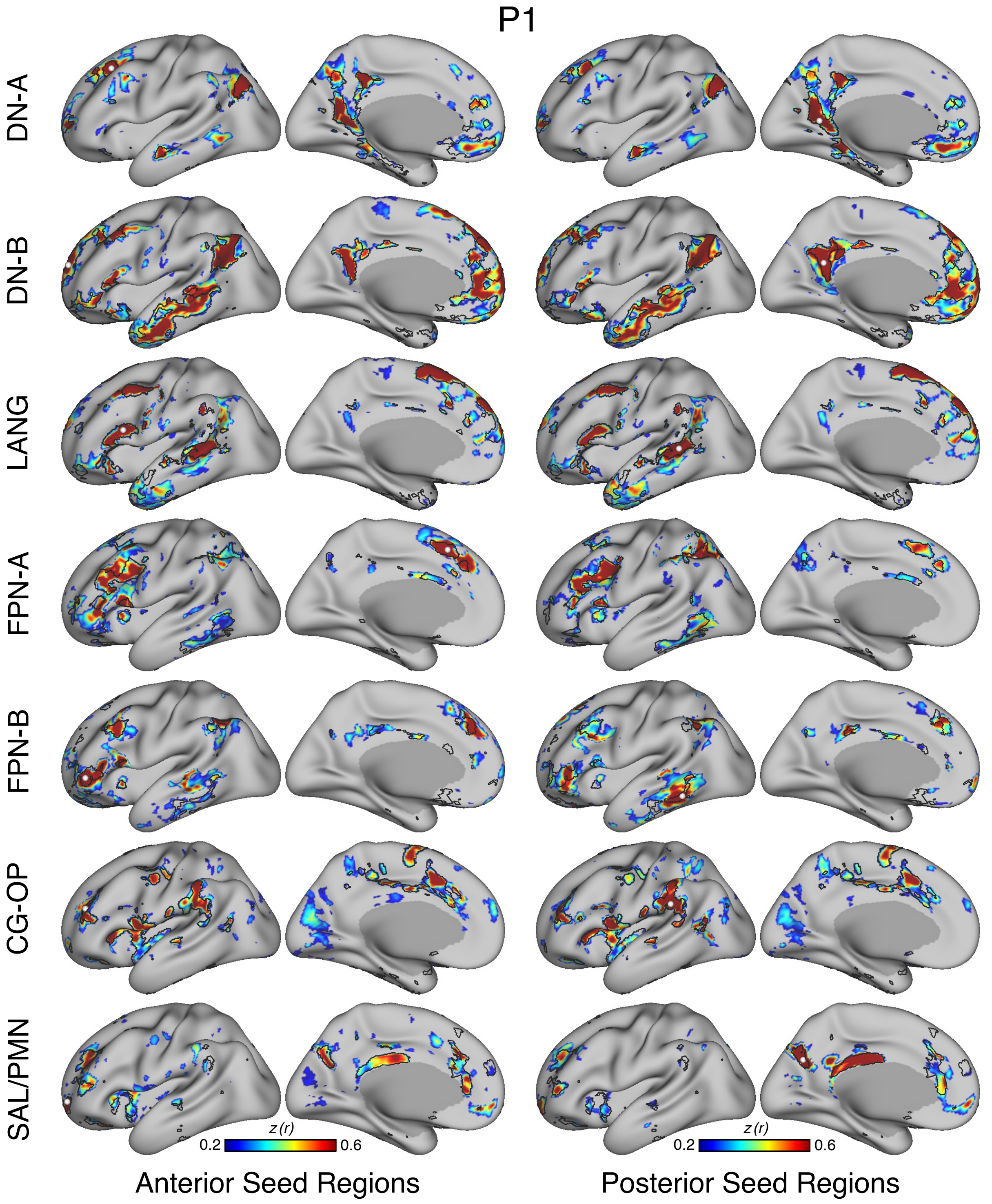
P1
DN-A
DN-B
LANG
FPN-A
FPN-B
CG-OP
SAL/PMN
z (r)
0.2
0.6
z (r)
0.2
0.6
Anterior Seed Regions
Posterior Seed Regions

Figure S22

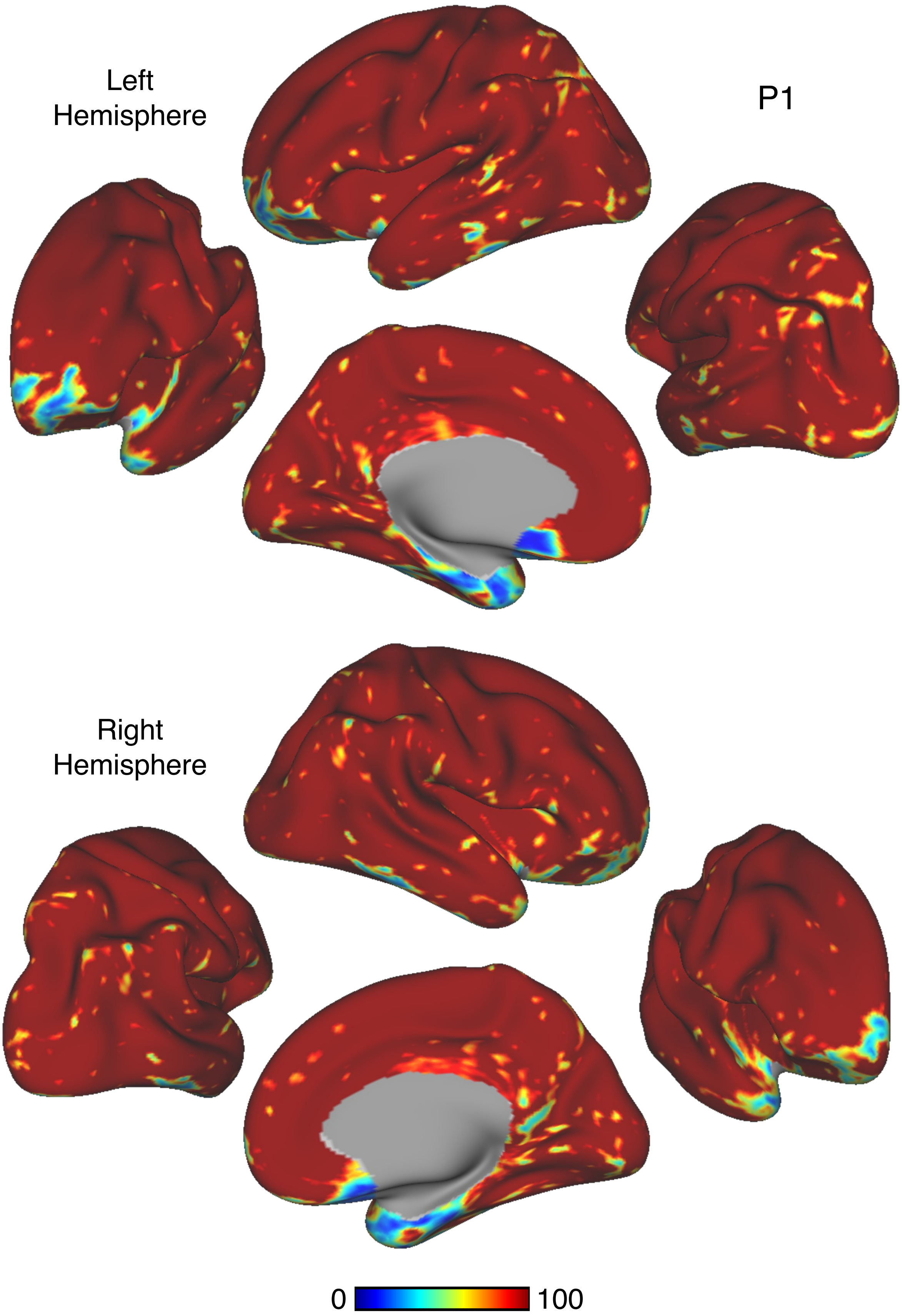


Figure S23

P1

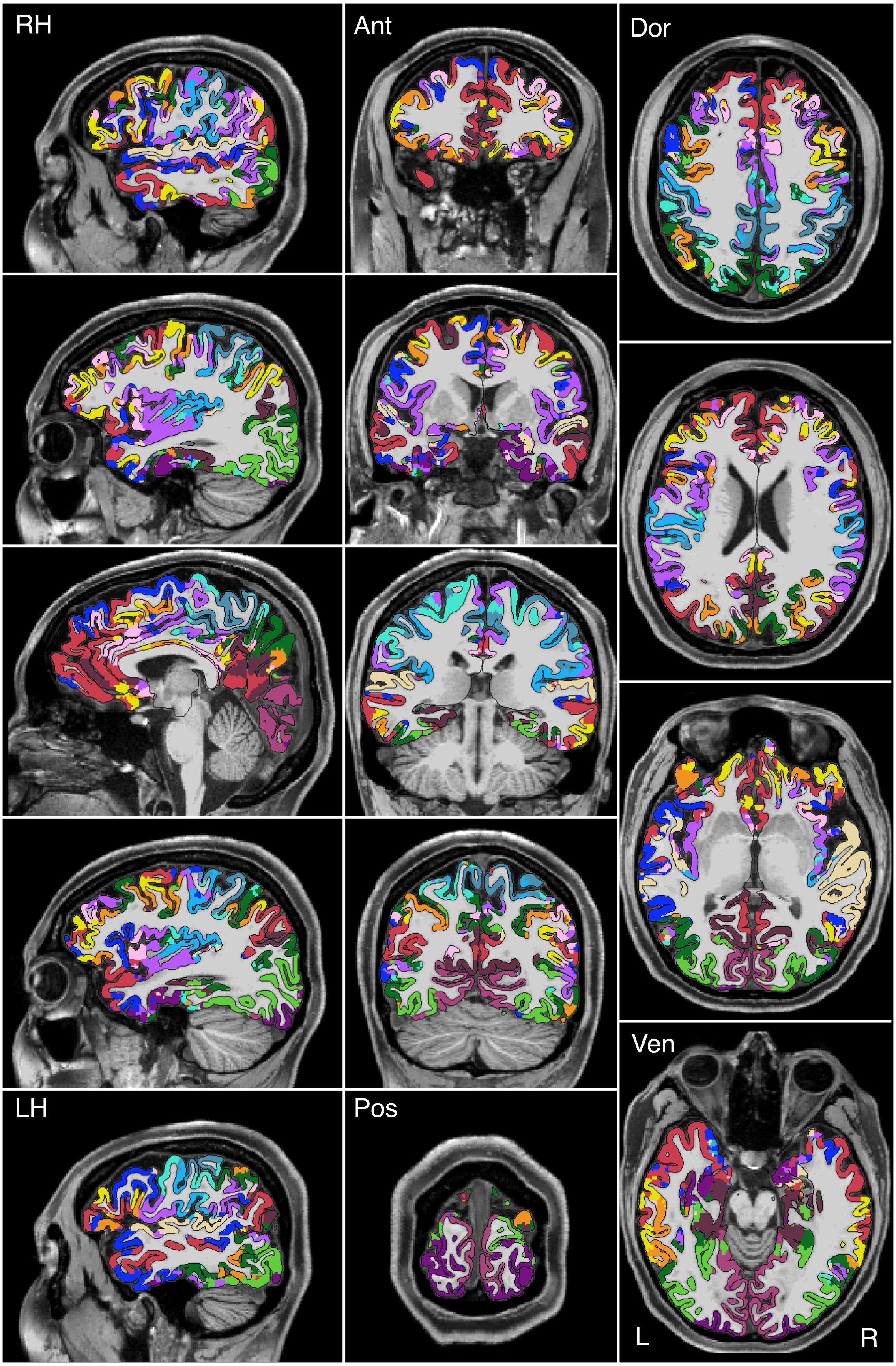


Figure S24

# P1

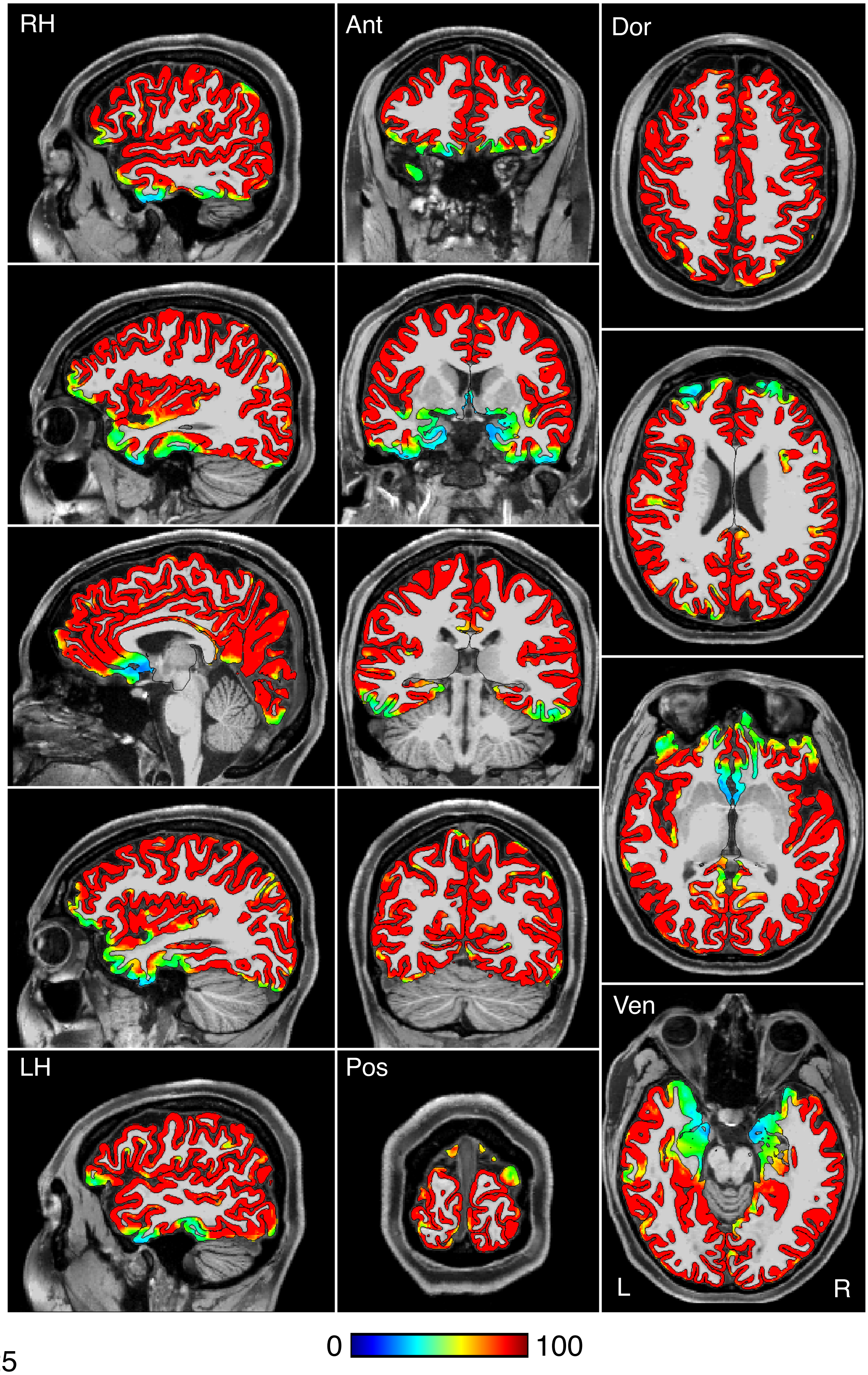


Figure S25

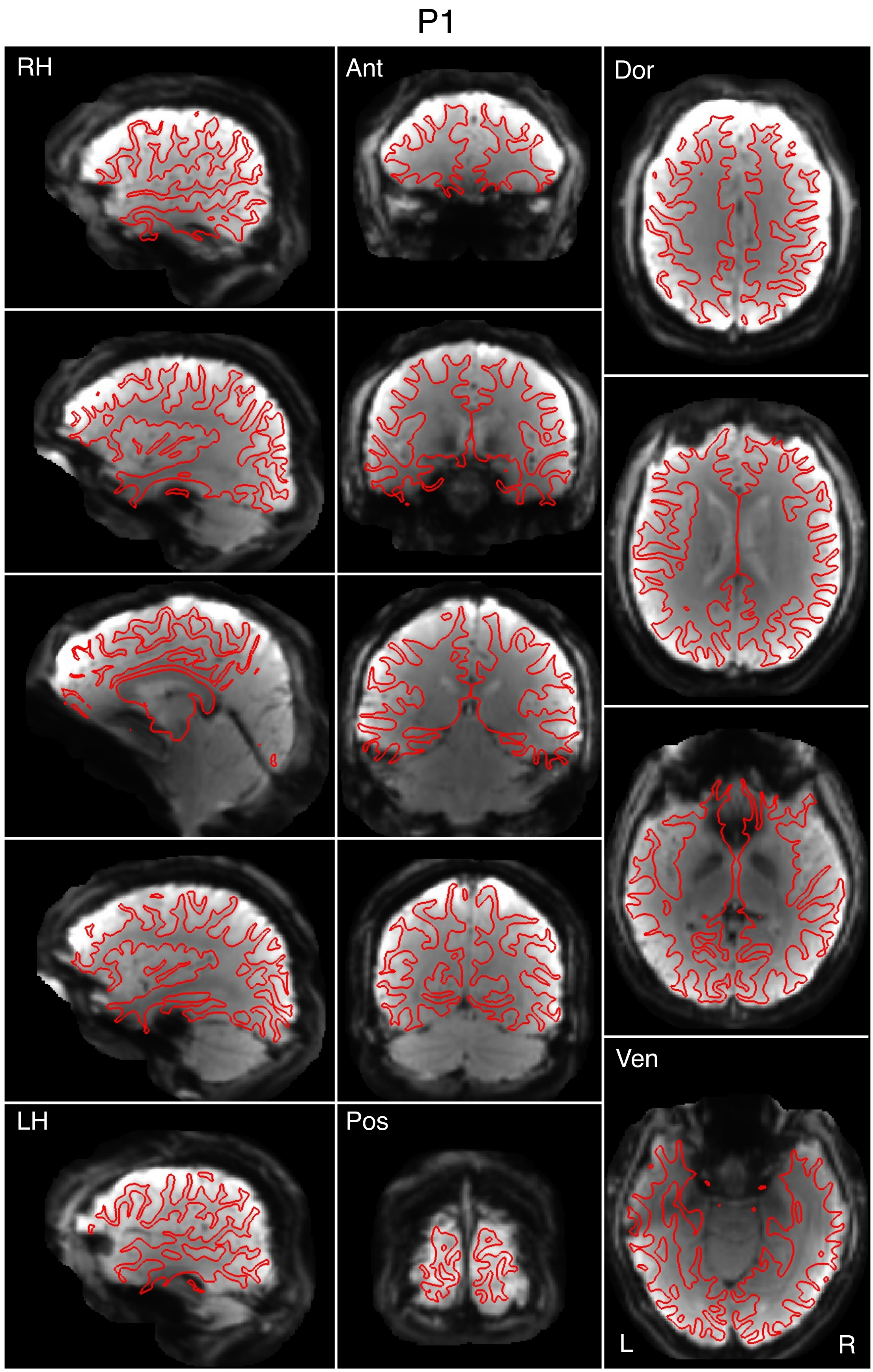


Figure S26

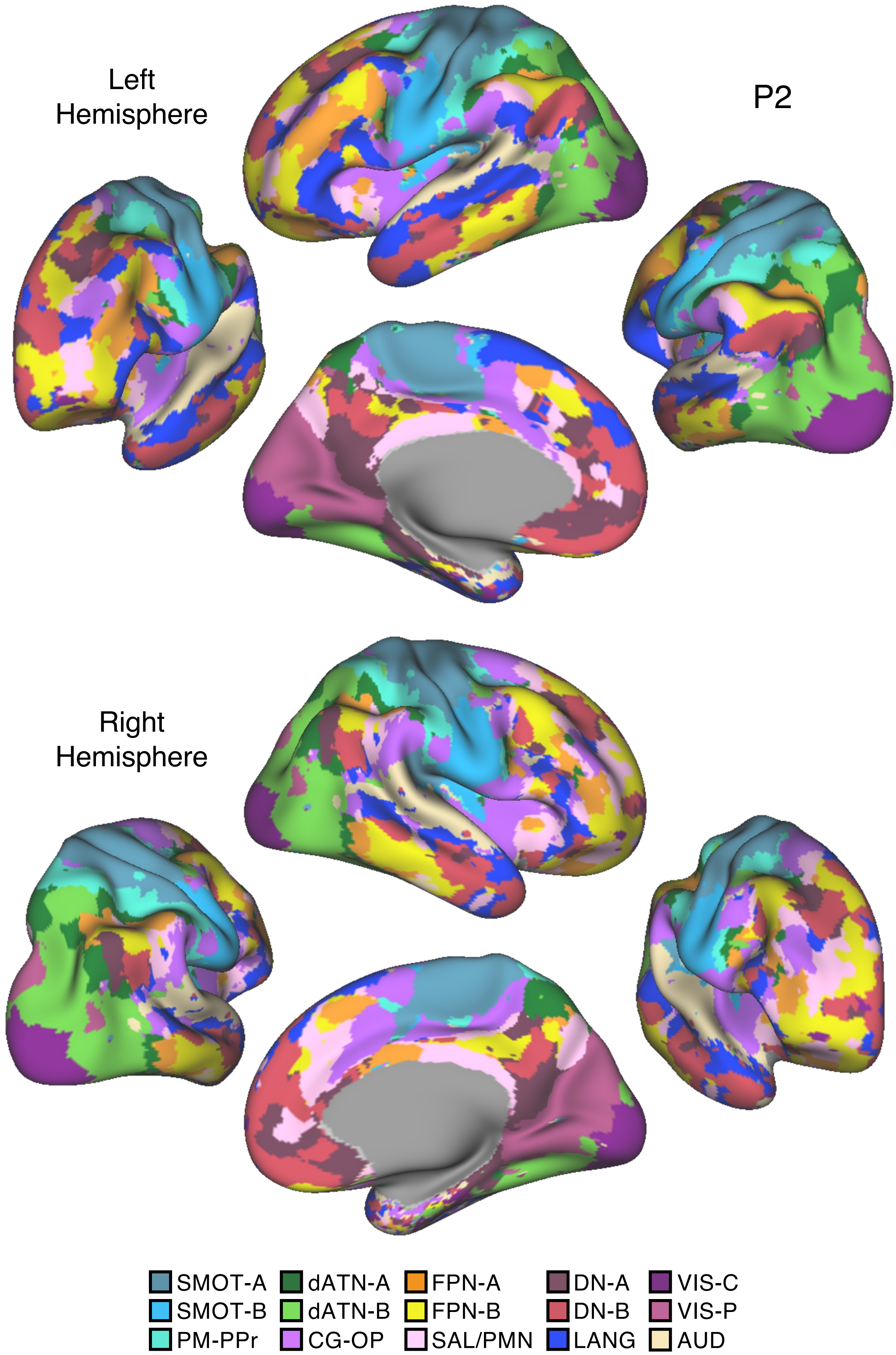
Left
Hemisphere
P2
Right
Hemisphere
SMOT-A
SMOT-B
PM-PPr
dATN-A
dATN-B
CG-OP
FPN-A
FPN-B
SAL/PMN
DN-A
DN-B
LANG
VIS-C
VIS-P
AUD

Figure S27

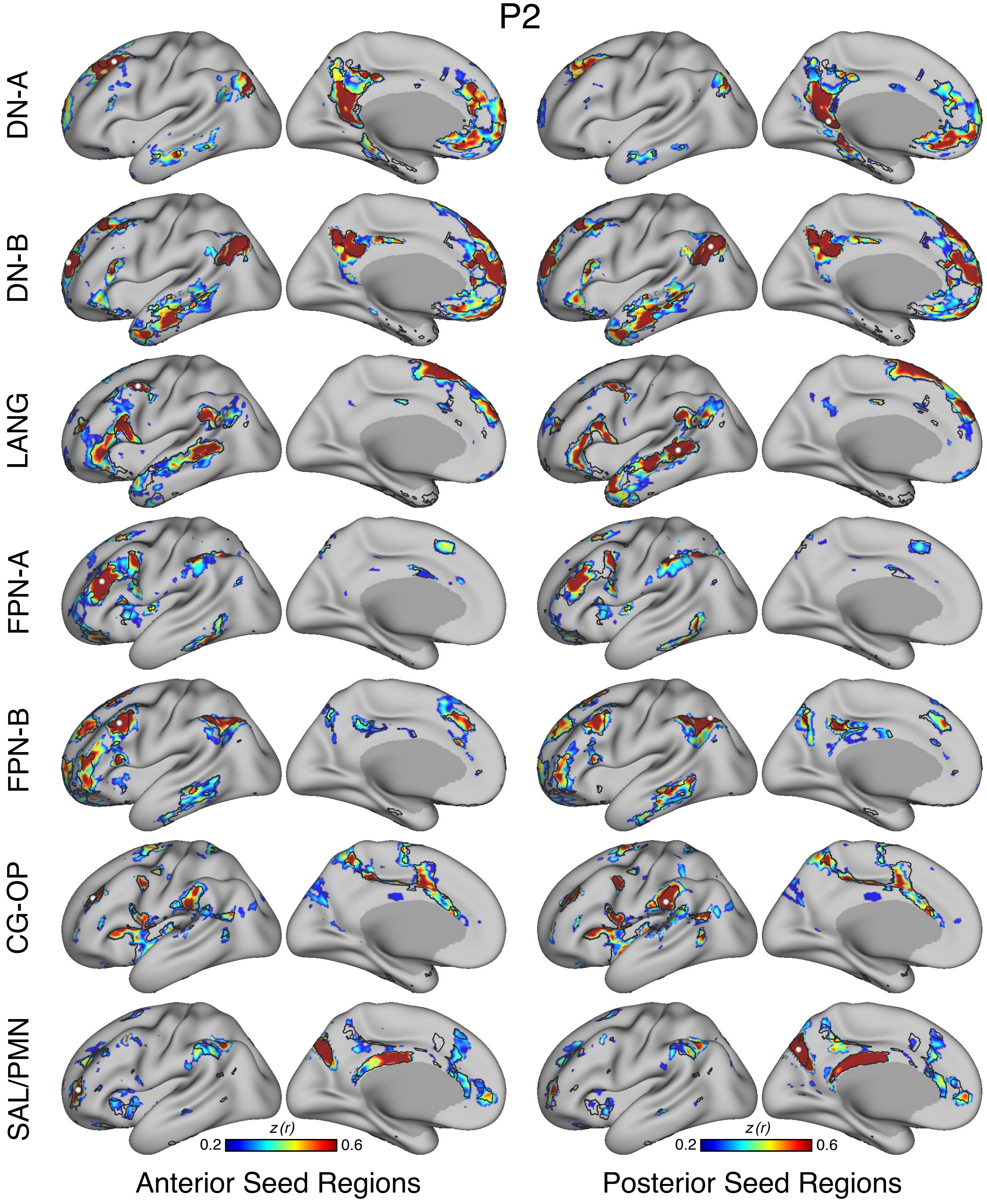
P2
DN-A
DN-B
LANG
FPN-A
FPN-B
CG-OP
SAL/PMN
z (r)
0.2
0.6
z (r)
0.2
0.6
Anterior Seed Regions
Posterior Seed Regions

Figure S28

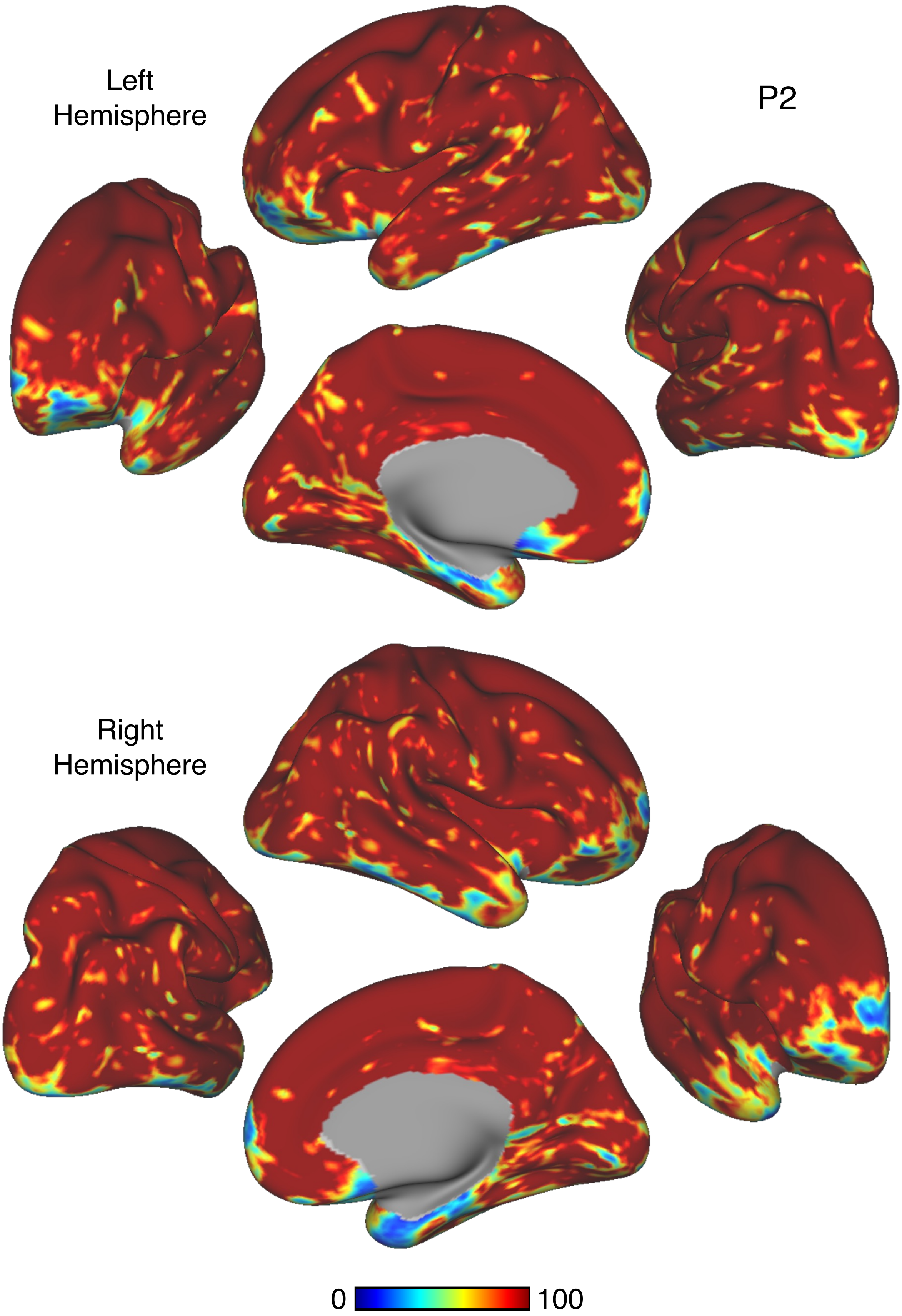


Figure S29

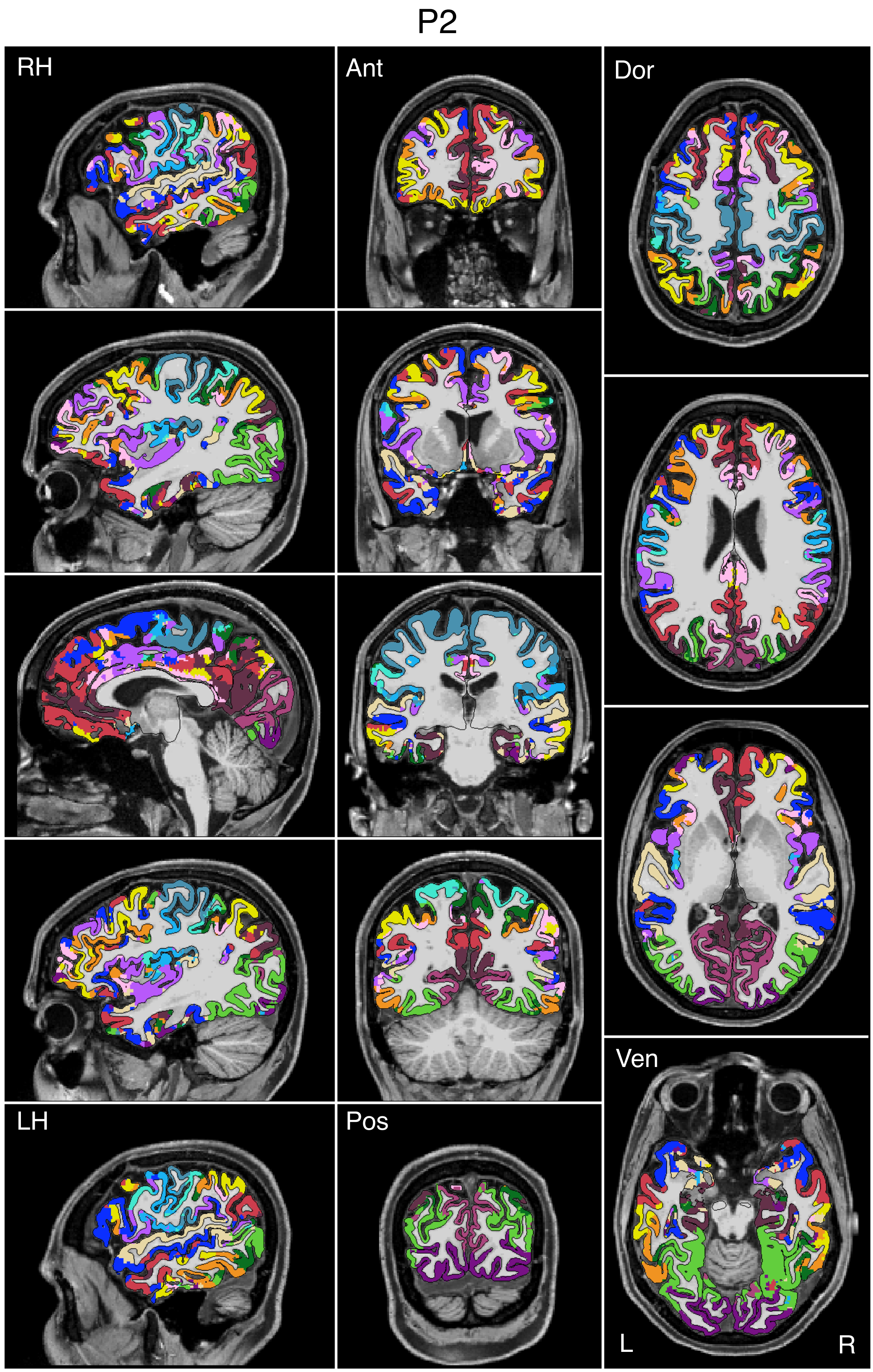


Figure S30

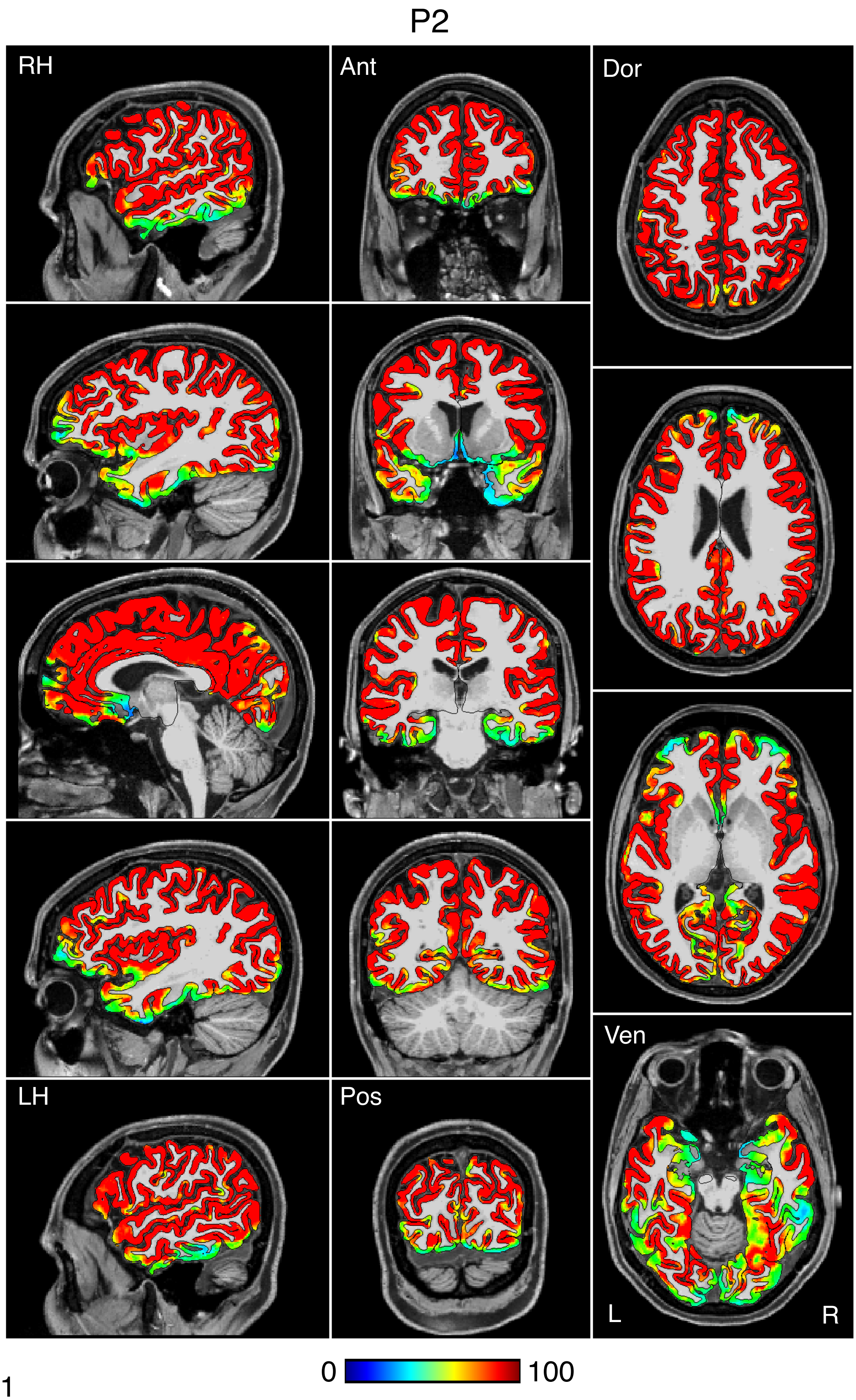


Figure S31

P2

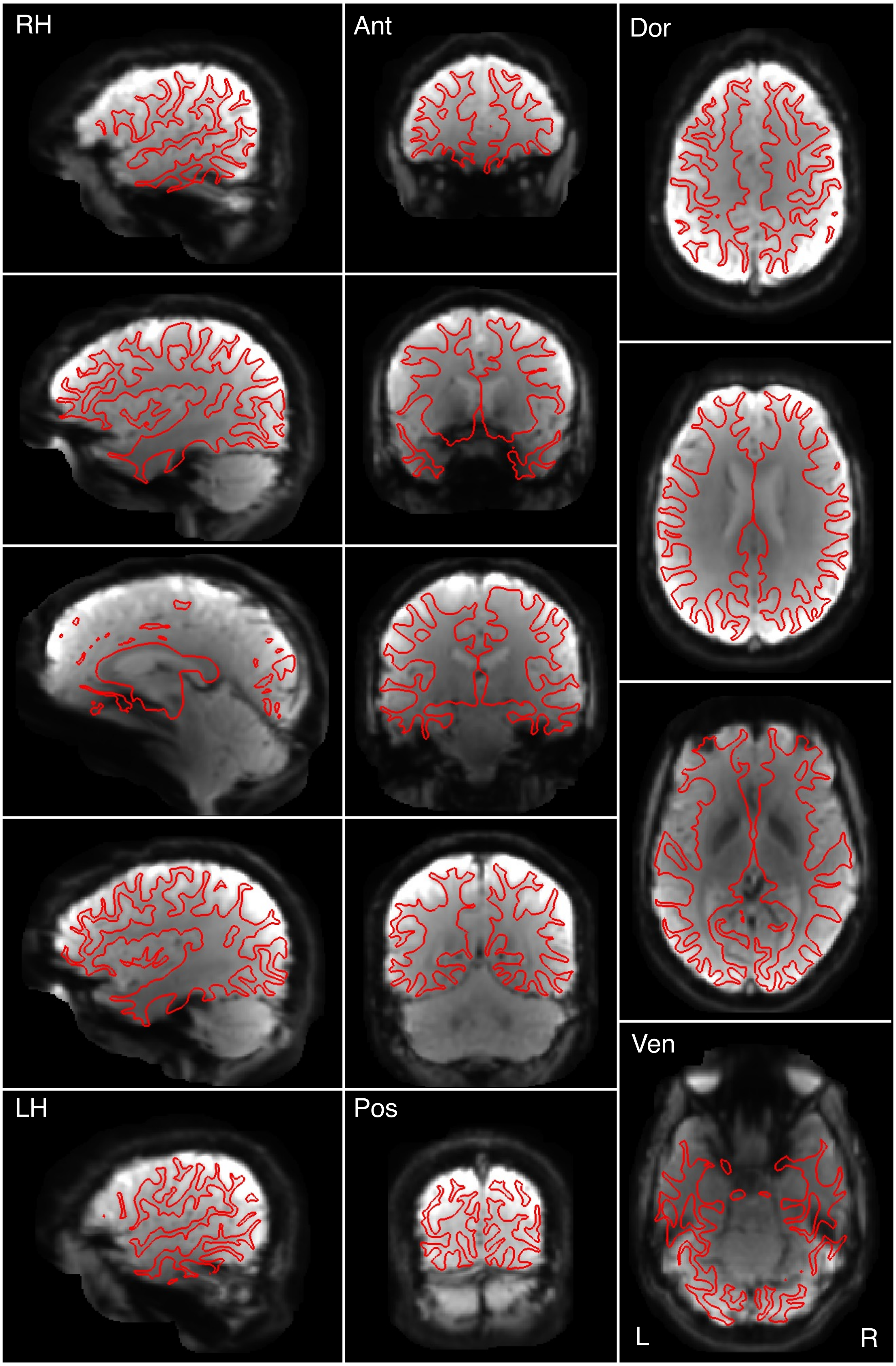


Figure S32

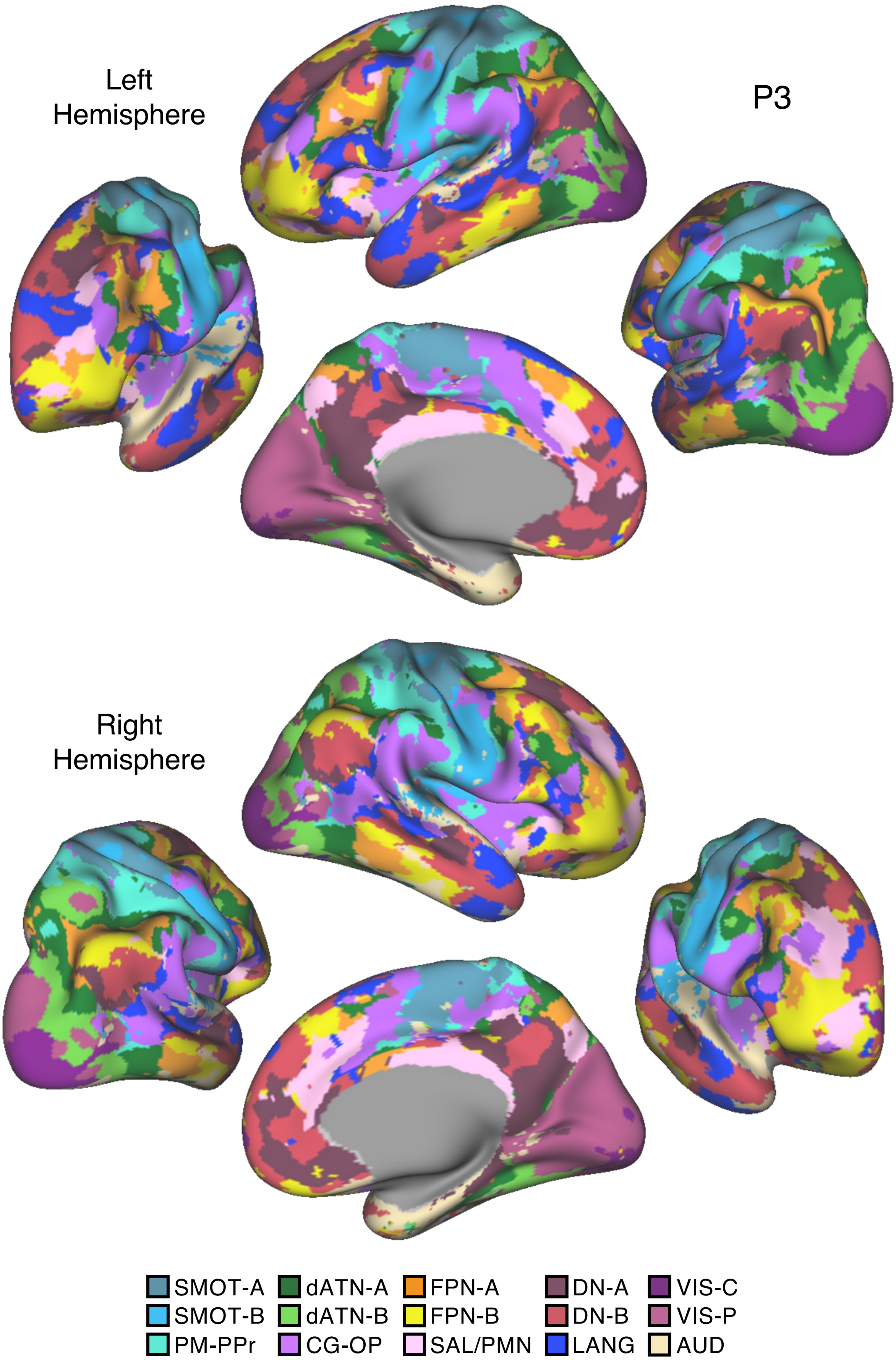
Left
Hemisphere
P3
Right
Hemisphere
SMOT-A
SMOT-B
PM-PPr
dATN-A
dATN-B
CG-OP
FPN-A
FPN-B
SAL/PMN
DN-A
DN-B
LANG
VIS-C
VIS-P
AUD

Figure S33

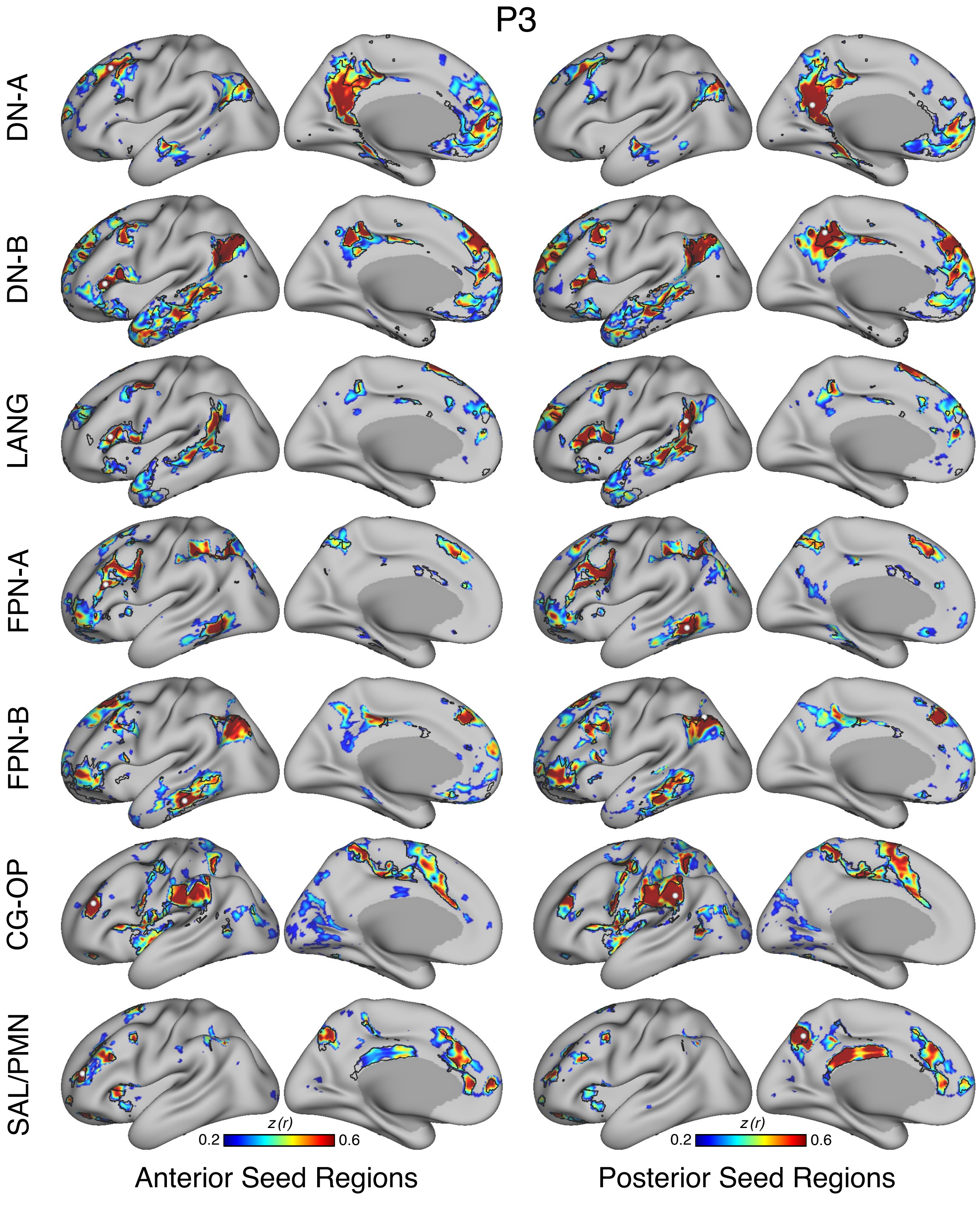
P3
DN-A
DN-B
LANG
FPN-A
FPN-B
CG-OP
SAL/PMN
z (r)
0.2
0.6
z (r)
0.2
0.6
Anterior Seed Regions
Posterior Seed Regions

Figure S34

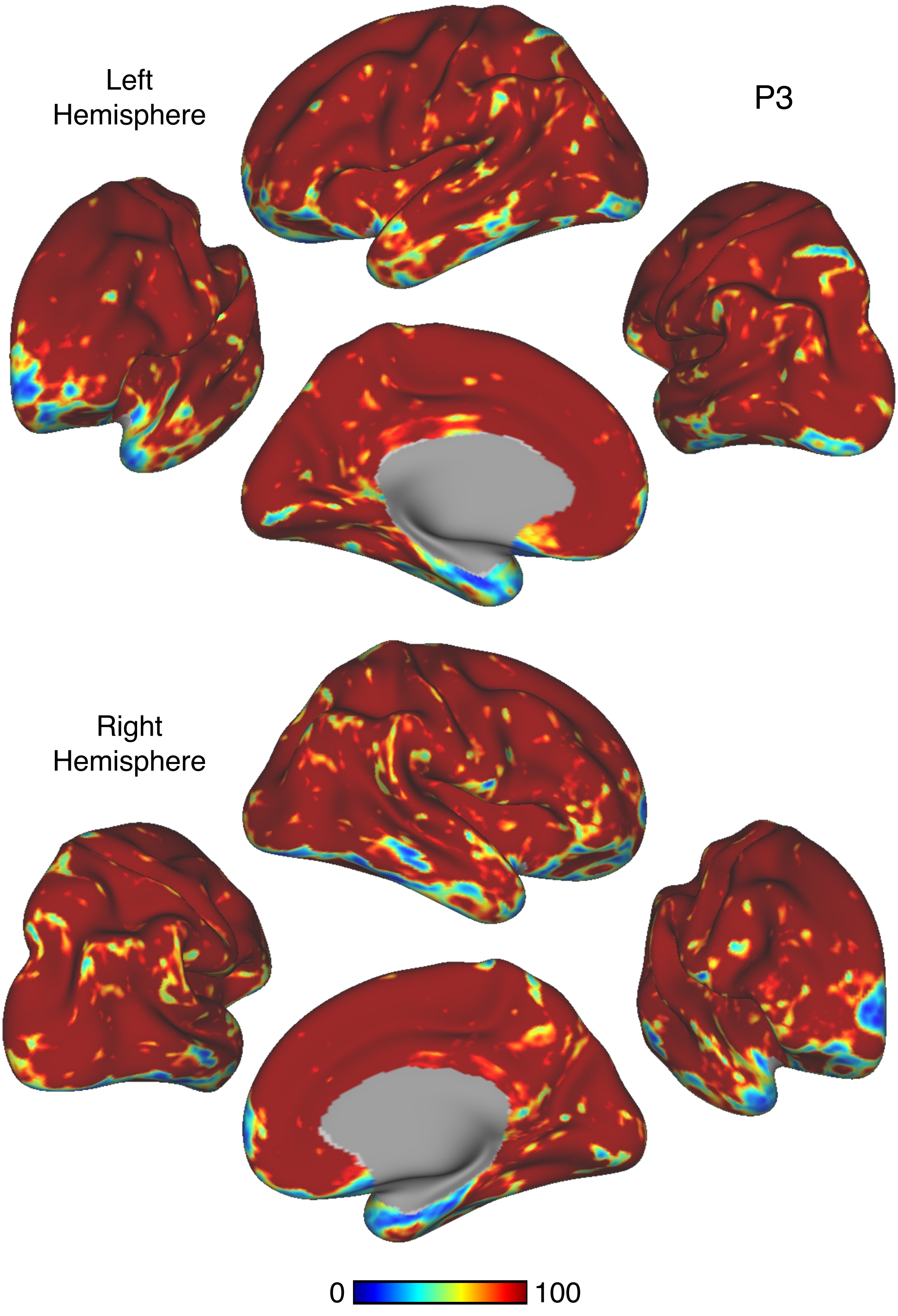


Figure S35

P3

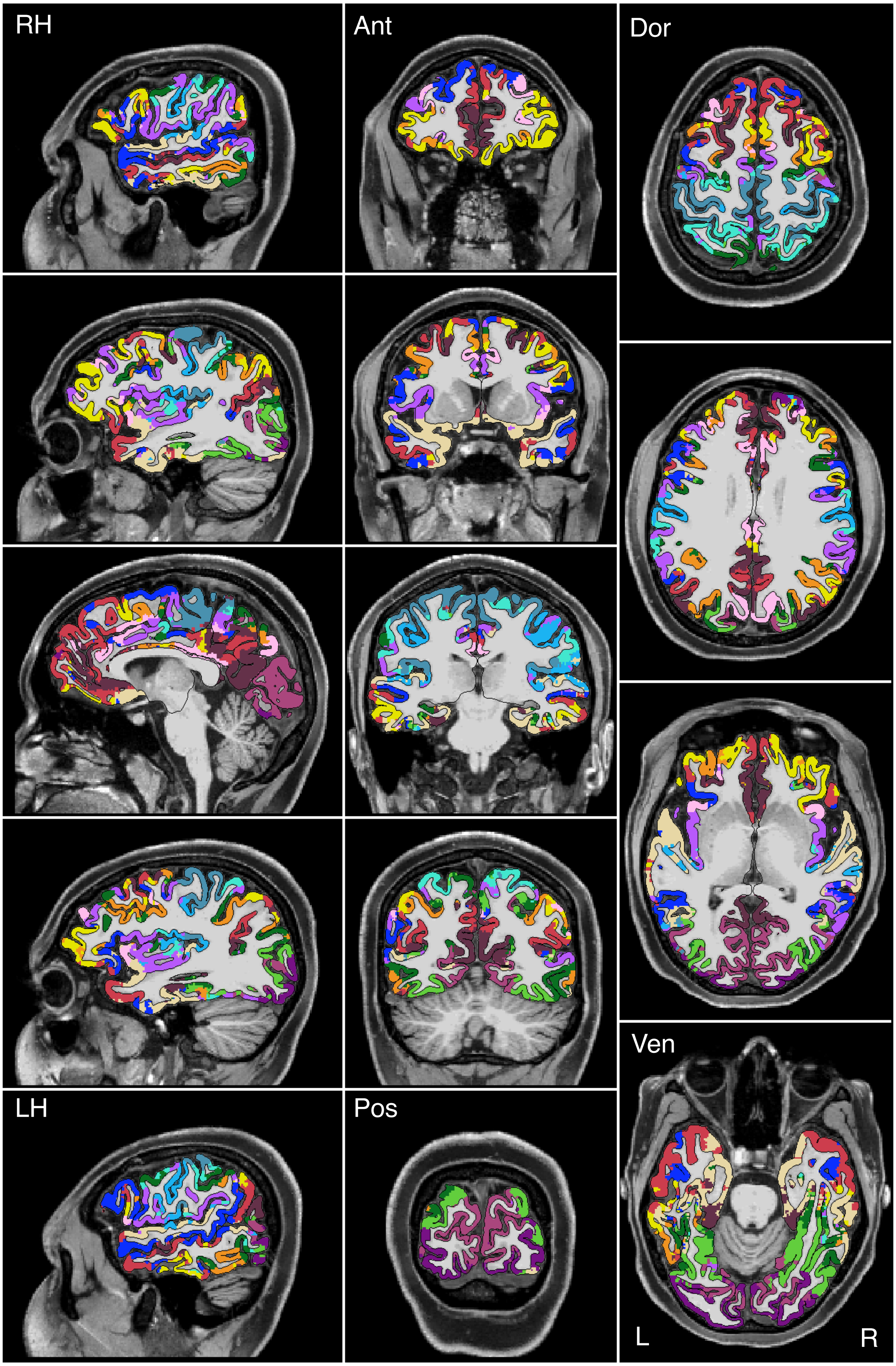


Figure S36

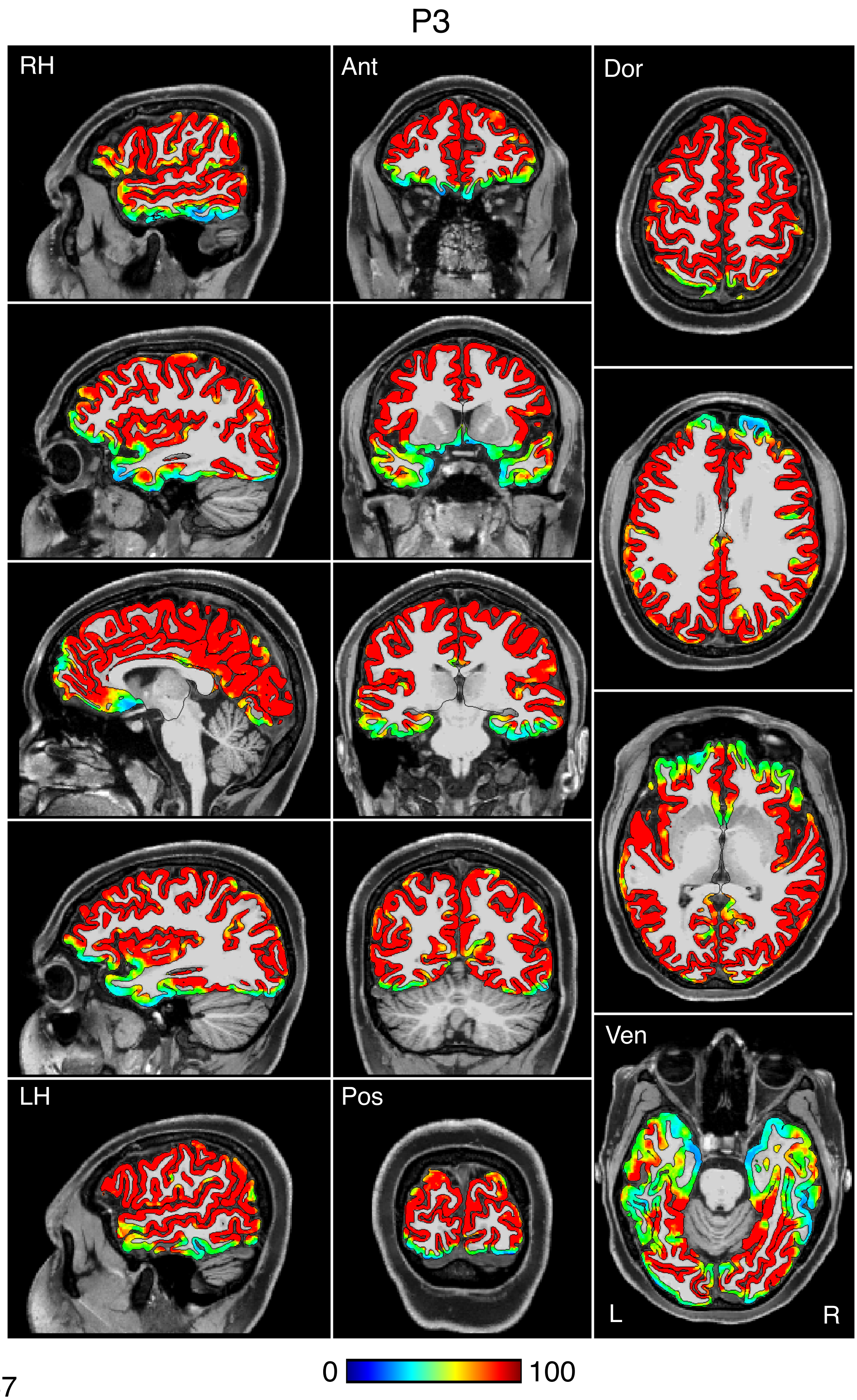


Figure S37

P3

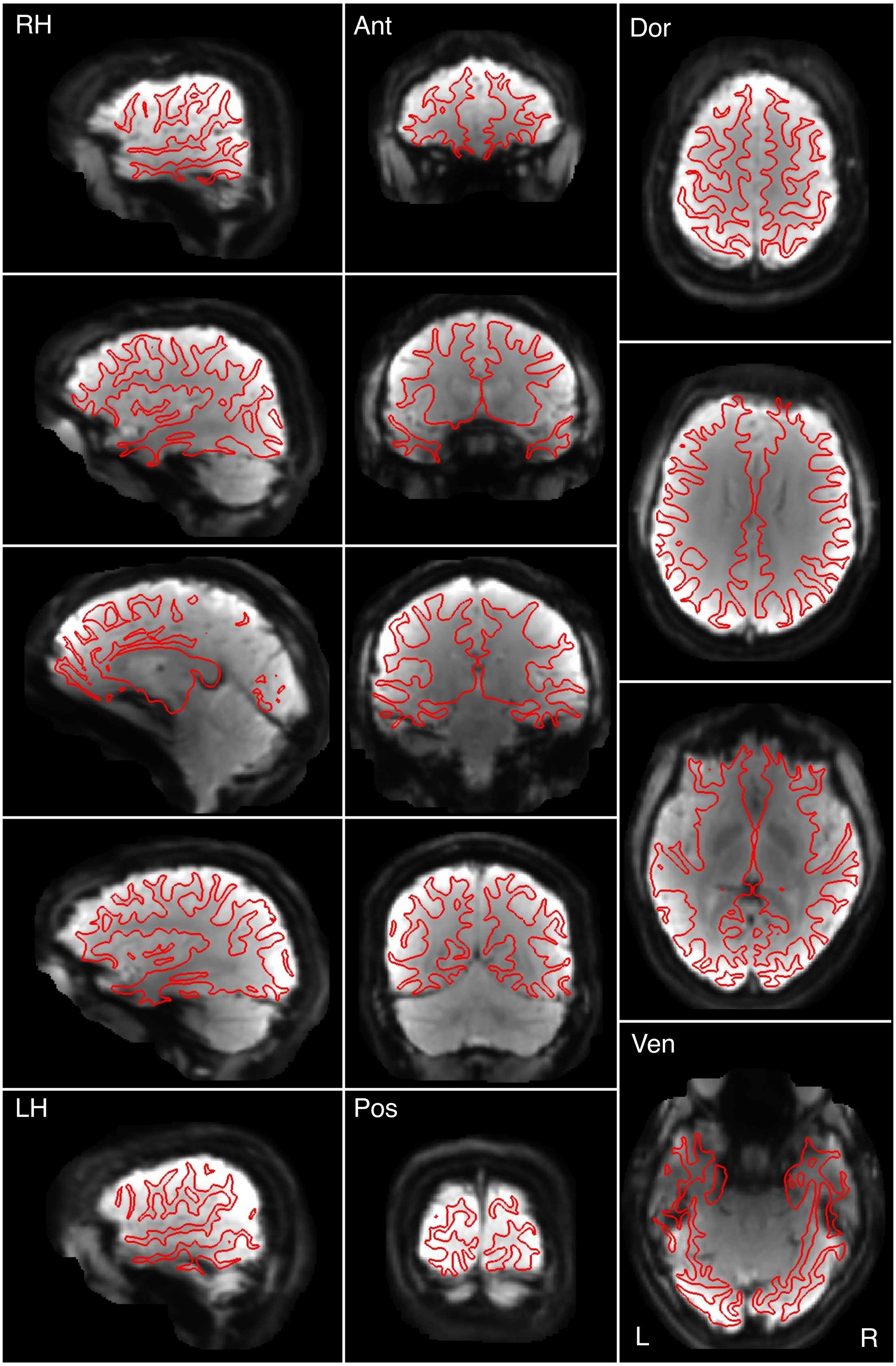


Figure S38

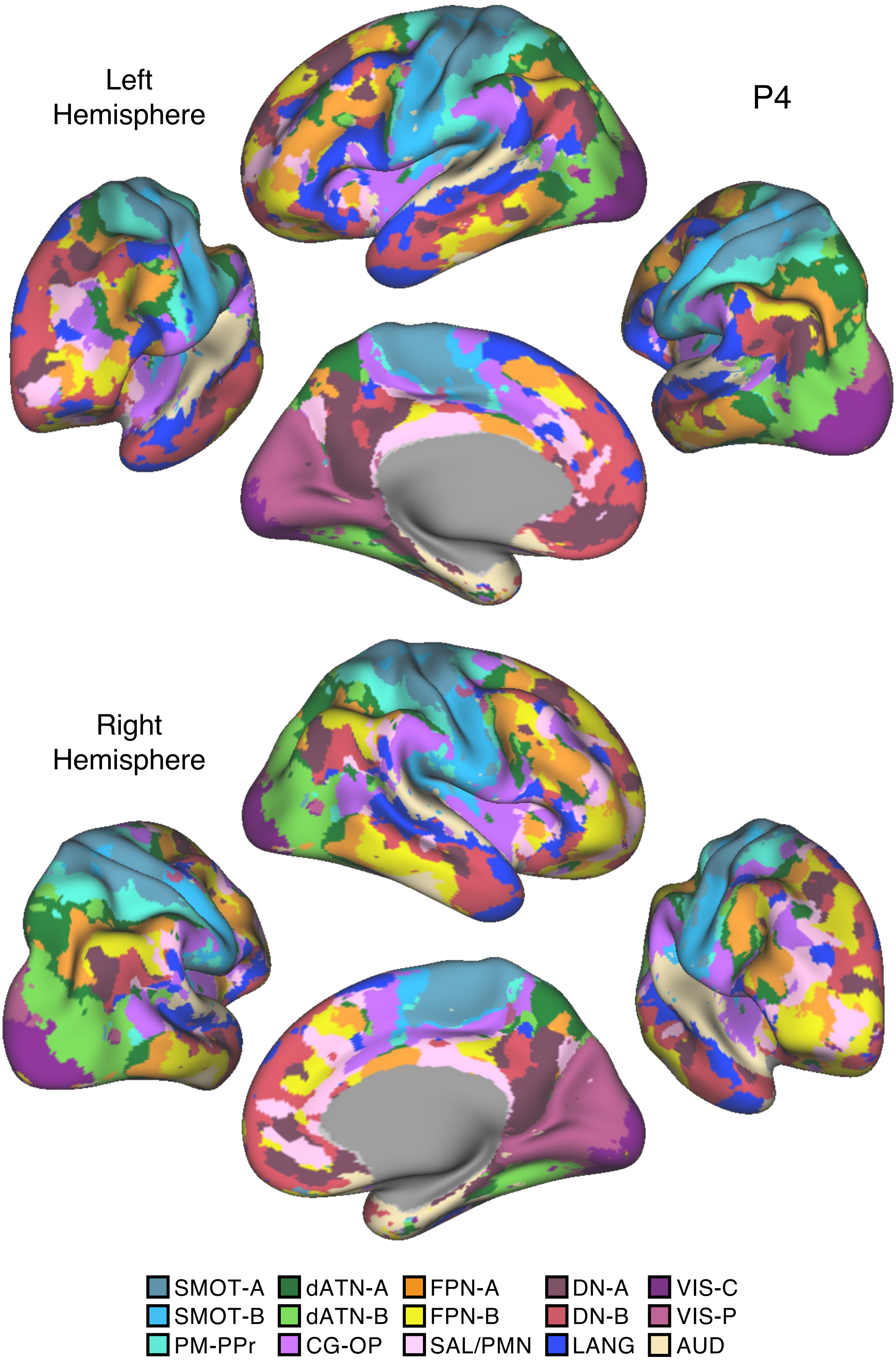
Left
Hemisphere
P4
Right
Hemisphere
SMOT-A
SMOT-B
PM-PPr
dATN-A
dATN-B
CG-OP
FPN-A
FPN-B
SAL/PMN
DN-A
DN-B
LANG
VIS-C
VIS-P
AUD

Figure S39

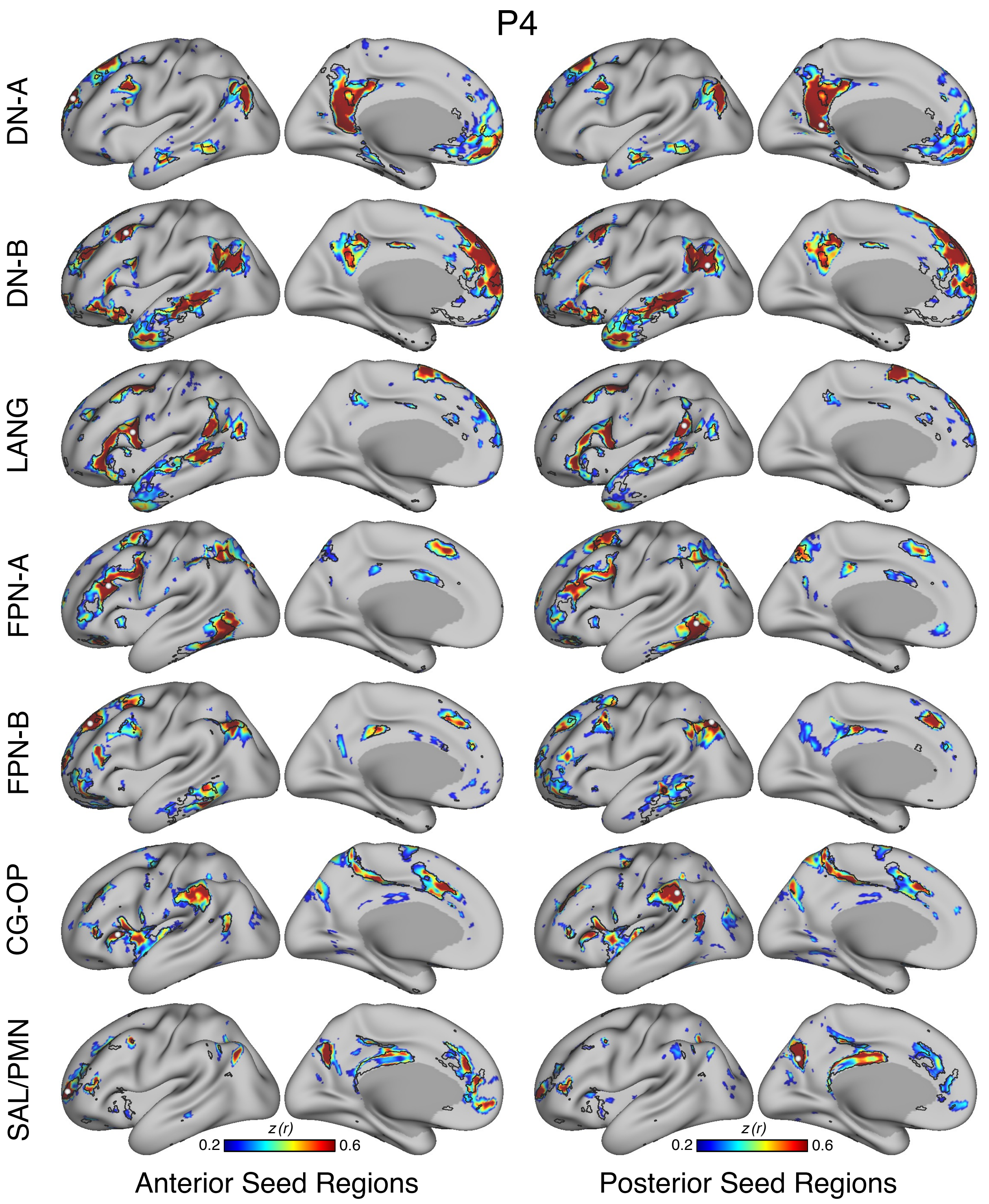
P4
DN-A
DN-B
LANG
FPN-A
FPN-B
CG-OP
SAL/PMN
z (r)
0.2
0.6
z (r)
0.2
0.6
Anterior Seed Regions
Posterior Seed Regions

Figure S40

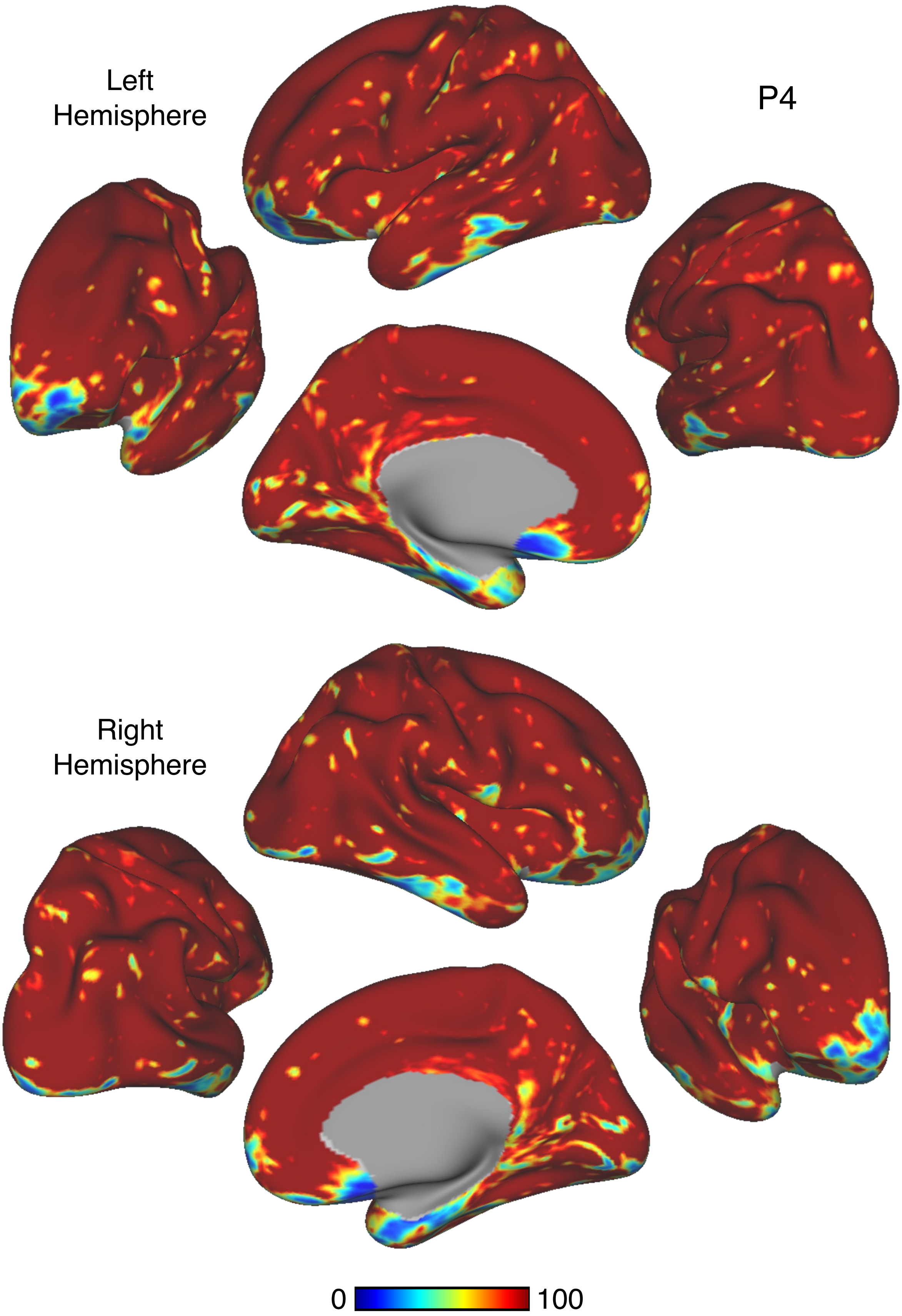


Figure S41

# P4

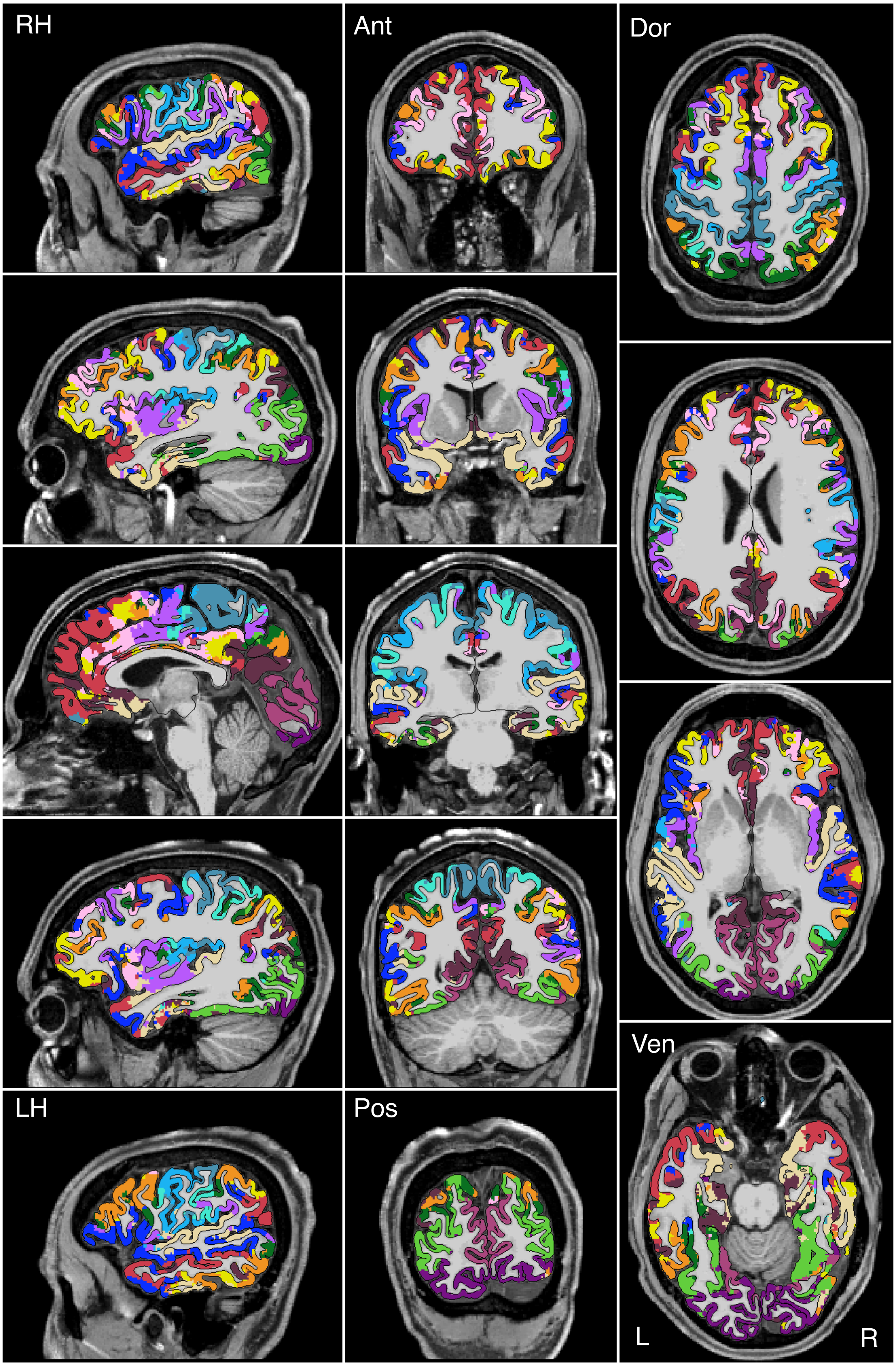


Figure S42

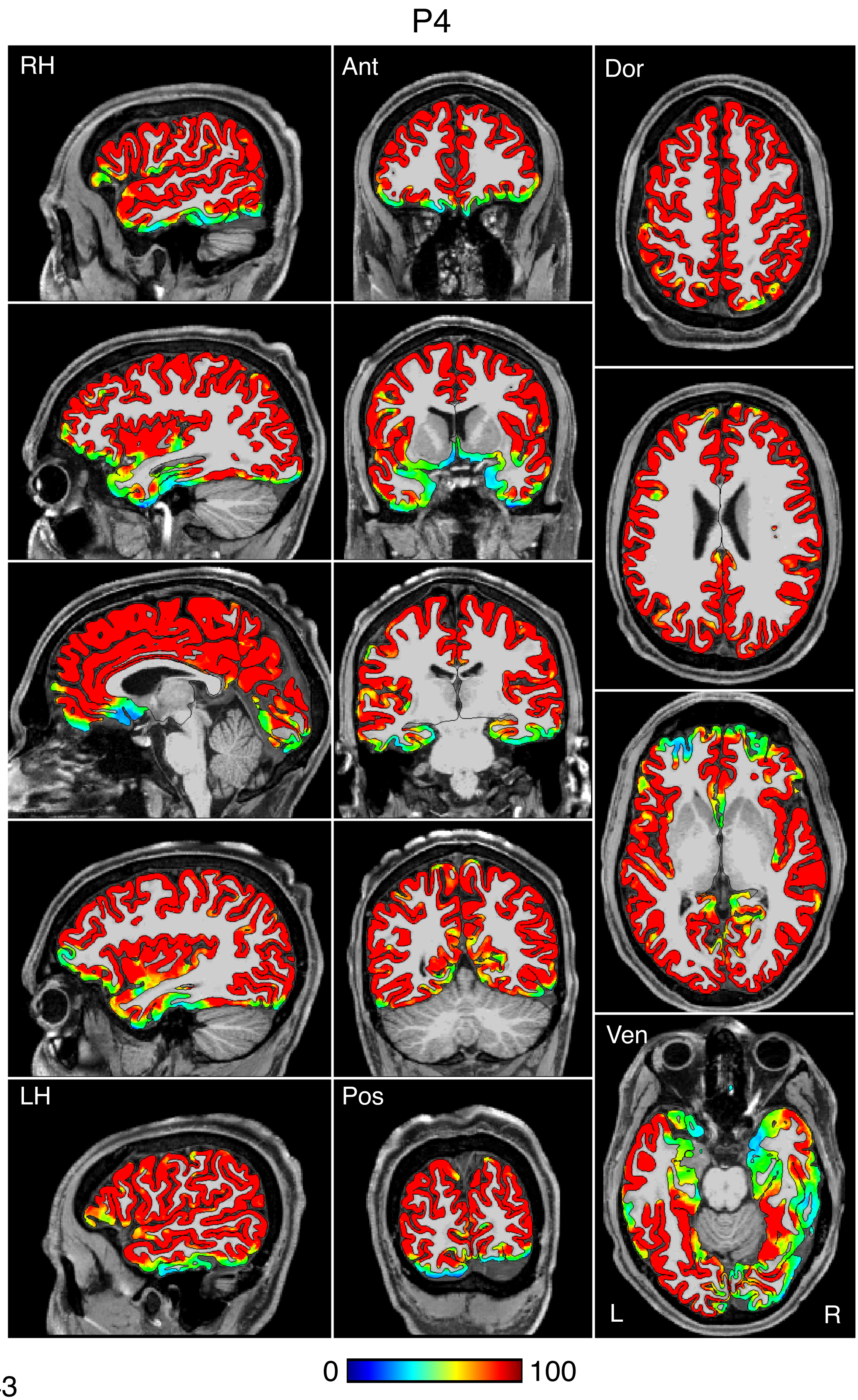


Figure S43

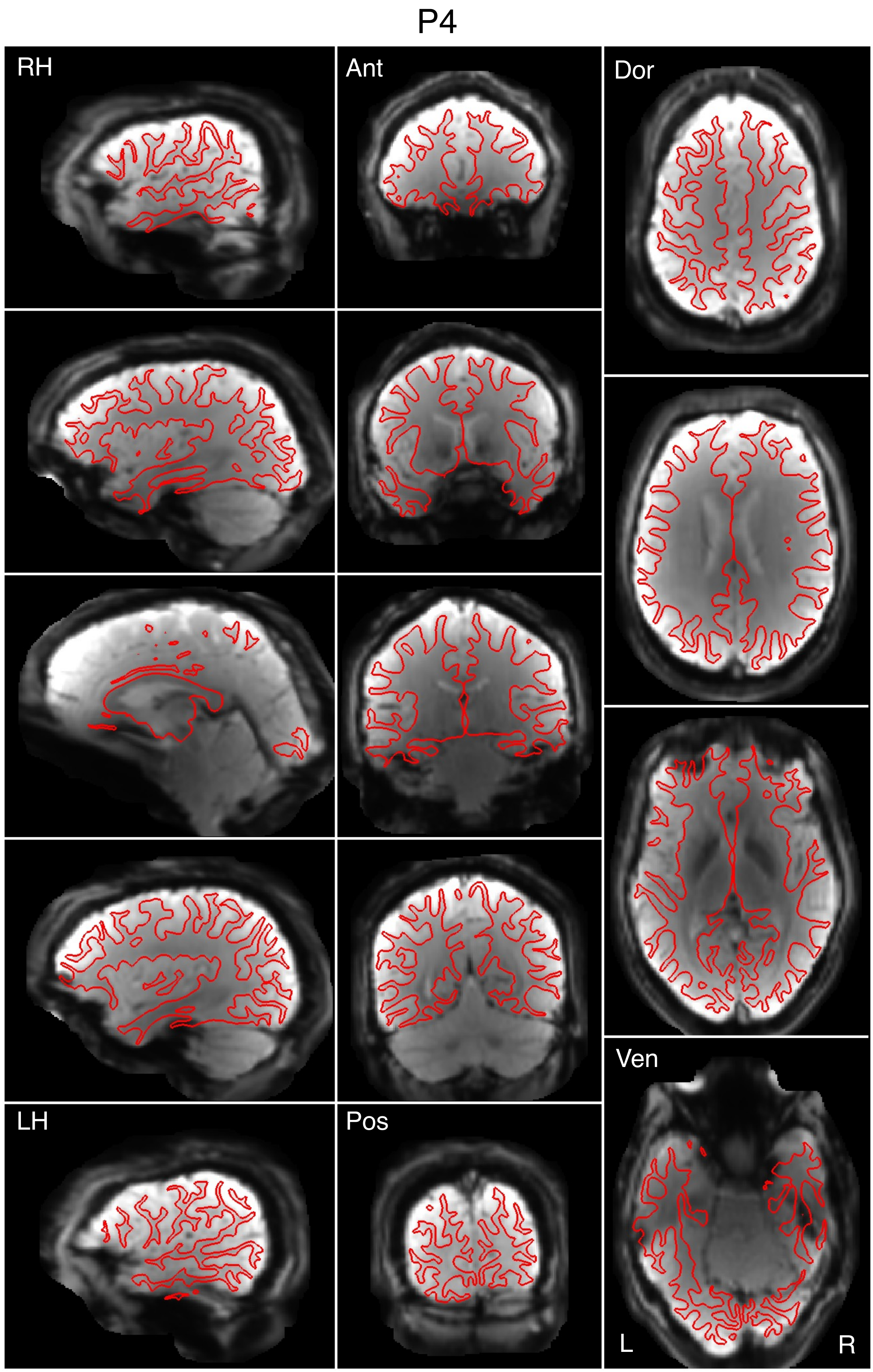


Figure S44

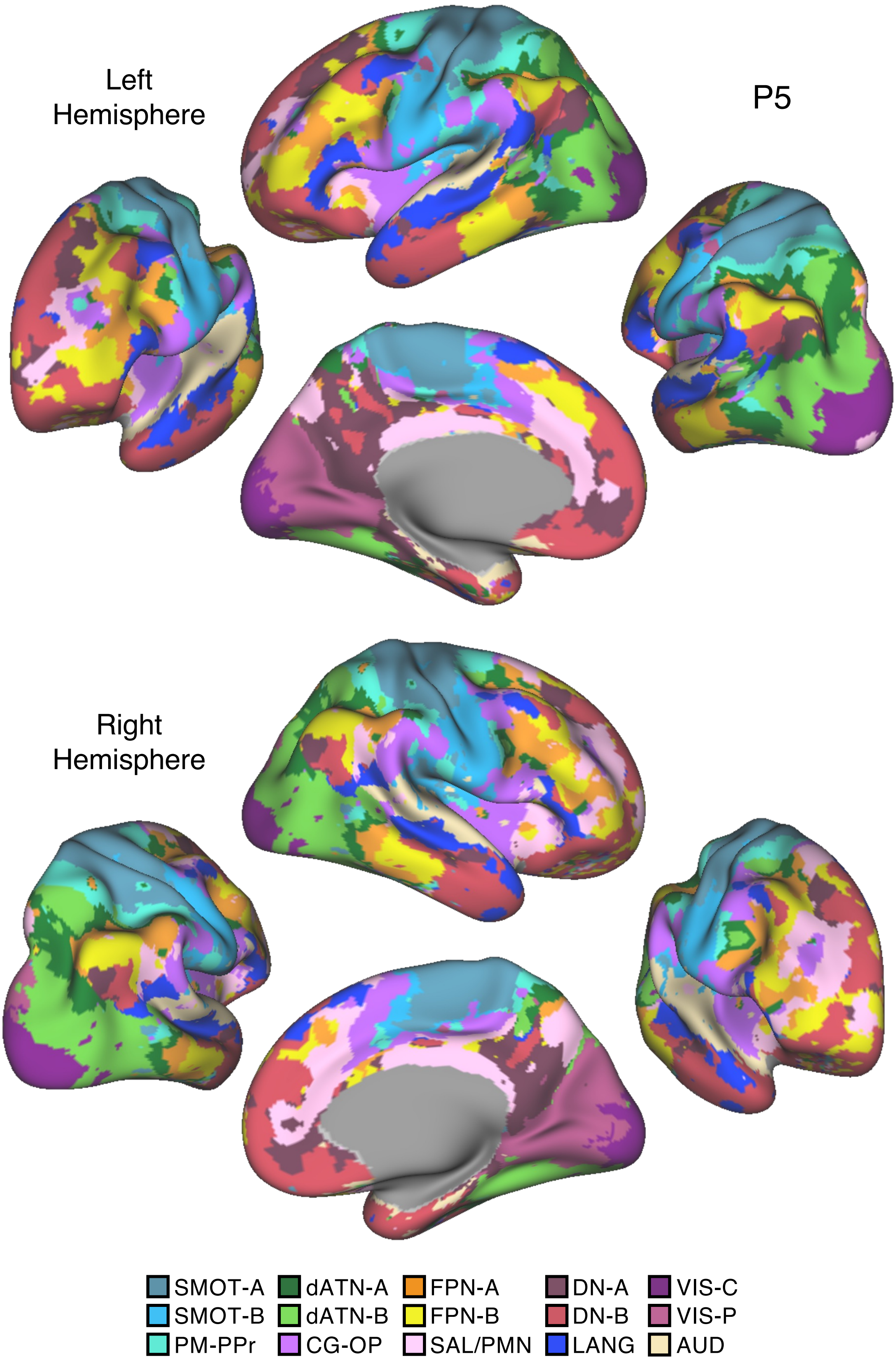
Left
Hemisphere
P5
Right
Hemisphere
SMOT-A
SMOT-B
PM-PPr
dATN-A
dATN-B
CG-OP
FPN-A
FPN-B
SAL/PMN
DN-A
DN-B
LANG
VIS-C
VIS-P
AUD

Figure S45

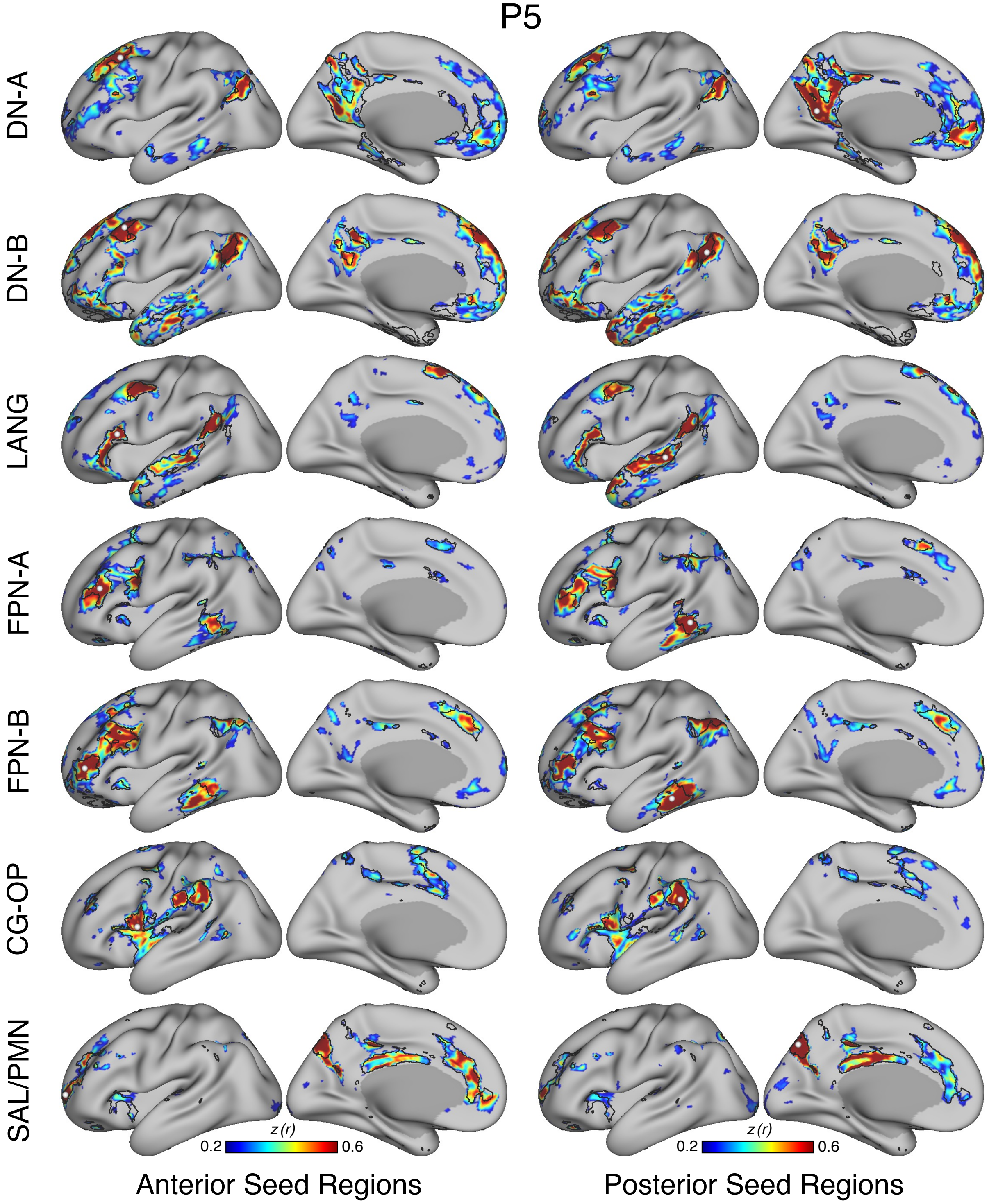
P5
DN-A
DN-B
LANG
FPN-A
FPN-B
CG-OP
SAL/PMN
z (r)
0.2
0.6
z (r)
0.2
0.6
Anterior Seed Regions
Posterior Seed Regions

Figure S46

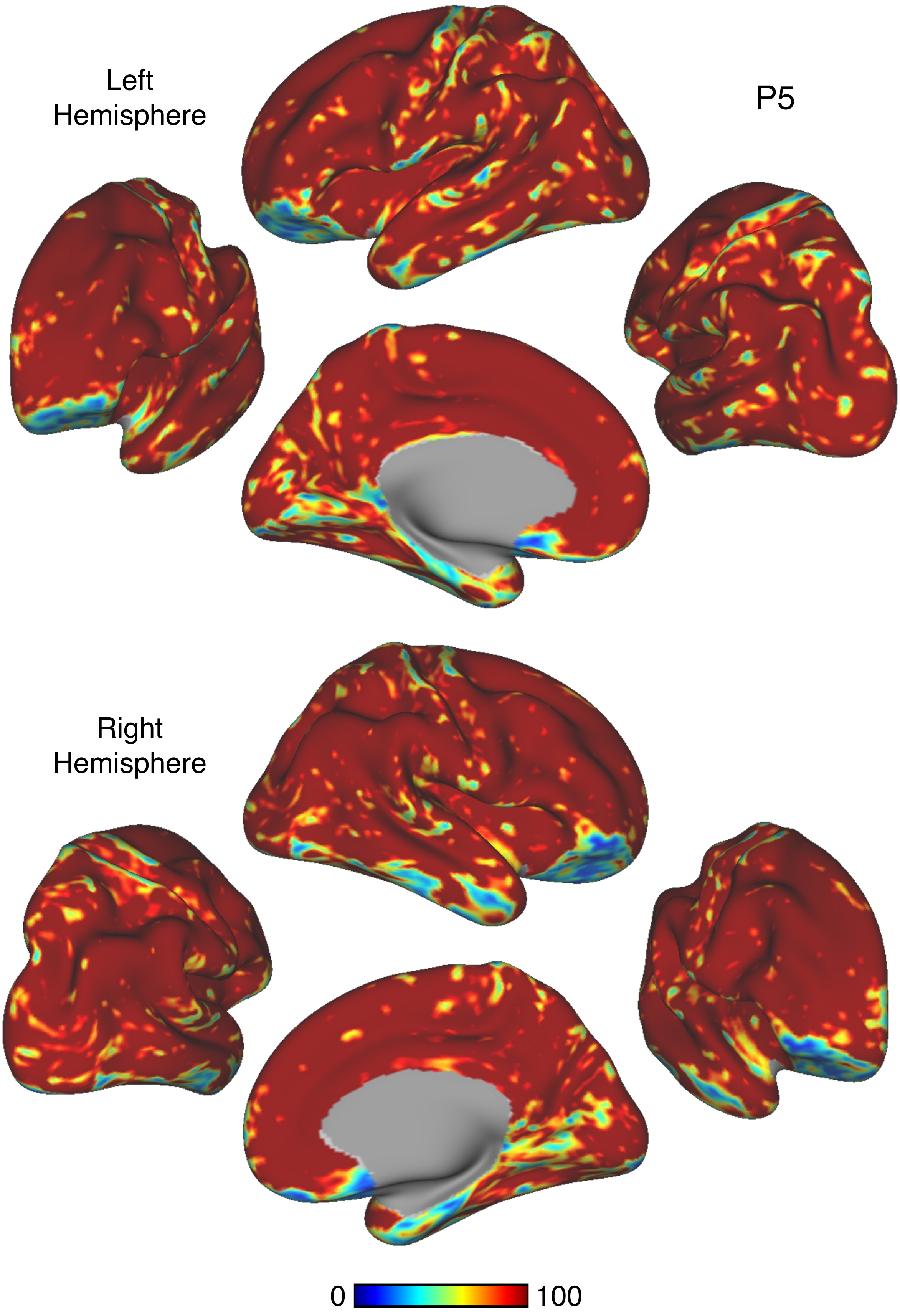


Figure S47

P5

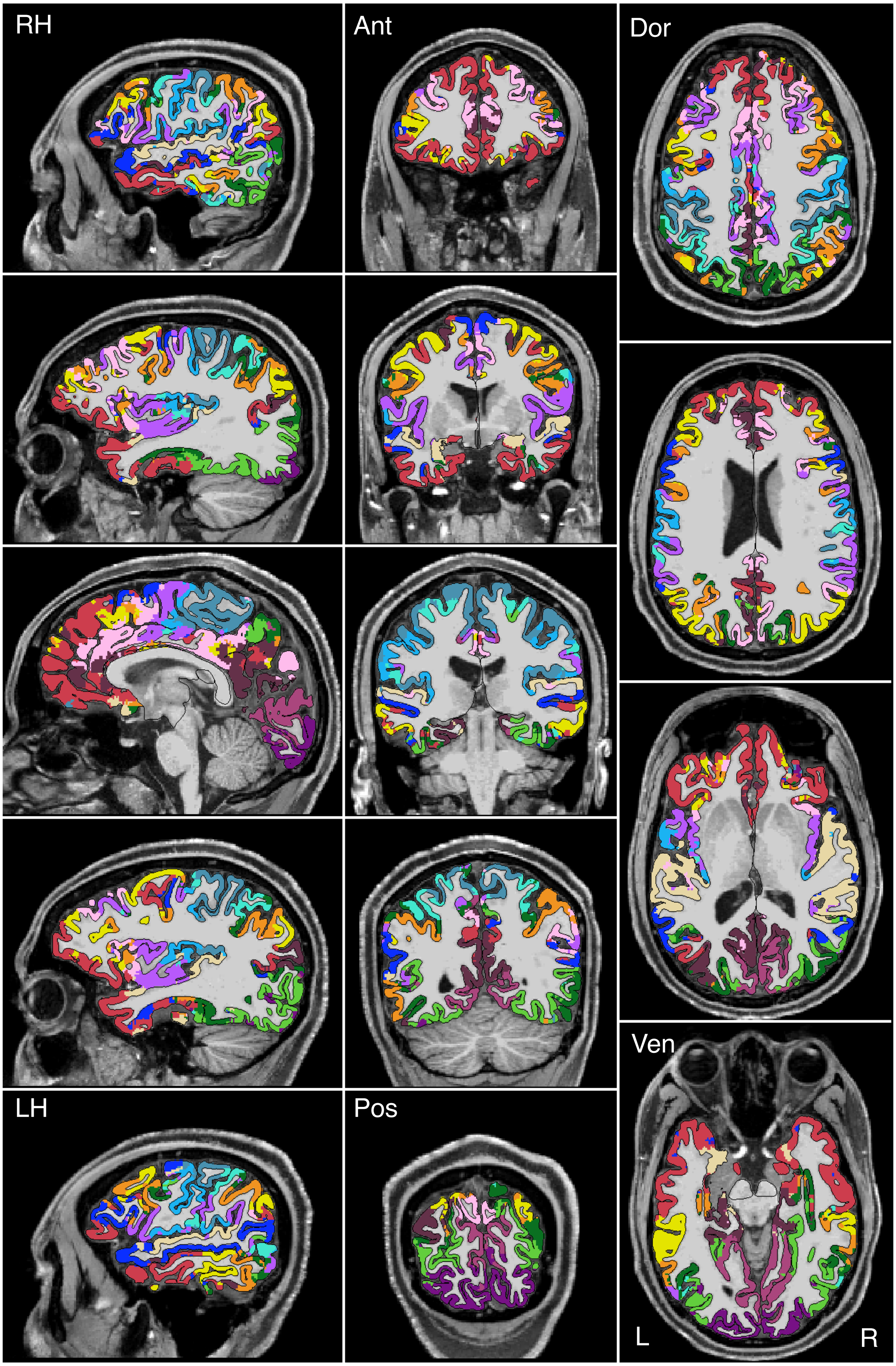


Figure S48

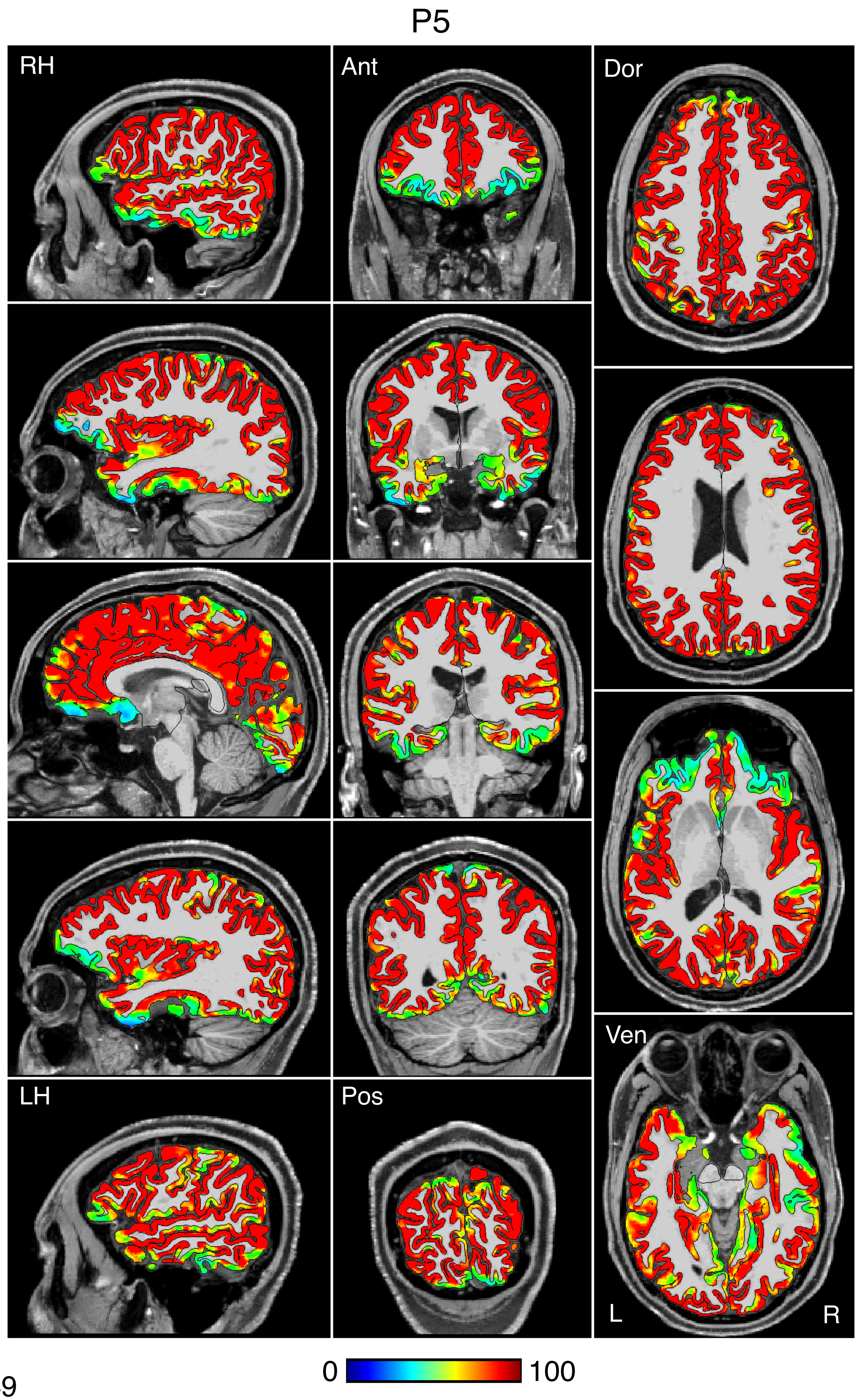


Figure S49

P5

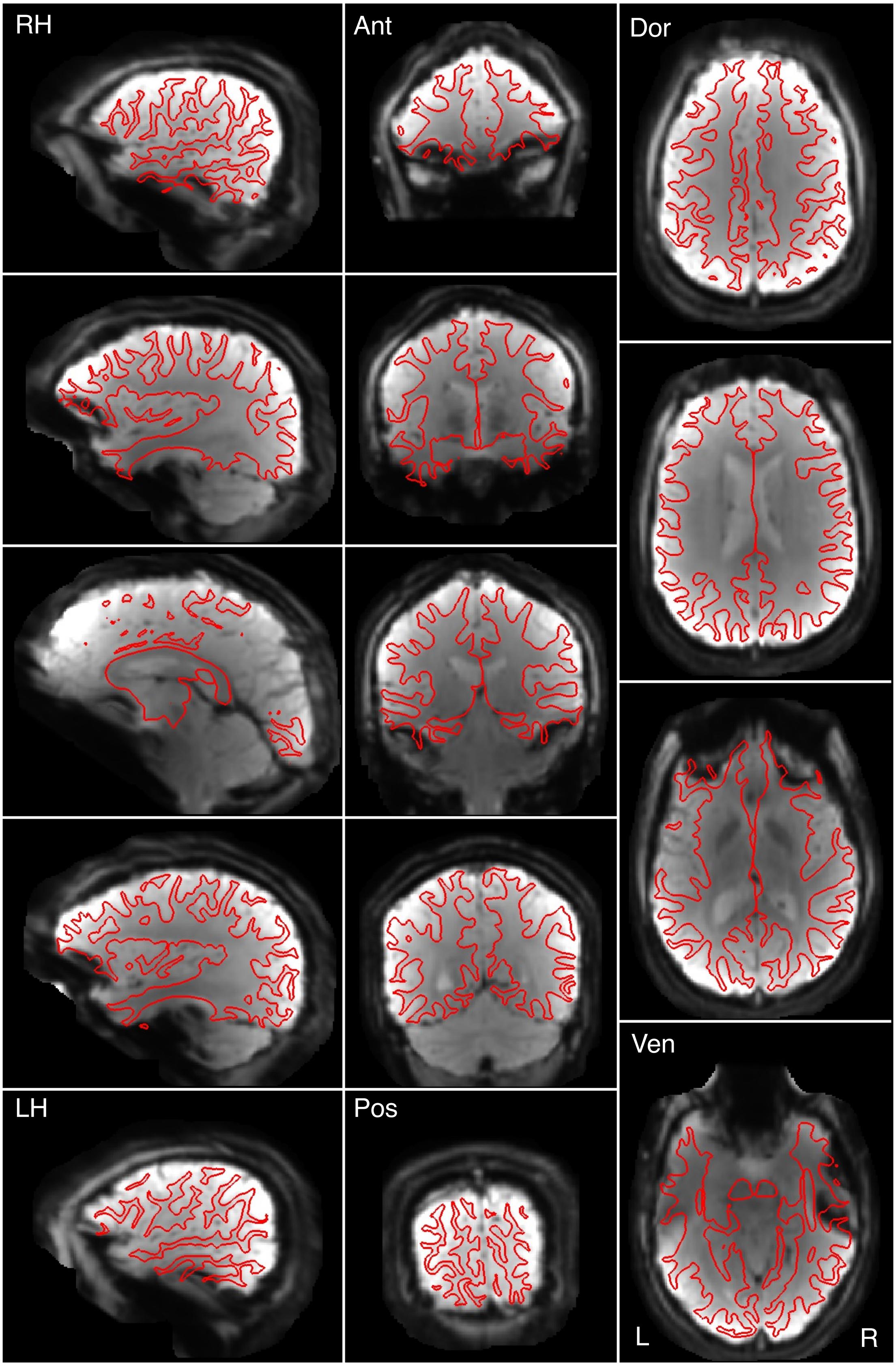


Figure S50

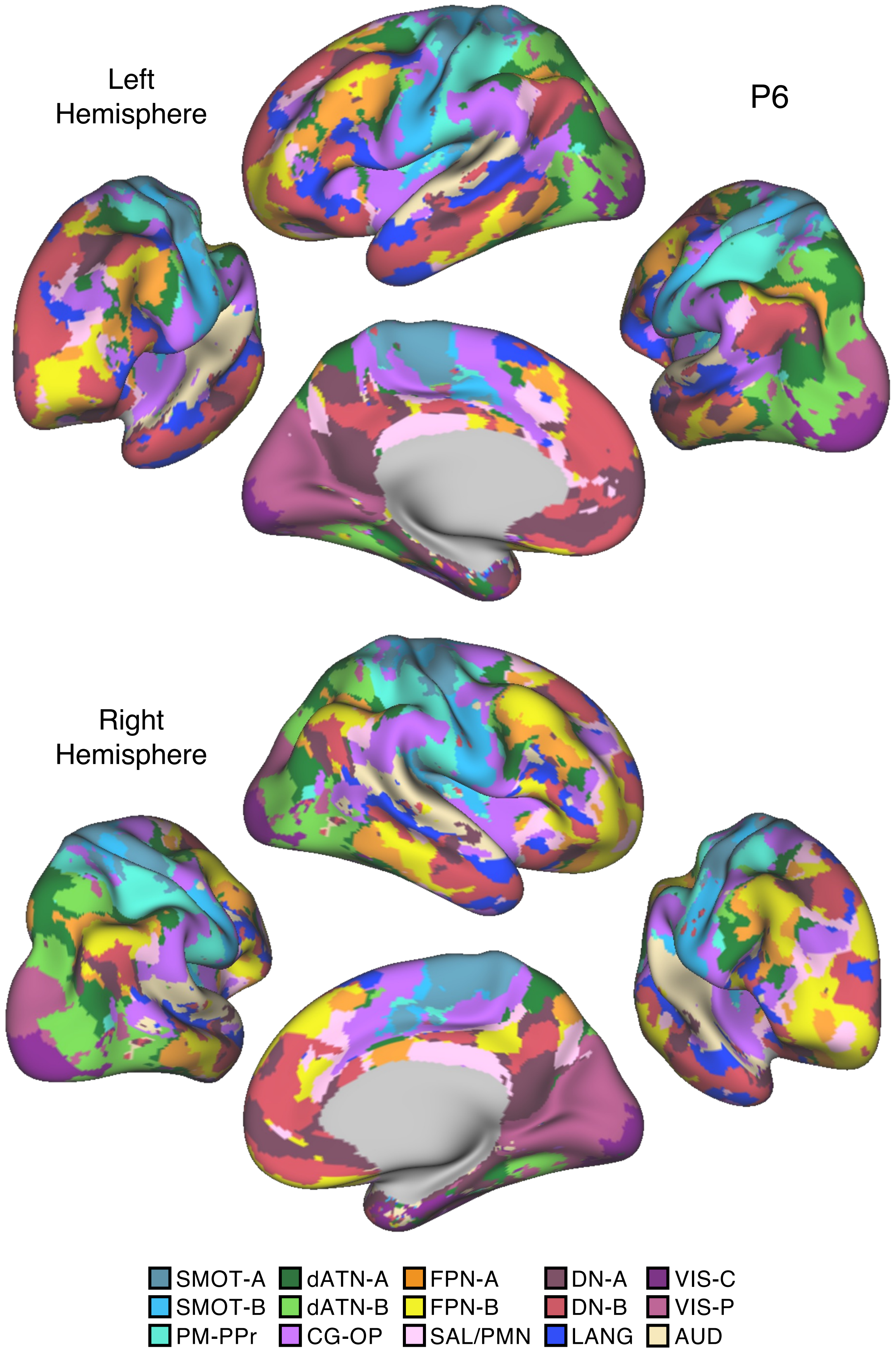
Left
Hemisphere
P6
Right
Hemisphere
SMOT-A
SMOT-B
PM-PPr
dATN-A
dATN-B
CG-OP
FPN-A
FPN-B
SAL/PMN
DN-A
DN-B
LANG
VIS-C
VIS-P
AUD

Figure S51

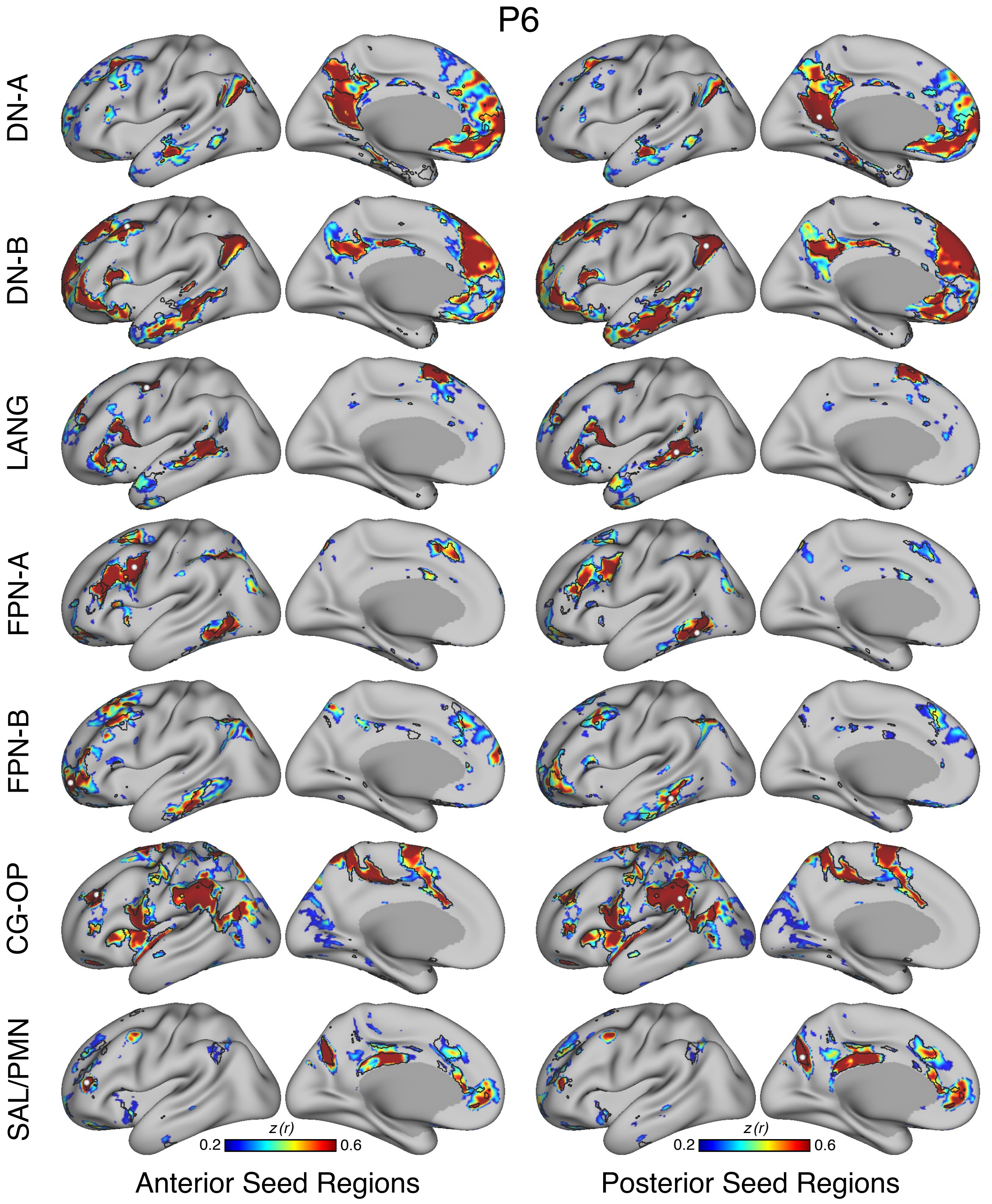
P6
DN-A
DN-B
LANG
FPN-A
FPN-B
CG-OP
SAL/PMN
z (r)
0.2
0.6
z (r)
0.2
0.6
Anterior Seed Regions
Posterior Seed Regions

Figure S52

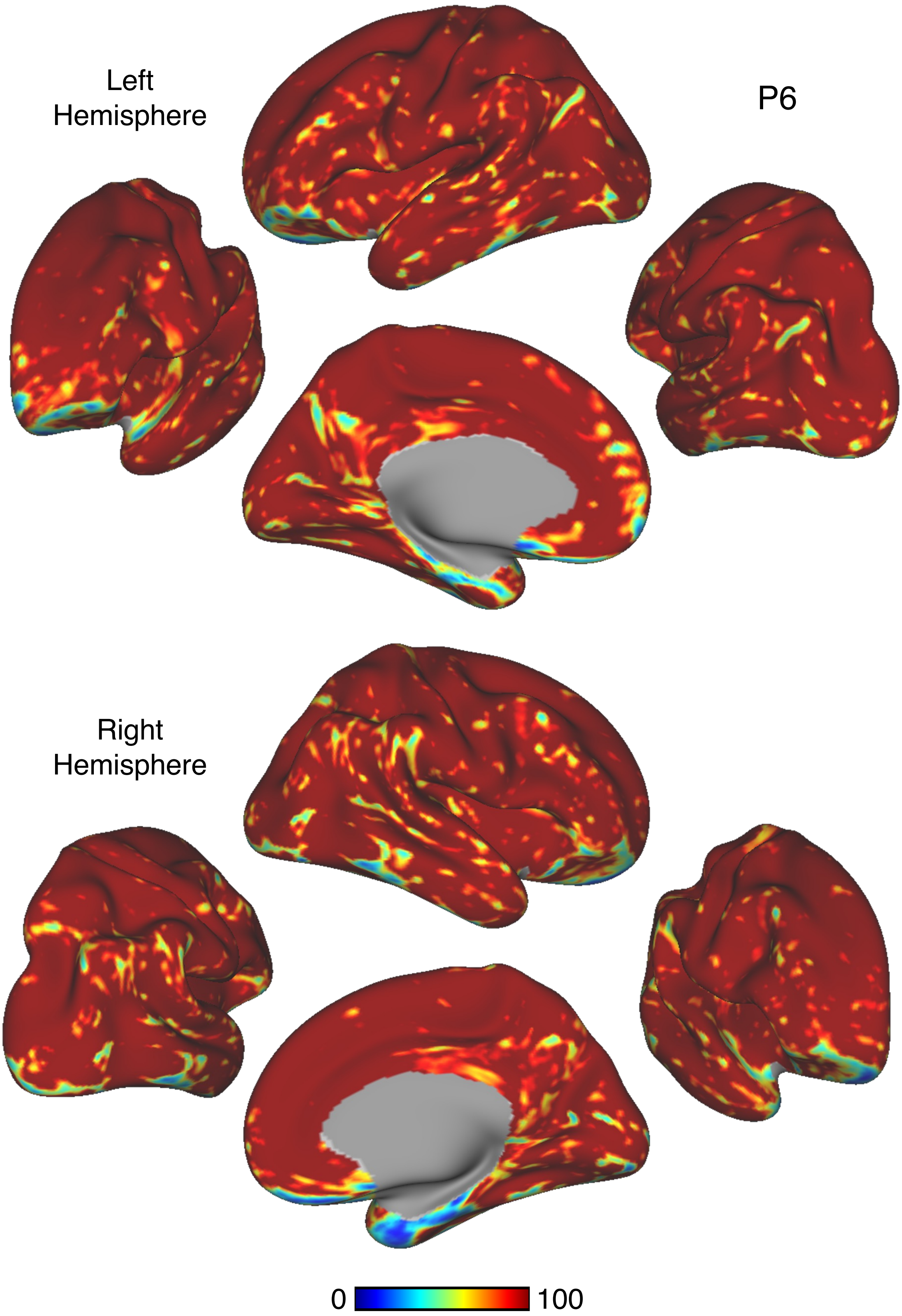


Figure S53

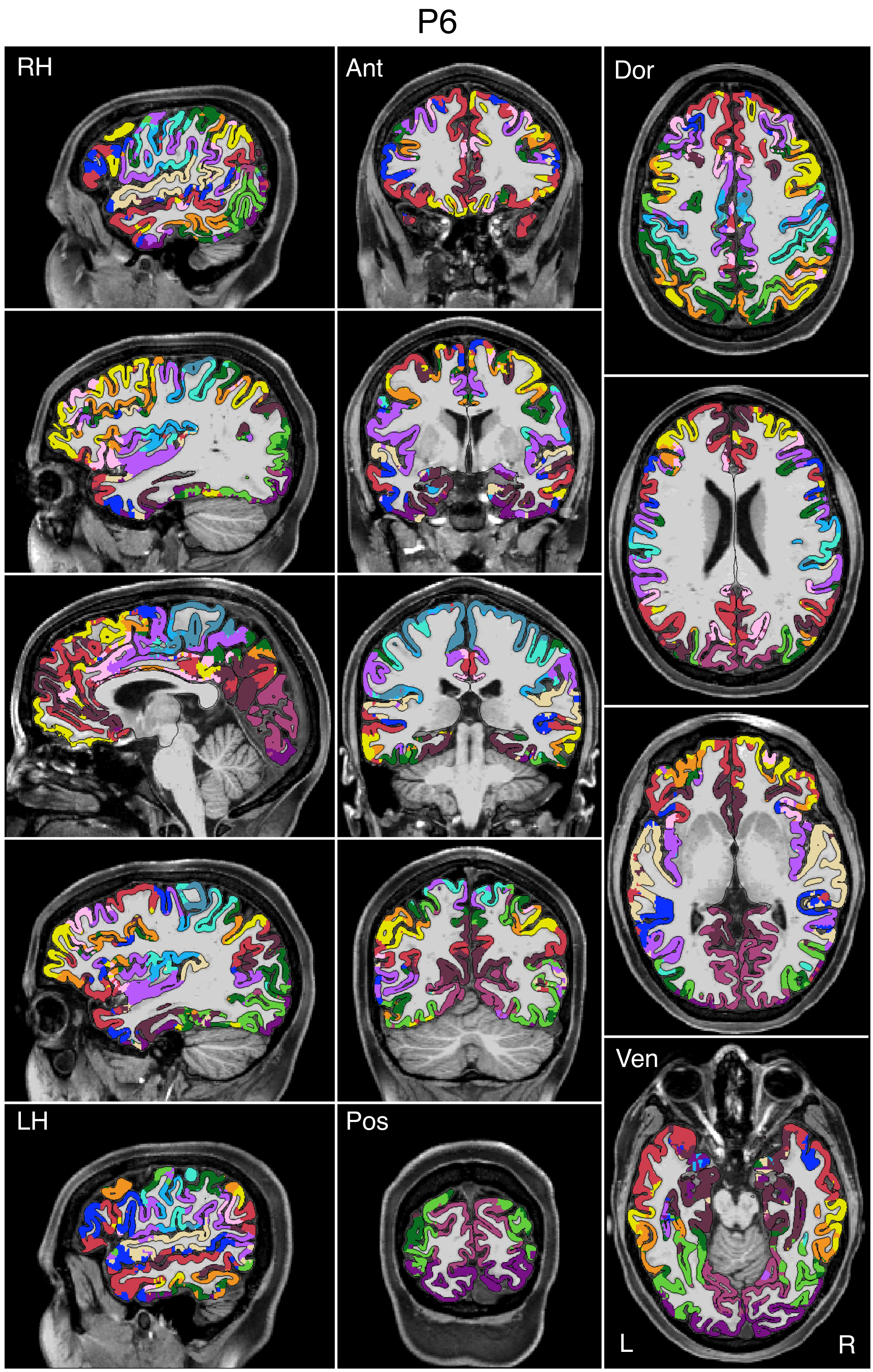


Figure S54

P6

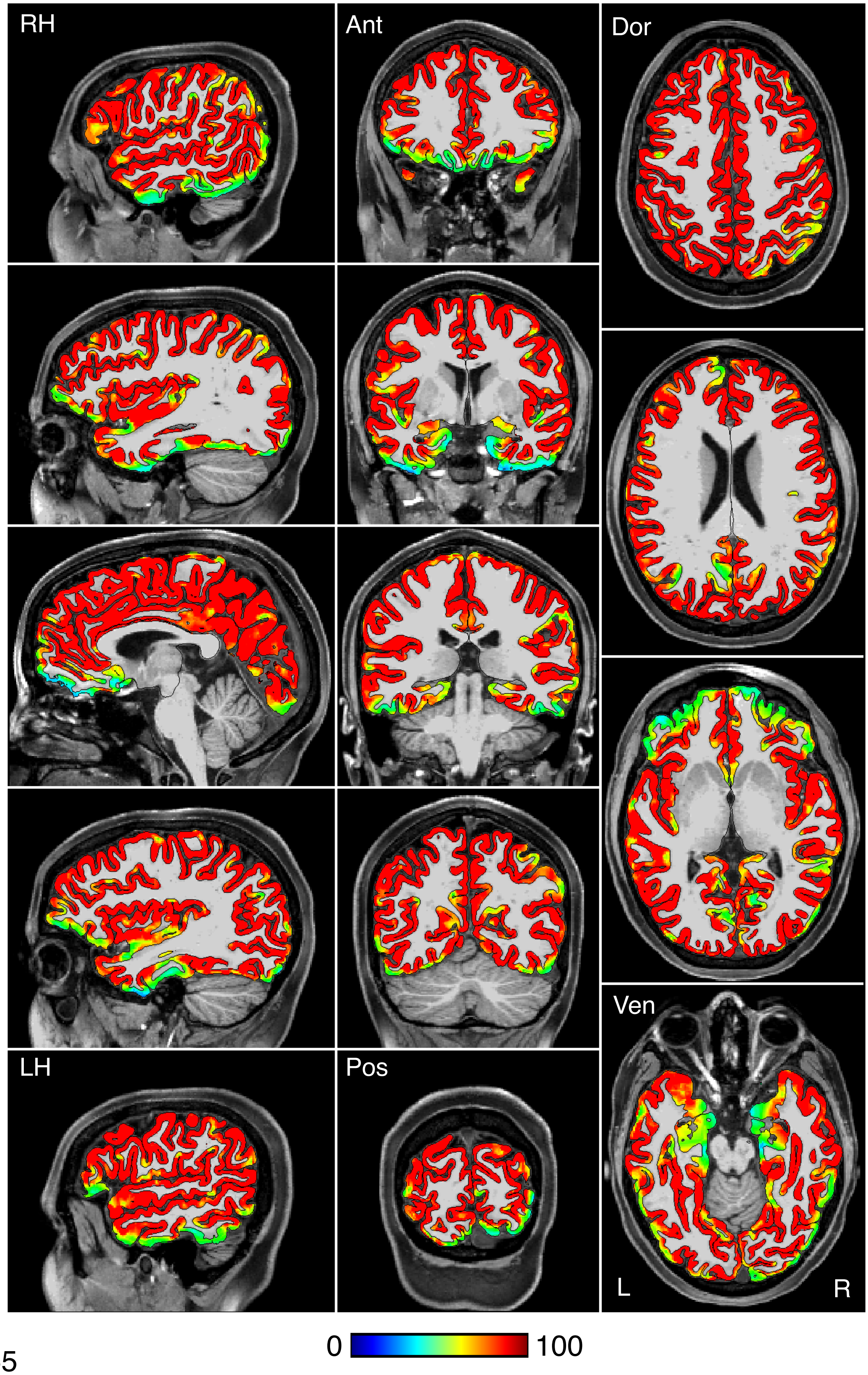


Figure S55

P6

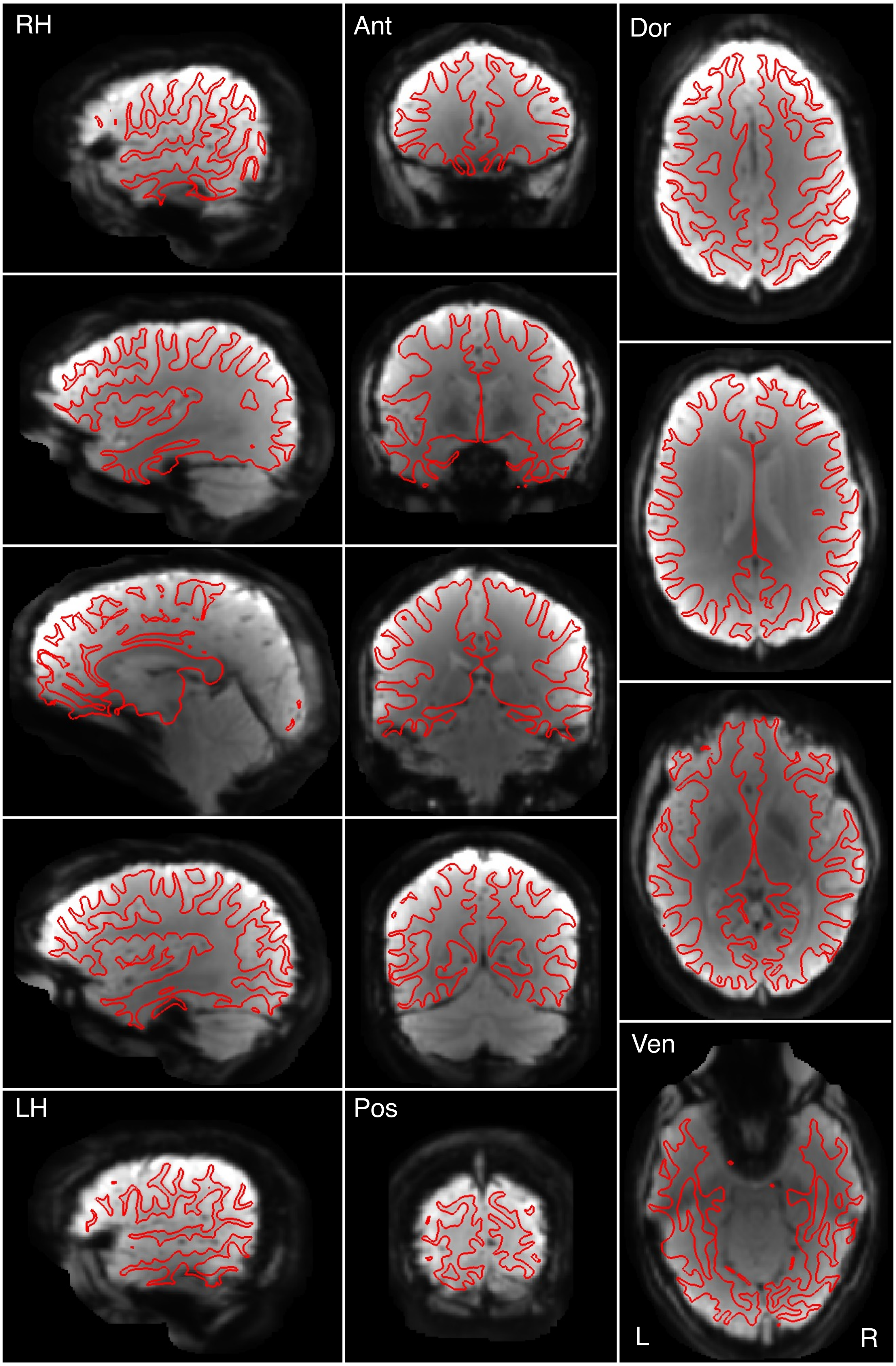


Figure S56

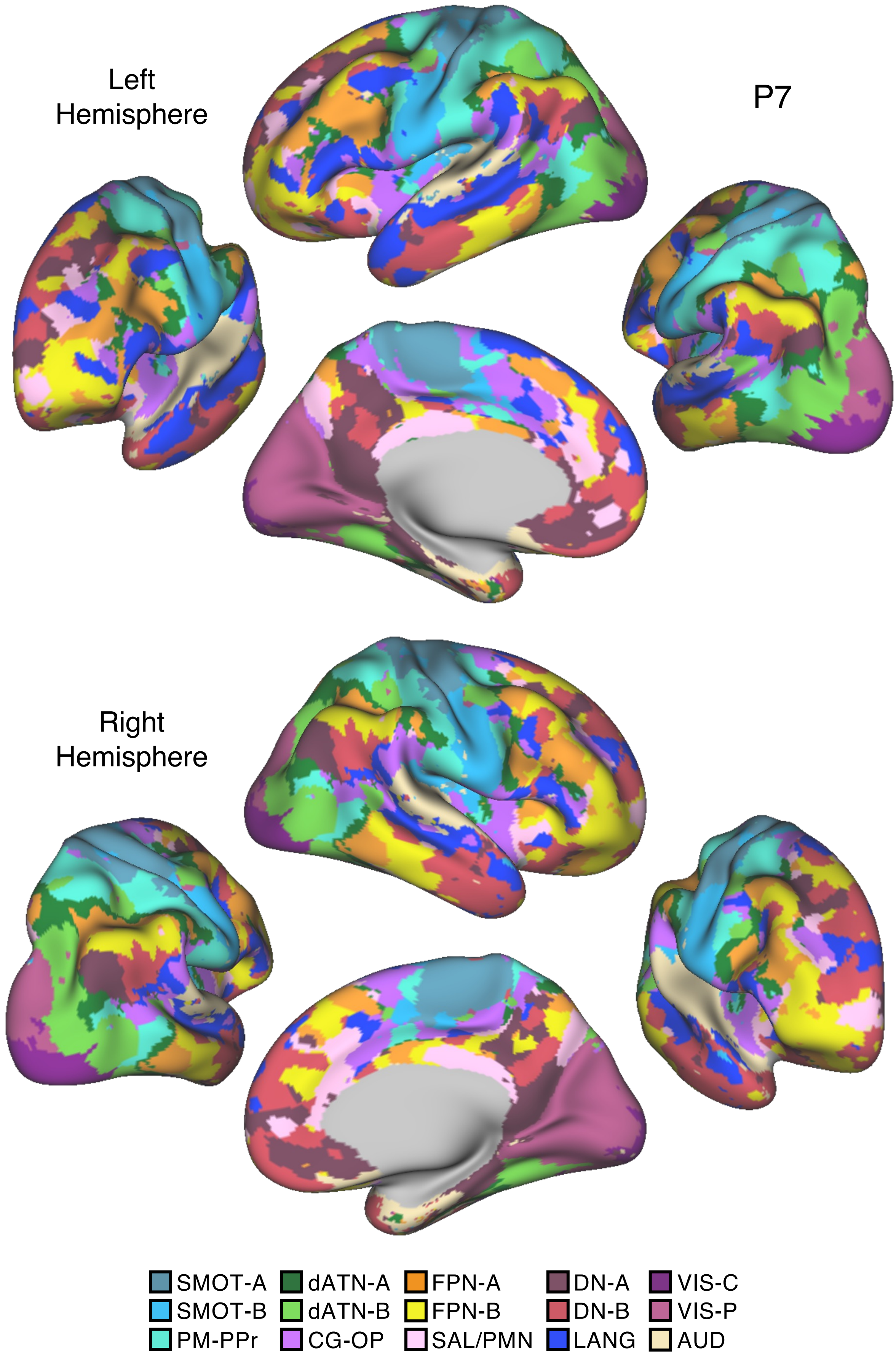

Left
Hemisphere
P7
Right
Hemisphere
SMOT-A
SMOT-B
PM-PPr
dATN-A
dATN-B
CG-OP
FPN-A
FPN-B
SAL/PMN
DN-A
DN-B
LANG
VIS-C
VIS-P
AUD


Figure S57

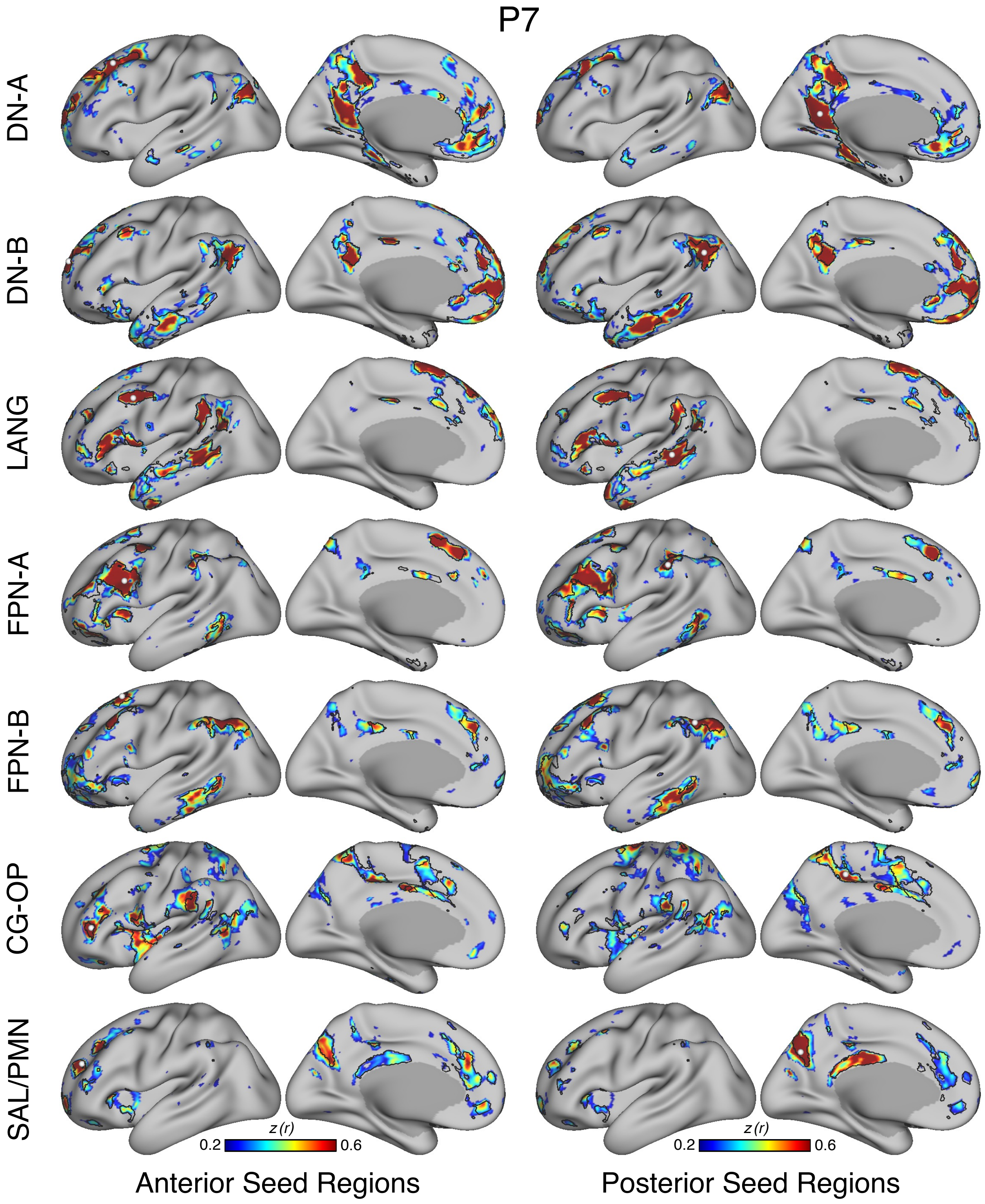

P7
DN-A
DN-B
LANG
FPN-A
FPN-B
CG-OP
SAL/PMN
z (r)
0.2
0.6
z (r)
0.2
0.6
Anterior Seed Regions
Posterior Seed Regions


Figure S58

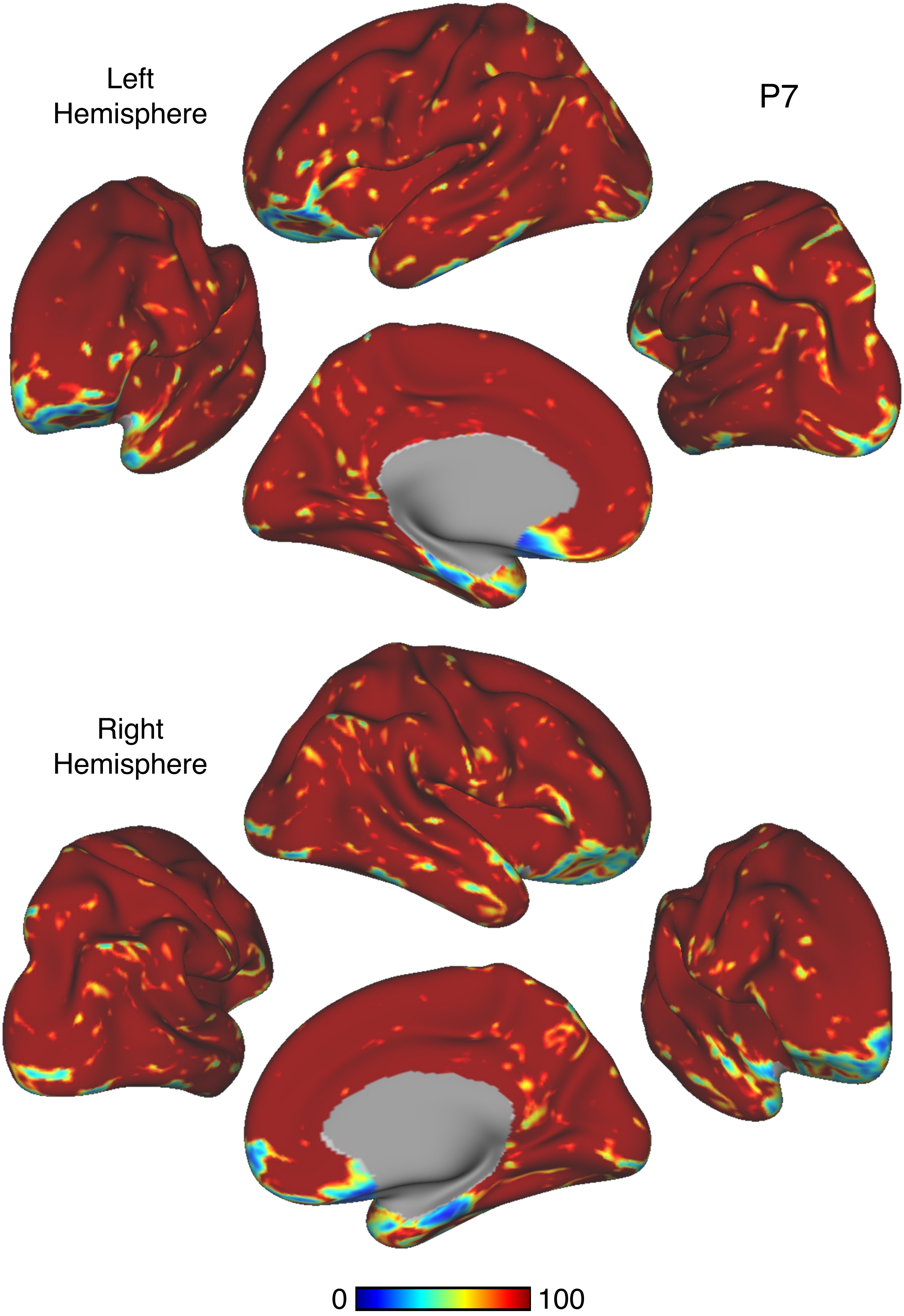


Figure S59

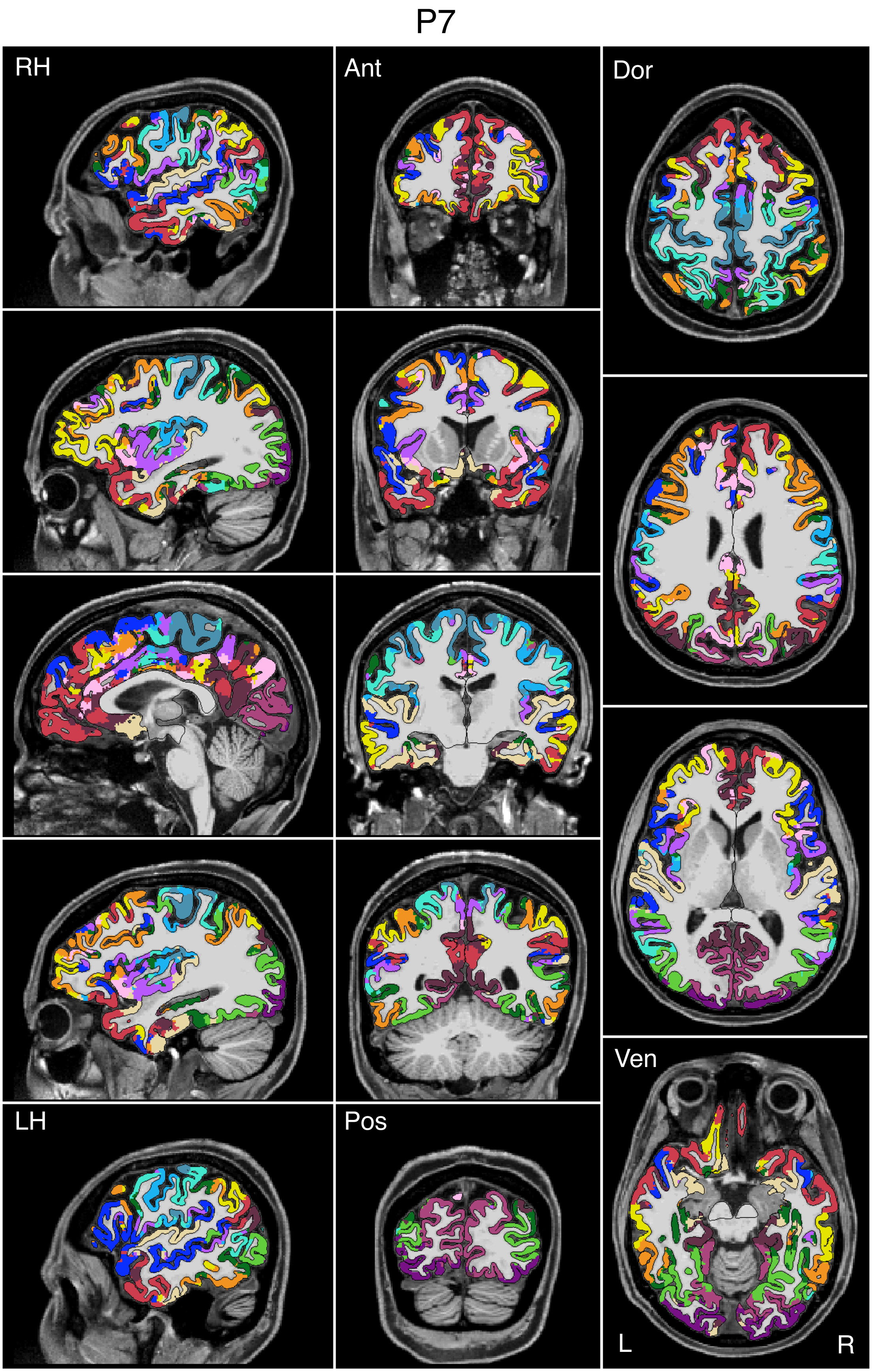


Figure S60

P7

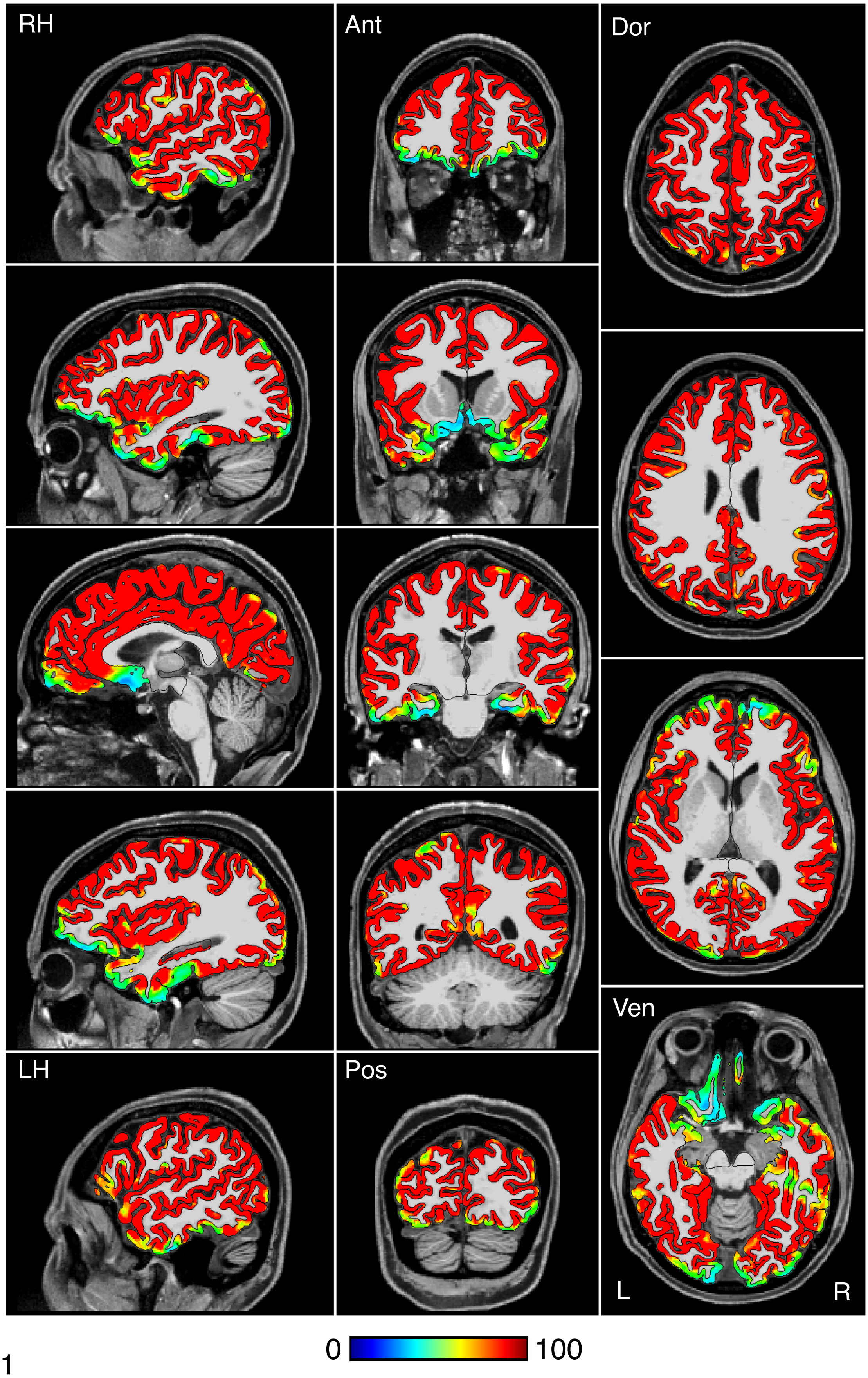


Figure S61

P7

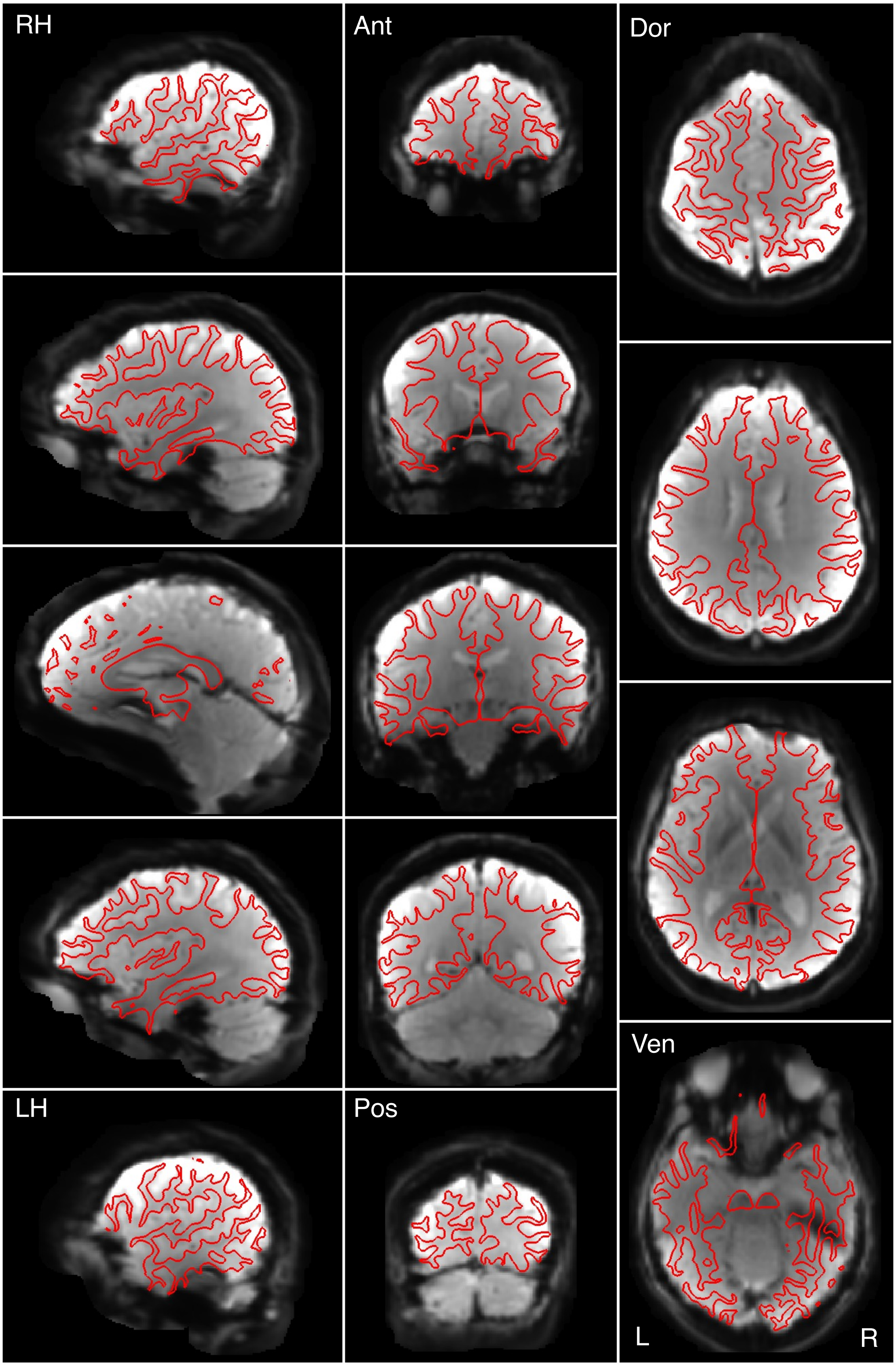


Figure S62

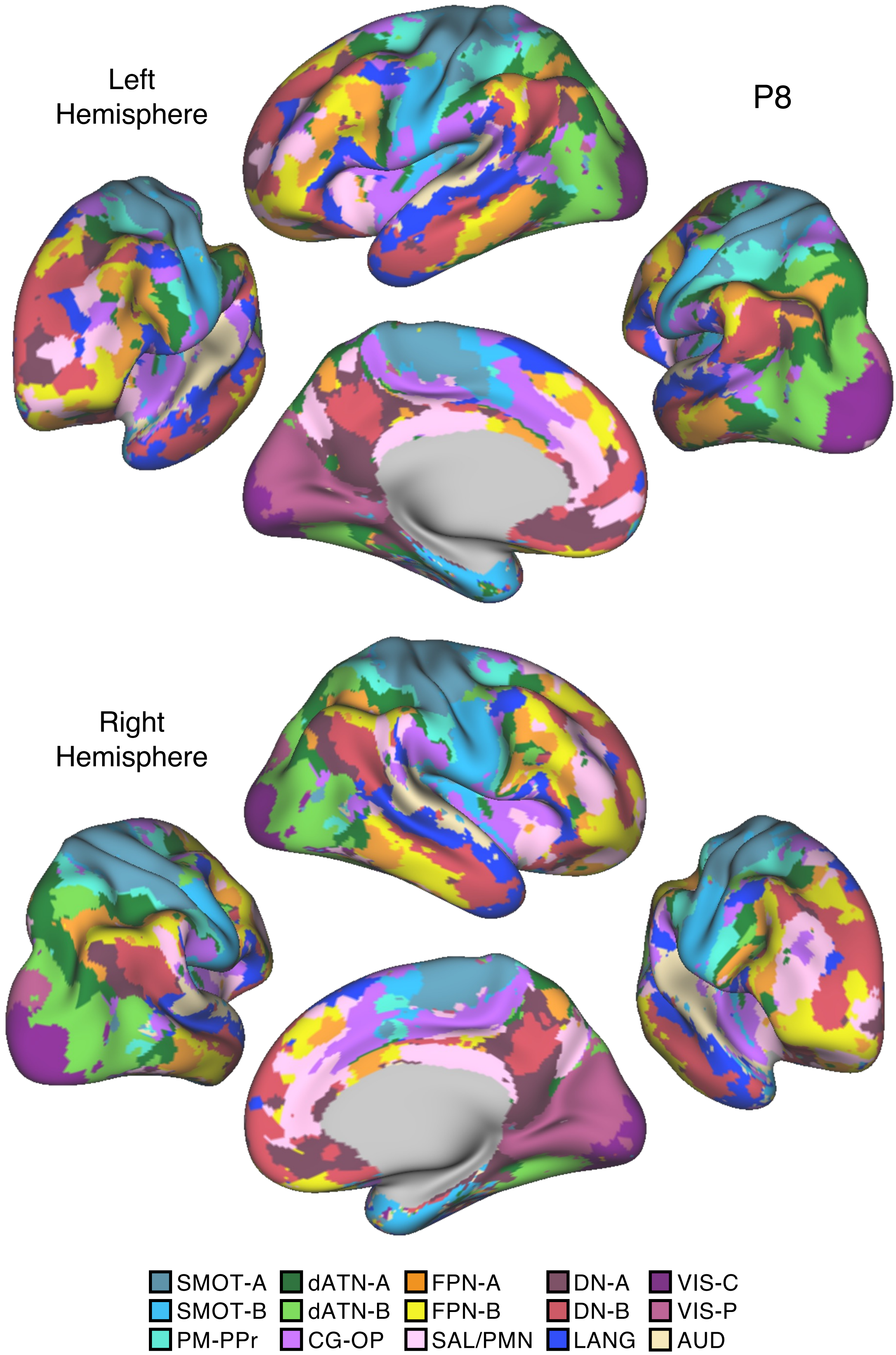
Left
Hemisphere
P8
Right
Hemisphere
SMOT-A
SMOT-B
PM-PPr
dATN-A
dATN-B
CG-OP
FPN-A
FPN-B
SAL/PMN
DN-A
DN-B
LANG
VIS-C
VIS-P
AUD

Figure S63

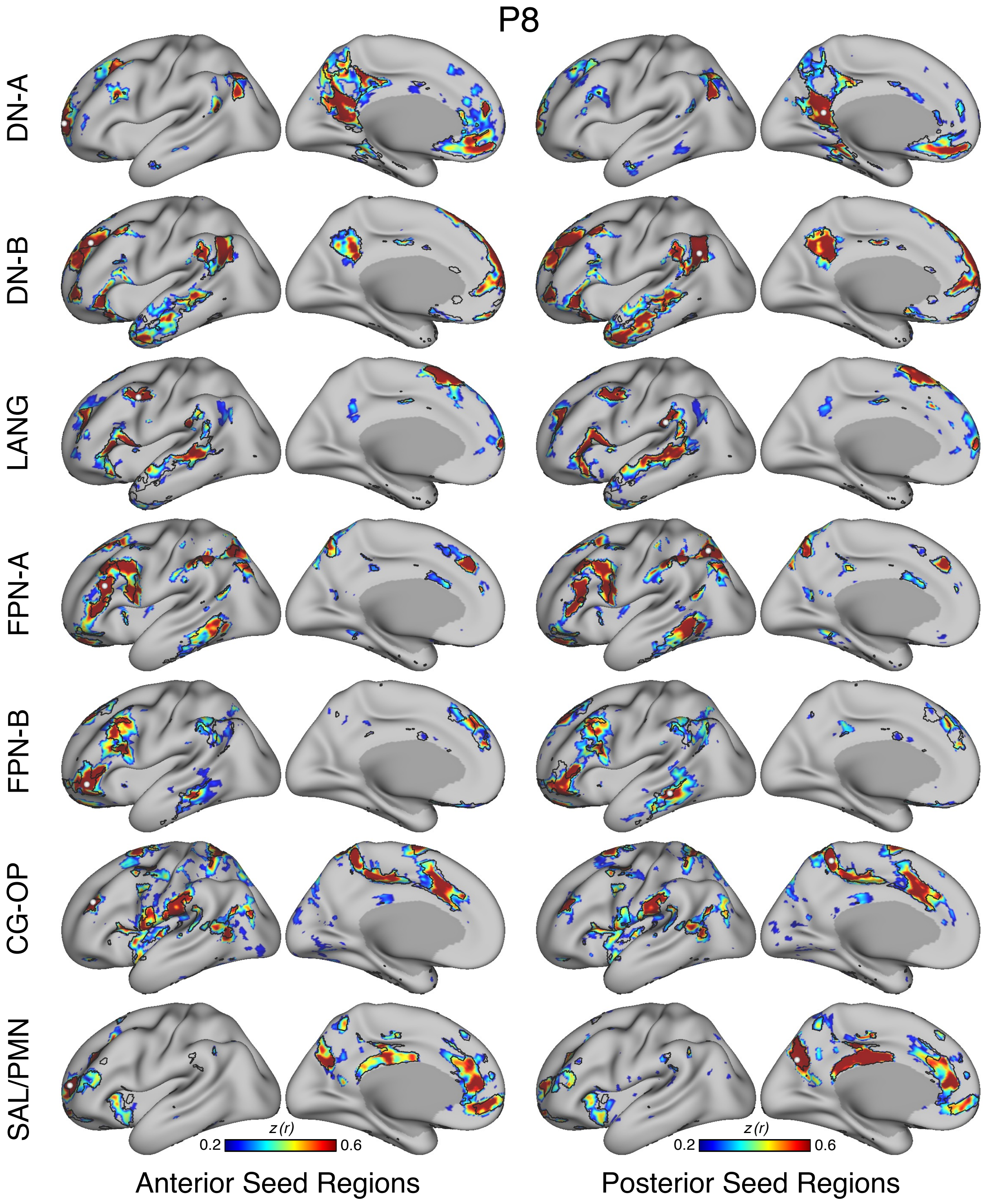
P8
DN-A
DN-B
LANG
FPN-A
FPN-B
CG-OP
SAL/PMN
z (r)
0.2
0.6
z (r)
0.2
0.6
Anterior Seed Regions
Posterior Seed Regions

Figure S64

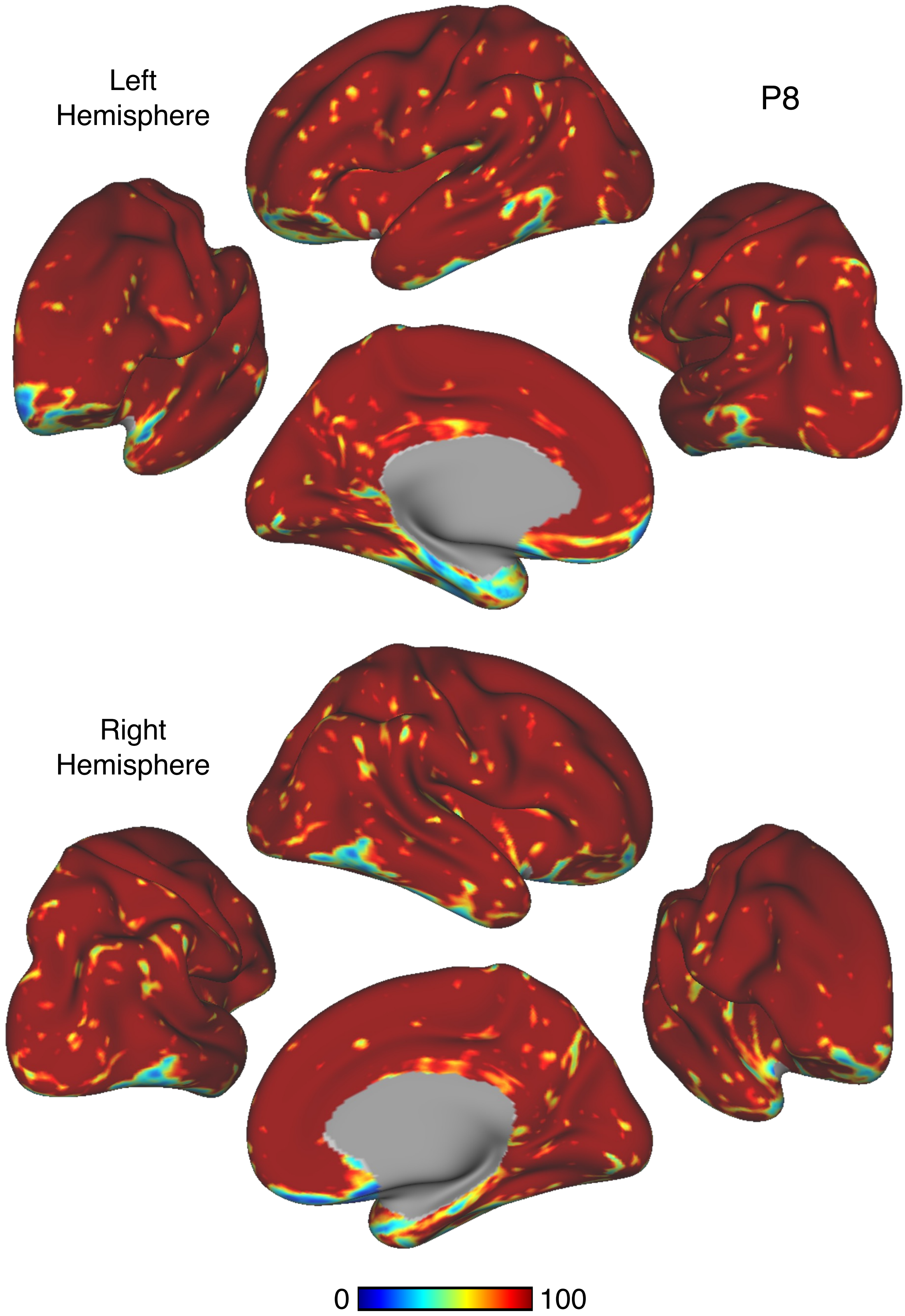


Figure S65

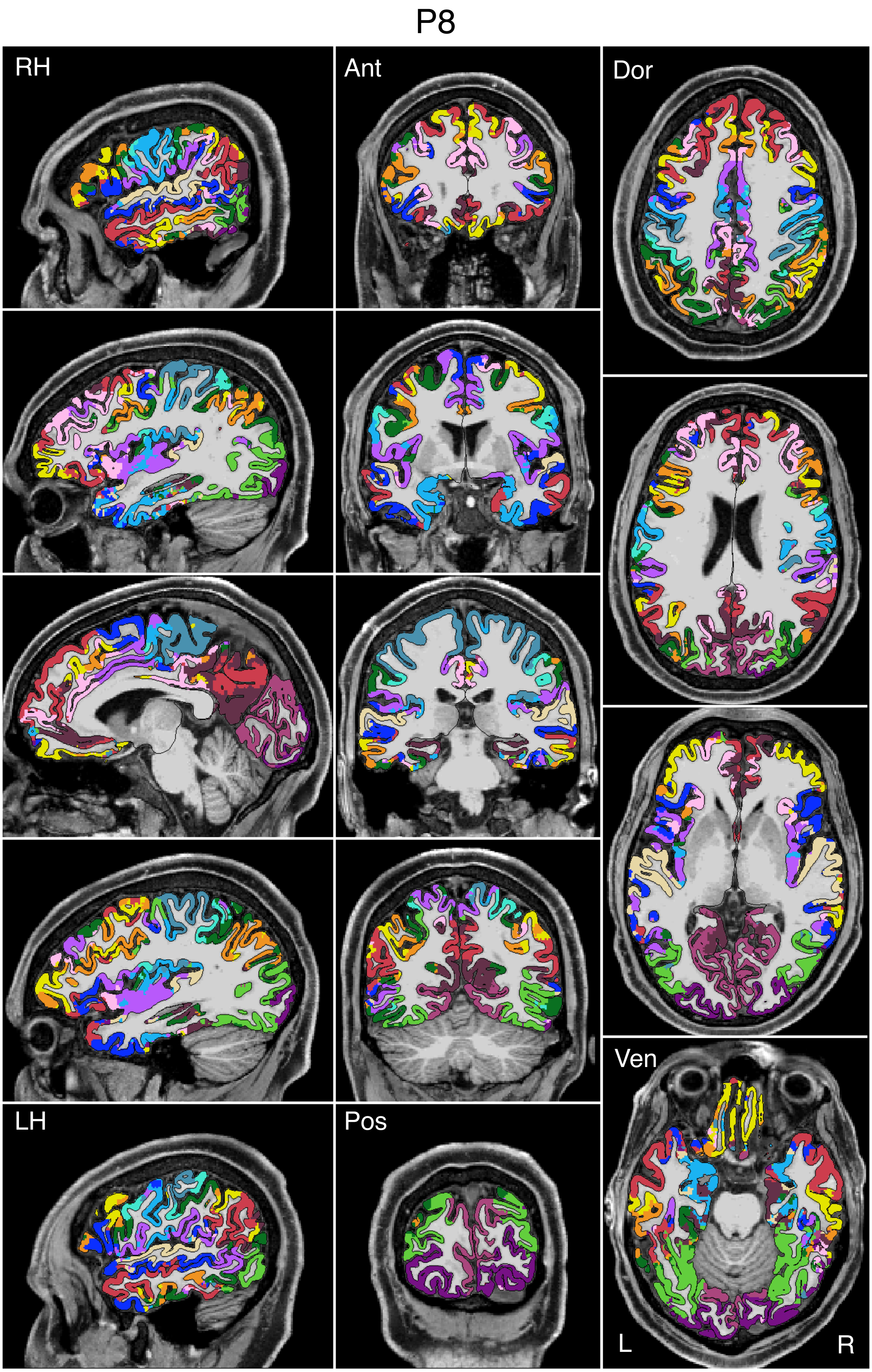


Figure S66

P8

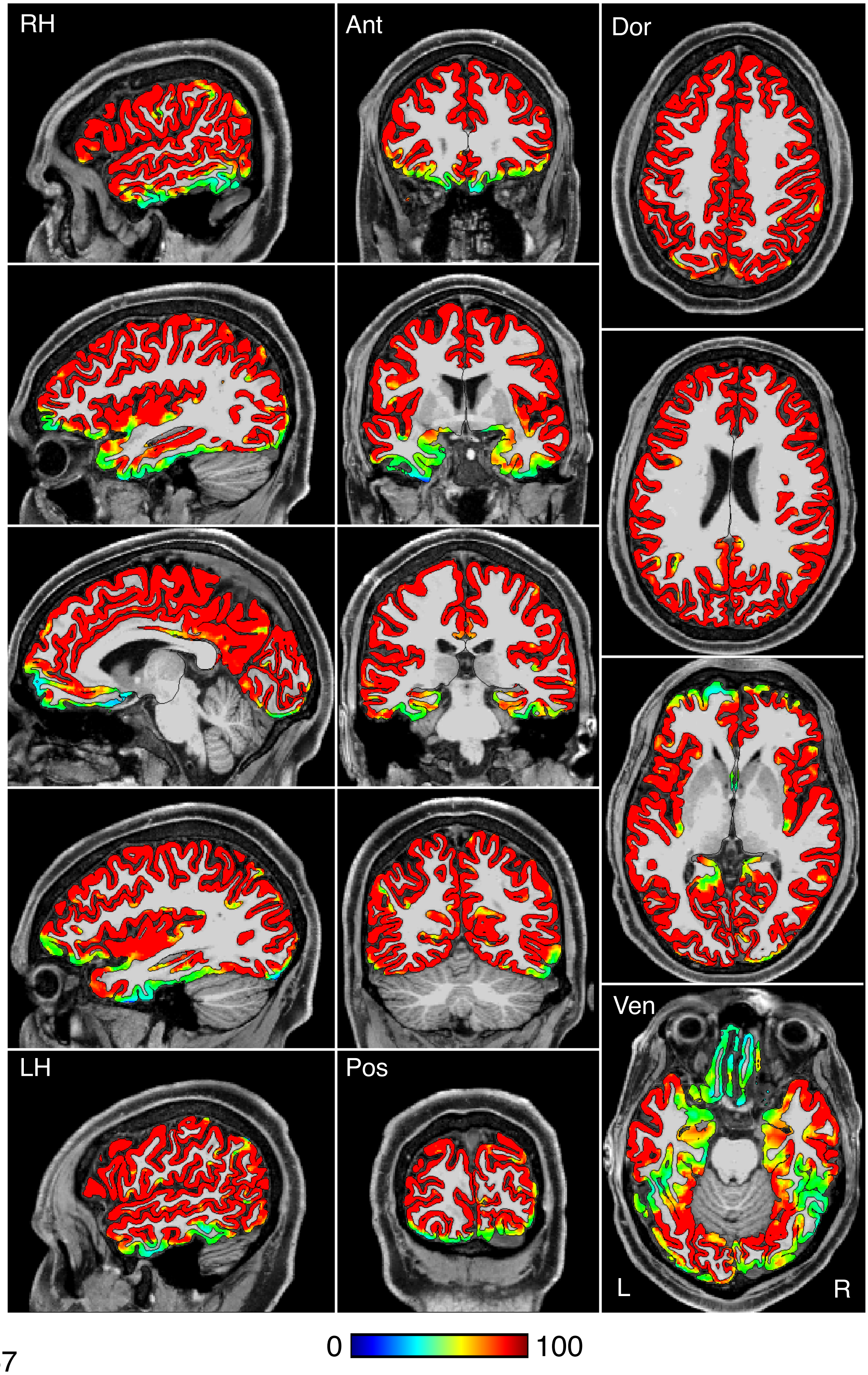


Figure S67

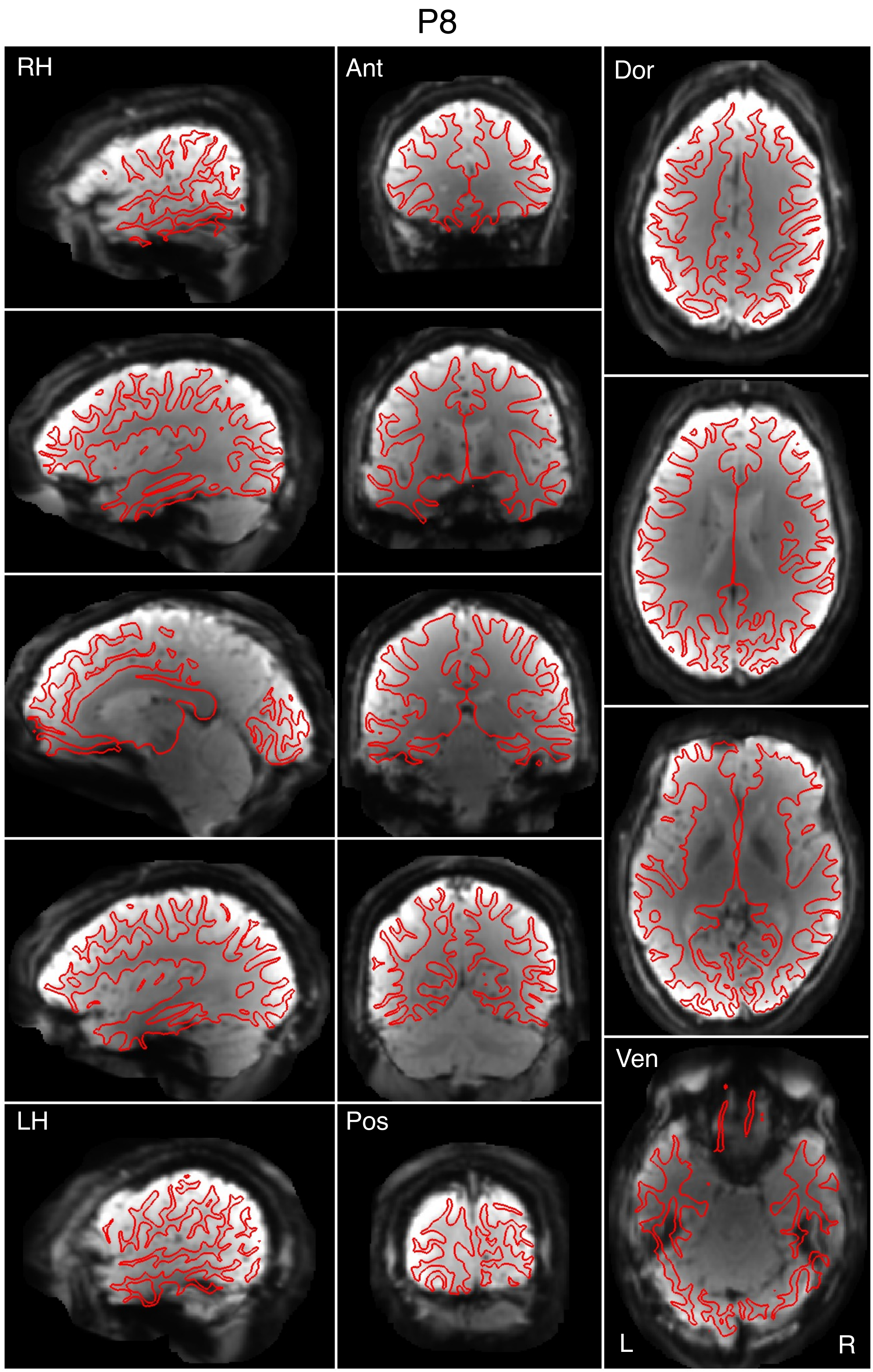


Figure S68

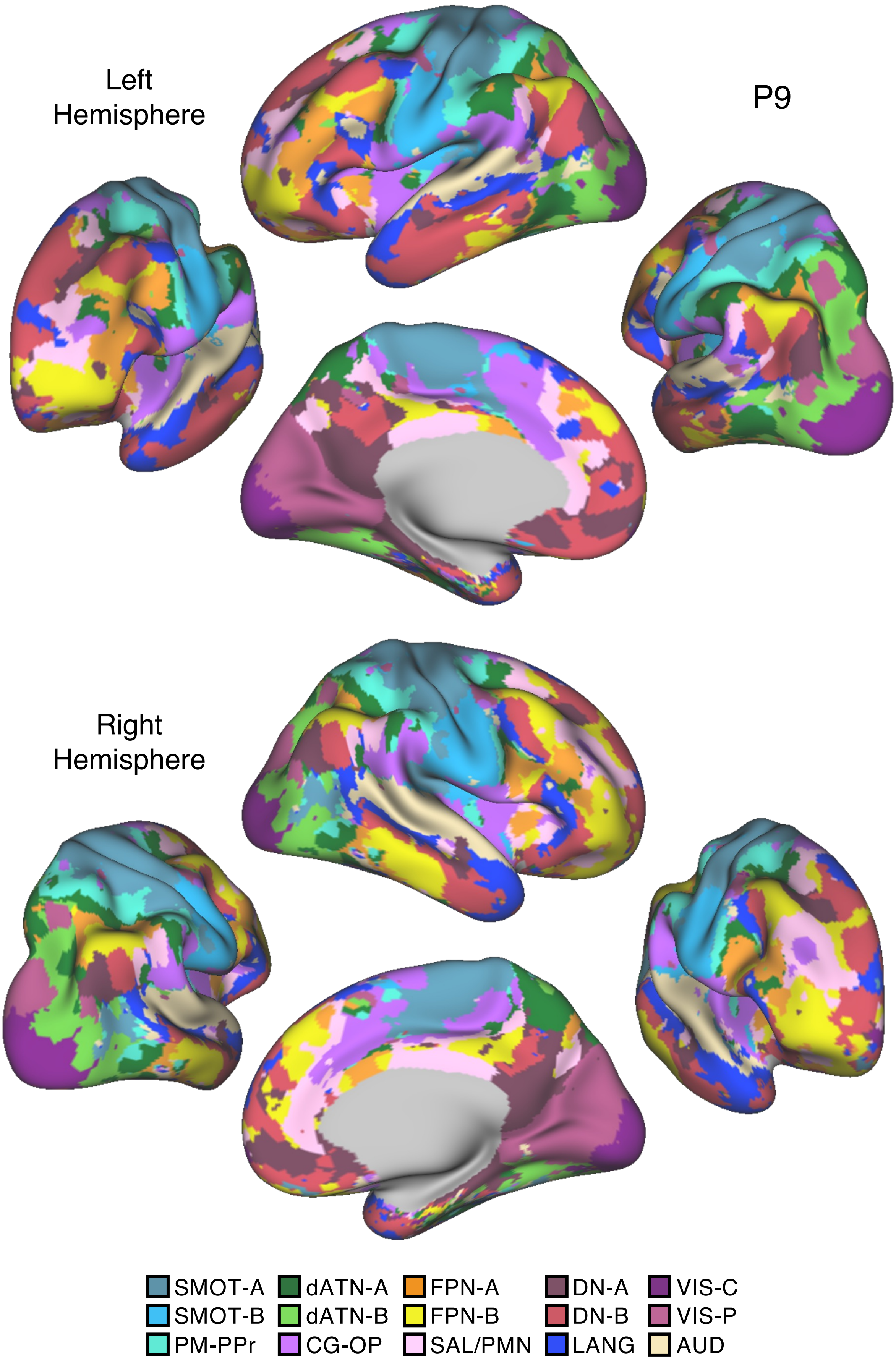

Left
Hemisphere
P9
Right
Hemisphere
SMOT-A
SMOT-B
PM-PPr
dATN-A
dATN-B
CG-OP
FPN-A
FPN-B
SAL/PMN
DN-A
DN-B
LANG
VIS-C
VIS-P
AUD


Figure S69

# P9

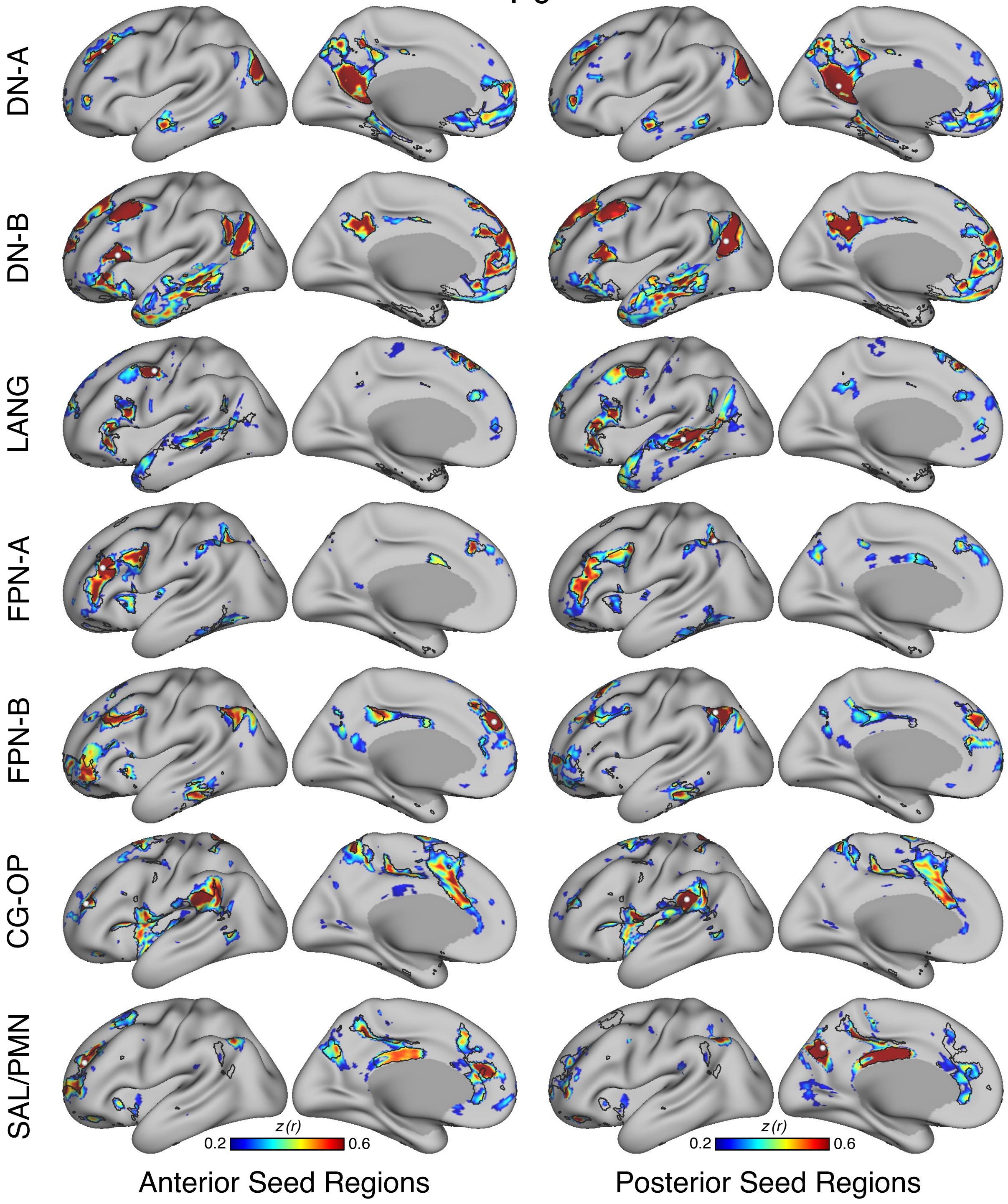


Figure S70

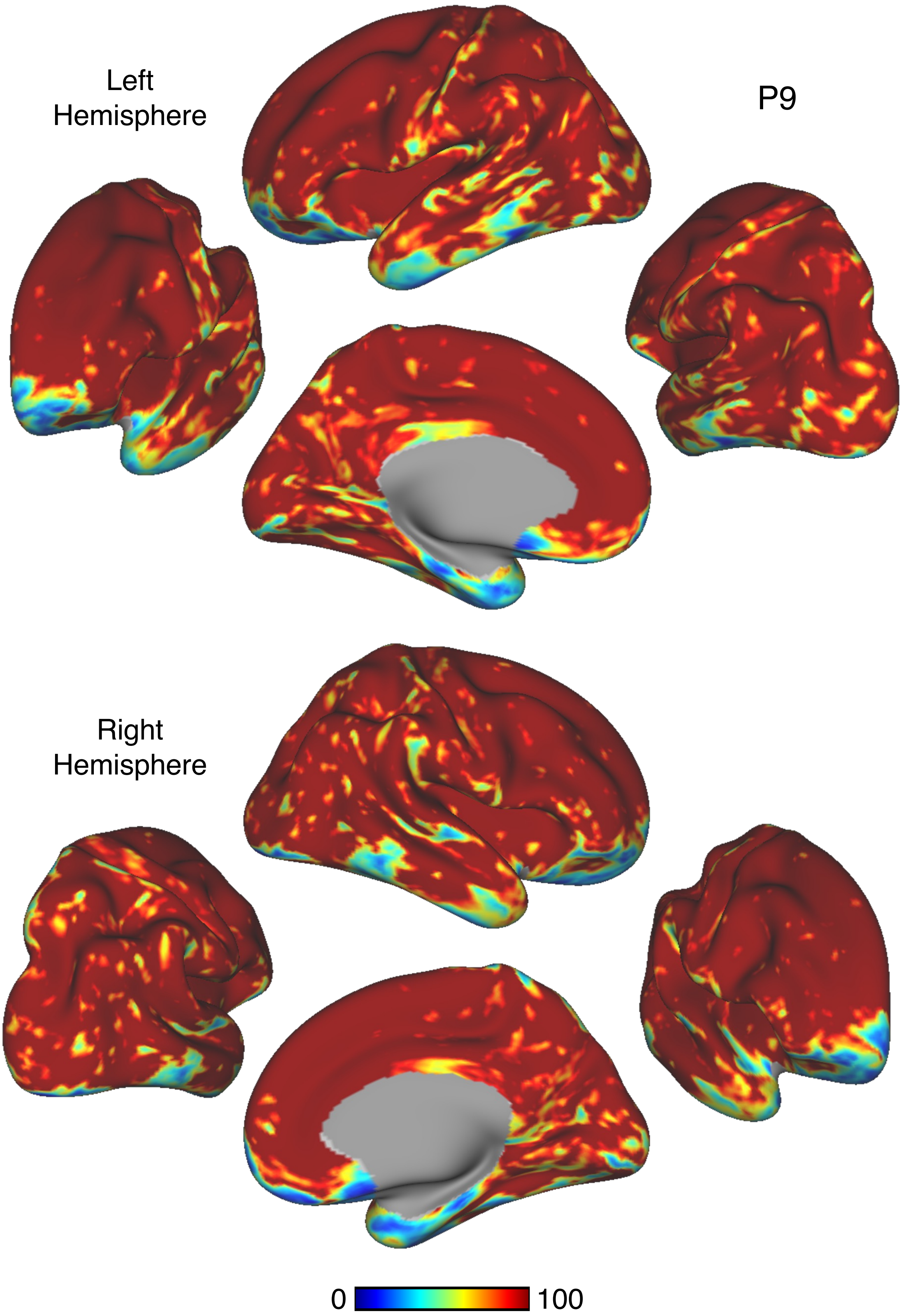


Figure S71

P9

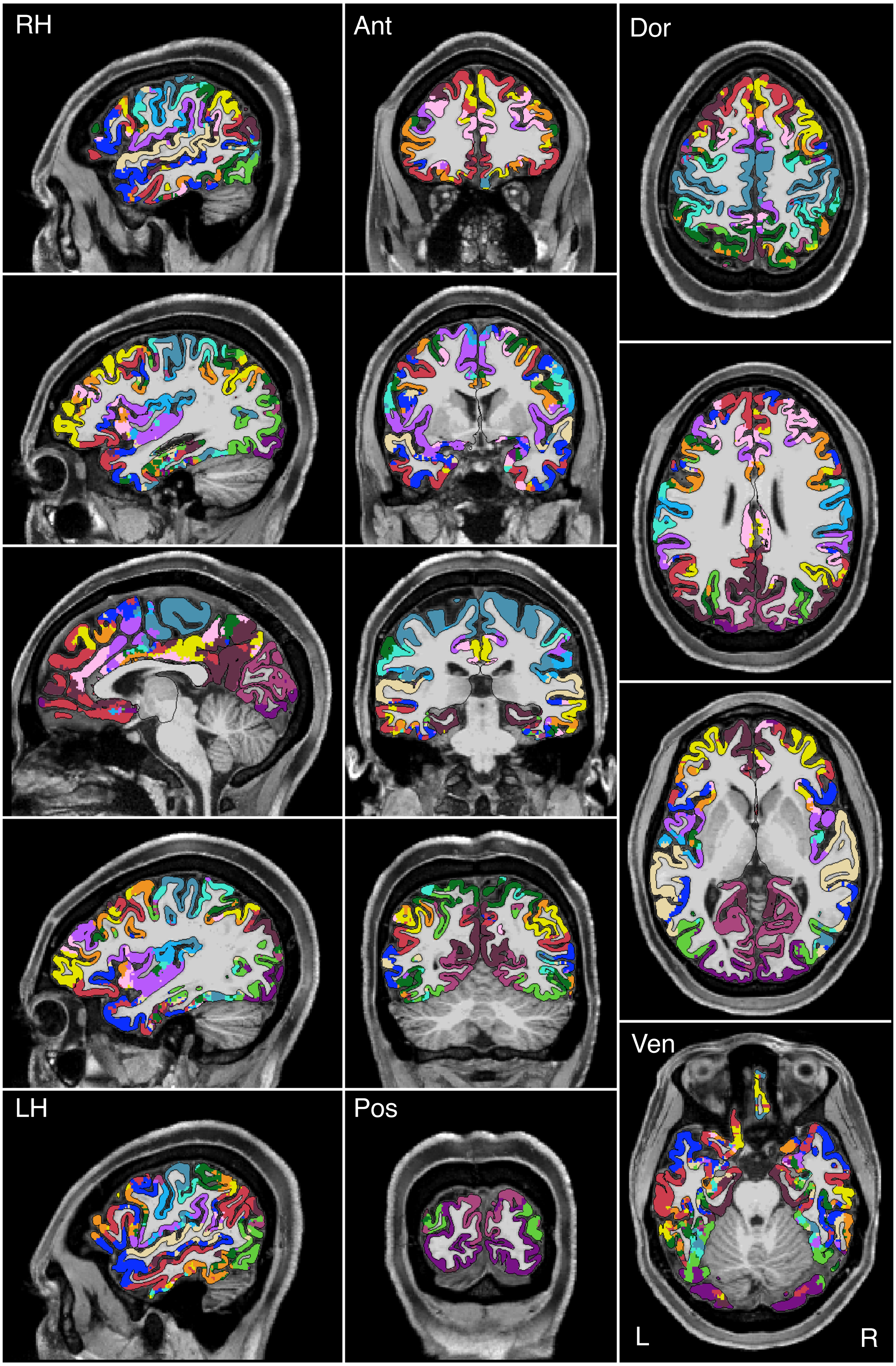


Figure S72

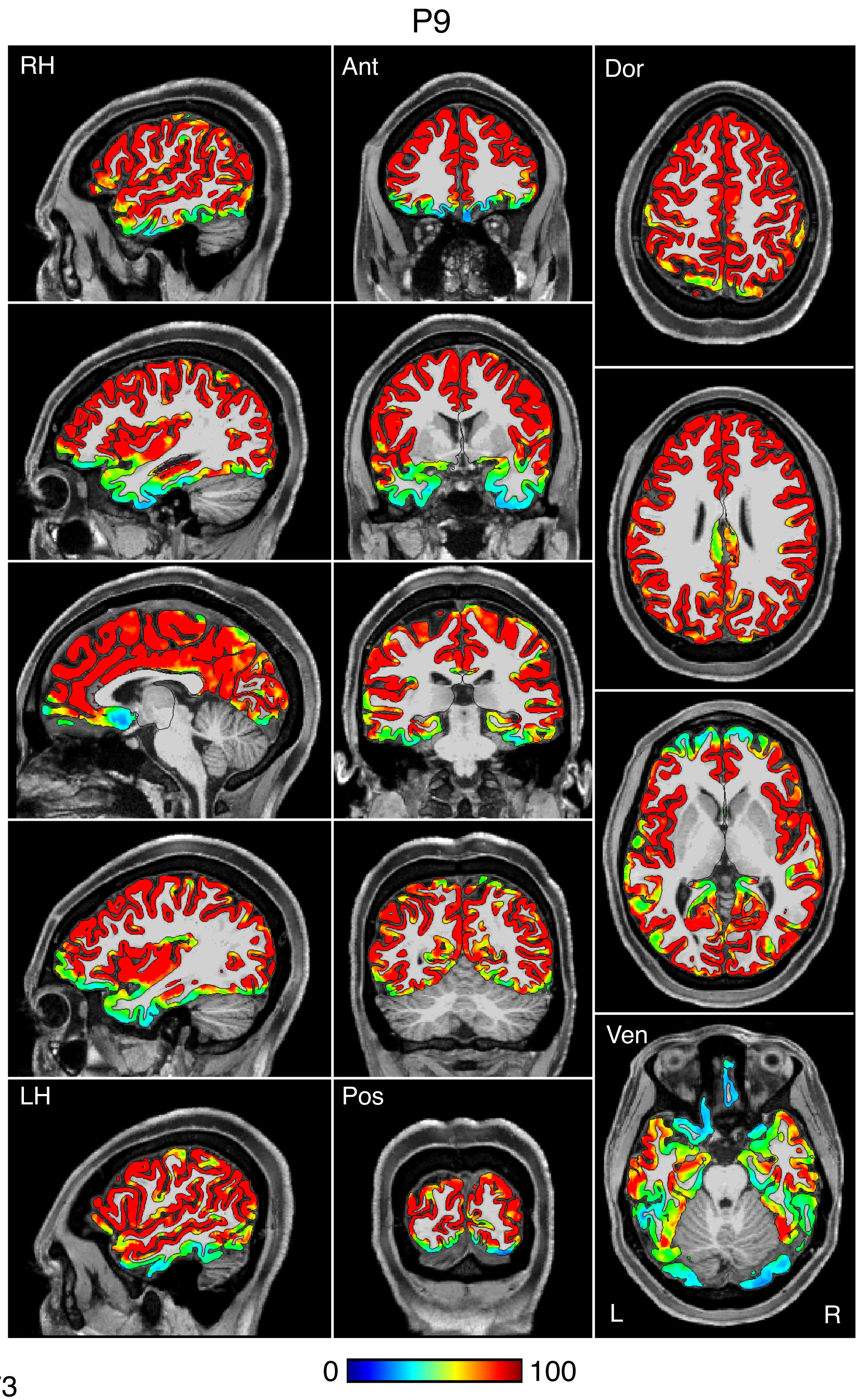

Figure S73

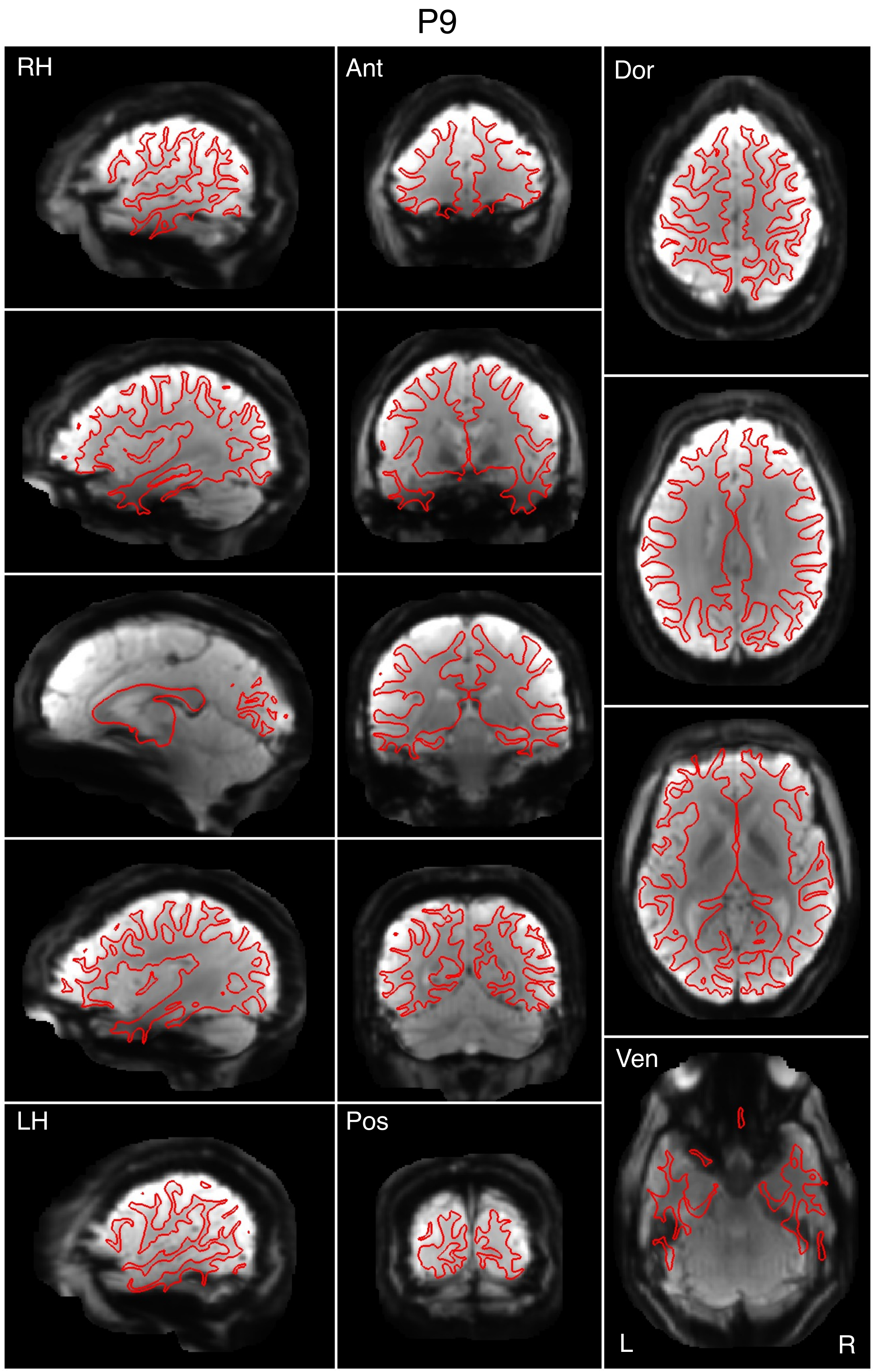


Figure S74

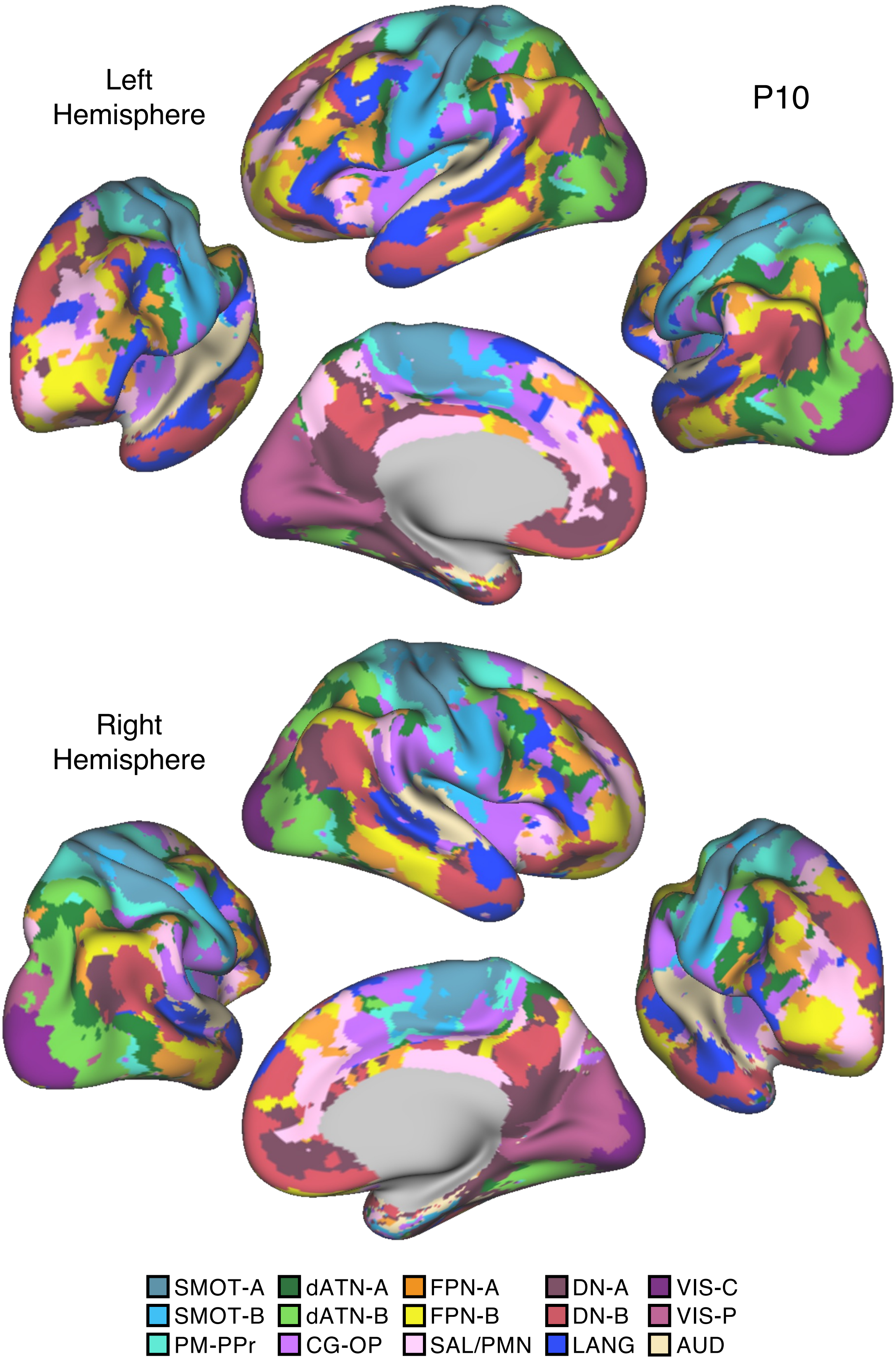

Left
Hemisphere
P10
Right
Hemisphere
SMOT-A
SMOT-B
PM-PPr
dATN-A
dATN-B
CG-OP
FPN-A
FPN-B
SAL/PMN
DN-A
DN-B
LANG
VIS-C
VIS-P
AUD


Figure S75

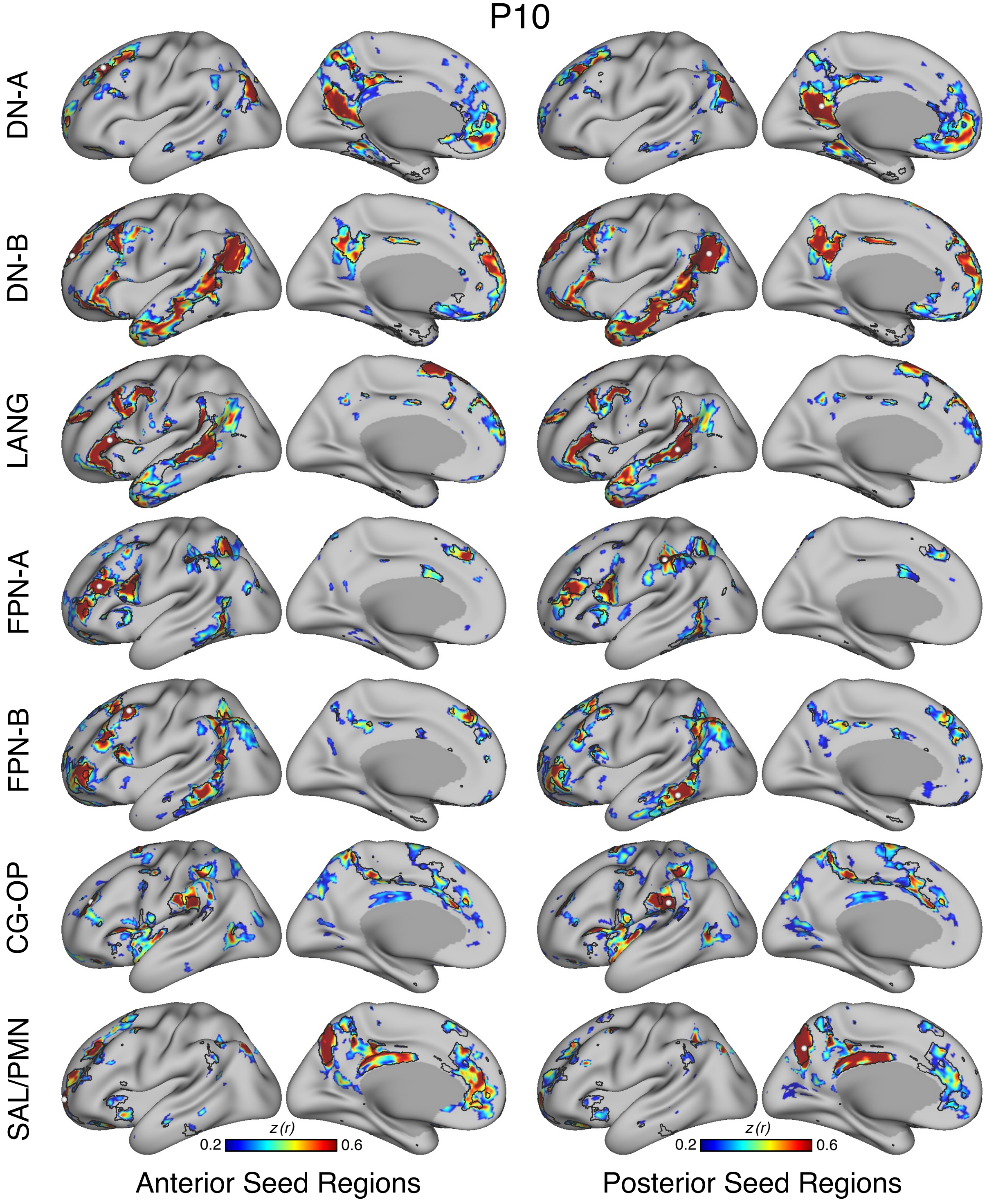
P10
DN-A
DN-B
LANG
FPN-A
FPN-B
CG-OP
SAL/PMN
z (r)
0.2
0.6
z (r)
0.2
0.6
Anterior Seed Regions
Posterior Seed Regions

Figure S76

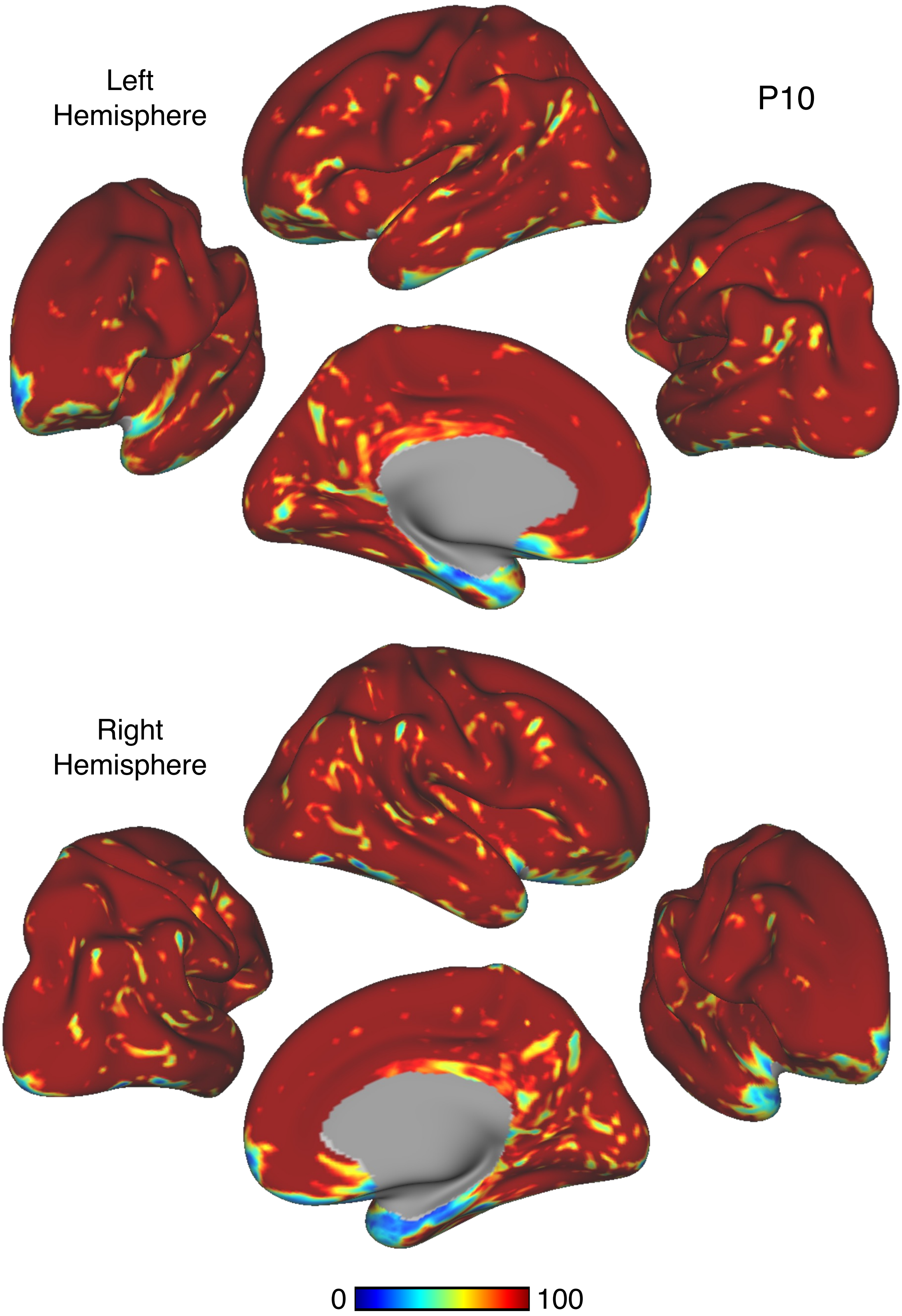


Figure S77

# P10

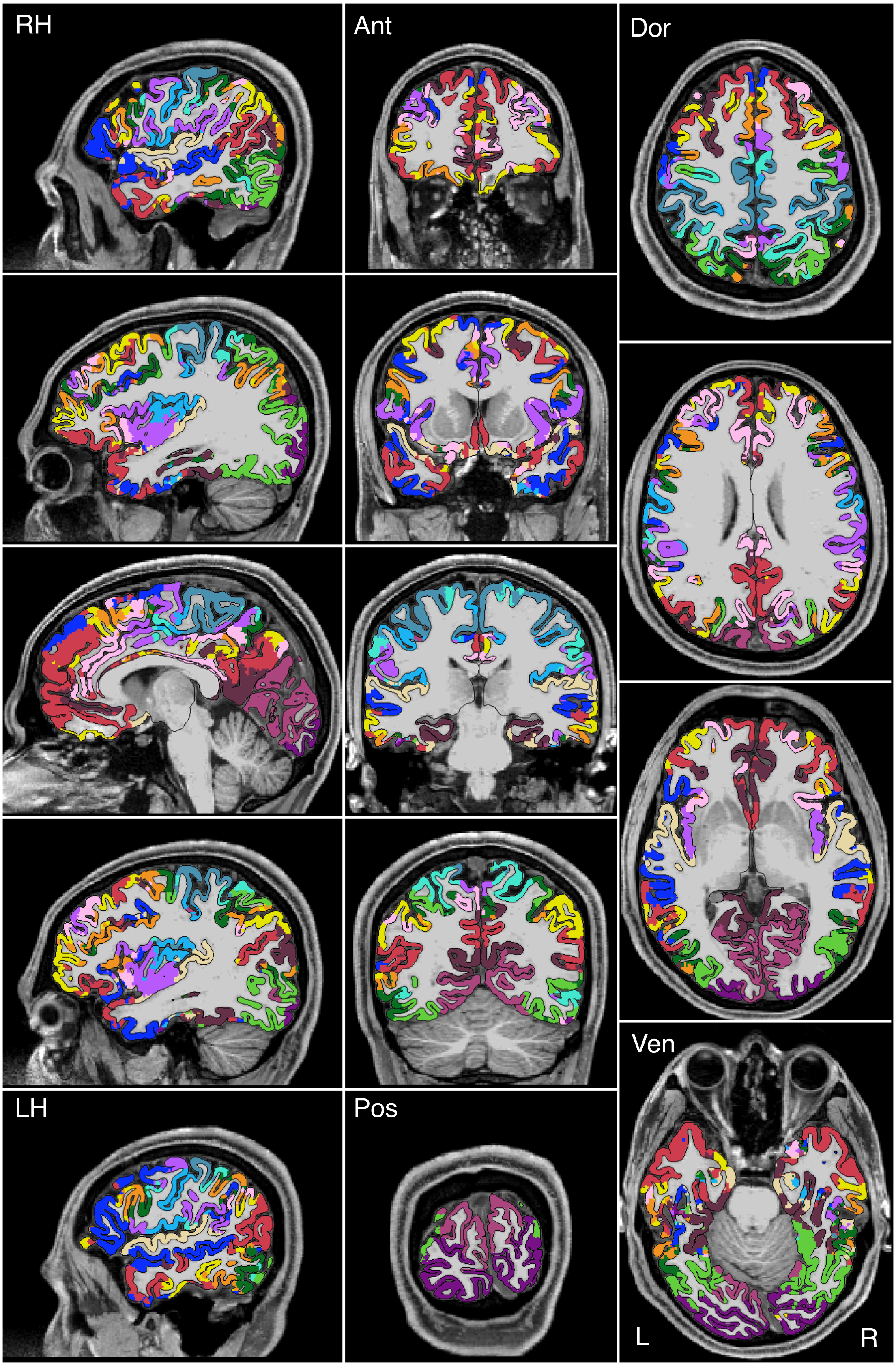


Figure S78

P10

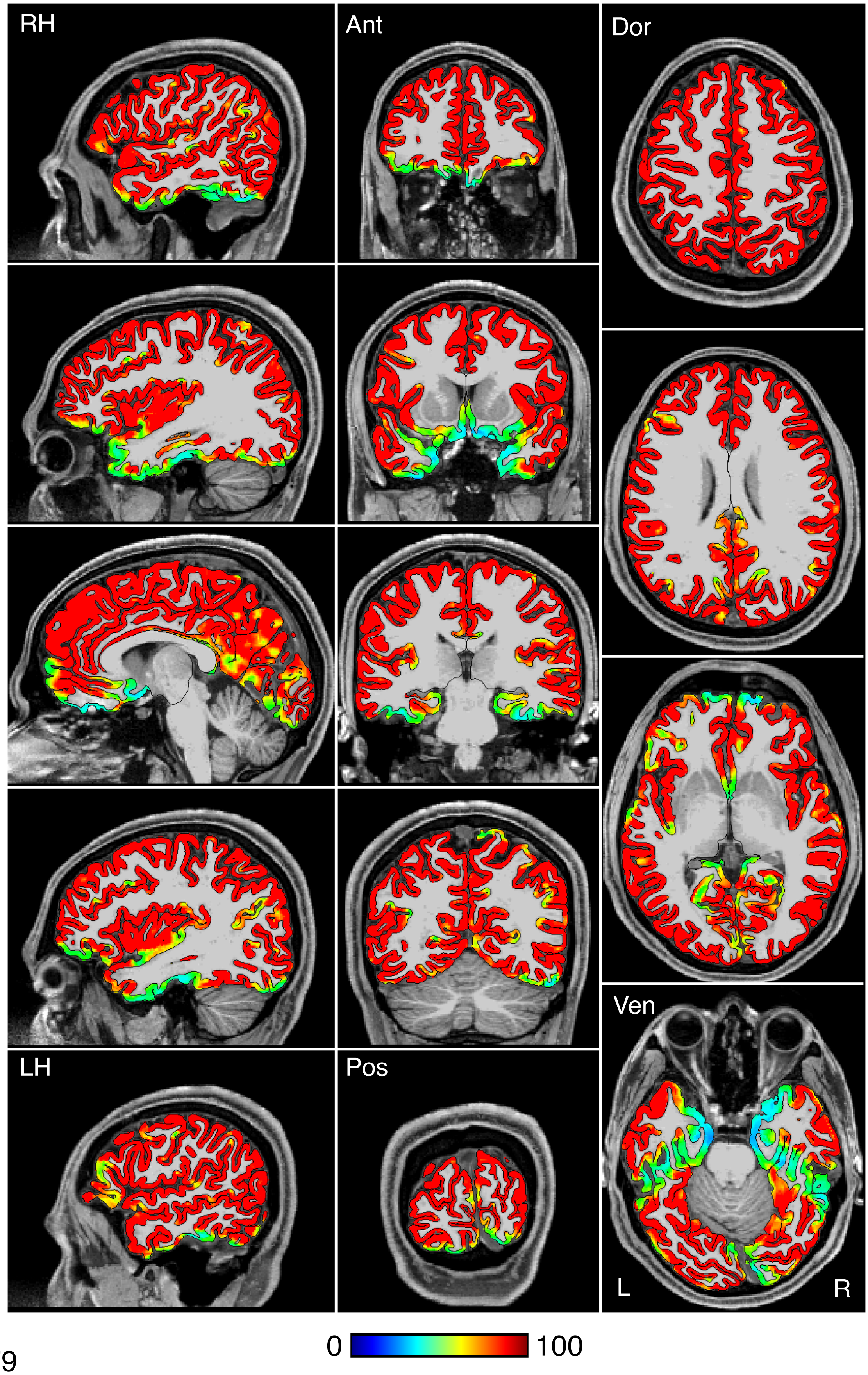


Figure S79

P10

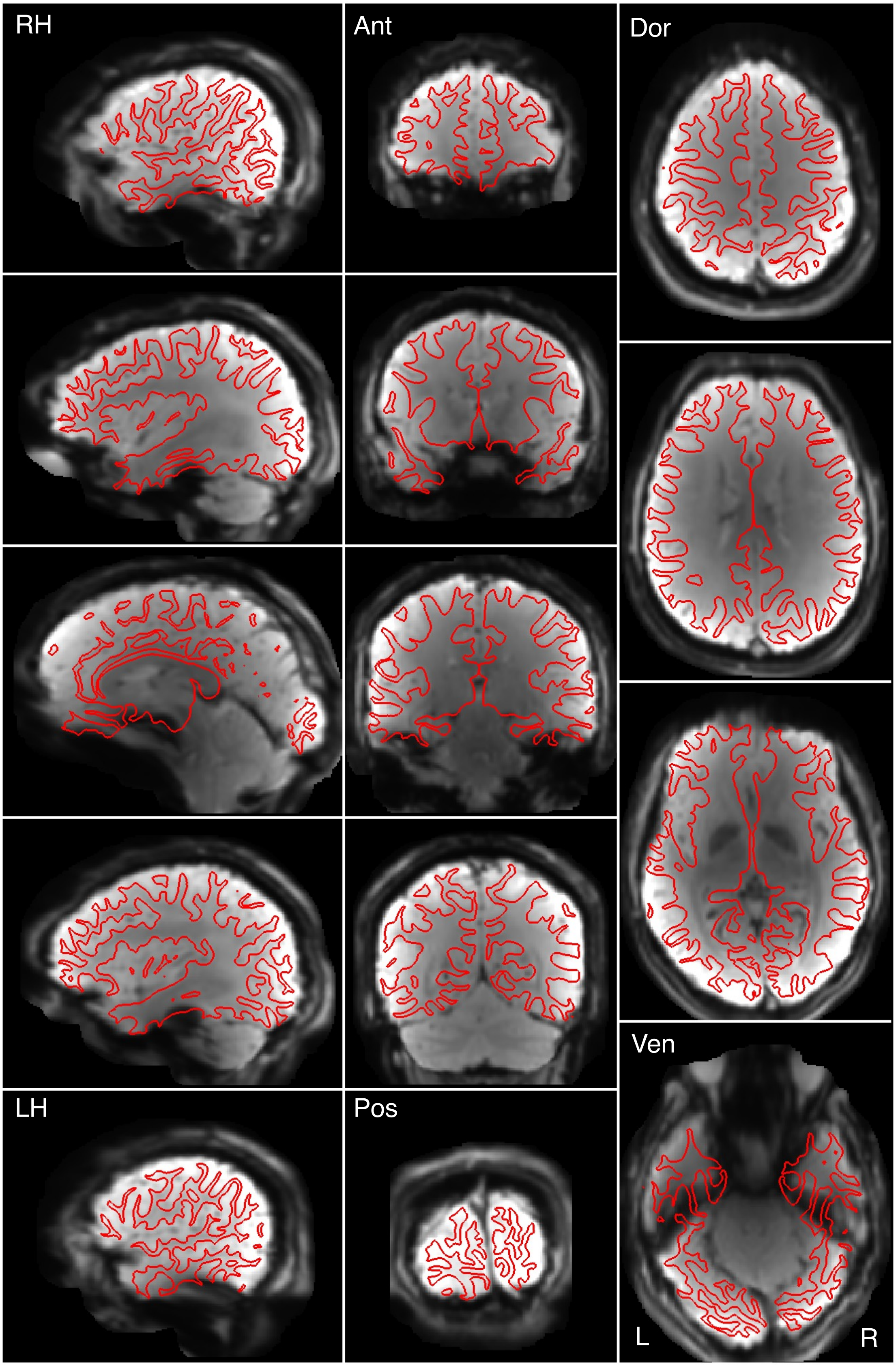


Figure S80

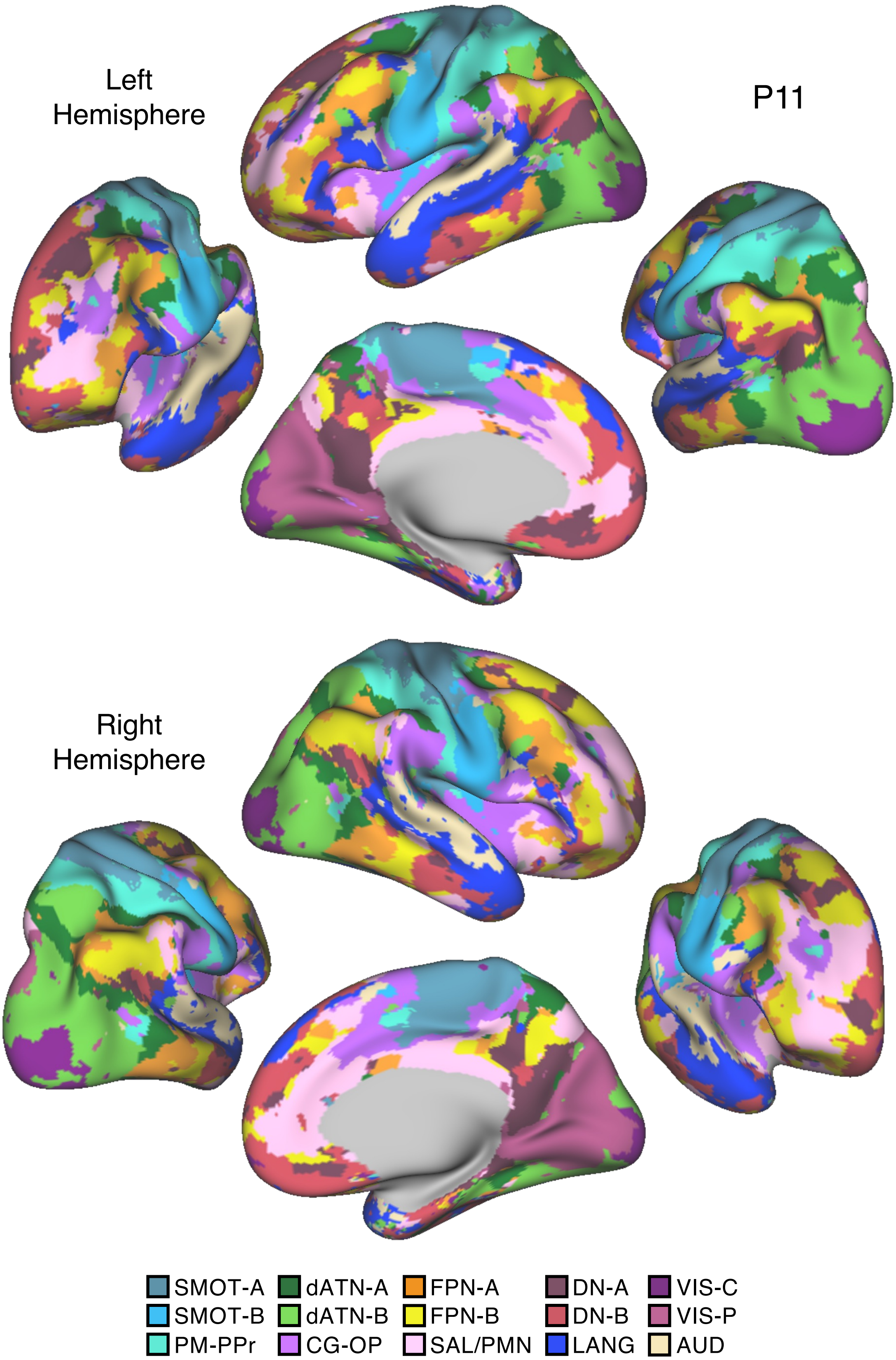

Left
Hemisphere
P11
Right
Hemisphere
SMOT-A
SMOT-B
PM-PPr
dATN-A
dATN-B
CG-OP
FPN-A
FPN-B
SAL/PMN
DN-A
DN-B
LANG
VIS-C
VIS-P
AUD


Figure S81

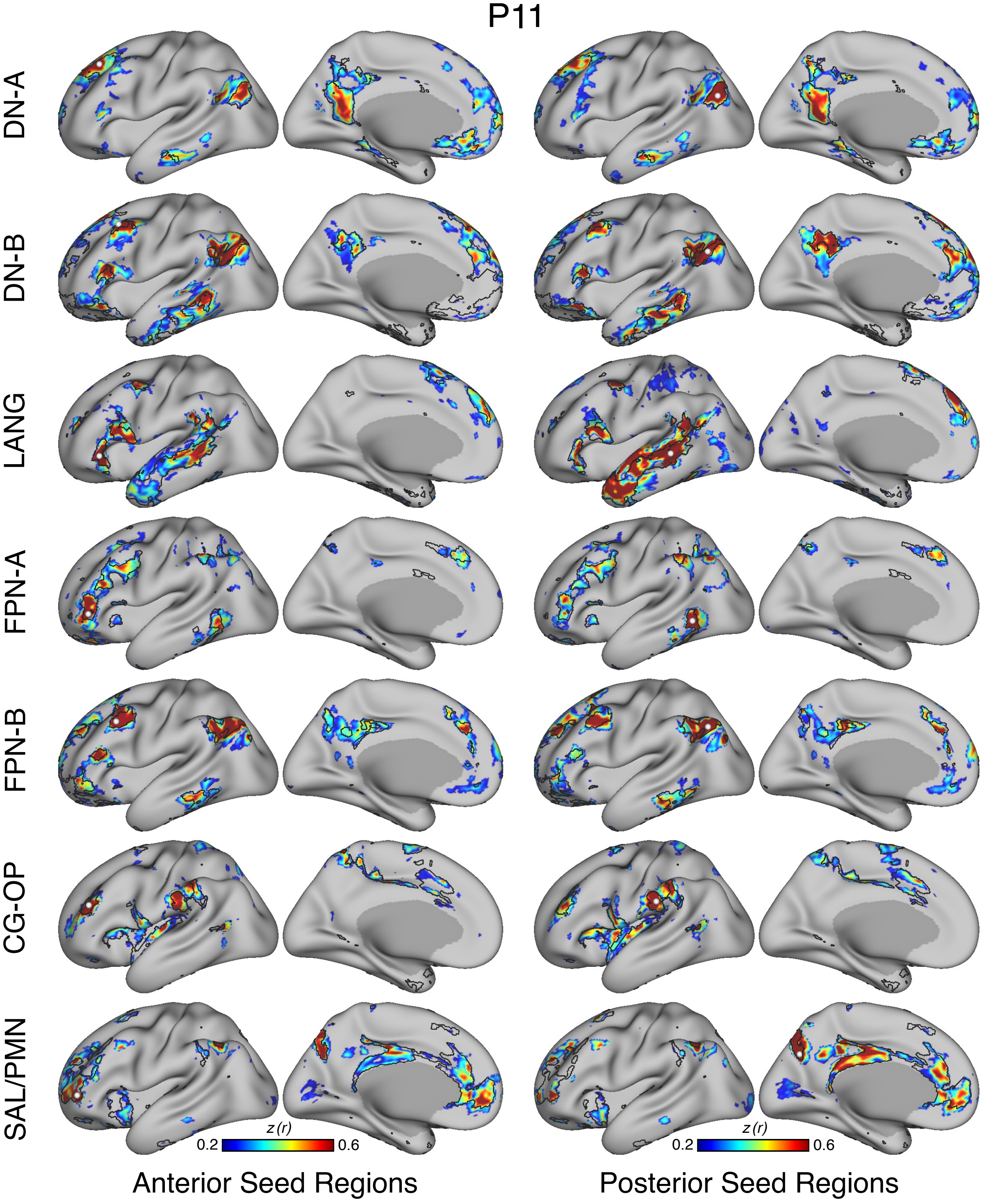
P11
DN-A
DN-B
LANG
FPN-A
FPN-B
CG-OP
SAL/PMN
z (r)
0.2
0.6
z (r)
0.2
0.6
Anterior Seed Regions
Posterior Seed Regions

Figure S82

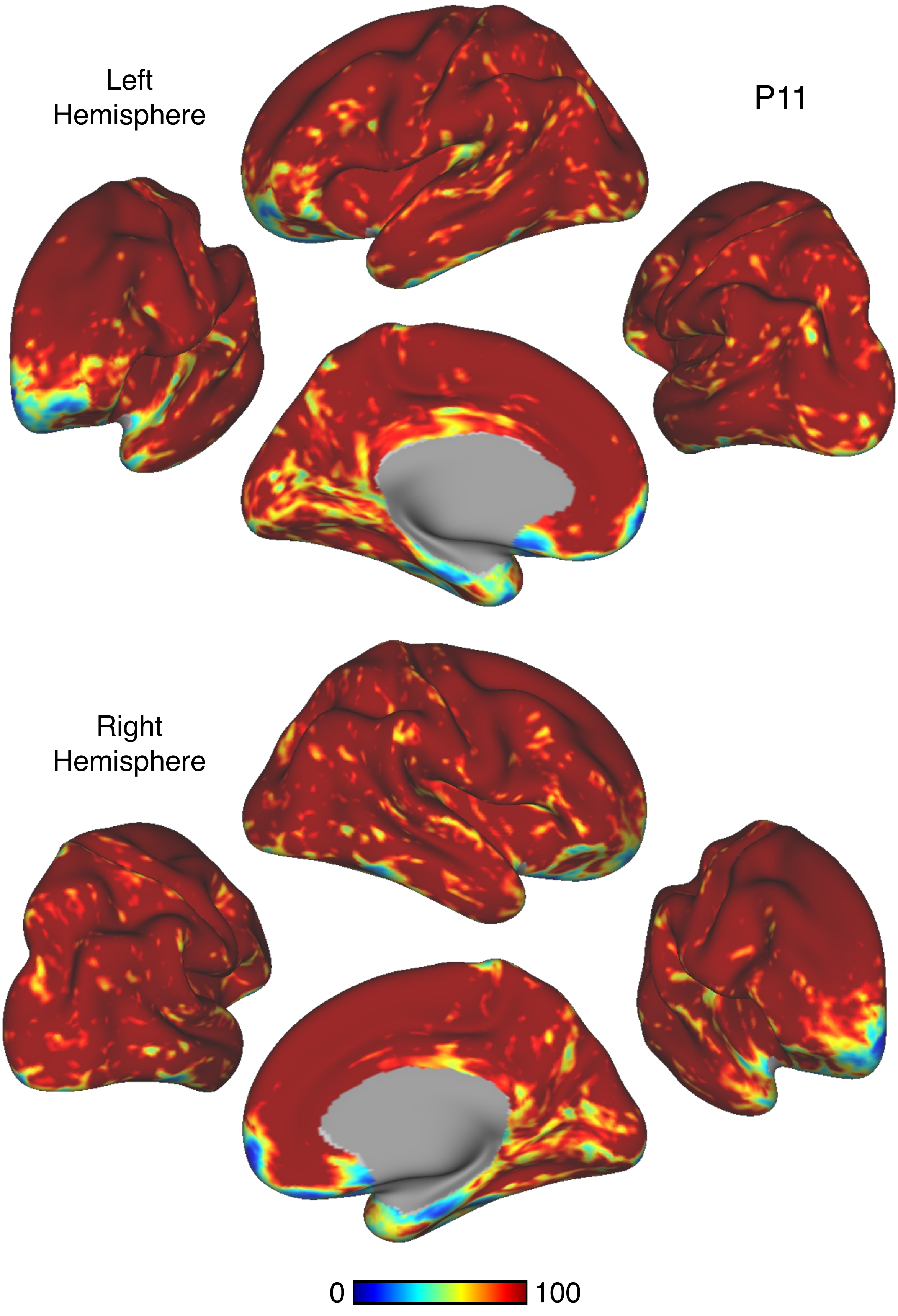


Figure S83

P11

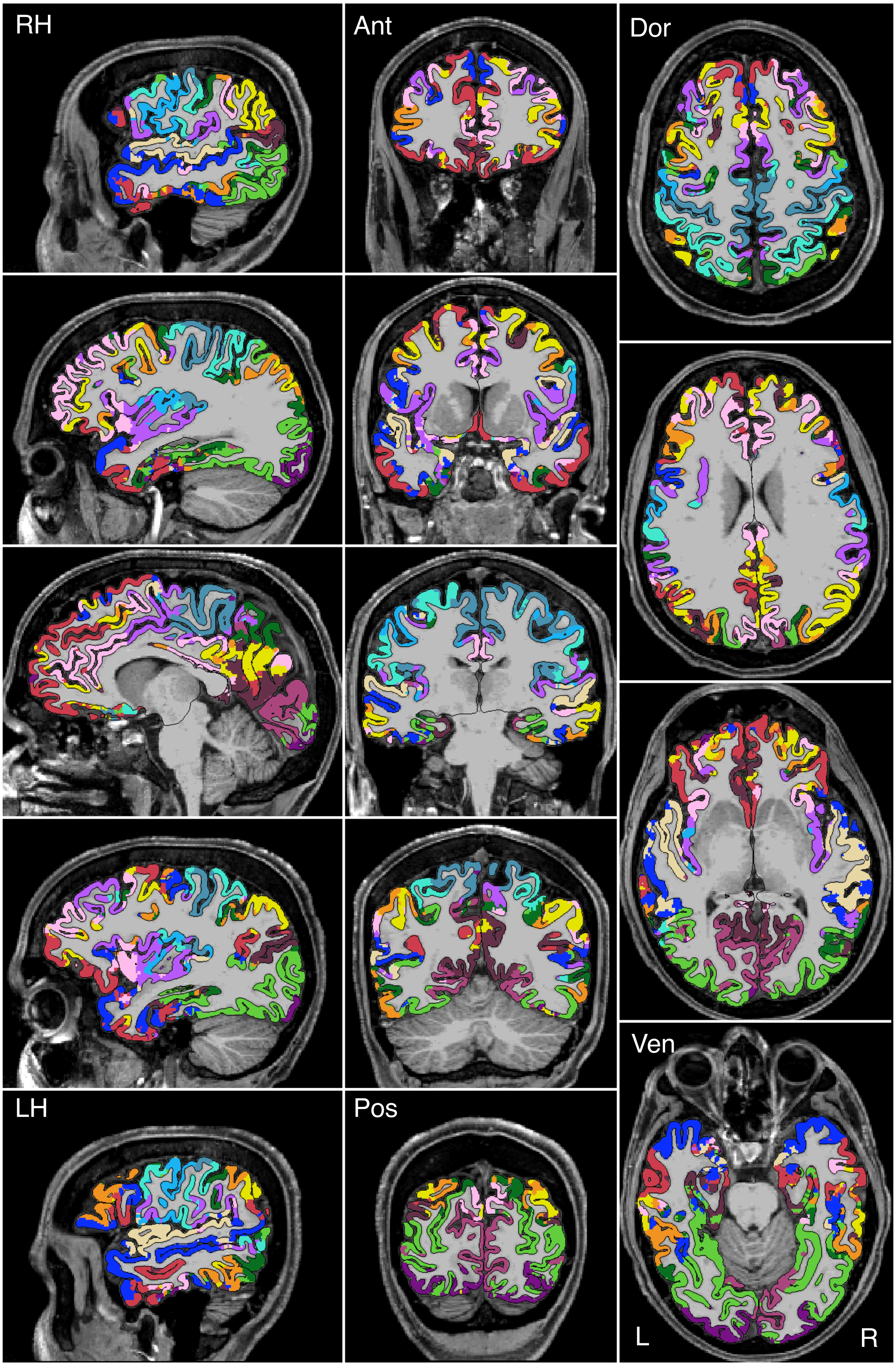


Figure S84

P11

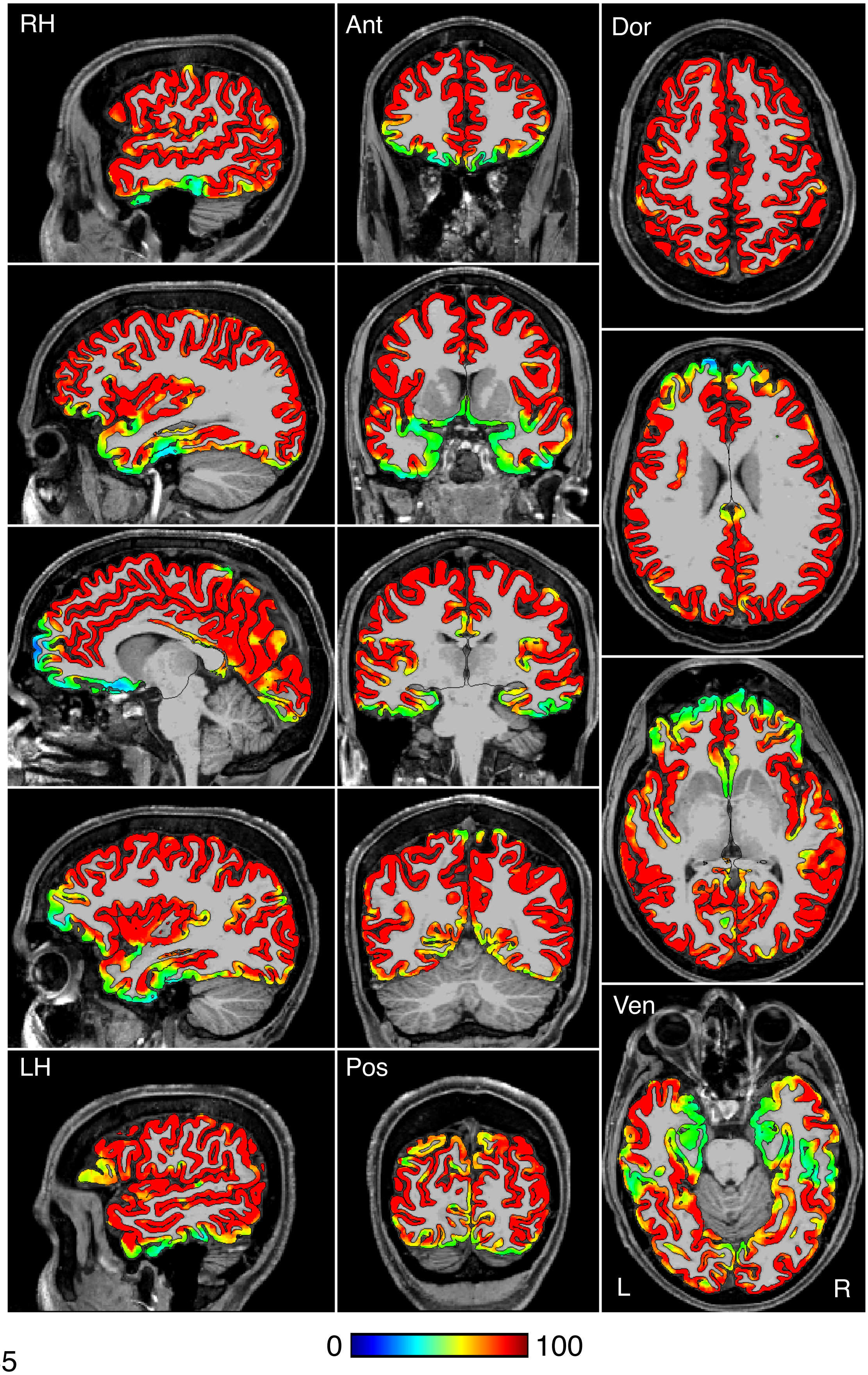


Figure S85

P11

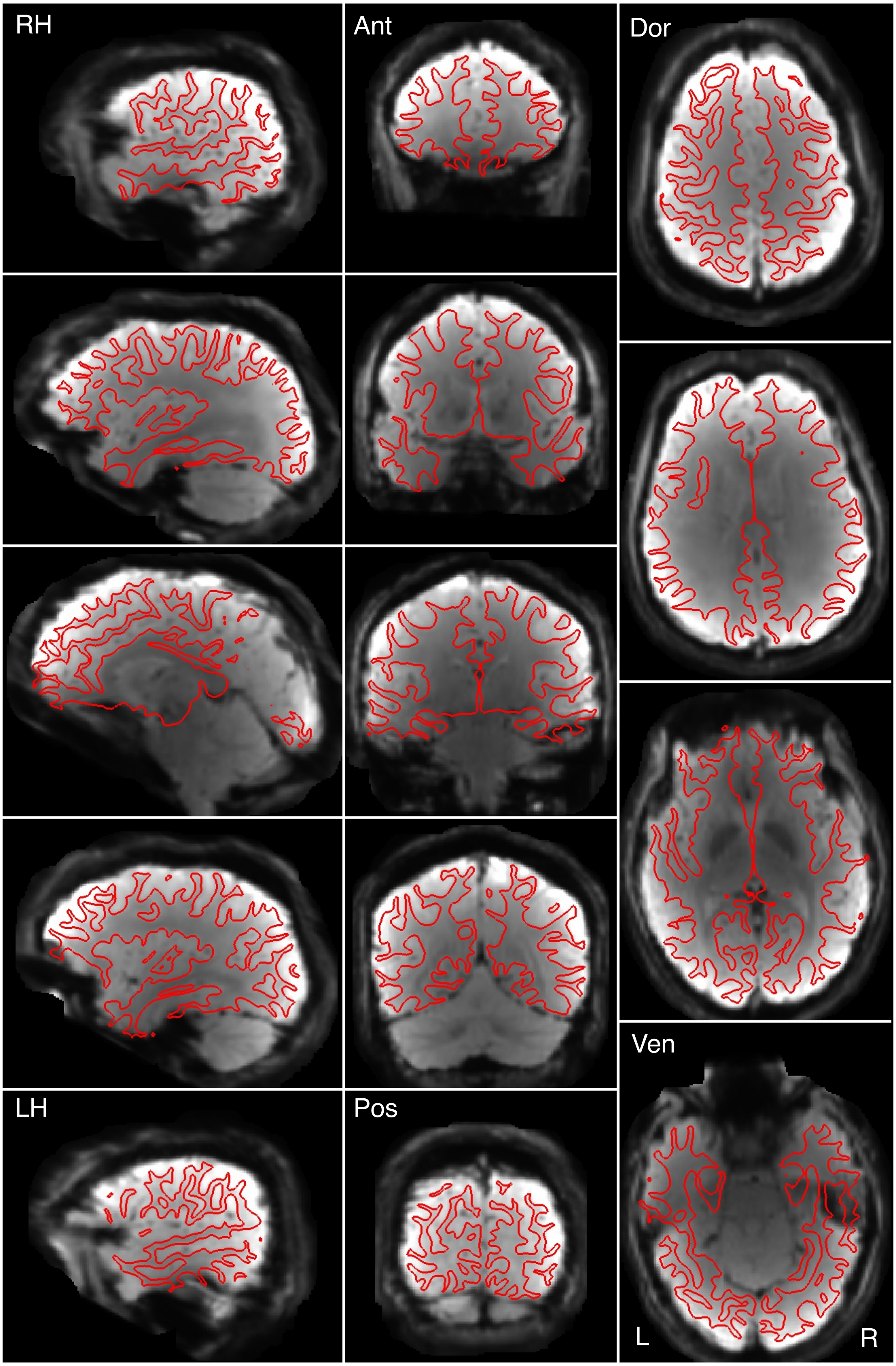


Figure S86

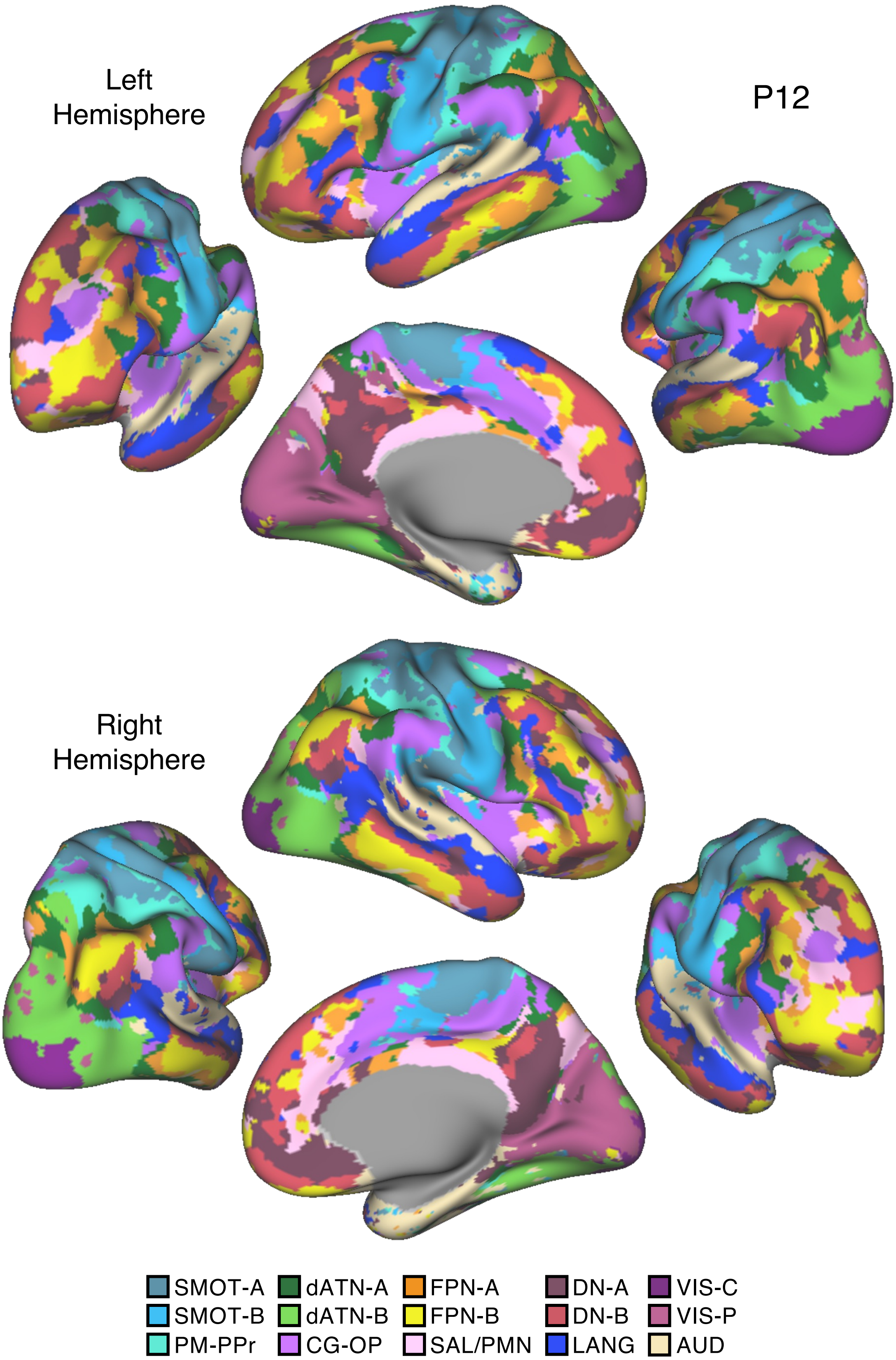
Left
Hemisphere
P12
Right
Hemisphere
SMOT-A
SMOT-B
PM-PPr
dATN-A
dATN-B
CG-OP
FPN-A
FPN-B
SAL/PMN
DN-A
DN-B
LANG
VIS-C
VIS-P
AUD

Figure S87

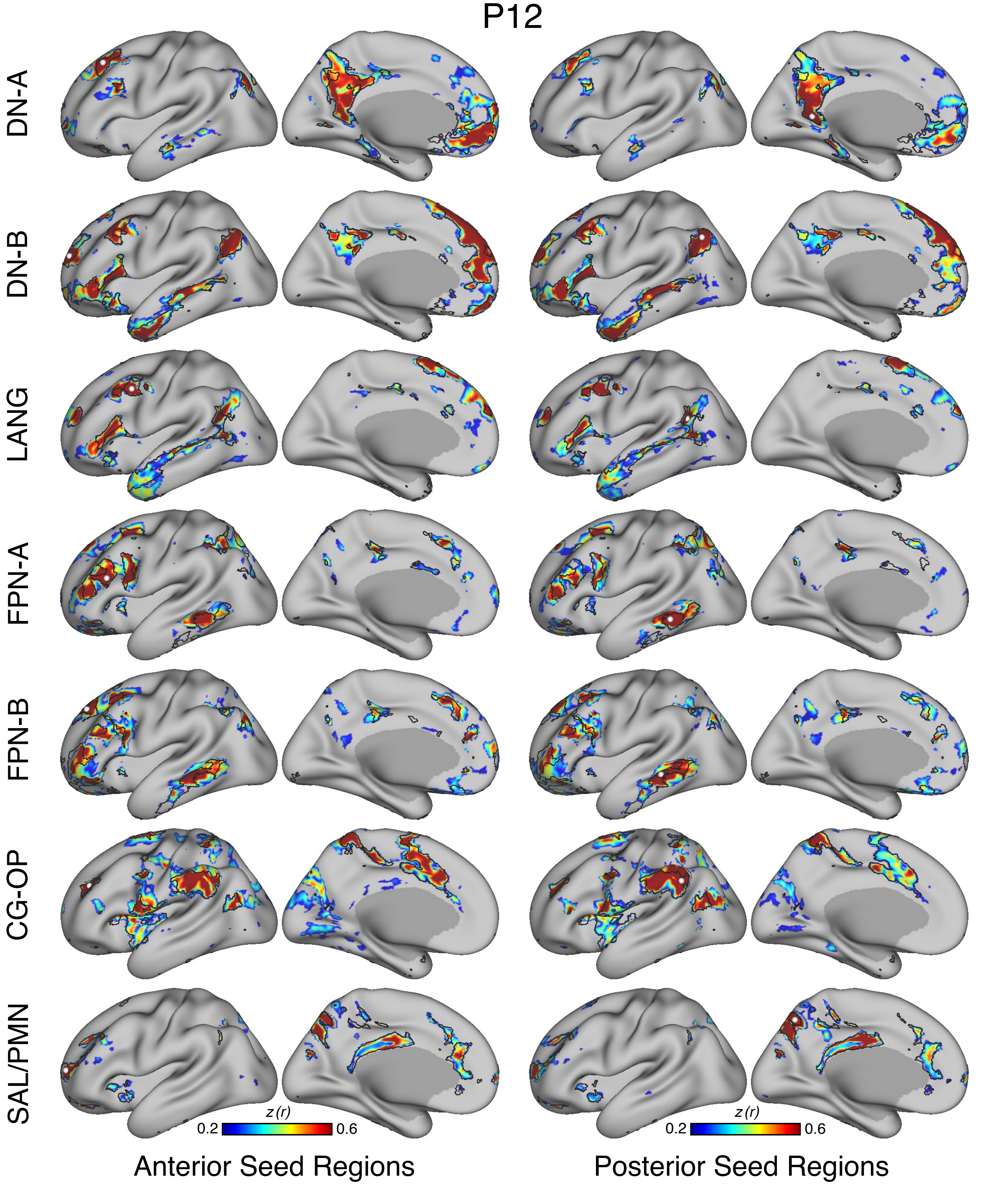
P12
DN-A
DN-B
LANG
FPN-A
FPN-B
CG-OP
SAL/PMN
z (r)
0.2
0.6
z (r)
0.2
0.6
Anterior Seed Regions
Posterior Seed Regions

Figure S88

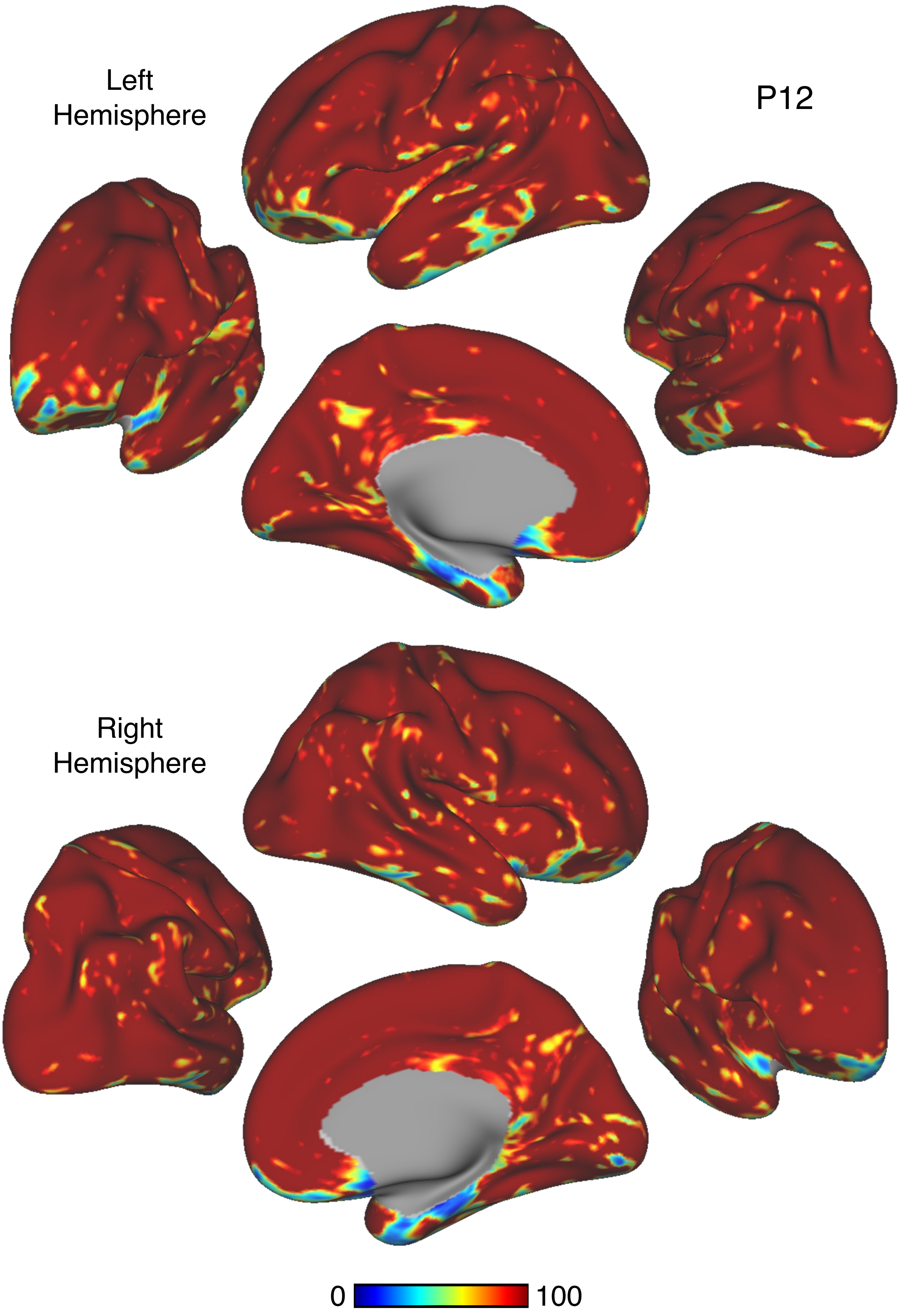


Figure S89

P12

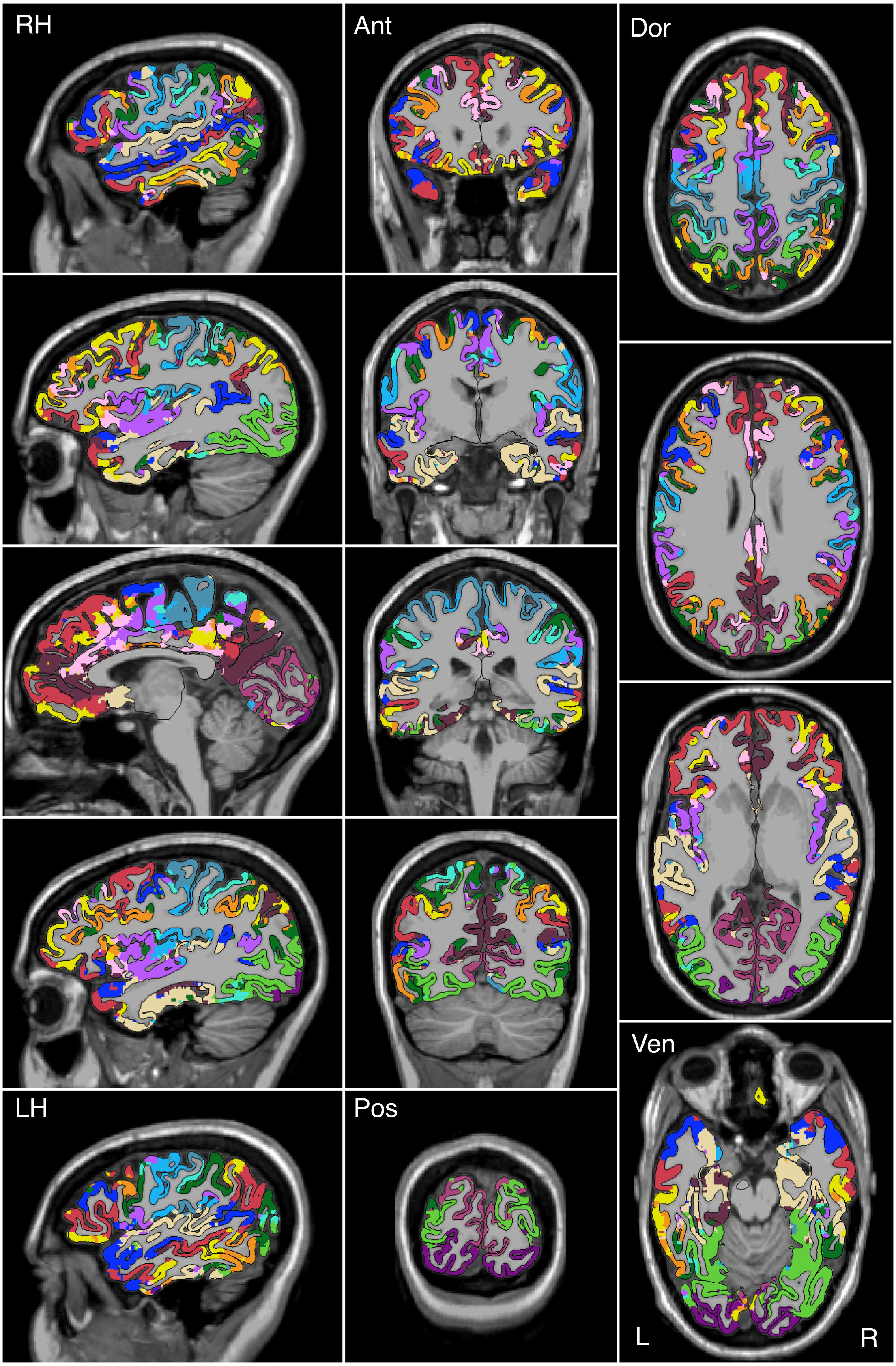


Figure S90

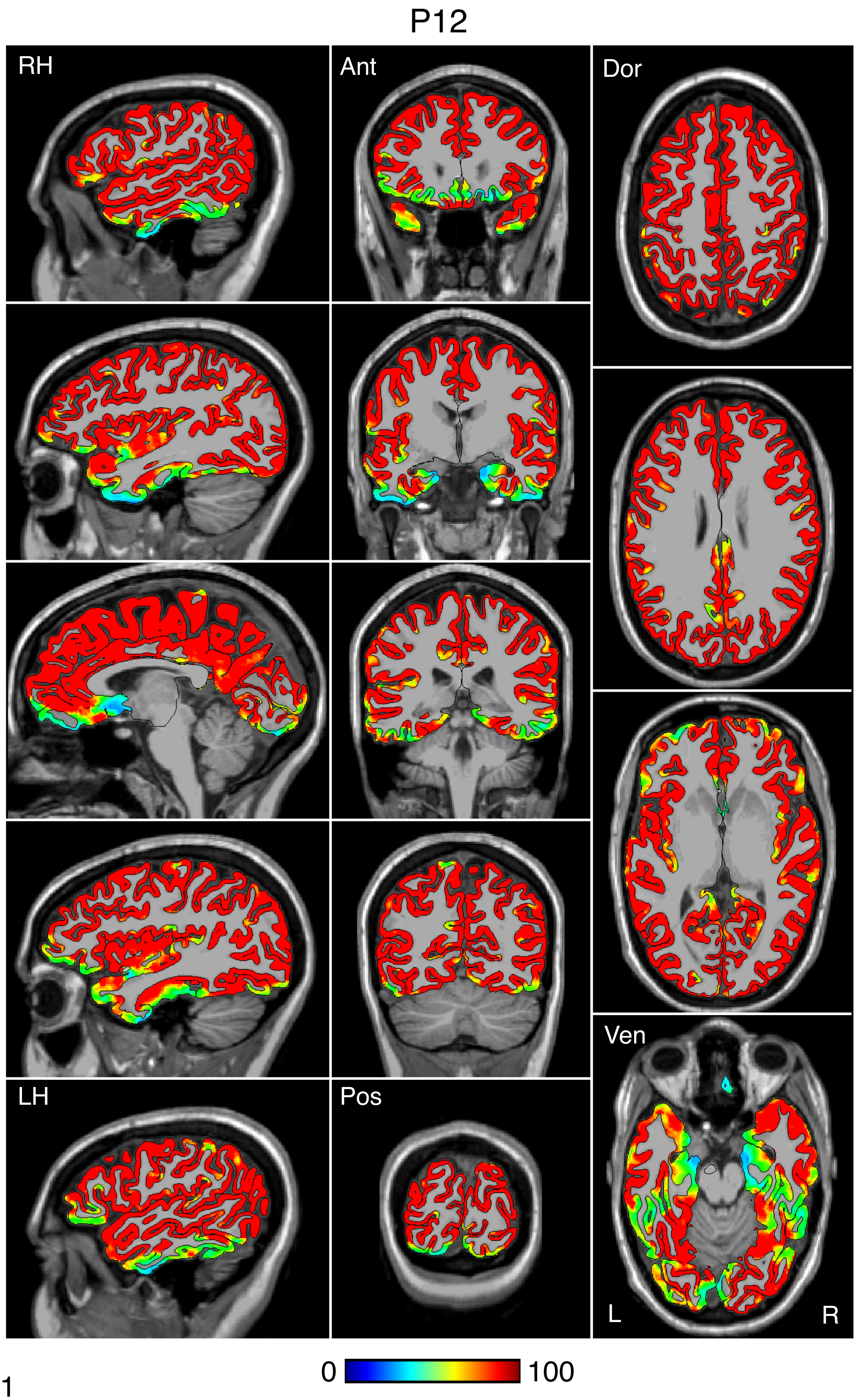


Figure S91

P12

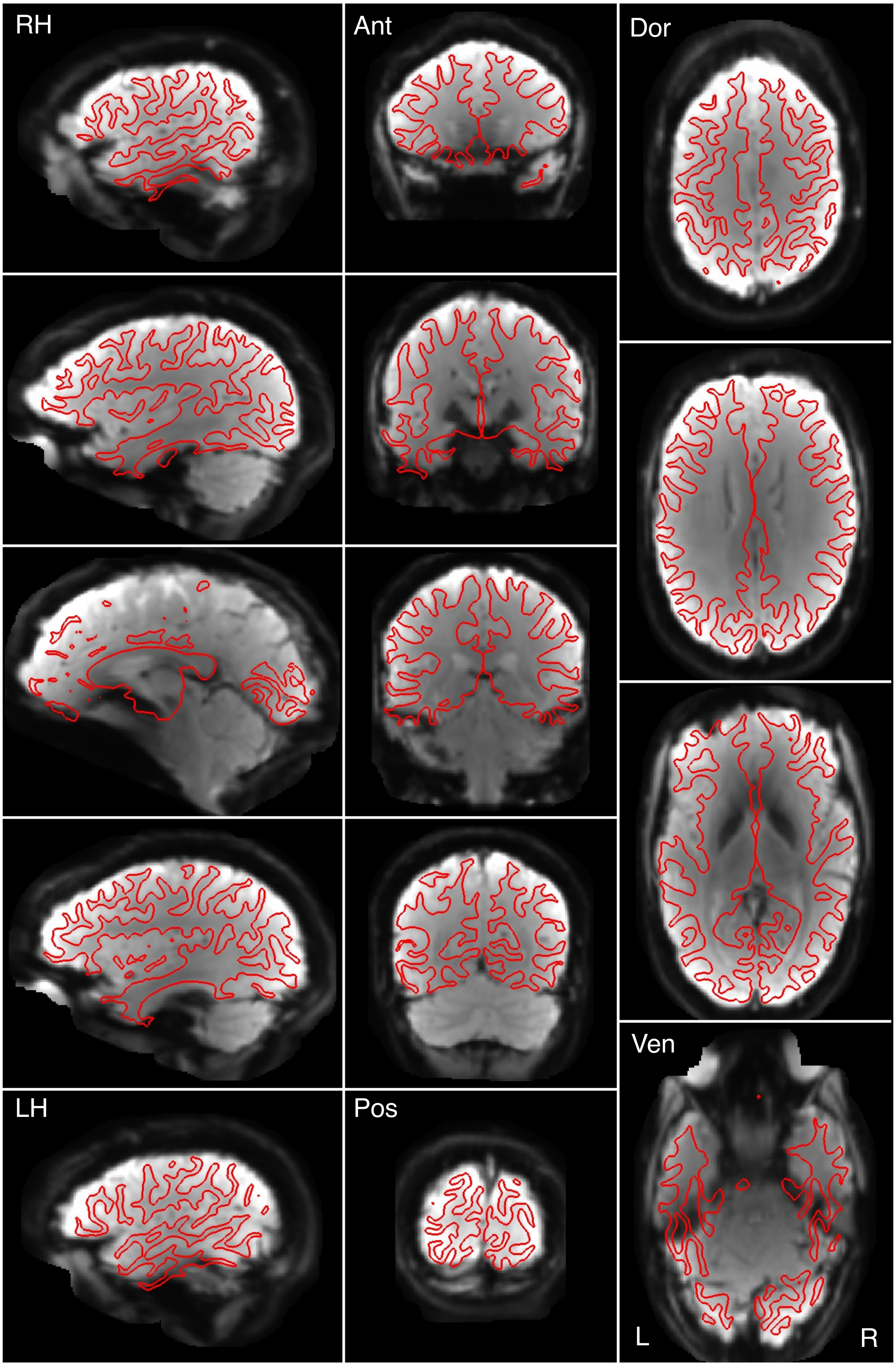


Figure S92

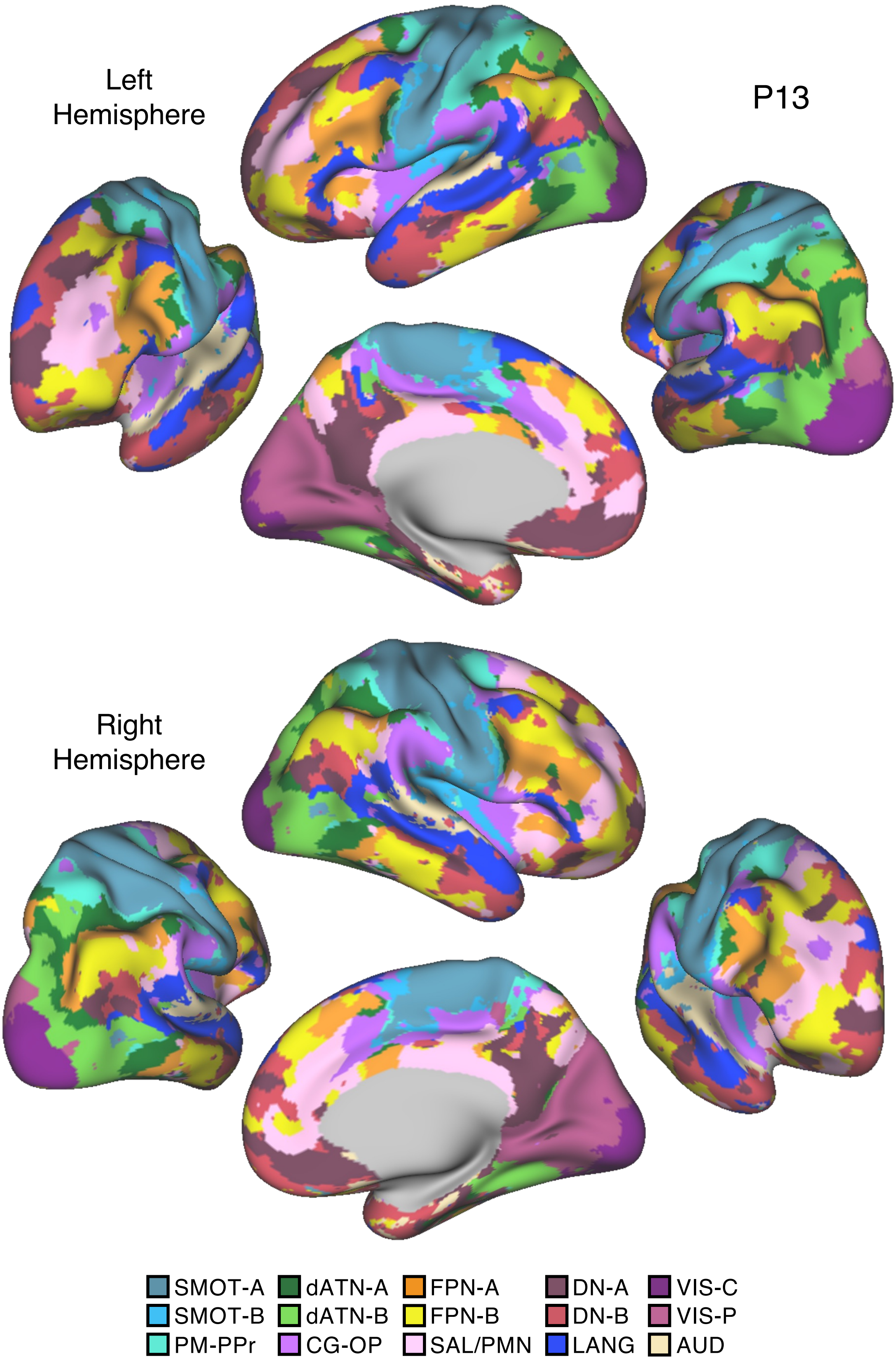

Left
Hemisphere
P13
Right
Hemisphere
SMOT-A
SMOT-B
PM-PPr
dATN-A
dATN-B
CG-OP
FPN-A
FPN-B
SAL/PMN
DN-A
DN-B
LANG
VIS-C
VIS-P
AUD


Figure S93

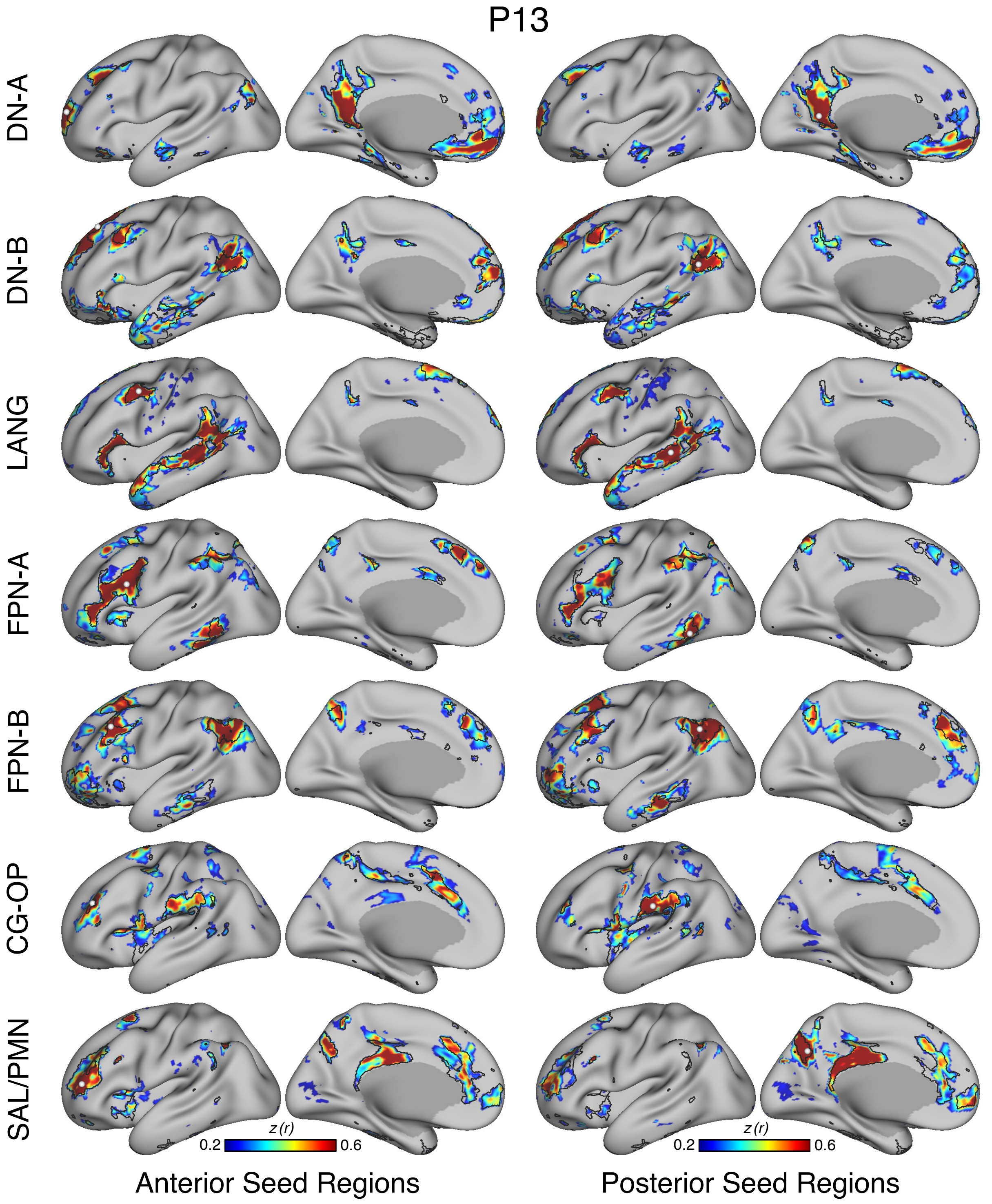
P13
DN-A
DN-B
LANG
FPN-A
FPN-B
CG-OP
SAL/PMN
z (r)
0.2
0.6
z (r)
0.2
0.6
Anterior Seed Regions
Posterior Seed Regions

Figure S94

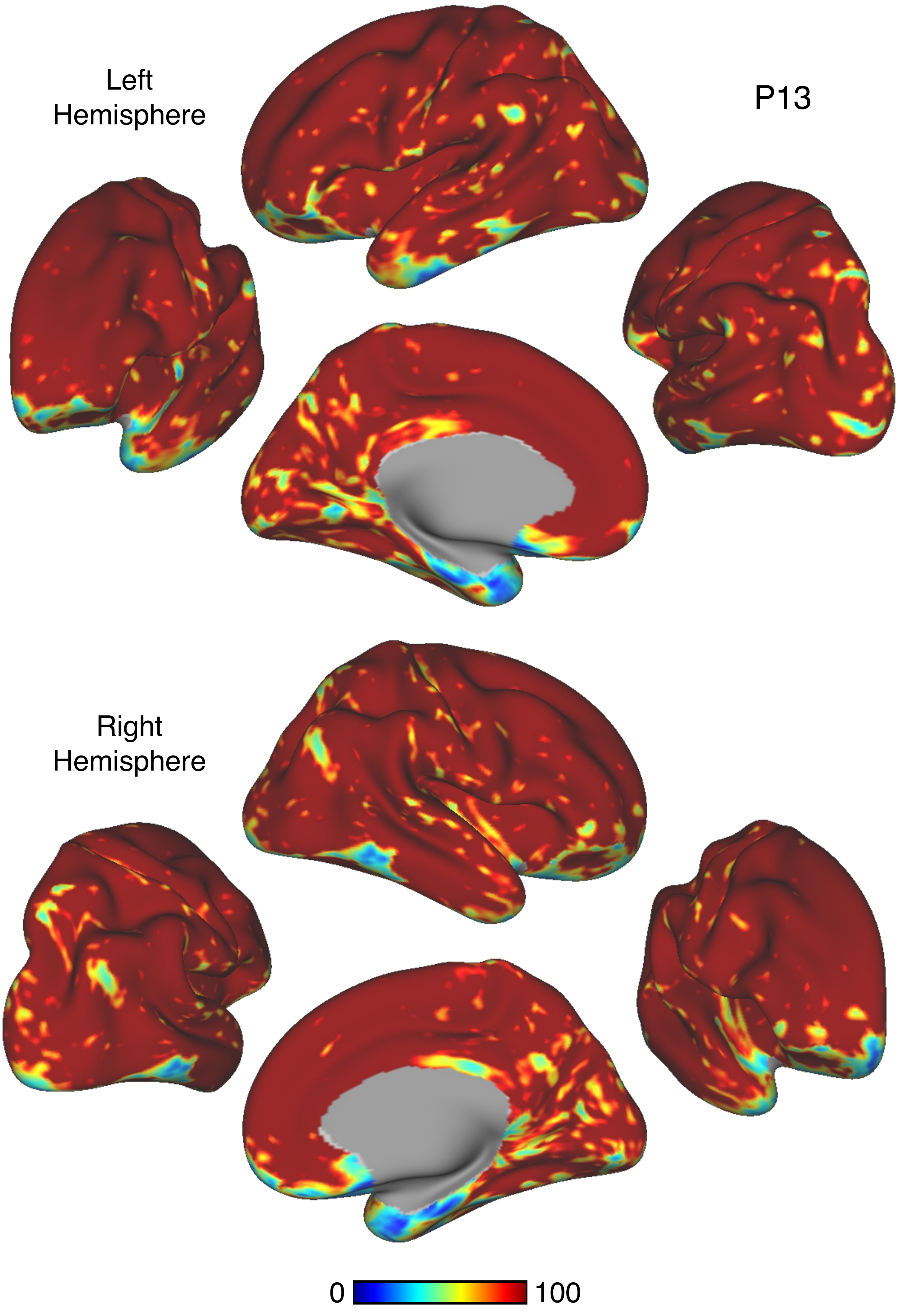


Figure S95

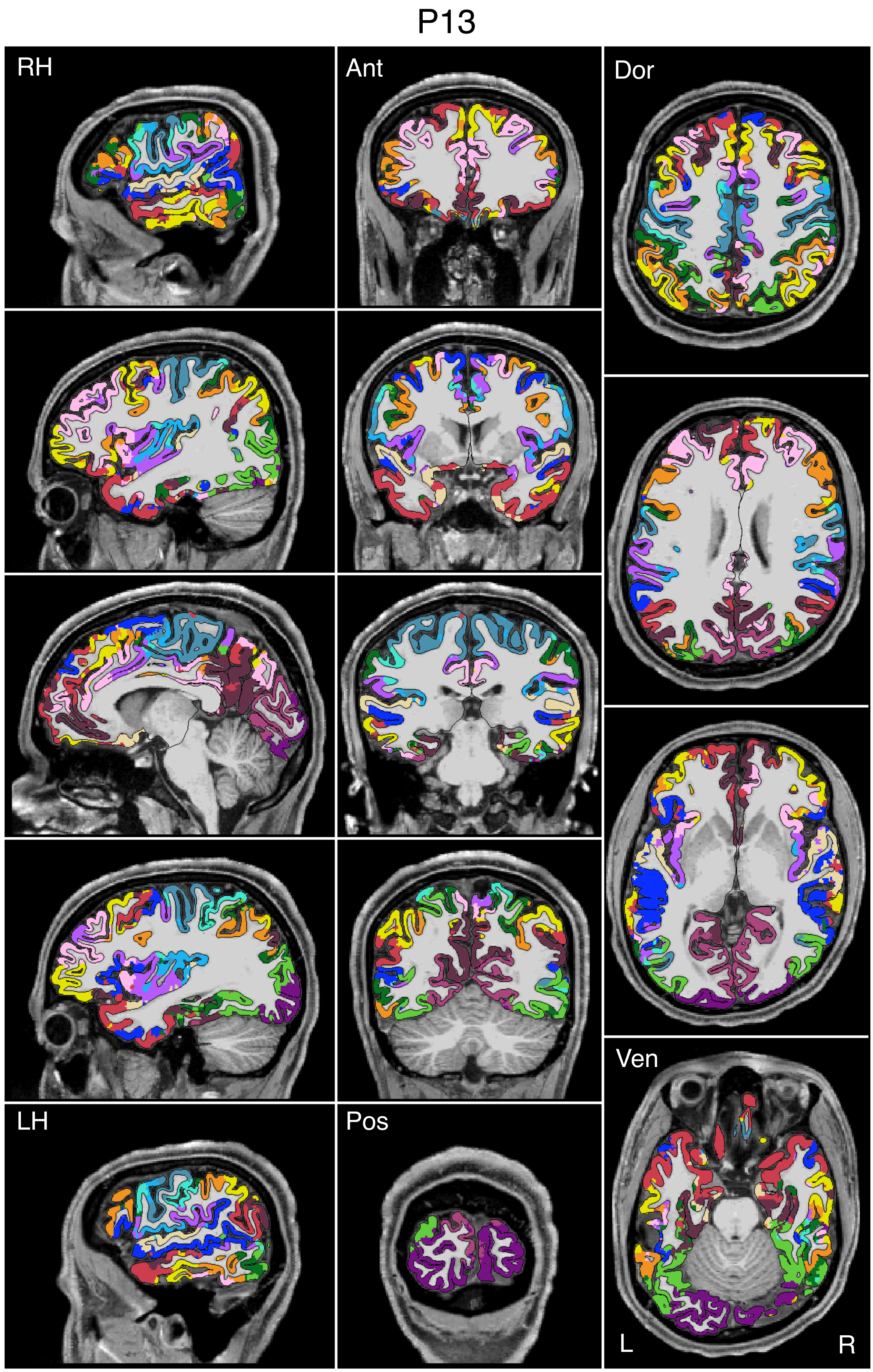


Figure S96

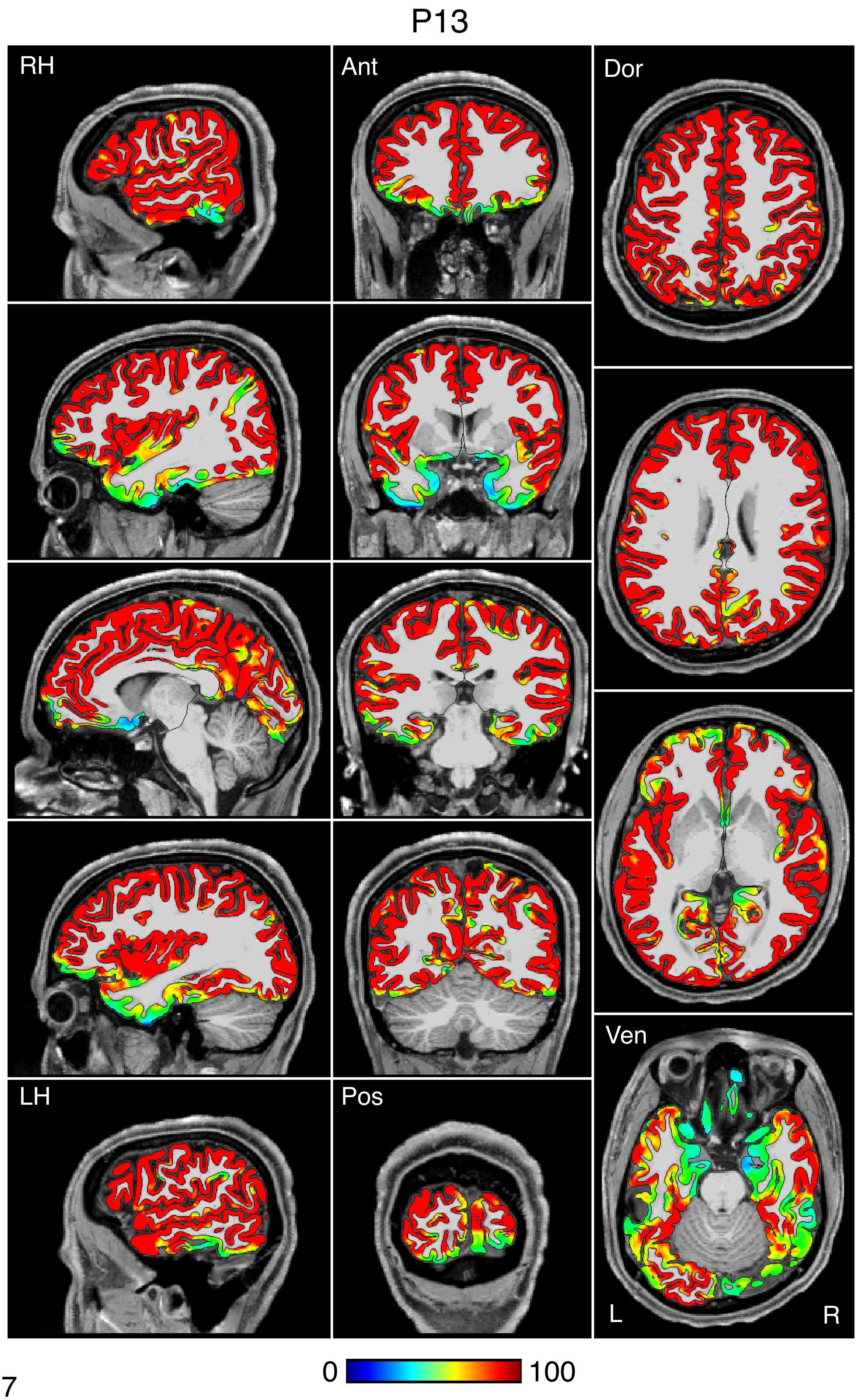


Figure S97

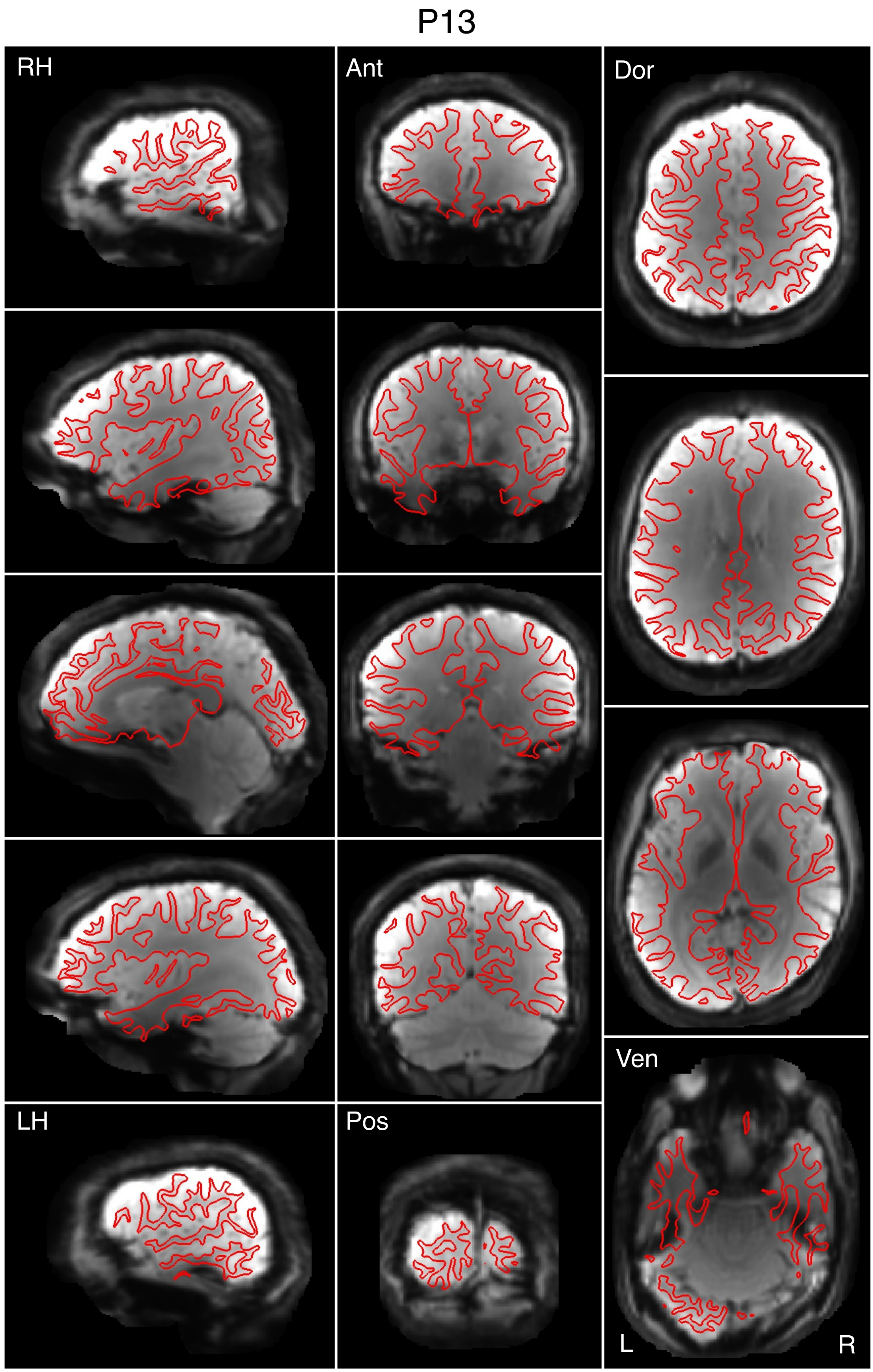


Figure S98

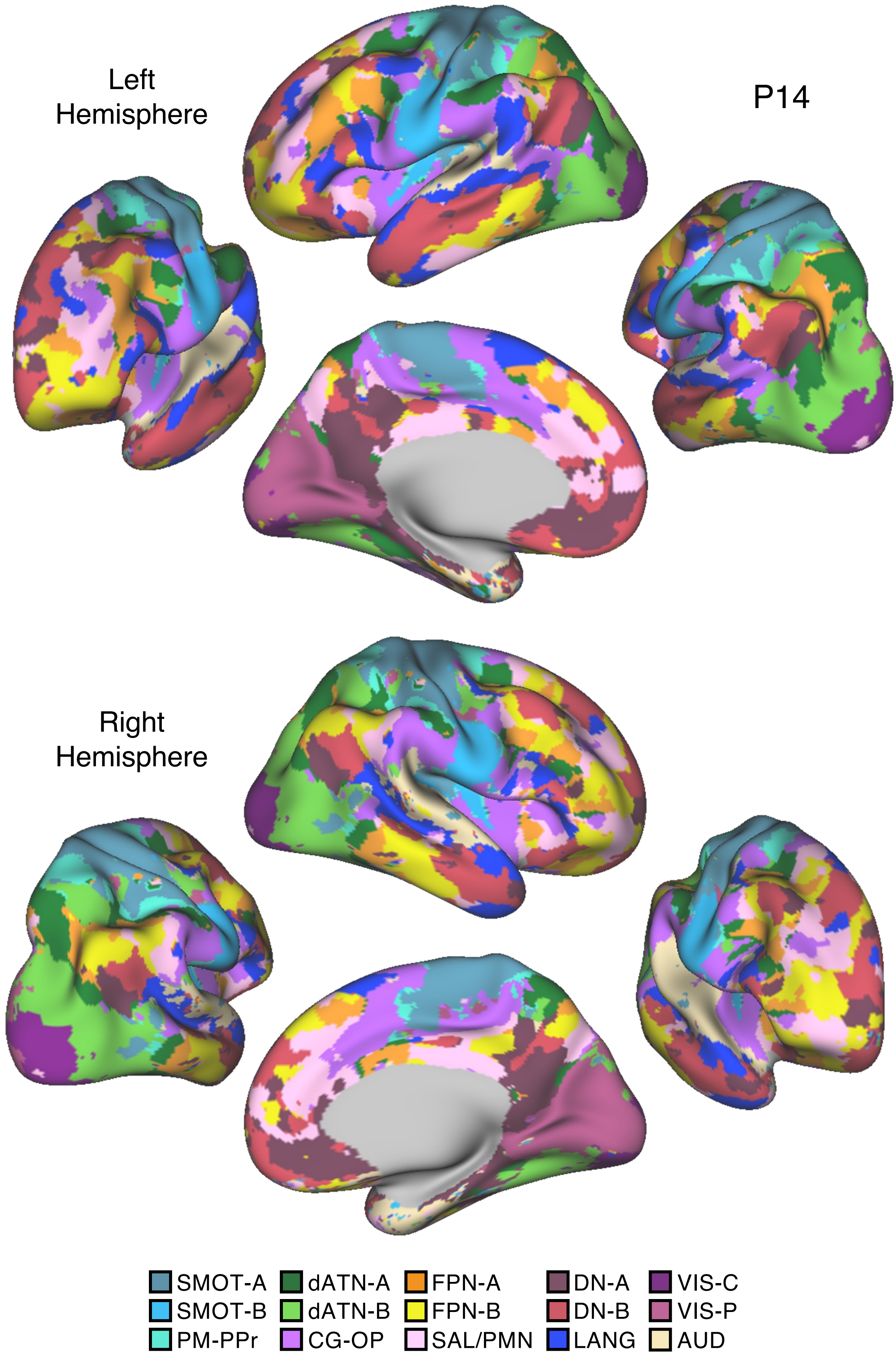
Left
Hemisphere
P14
Right
Hemisphere
SMOT-A
SMOT-B
PM-PPr
dATN-A
dATN-B
CG-OP
FPN-A
FPN-B
SAL/PMN
DN-A
DN-B
LANG
VIS-C
VIS-P
AUD

Figure S99

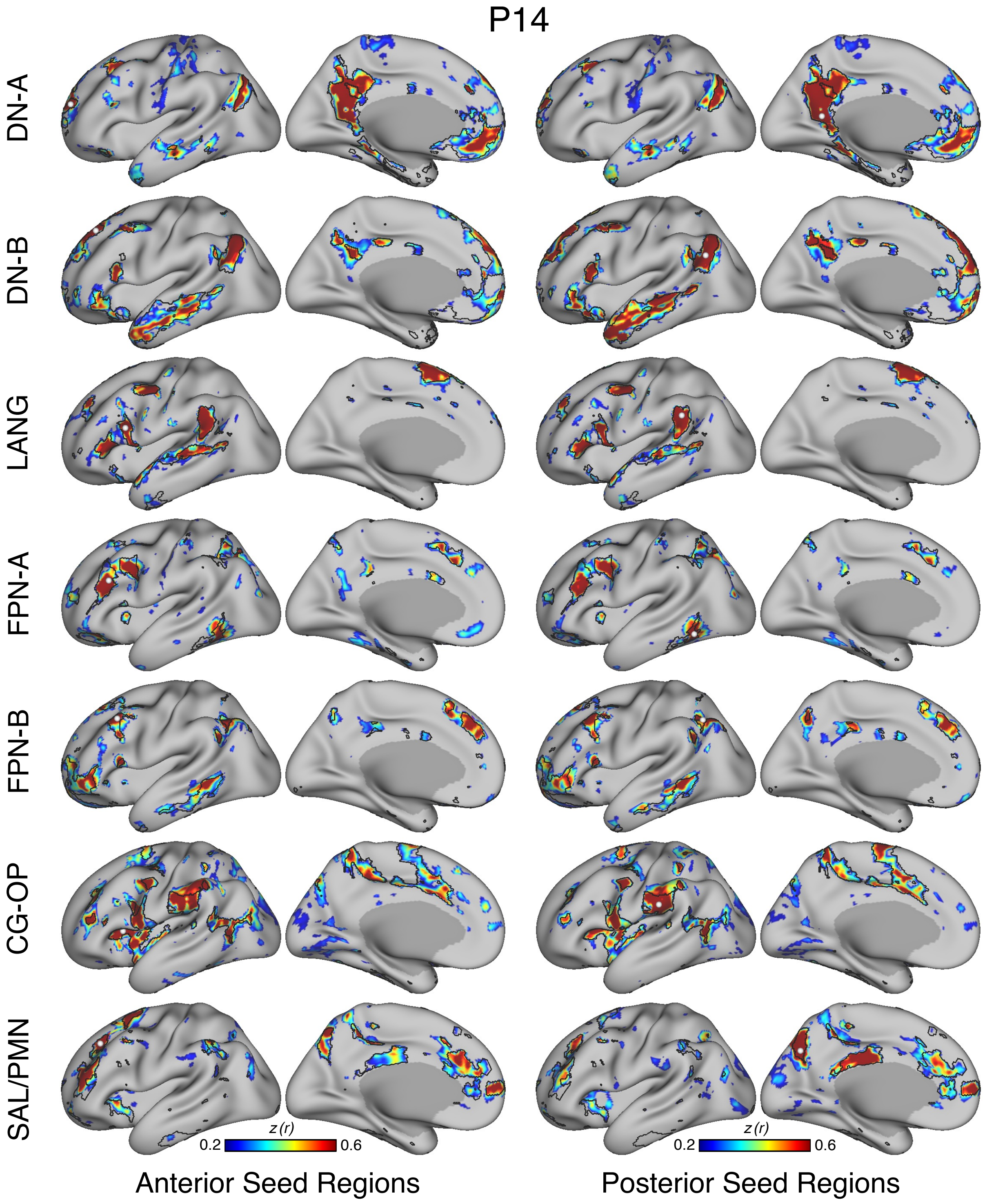

P14
DN-A
DN-B
LANG
FPN-A
FPN-B
CG-OP
SAL/PMN
z (r)
0.2
0.6
z (r)
0.2
0.6
Anterior Seed Regions
Posterior Seed Regions


Figure S100

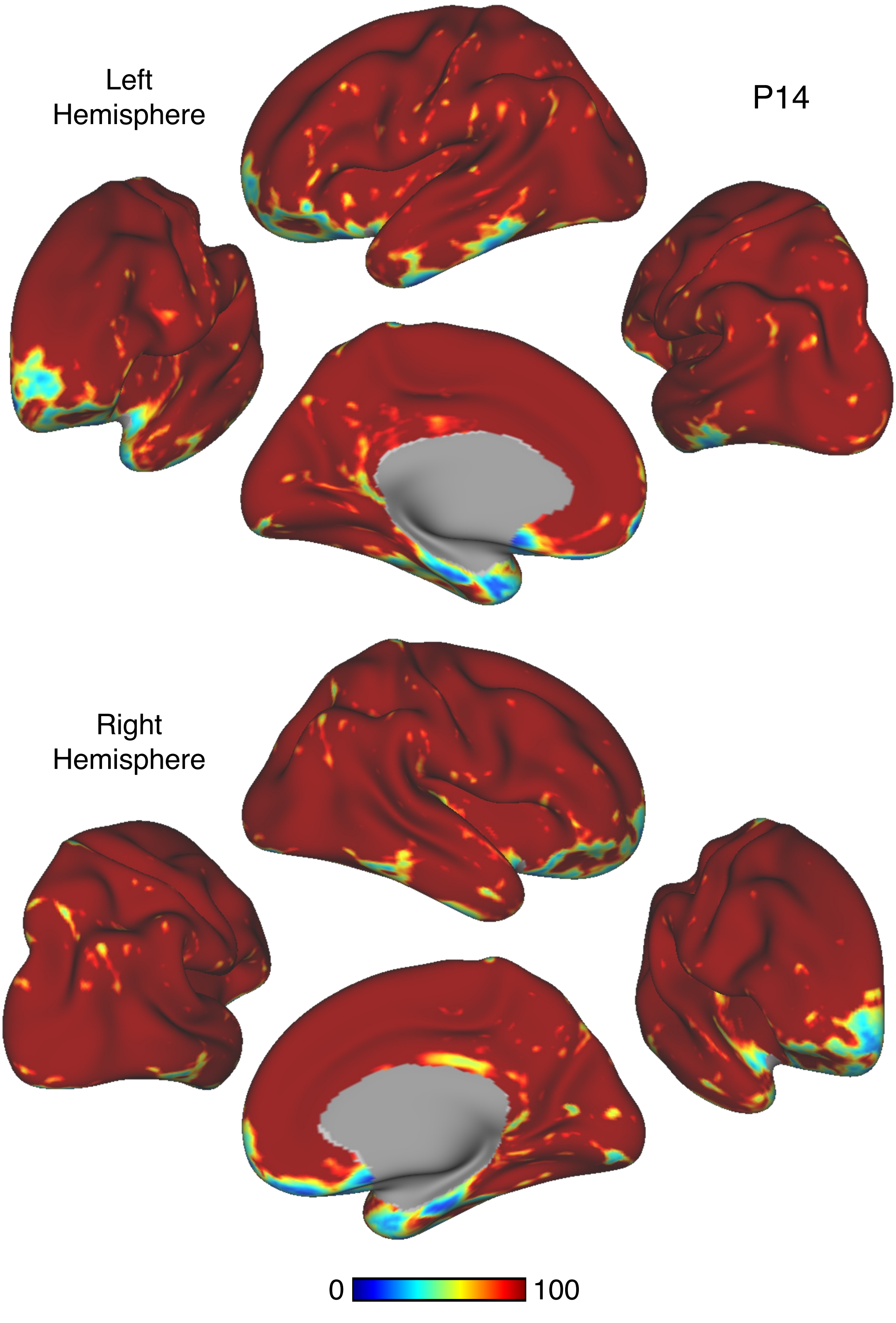


Figure S101

# P14

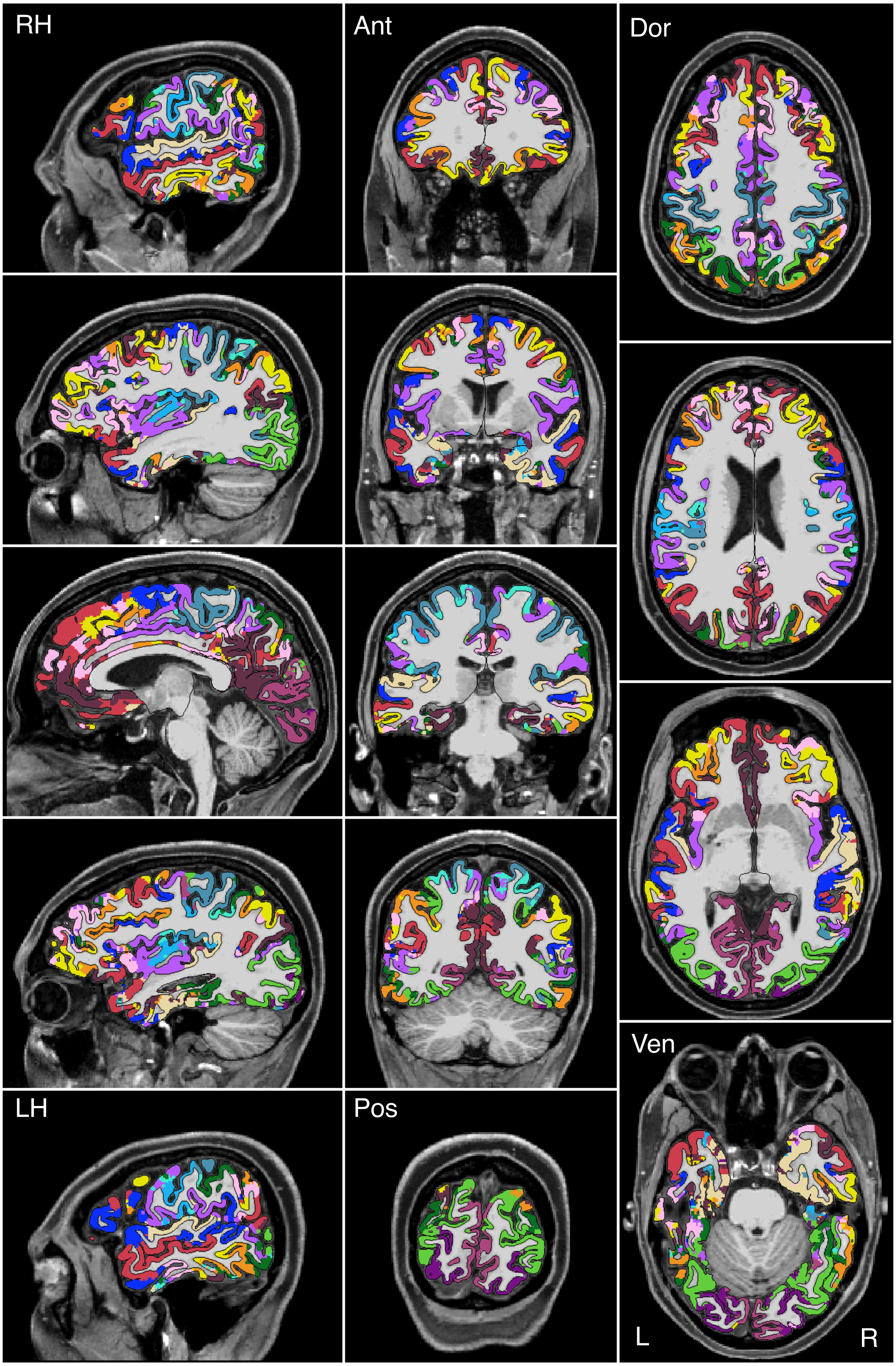


Figure S102

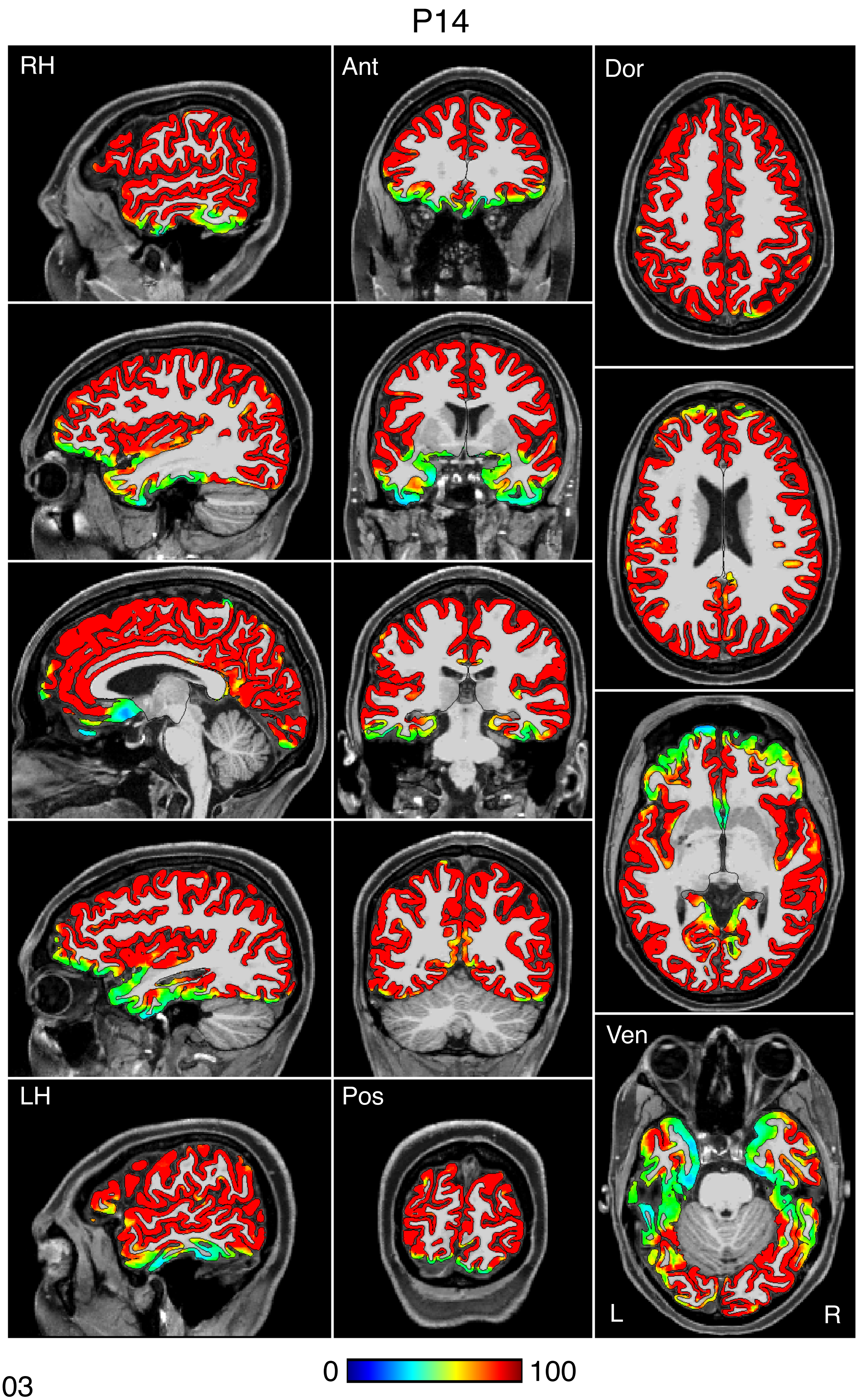


Figure S103

P14

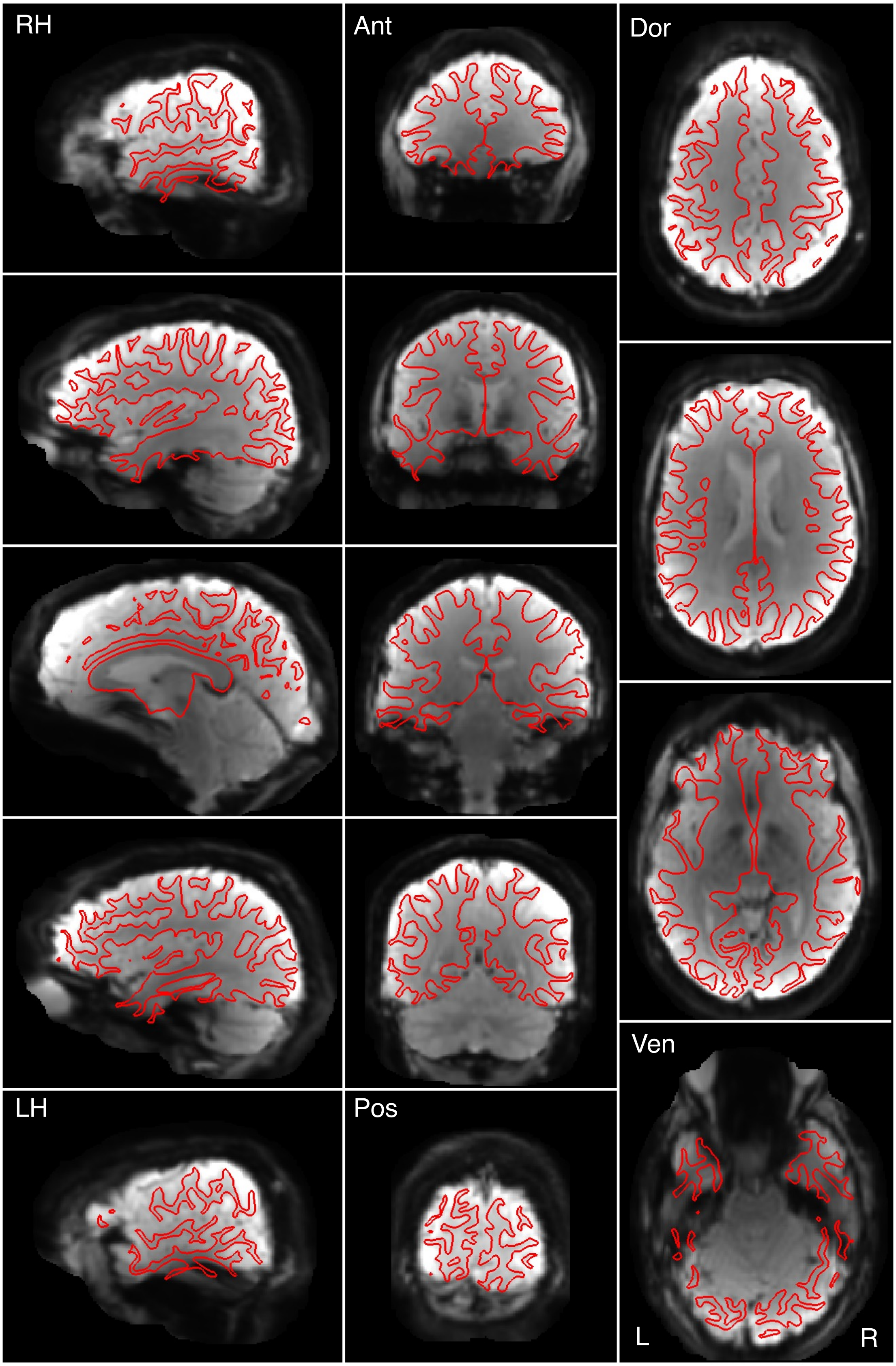


Figure S104

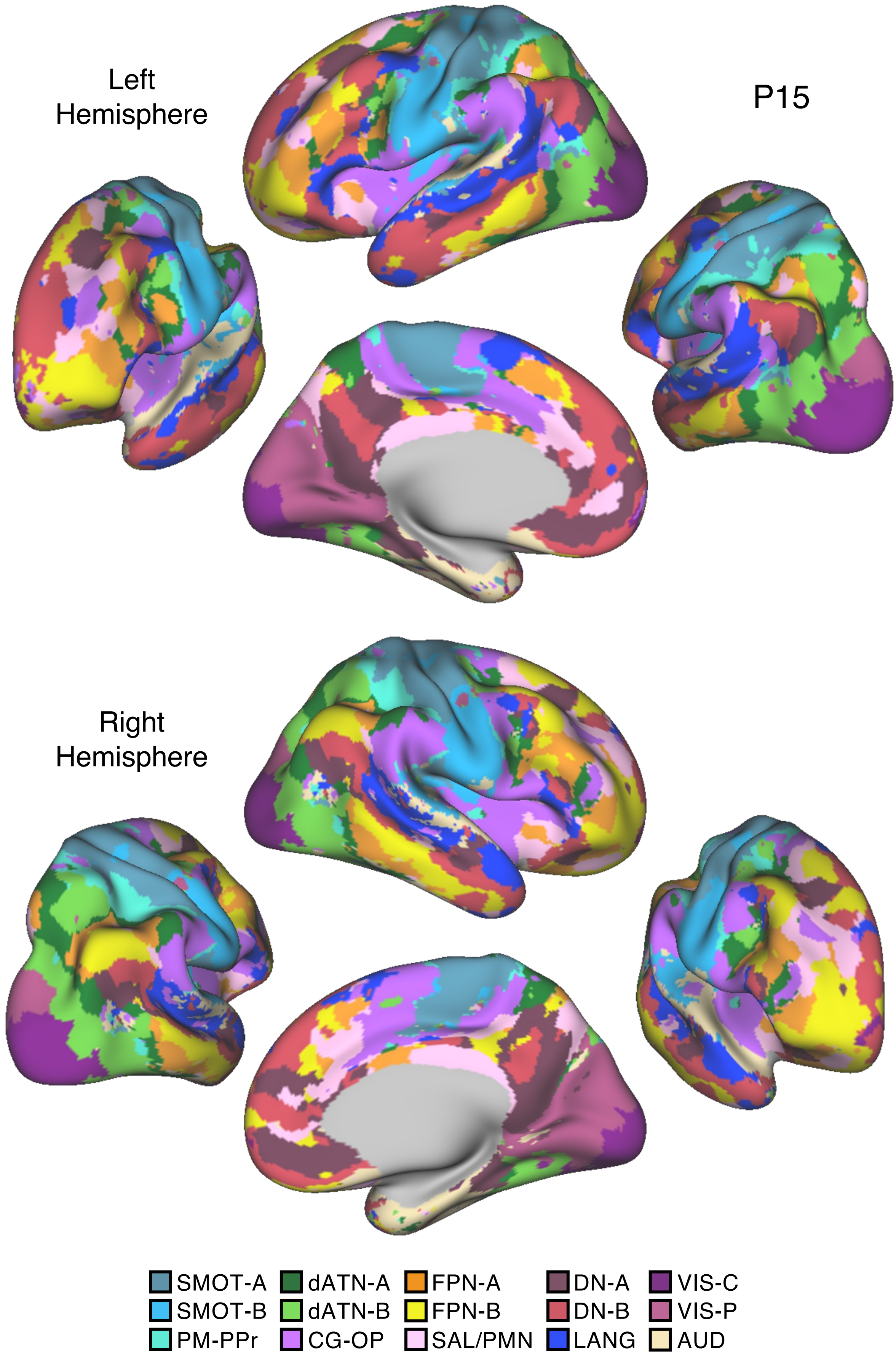

Left
Hemisphere
P15
Right
Hemisphere
SMOT-A
SMOT-B
PM-PPr
dATN-A
dATN-B
CG-OP
FPN-A
FPN-B
SAL/PMN
DN-A
DN-B
LANG
VIS-C
VIS-P
AUD


Figure S105

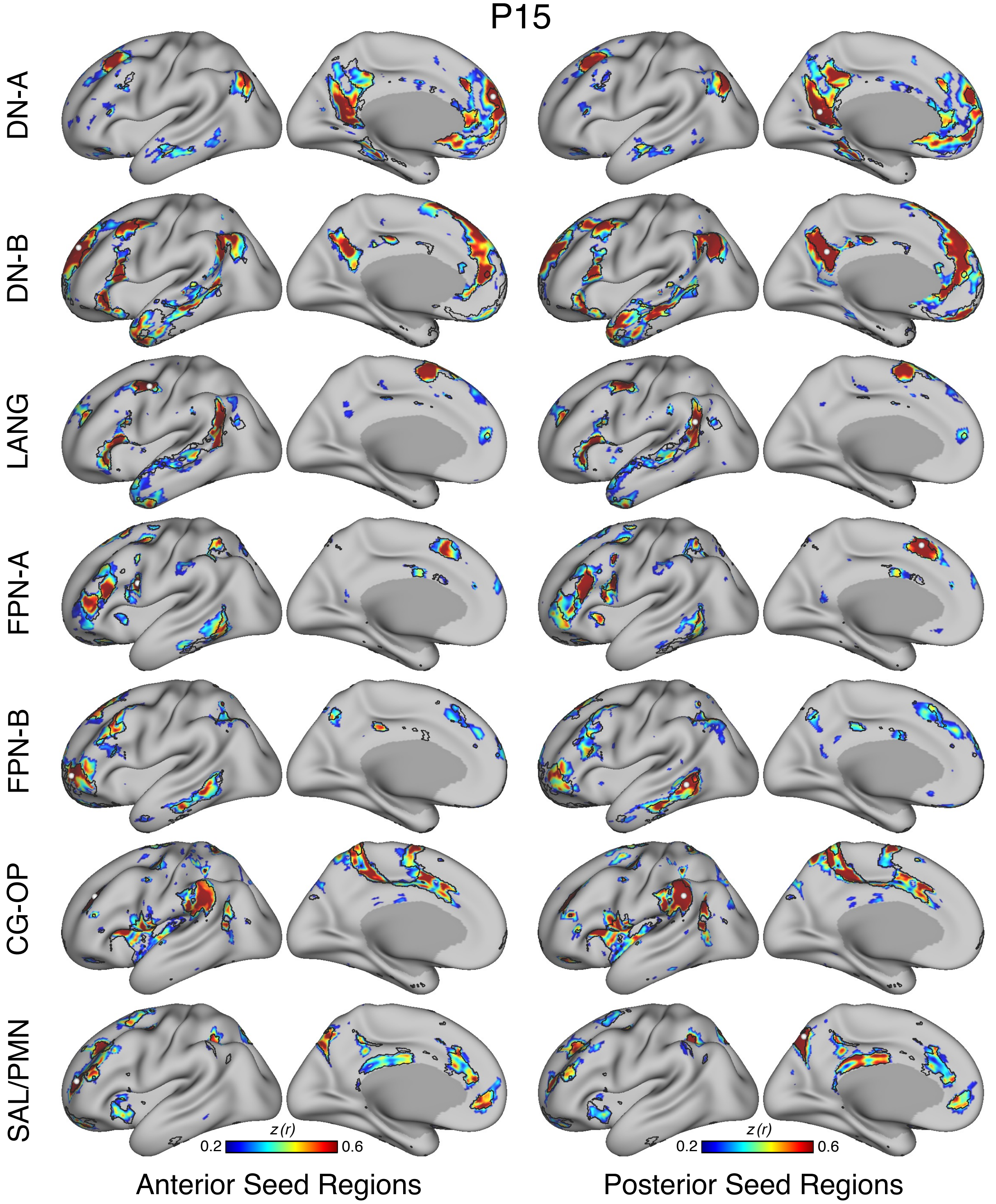
P15
DN-A
DN-B
LANG
FPN-A
FPN-B
CG-OP
SAL/PMN
z (r)
0.2
0.6
z (r)
0.2
0.6
Anterior Seed Regions
Posterior Seed Regions

Figure S106

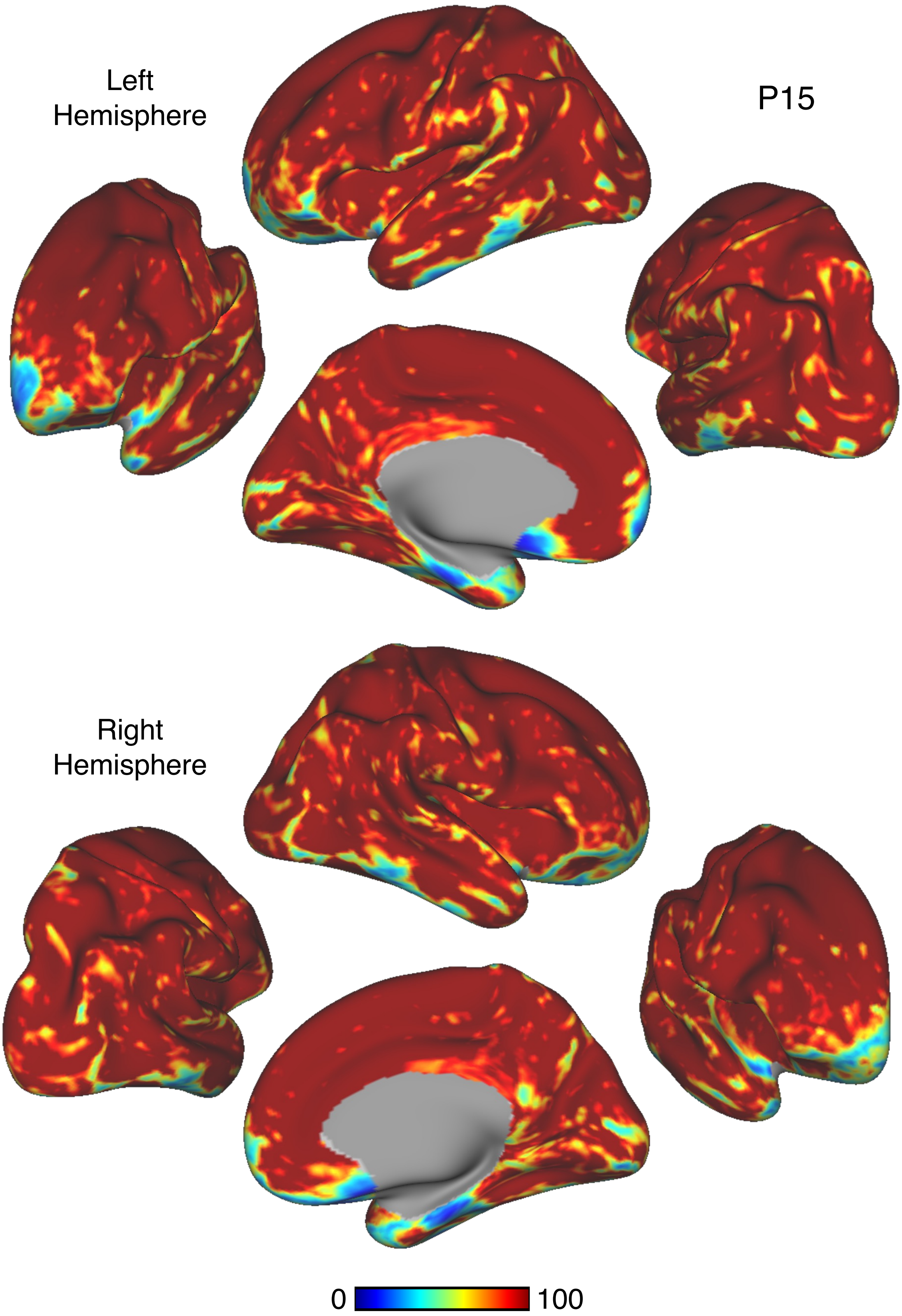


Figure S107

P15

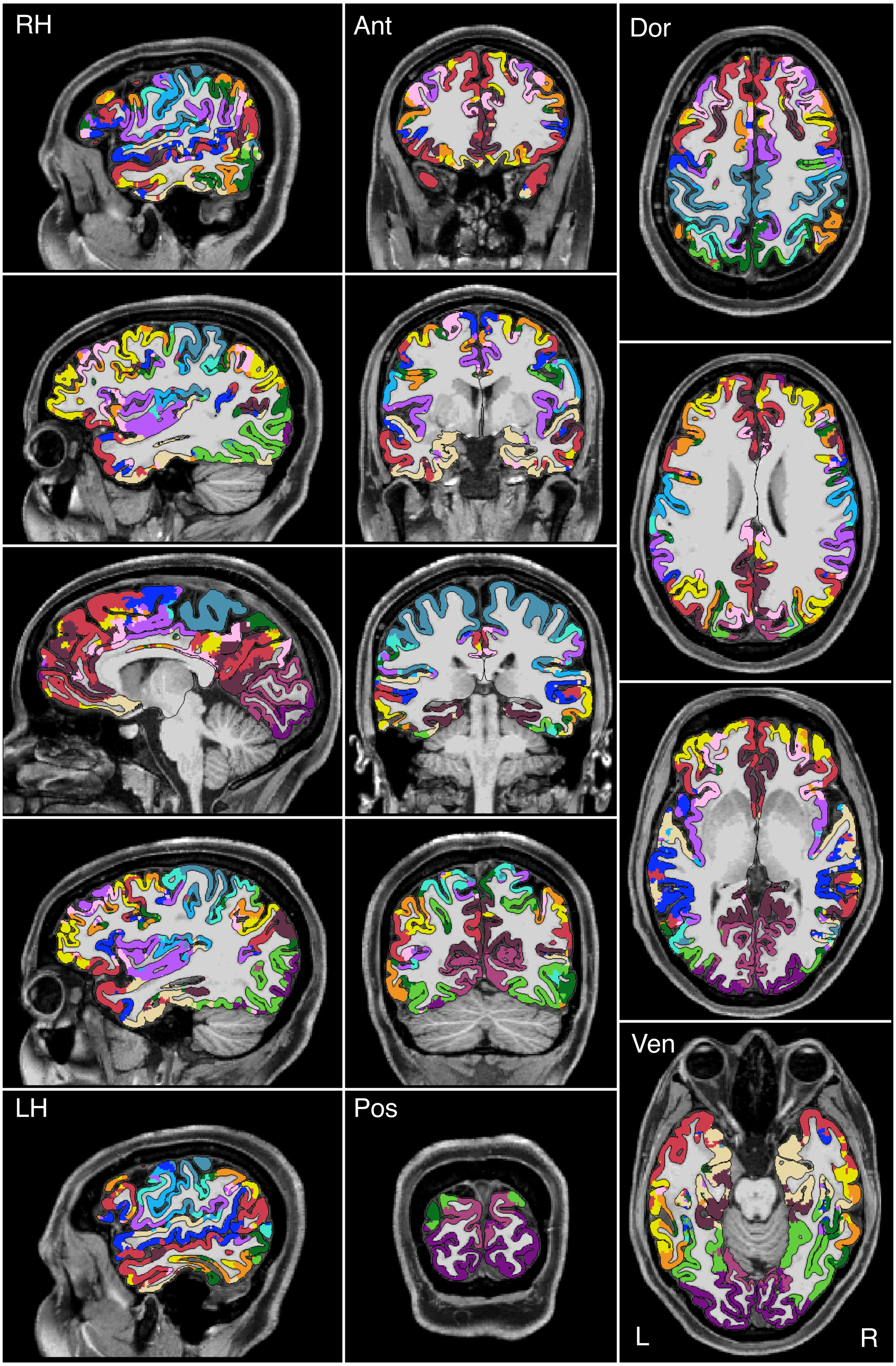


Figure S108

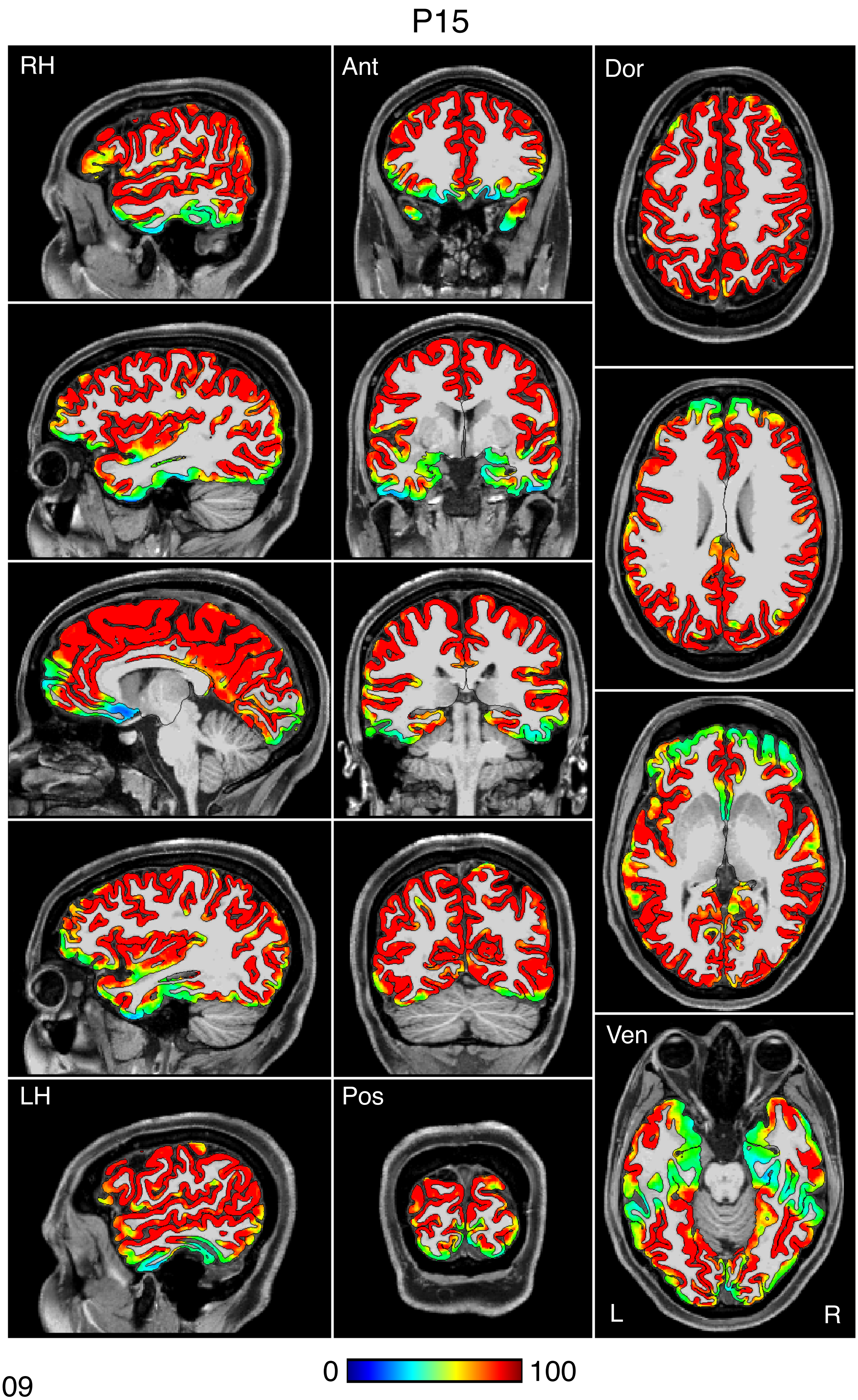


Figure S109

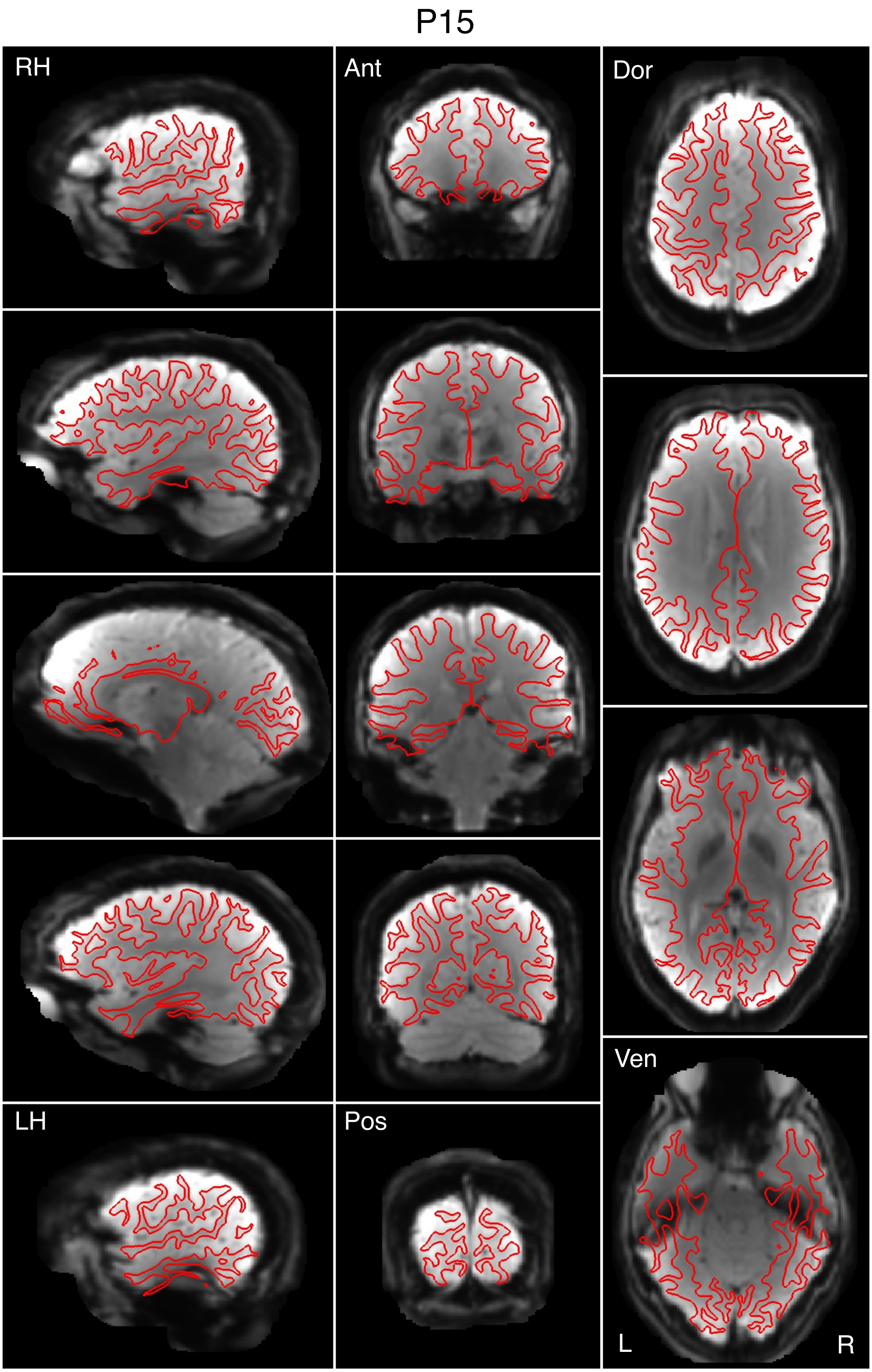


Figure S110
